# The VE-cadherin/AmotL2 mechanosensory pathway suppresses aortic inflammation and the formation of abdominal aortic aneurysms

**DOI:** 10.1101/2021.05.07.443138

**Authors:** Yuanyuan Zhang, Evelyn Hutterer, Sara Hultin, Otto Bergman, Maria J. Forteza, Zorana Andonovic, Daniel F.J. Ketelhuth, Joy Roy, Per Eriksson, Lars Holmgren

**Affiliations:** Department of Oncology-Pathology, BioClinicum, Karolinska Institutet, Stockholm, Sweden; Department of Medicine Solna, BioClinicum, Karolinska Institutet, Karolinska University Hospital, Stockholm, Sweden; Department of Cardiovascular and Renal Research, Institutet of Molecular Medicine, Univ. of Southern Denmark, Odense, Denmark; Department of Molecular Medicine and Surgery, Karolinska Institutet, Karolinska University Hospital, Stockholm, Sweden

**Author notes:** To whom correspondence should be addressed. Lars Holmgren.

## Abstract

Arterial endothelial cells (ECs) have the ability to respond to mechanical forces exerted by fluid shear stress. This response is of importance, as it is protective against vascular diseases such as atherosclerosis and aortic aneurysms. Mechanical forces are transmitted at the sites of adhesion to the basal membrane as well as cell-cell junctions where protein complexes connect to the cellular cytoskeleton to relay force into the cell. Here we present a novel protein complex that connects junctional VE-cadherin and radial actin filaments to the LINC complex in the nuclear membrane. We show that the scaffold protein AmotL2 is essential for the formation of radial actin filaments and the flow-induced alignment of aortic and arterial ECs. The deletion of endothelial AmotL2 alters nuclear shape as well as subcellular positioning. Molecular analysis shows that VE-cadherin is mechanically associated with the nuclear membrane via binding to AmotL2 and Actin. Furthermore, the deletion of AmotL2 in ECs provokes a pro-inflammatory response and abdominal aortic aneurysms (AAA) in the aorta of mice on a normal diet. Remarkably, transcriptome analysis of AAA samples from human patients revealed a negative correlation between AmotL2 expression and aneurysm diameters, as well as a positive correlation between AmotL2 and YAP expression. These findings provide a conceptual framework regarding how mechanotransduction in the junctions is coupled with vascular disease.

## Introduction

The blood vessel wall is lined with a thin layer of vascular endothelial cells (ECs), which form a barrier between the blood and tissues. These cells differ in biochemical characteristics depending on their localization in arteries or veins, as well as on the organ in which they reside. The endothelium is continuously exposed to the shear stress exerted by the blood flow. Understanding how ECs respond to shear stress is of importance as it has implications for the development of vascular diseases. Indeed, since the 1870’s it has been postulated that mechanical stress exerted on the blood vessel wall may be a trigger of atherosclerosis. Also, low wall shear stress has been associated with abdominal aortic aneurysm (AAA) rupture(1). AAA is characterized by localized medial and adventitial inflammation and dilatation of the abdominal aorta and is prevalent in men over 65 with significant morbidity and mortality(2). By contrast, areas of laminar flow appeared relatively protected against the development of the inflammatory disease(3).

To explore how the mechanotransductive pathways mediate protection or activation of the vascular disease is of clear importance(4-18). To date, several mechanosensory pathways have been identified that relay external mechanical forces to the endothelial lining(19). *In vitro*, flow-induced endothelial alignment is dependent on the activation of GTPases and consequent actin reorganization. *In vivo*, it has been shown that endothelial junctional protein complexes including PECAM-1, VE-cadherin, and VEGFR2 play an important role in the adaptive response to shear stress(18). However, it is still not clear how mechanical forces exerted by the blood flow are transmitted from the junctions via the cytoskeleton into the cell.

Studies of the (Angiomotin) Amot protein family may provide important insights into this aspect. This is a family of scaffold proteins that link membrane receptors to the actin cytoskeleton, polarity proteins, and are implicated in modulating the Hippo pathway(20-26). We have recently shown that one of its members, AmotL2 (p100 isoform), is associated with the VE-cadherin complex in ECs and E-cadherin in epithelial cells(25, 27). Silencing of AmotL2 in zebrafish, mouse or cells *in vitro* results in a loss of radial actin filaments that run perpendicular to the outer cell membrane. These actin filaments mechanically connect cells via binding to junctional cadherins, and thereby transmit force. Conditional silencing of AmotL2 in the endothelial lineage of mice inhibits expansion of the aorta during the onset of circulation resulting in death *in utero* at embryonic day 10(25). In this report, we have analyzed the role of AmotL2 in controlling junctional and cytoskeletal components during the alignment of arterial ECs exposed to laminar flow. Here we present a novel mechanical transduction pathway active in arteries that is protective against vascular inflammation as well as the formation of AAAs.

## Results

### Amotl2 is essential for arterial alignment

We analyzed the expression pattern of AmotL2 in mouse descending aorta (DA) and the inferior vena cava (IVC) as shown in **Fig. 1a**,**b**. ECs of the DA were typically elongated and aligned with the direction of blood circulation and contained radial actin filaments that were connected to the cellular junctions (**Fig. 1c**). AmotL2 localized to EC junctions as previously reported(25). In contrast, ECs of the IVC exhibited a more rounded cellular shape with no or few detectable radial actin filaments as well as a significantly lower expression level of AmotL2. Box plots in **Fig. 1d-f** represents the statistically significant difference between the DA and IVC with regards to AmotL2 expression, cellular shape and the presence of radial actin filaments. In addition, we mapped AmotL2 expression in mouse retinal vasculature at different ages (**Supplementary Fig. 1**). AmotL2 was expressed at similar levels in arteries and veins of post-natal day 6 mice, whereas in mature mice (three months) Amotl2 was primarily expressed in retinal arteries. Interestingly, AmotL2 was barely detectable in blood vessels of mice older than 15 months.

**Figure 1.**
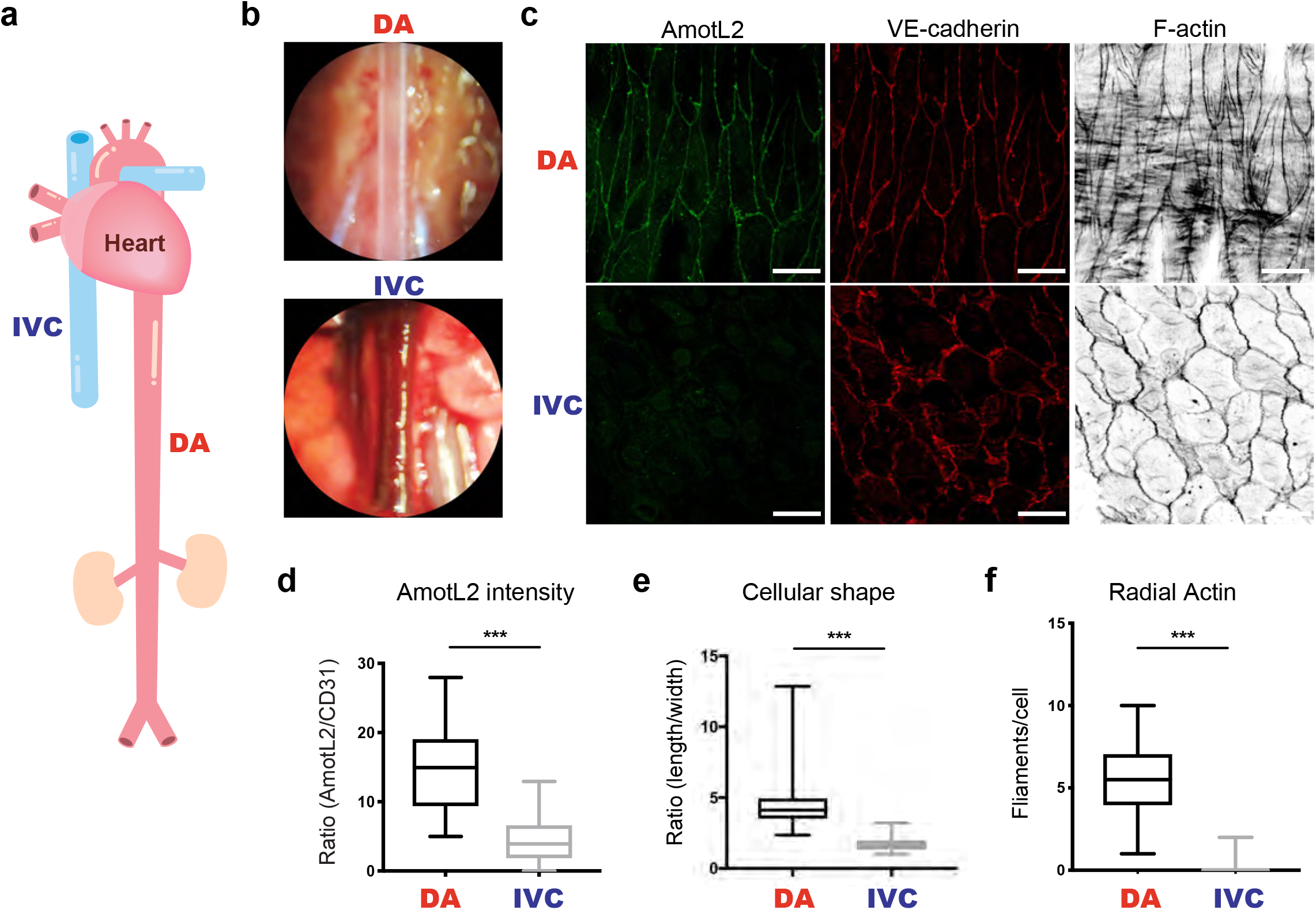
AmotL2 is primarily expressed in DA rather than IVC. **a**, Schematic of major blood vessels connected to the heart. DA indicates descending aorta leaving from the heart and IVC is for inferior vena cava. **b**, Anatomical images of the DA and IVC under microscopy during dissection process. DA image was taken after PFA perfusion. **c**, Representative images of DA and IVC used for whole-mount staining. Both vessels were stained with immunoaffinity purified AmotL2 antibodies (in green), VE-cadherin (in red) and phalloidin (in gray scale). **d**, Graph of box plot showing quantification of fluorescence intensity of AmotL2 in co-localization with CD31 in *amotl2*^ec+/ec+^ DA, using ImageJ. **e**, Box plot depicting the difference of cell length/width ratio in *amotl2*^ec+/ec+^ DA and IVC. **f**, Quantification of macroscopic radial actin filaments per EC in the DA and IVC. At least 4 images per mouse (n>3) were analyzed for each group. ^***^*P*<0.001. Scale bars, 20 µm.

We have previously shown that AmotL2 is required for the formation of radial actin filaments both in epithelial and endothelial cells(25, 27). The preferential expression of AmotL2 in EC of the aorta raised the possibility that AmotL2 controlled arterial EC shape via the formation of radial actin fibers. To address this question, we used a genetic deletion approach to silence *amotl2* gene expression specifically in the endothelial lineage as previously reported(28). In this model system, *amotl2*^flox/flox^ mice were crossed with Cdh5(PAC)^CreERT2^ transgenics as well as ROSA26-EYFP reporter mice(25), from now on referred to as *amotl2*^*ec/ec*^. This crossing enables efficient inducible conditional recombinase expression and subsequent *amotl2* knockout in ECs (*amotl2*^*ec-/ec-*^) after tamoxifen injections as well as quantification of recombination efficiency by YFP expression (**Supplementary Fig. 2a**,**b**). Adult mice (7-9 months old) were euthanized one month after tamoxifen injections, and aortae were dissected and analyzed by whole-mount immunostaining. Inactivation of AmotL2 in the DA resulted in the loss of radial actin filaments and altered cell shape (**Fig. 2a**, quantification in **2b**,**c**). No detectable changes in the actin cytoskeleton and cellular shape were observed in the endothelium of the IVC of *amotl2*^ec-/ec-^ mice. This observed phenotype change appeared to be arterial-specific as similar effects were observed in arterial, but not venous ECs of other organs such as the urinary bladder (**Fig. 2d**, quantification in **2e**,**f**).

**Figure 2.**
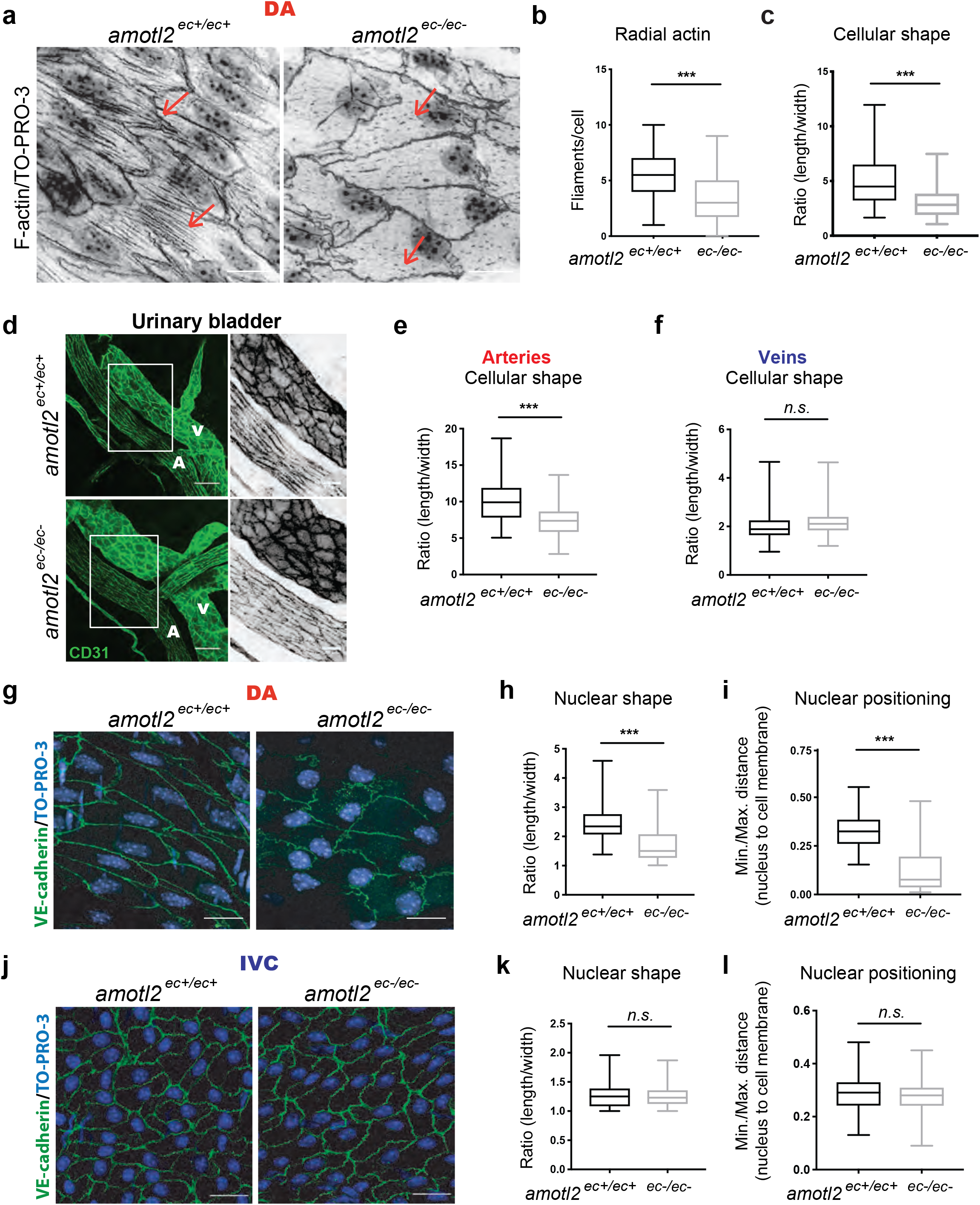
Deletion of AmotL2 resulted in aortic endothelial cellular and nuclear shape change. **a**, Whole-mount IF staining of *amotl2*^ec+/ec+^ and *amotl2*^ec-/ec-^ DA with phalloidin and TO-PRO-3 showing F-actin and nuclei (in gray scale), respectively. **b**, Box plot showing quantification of radial actin filaments per EC in DAs from *amotl2*^ec+/ec+^ and *amotl2*^ec-/ec-^ mice (n=6 in each group). **c**, Quantification of cell length/width ratio in *amotl2*^ec+/ec+^ and *amotl2*^ec-/ec-^ DAs (n=6 in each group). **d**. IF staining of bladder vasculature in *amotl2*^ec+/ec+^ (n=6) and *amotl2*^ec-/ec-^(n=6) mice, with PECAM-1 indicating ECs (in green). Quantification of EC shape in bladder arteries and veins were presented in **e** and **f**, respectively. Boxed area in the left panel was magnified in the middle and right panels. White character V and A were abbreviations of vein and artery respectively. **g**, Whole-mount IF staining of *amotl2*^ec+/ec+^ and *amotl2*^ec-/ec-^ DAs with VE-cadherin (in green) and TO-PRO-3 showing nuclei (in blue,). **h**, Quantification of nuclear length/width ratio in *amotl2*^*ec+/ec+*^ and *amotl2*^*ec-/ec-*^ DAs (n ≥ 6 in each group). Along with the long axis of the EC, the closest distance between nuclei to one end of the cell was measured and normalized to the full cell length. The ratio was depicted in box plot **i. j**, Representative images on IVC of *amotl2*^ec+/ec+^ and *amotl2*^ec-/ec-^ mice with VE-cadherin (in green) and TO-PRO-3 (in blue). Quantification of nuclear shape and nuclear positioning of ECs in IVC were presented in **k** and **l** (n=5 in each group). *n*.*s*., not significant. ^***^*P*<0.001. Scale bars: (**a**), (**g**) and (**j**) are 10 µm, (**d**) is 20 µm.

The nucleus is the largest organelle of the EC. As such it is exposed to the hemodynamic drag by the blood flow. In response to shear stress, EC as well as EC nuclei elongate, and nuclei orient themselves relative to the direction of flow. Nuclear positioning as well as alignment has previously been shown to be dependent on the association to microfilaments as well as the tubulin network(29). In *amotl2*^ec+/ec+^ mice, nuclei of ECs of the DA were elongated and orientated in parallel with cell alignment in the direction of blood flow. However, in AmotL2 deficient ECs, the nuclei were more rounded with irregular shapes and positioned close to the cell edge downstream of the flow direction (**Fig. 2g**, quantification in **2h**,**i**). These changes in nuclear shape and positioning were again not observed in the IVC (**Fig. 2j**, quantification in **2k**,**l**). Taken together, these data showed that AmotL2 is required for EC elongation as well as positioning of the EC nucleus.

### AmotL2 expression is required for arterial response to flow

Next we investigated whether AmotL2 is required for arterial endothelial compliance to laminar flow *in vitro*. For this purpose, we used a short hair-pin Lentiviral approach to deplete AmotL2 in Human Aortic ECs (HAoEC, **Fig. 3a**). No differences between control and AmotL2 depleted cells in cellular and nuclear shape were detectable under static conditions (**Fig. 3b-d**). To recapitulate arterial flow conditions, cells were exposed to 14dynes/cm^2^ for 48h in a flow chamber as described in materials and methods. Control HAoECs exhibited an elongated phenotype and aligned in the direction of flow; however, the depletion of AmotL2 resulted in failure to elongate and align in the direction of flow (**Fig. 3b**, quantification in **3e**,**f**). We could further show that AmotL2 was required for controlling nuclear shape and positioning. Consistent with the cellular shape change, nuclei also exhibited a rounder shape and could not maintain the positioning at the center of the cells when compared to control HAoECs (**Fig. 3g,h**).

**Figure 3.**
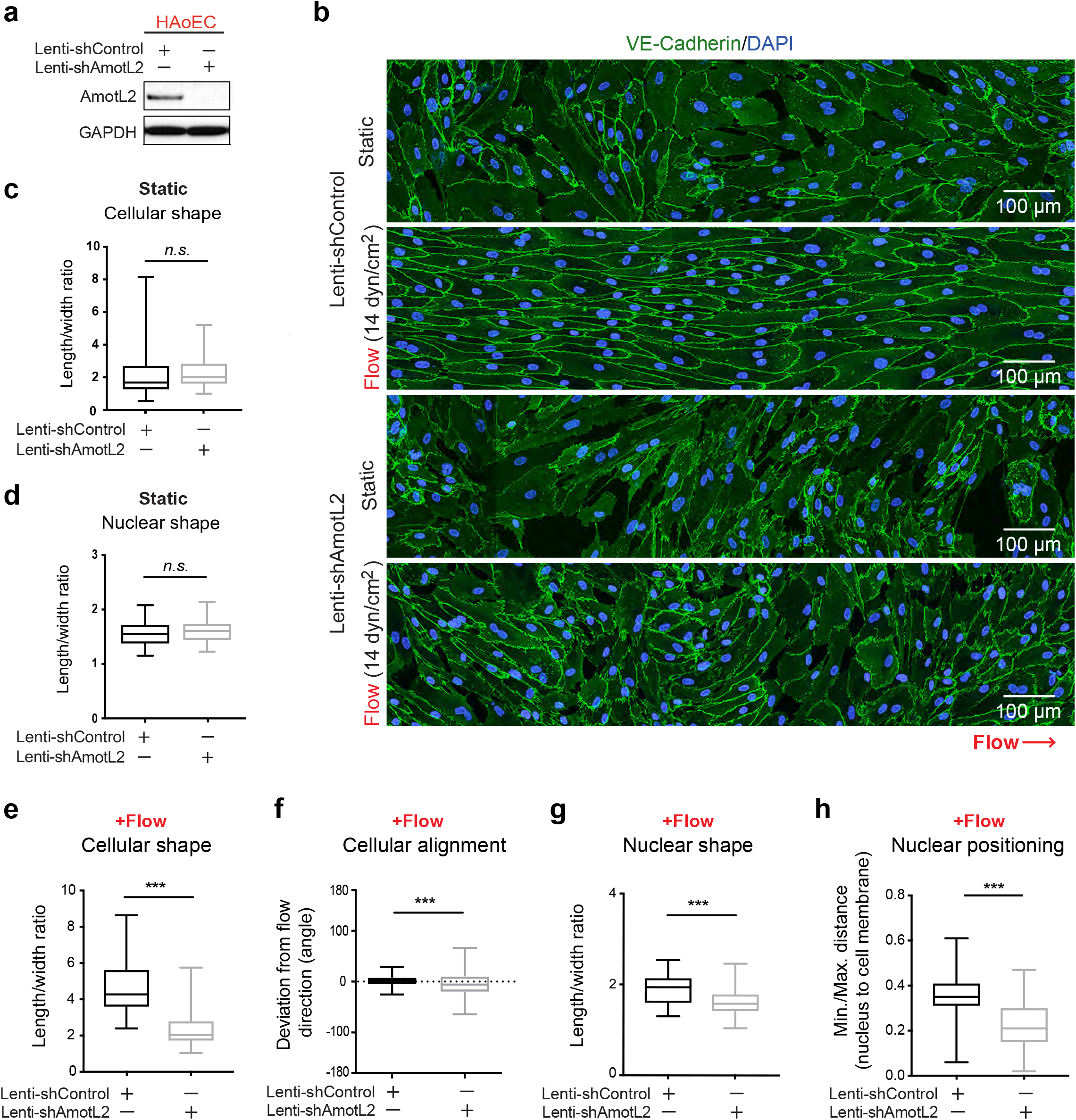
Effect of AmotL2 knockdown on HAoEC *in vitro*. **a**, WB analysis of whole cell lysates of confluent HAoEC cells treated with scrambled control or AmotL2 shRNA Lentivirus probed for AmotL2 and GAPDH. **b**, IF staining for VE-cadherin (in green) and DAPI (in blue) of HAoEC cells infected with scrambled control or AmotL2 shRNA Lentivirus under static or flow (14 dyn/cm^2^) conditions. Flow direction is indicated below the images. **c**, Quantification of cellular length/width ratio and **d**, nuclear length/width ratio in scrambled control and AmotL2 shRNA Lentivirus of HAoEC cells under static conditions. Representative quantification analysis of cell length/width ratio (**e**, n>80 cells), cell alignment (**f**, n>250 cells), nuclear length/width ratio (**g**, n>60) and nuclear positioning (**h**, n>55 cells) were performed in both control and KD HAoEC cells. Scale bars: (**b)** 100 µm. *n*.*s*., not significant. ^***^*P*<0.001.

### AmotL2 couples VE-cadherin to the nuclear lamina

Actin filaments are coupled to the nuclear membrane through the LINC complex(29-32). This complex consisting of Sun-domain proteins (SUN-1 and -2) and KASH domain proteins (Nesprin-2) connects to Lamin A/C of the nuclear lamina. Therefore, we next investigated a possible connection between VE-cadherin, AmotL2, actin and the LINC complex.

We used a Co-IP approach to obtain AmotL2 associated immunocomplexes from both murine endothelial cell line (MS1) and primary Bovine Aortic Endothelial (BAE) cells. By spectrometry analysis (raw data in **Supplementary Table 1**,**2**), we identified cellular membrane protein VE-cadherin and α-, β-, δ-catenin, as well as the nuclear laminal proteins such as SUN2 and Lamin A, in all triplicate IP samples from MS1 cells (**Fig. 4a)**. Consistently, the majority of key membrane proteins were recognized in BAE AmotL2 immunoprecipitates **(Supplementary Fig. 3a**). Mouse lungs consist of approximately 10-20% ECs. We performed immunoprecipitation using AmotL2 antibodies and could verify that Amotl2 associates with VE-cadherin, B-catenin, actin, and Lamin A also in the murine lung tissue *in vivo* (**Fig. 4b**). It was not possible to analyze Nesprin-2 by western blot due to its high molecular mass (<800 kDa). The association of AmotL2 was dependent on myosin contraction and actin filaments as treatment with the myosin II inhibitor blebbistatin or the actin polymerization inhibitor cytochalasin D both decreased the association of AmotL2 to Lamin A/C and SUN2 (**Fig. 4c**). In addition, immunoprecipitation of VE-cadherin confirmed the association to AmotL2 and Lamin A in HAoECs (**Fig. 4d**). Interestingly, shRNA depletion of AmotL2 protein in ECs abrogated the association of VE-cadherin to both actin and Lamin A (**Fig. 4d**).

**Figure 4.**
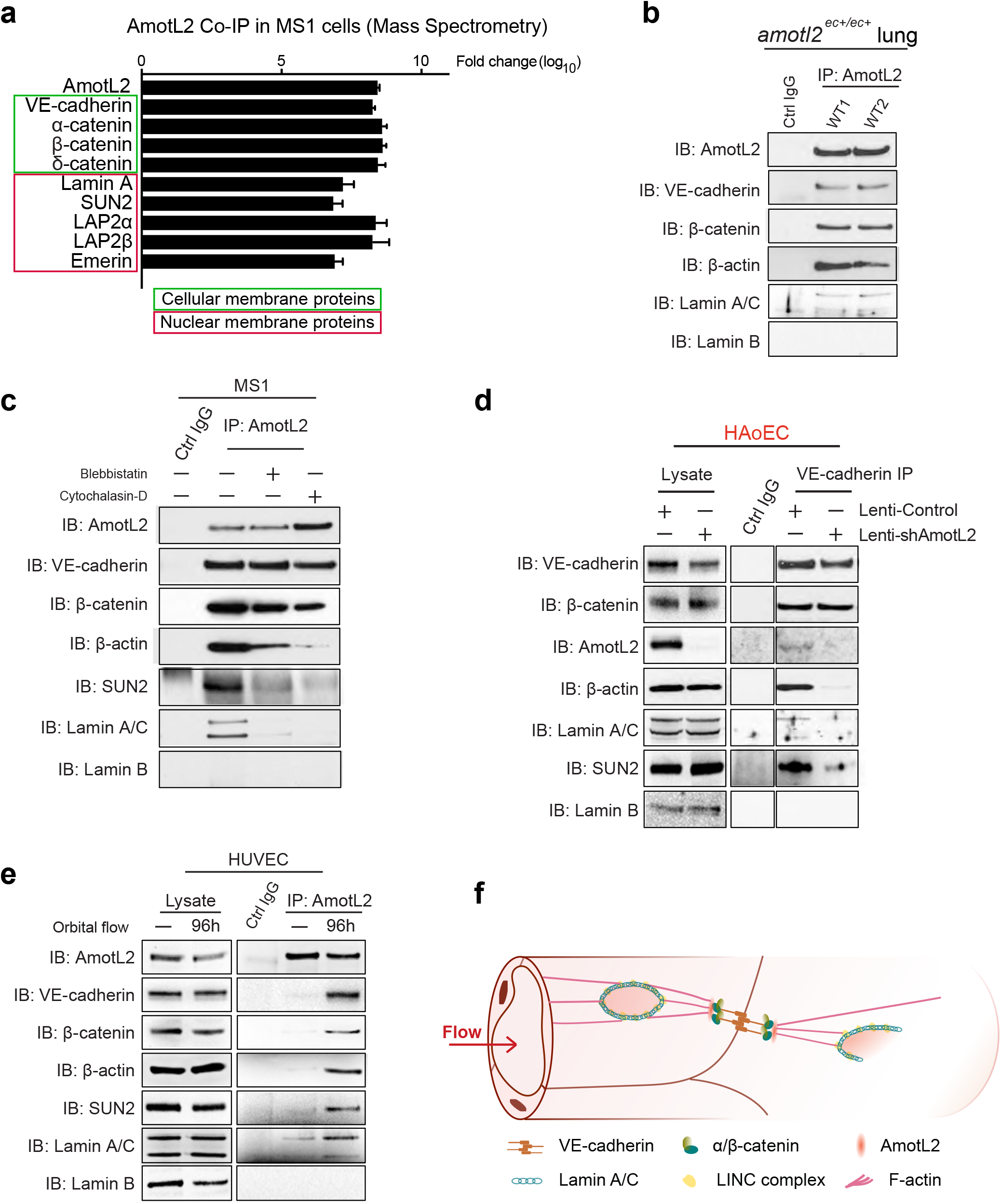
AmotL2 links VE-cadherin to nuclear membrane protein Lamin A/C through F-actin. **a**, AmotL2, together with cellular membrane proteins (framed in green box) and nuclear membrane proteins (framed in red box) were identified from AmotL2 immunoprecipitates from MS1 cells analyzed by MS. The data was displayed with fold change (log_10_) as compared to control immunoprecipitate samples. **b**, Mouse lung tissues from two randomly picked *amotl2*^ec+/ec+^ mice were lysed, subjected to IP with rabbit IgG or AmotL2 antibody and analyzed by WB. **c**, Murine endothelial cells (MS1) either treated with 25µM blebbistatin for 2h or 1µM cytochalasin D for 1h. Cell immunoprecipitates with rabbit IgG or Amotl2 antibody were analyzed by WB. **d**, Immunoblotting (IB) of VE-cadherin associated protein complexes in HAoEC with either Lenti-Control or Lenti-shAmotL2. Whole lysate harvested from those cells were used as positive input. **e**, WB analysis of AmotL2 IP samples in HUVECs, either under the static condition or 96h after orbital shaking (300 rpm). Only the cells that grew at the periphery (6cm to the edge) of a 15cm dish were harvested. **f**, Hypothetical schematic of AmotL2 associated mechano-responsive complex forming under flow condition.

The higher expression levels of AmotL2 detected in the DA as compared to the IVC *in vivo* raised the question of whether AmotL2 associated complex is shear stress responsive. We placed primary Human Umbilical Venous ECs (HUVECs) in a 15cm culture dish on an orbital shaker to apply the circulatory flow to the cells (schematic in **Supplementary Fig. 3b**). As previously described(33), the alignment of the cells at the periphery area was observed after 96h (**Supplementary Fig. 3c,d**). Analyses of the AmotL2 IP revealed a dramatically increased binding with VE-cadherin, actin and nuclear membrane proteins compared to the static condition (**Fig. 4e**).

Taken together, MS and IP data suggest a model where VE-cadherin/AmotL2 forms a complex that is connected to the nuclear LINC complex via actin filaments (schematic in **Fig.4f**).

### Deletion of AmotL2 promotes vascular inflammation

EC alignment and cytoskeletal reorganization in response to laminar blood flow is protective against inflammation(3). The change in the mechano-response of *amotl2*^ec-/ec^ EC raised the question whether this was accompanied by a pro-inflammatory phenotype. The aortic arch is exposed to turbulent blood flow and the descending aorta is exposed to laminar flow. For this reason, we analyzed these areas of the aorta separately (**Fig. 5a**). mRNA isolated from DAs in both *amotl2*^ec+/ec+^ (n=3) and *amotl2*^ec-/ec-^ (n=5) mice were analyzed by mRNA-seq. Due to high variability between individual mice, only 65 genes were identified to be differentially expressed (*adjusted p value<0*.*05*) between *amotl2*^*ec+/ec+*^ and *amotl2*^*ec- /ec-*^groups (**Supplementary Table 3**). However, those genes were obviously enriched to immuno-related GO terms, such as “Neutrophil activation involved in immune response”, “inflammatory response” and “regulation of immune effector process” (**Fig. 5b**). Gene expression profile analysis on MS1 cells *in vitro* treated with control or AmotL2 siRNA was also performed by RNA-seq. Interestingly, we did not observe any up-regulation of inflammatory gene signatures *in vitro* (**Supplementary Fig. 4a** and **Supplementary Table 4**), indicating that the *in vivo* environment was required for this to occur. We therefore performed whole-mount immunostaining to analyze the presence of CD45^+^ inflammatory cells in the aortic arch. The whole mount immunofluorescence staining showed the presence of spindle-like CD45^+^ cells in the inner curvature of the aortic arch in wild-type animals and in arterial bifurcations (**Fig. 5c,d**). Analyses of the *amotl2*^ec-/ec-^ mice 1-month post-injection of tamoxifen revealed the infiltration of CD45^+^ round monocytes in the outer curvature of the aortic arch (**Fig. 5c,d**). The CD45^+^ monocyte-like cells were detected in the sub-endothelial compartment of the aorta (**Fig. 5e,f**). In total, 45% (9/20) *amotl2*^ec-/ec-^ arches contained more than one CD45^+^ lesion.

**Figure 5.**
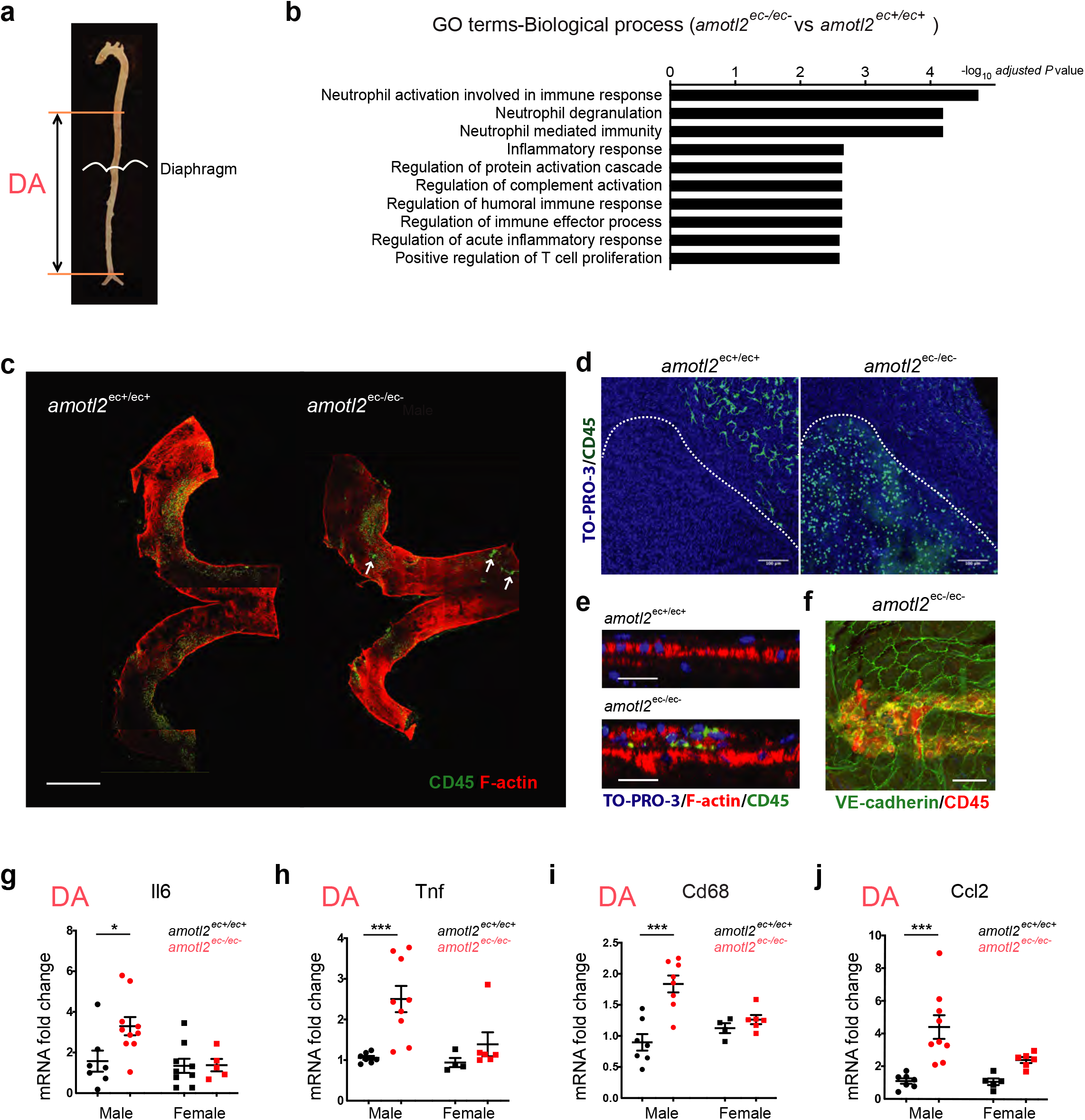
Absence of AmotL2 resulted in pro-inflammatory phenotype of aorta. **a**, Schematic of the full-length aorta. mRNA was isolated from “DA” indicating descending aorta (including part of thoracic aorta and whole abdominal aorta). **b**, Enriched GO terms (Biological process 2018) analyzed by Enrichr (**Supplementary Table 3**) and presented in the graph ranking by -log_10_ *adjusted P* value. mRNA isolated from DAs of *amotl2*^ec+/ec+^ (n=3) and *amotl2*^ec-/ec-^ (n=5) mice were sent for RNA-sequencing analysis. 65 significant genes which were differentially expressed were subjected to GO terms matching. **c**, Whole-mount staining on aortic arch of *amotl2*^ec+/ec+^ and *amotl2*^ec-/ec-^ mice, stained with CD45 (in green) and phalloidin (in red). White arrows are pointing at the cluster of CD45 positive cells, which indicates the endothelial lesions. **d**, Representative images of CD45 positive cell clusters (in green) in *amotl2*^ec-/ec-^ aortic arches within the area outlined by white dash line, which were not present in the *amotl2*^ec+/ec+^ arch. **e**, Orthogonal view of *amotl2*^ec+/ec+^ arch and *amotl2*^ec-/ec-^ lesion area stained with CD45 (in green), phalloidin (in red) and TO-PRO-3 (in blue). The luminal cells are on the top layer in the images. **f**, 3D-projection view of CD45+ (in red) cells invading in the endothelium (in green) in *amotl2*^ec-/ec-^ arch. The image was processed in ImageJ. **g-j**, The relative mRNA expression levels of *Il6, Tnf, Cd68*, and *Ccl2* were determined with TaqMan probes. The mRNA was isolated from DA (**g-j**) tissues of 13 *amotl2*^ec+/ec+^ mice (in black dots, male n=7, female n=4-9) and 15 *amotl2*^ec-/ec-^ mice (in red dots, male n=8-10 and female n=5-6). Relative expression levels were normalized versus *Hprt*. Fold-changes are quantified, and data shown are mean ± S.E. Scale bars: **(c)** 1000 µm, **(d)** 100 µm, and **(e)** 20 µm. ^*^*P*<0.05. ^***^*P*<0.001.

### Inflammatory markers are up-regulated in *amotl2*^*ec-/ec-*^ mouse aortae

Next, we quantified mRNA expression of inflammatory markers in DAs. The mRNA expression level analyzed by TaqMan qRT-PCR showed a significant increase of inflammatory markers, such as *Il6, Tnf*, and *Cd68a* in *amotl2*^ec-/ec-^ DA, suggesting a provocation of a general immune response. Interestingly, up-regulation of pro-inflammatory markers was more pronounced in male animals (**Fig. 5g-j**).

In response to pathogenic stimuli, expression of various leukocyte adhesion molecules can be augmented as the initial sign of vascular inflammation, such as ICAM-1 and VCAM-1. Indeed, in *amotl2*^ec-/ec^ DA, an increase in *Vcam-1* expression on transcriptional level was detected by qPCR, but the change of *Icam-1* was less obvious (**Supplementary Fig. 4b,c**), revealing the activation of endothelium of the DAs in the absence of AmotL2. Remarkably, Vcam-1 up-regulation was also more profound in male mice (**Supplementary Fig. 4b**). Once leukocytes attach and migrate into underlying intima, the process of lesion formation begins. This requires the participation of chemoattractant cytokines such as Ccl2, also known as monocyte chemoattractant protein-1 (MCP-1), which is especially important for the recruitment of mononuclear immune cells. Compared to the *amotl2*^*ec+/ec+*^ aortic tissues, *amotl2* deletion caused an increase of *Ccl2* expression in the descending regions (**Fig. 5j**), as well as *Cxcl10*, which is able to selectively attract NK cells as well as T and B lymphocytes. (**Supplementary Fig. 4d**). However, other chemokines such as *Ccl5* had no significant changes (**Supplementary Fig. 4e**), suggesting that the induced inflammatory response was mainly affecting innate immune cells.

By contrast, the T and B cell markers *Cd4, Cd8*, and *Cd19*, as adaptive immune markers, showed no differences in expression level of these markers on average was detected in the DAs with or without AmotL2 (**Supplementary Fig. 4f-h**).

In conclusion, the expression profile of inflammatory-related genes indicates that AmotL2 depletion results in an innate immune response rather than an activation of adaptive immunity. Of note is the preferential upregulation of pro-inflammatory markers in *amotl2*^*ec-/ec-*^males.

### AmotL2 depletion promotes the formation of abdominal aortic aneurysms in male mice

Aortic inflammation is a predictor of the development of arterial aneurysm(34). Inflammatory aortic aneurysms account for up to 10% of all AAA cases(35). After dissection of the aorta, we could detect the formation of abdominal aortic aneurysms (AAA, dilatation>1.5 times normal size) in the proximity of the renal arterial branch (**Fig. 6a**), but not in ascending or thoracic aortae. Interestingly, 20 % of male *amotl2*^ec-/ec-^ (5/25) mice developed an AAA; however, no aneurysm was detected in the females (0/20). Furthermore, no aneurysms were observed in the *amotl2*^ec+/ec+^ mice (36 mice, 20 male and 16 female). Imaging analysis of a typical AAA revealed damage to the endothelium as well as the vessel wall (**Fig. 6b,c**). The insult to the vessel wall was further verified by histological sectioning. The intima of the vessel wall was infiltrated by CD45+ leukocytes (**Fig. 6d**) and elastin fibers were degraded (**Fig. 6e**).

**Figure 6.**
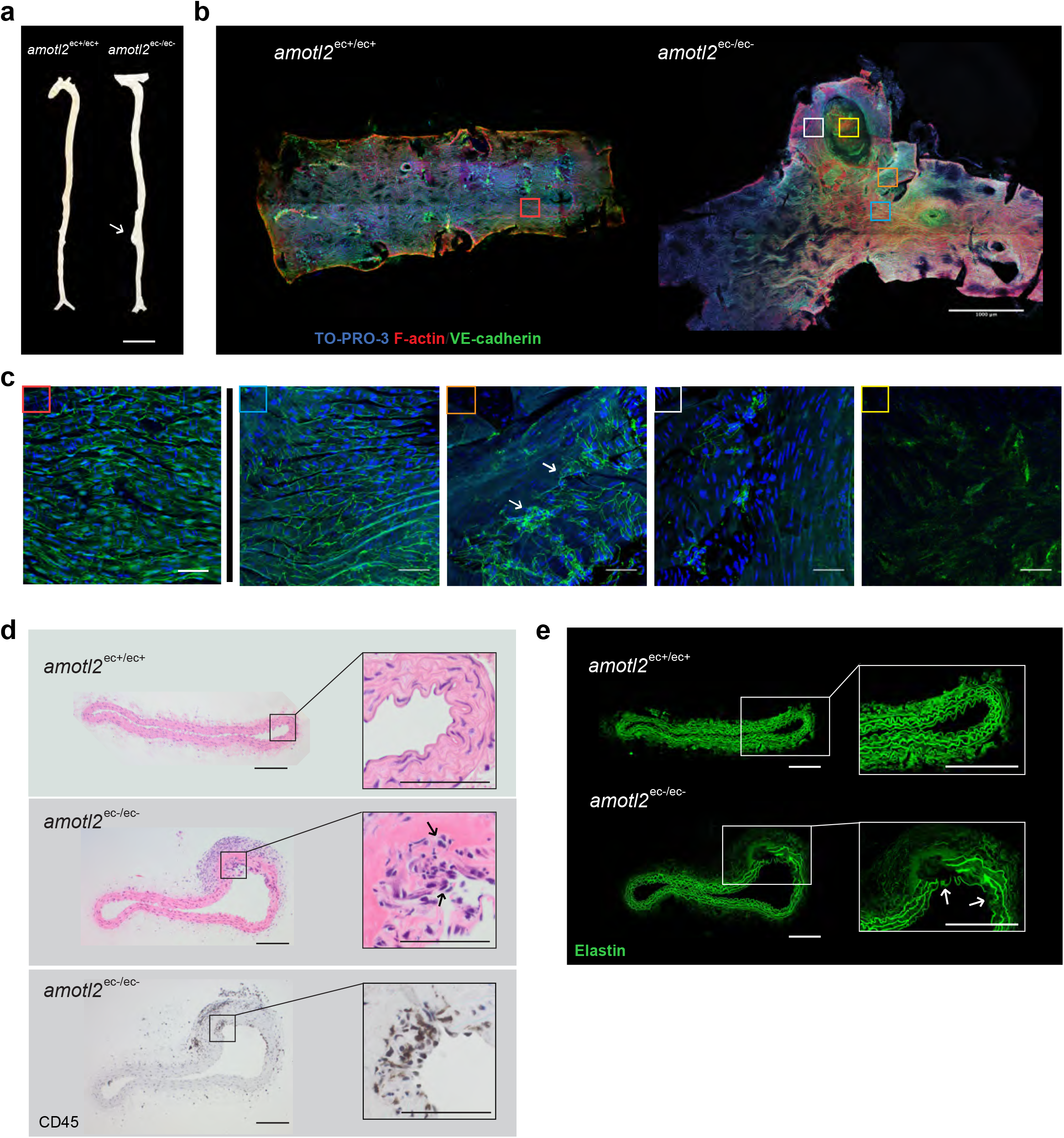
Aortic aneurysms were induced in male *amotl2*^*ec-/ec-*^ mice. **a**, The dissected full-length aorta from *amotl2*^ec+/ec+^ and *amotl2*^ec-/ec-^ mice. White arrow indicates the abdominal aortic aneurysm (AAA), which occurred close to the renal artery bifurcations. **b**, Representative images of *amotl2*^ec+/ec+^ DA and aneurysm in *amotl2*^ec-/ec-^ mice (VE-cadherin in green, phalloidin in red and TO-PRO-3 in blue). Selective area (boxed) was magnified and presented in **c**. The red box shows wild-type EC morphology in *amotl2*^ec+/ec+^ DA, while the blue box, orange box, white box, and yellow box sequentially outlined in aneurysm reveal the impact on endothelium during aneurysm formation. In orange box, endothelium was damaged and regressed, indicated by white arrows. The DAs from *amotl2*^ec+/ec+^ and *amotl2*^ec-/ec-^ mice with aneurysm were fixed and embedded by paraffin. H&E (upper and middle panel) and CD45 (lower panel) staining was applied on the paraffin sections in **d**. Additionally, aneurysm section was stained by CD45 and showed in the bottom panel. Boxed areas are magnified and placed on the right. In the aneurysm section, abnormal nuclei penetrated were observed along the lumen side indicated by black arrows. **e**, Elastin structure visualized by auto-fluorescence captured under UV light. Layers of elastin were presented in magnified images in white boxes on the right. White arrows indicate normal elastin structure in *amotl2*^ec+/ec+^ DA and discontinuation in *amotl2*^ec-/ec-^ aneurysm, respectively. Scale bars: 1000 µm (**a**), 25 µm (**c**) and 100 µm (**d** and **e**).

### AmotL2/YAP correlation in human AAA samples

Next, we assessed whether AmotL2 gene expression could be correlated to disease progression in human AAA patients. We analyzed mRNA expression in surgically resected materials from both healthy donors and patients diagnosed with AAA and undergoing aneurysm repair at the Karolinska University Hospital. mRNA samples were taken from both medial and adventitial layers of the intact aorta (13 donors) or AAA tissues (35 patients). Interestingly, AmotL2 expression level in the media tissue (containing endothelial cells, schematic in **Fig. 7a**) in AAA samples was significantly down regulated compared to the intact aortic media, which was not found in adventitia tissues (**Fig. 7b**). The expression pattern of the first exon of AmotL2 was specifically analyzed since this exon is specific for the p100-AmotL2 isoform that we have previously shown interacts with VE-cadherin(25). In line with the studies in mice, there was an inverse correlation between the expression of the first exon of AMOTL2 and the luminal/external diameter of AAA (**Fig. 7c**).

**Figure 7.**
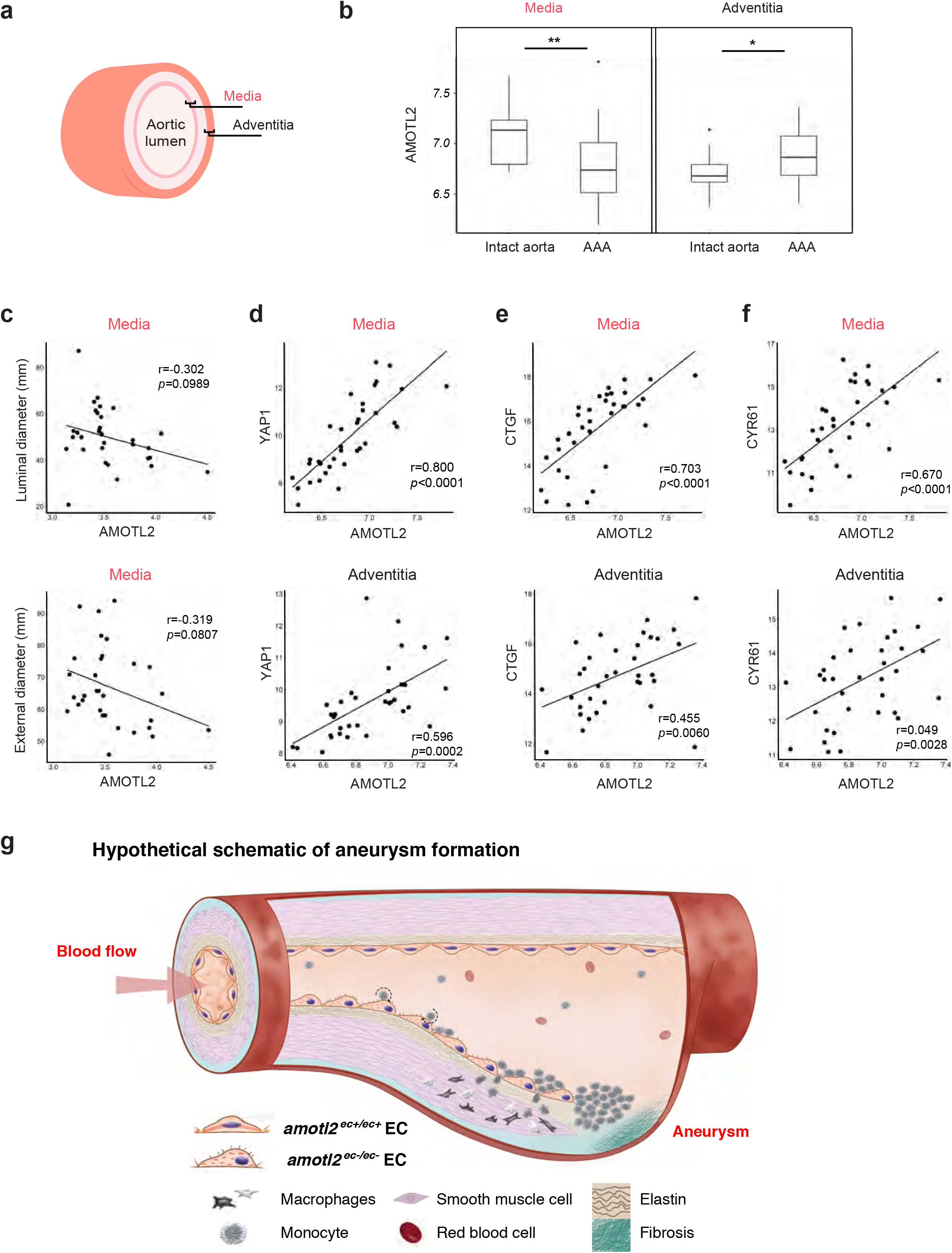
YAP1 and AMOTL2 correlates in human AAA patients. AAA samples of intima/media region of aortic wall not covered by an intraluminal thrombus i.e., aneurysm wall covered by an EC layer were obtained at open AAA surgery, and mRNA were isolated from both media and adventitia. **a**, Schematic depicting anatomical view of the media and adventitia tissue in the aorta. **b**, Quantification of AmotL2 mRNA expression in medial and adventitial tissues from both intact aortae (13 healthy donors) and dilated aortae (35 AAA patients). ^*^*P*<0.05. ^**^*P*<0.01. **c**, The correlations between AMOTL2 mRNA level of expression and the luminal (upper panel) / external (lower panel) diameter of aneurysms in AAA patients (n=35). AMOTL2 expression level was based on the expression of the first exon from 3’ end, detected by specific exon probe, which represents the full-length isoform of AMOTL2. AMOTL2 correlation with YAP1, CTGF and CYR61 of mRNA level of expression were shown in **d-f**, respectively. AMOTL2 expression level was calculated with mean value of every exon expression detected. For each correlation analysis, samples from media were present on top while samples from adventitia on bottom. Correlation coefficient *r* and *p-value* are labelled in each individual figure. 21 male and 10 female patients were analyzed in **c**, while 25 male and 10 female patients were enrolled in **d-f. g**, Hypothetical schematic of the formation of an AAA. The step-by-step formation of an AAA includes EC activation, immune cell rolling and attachment, leukocyte extravasation, macrophage differentiation, thrombus formation, and finally, lumen dilation (aneurysm).

YAP (Yes Associated Protein 1) has been reported to act as a rheostat of external mechanical force(36). Of interest is that YAP binds to the AmotL2 promoter and thus activates its transcription(37). Consistent with these findings, a significant positive correlation was detected between AMOTL2 and YAP mRNA levels (*r = 0*.*800, P < 0*.*0001*) in media tissues containing endothelium in AAA patients of both genders (**Fig. 7d**). We also found a weaker positive correlation of AMOTL2/YAP in adventitia tissues that did not contain ECs (*r=0*.*592, p=0*.*0002*). A robust correlation of the well-established YAP target genes CTGF and CYR61 with AMOTL2 was also observed, as shown in **Fig. 7e** and **7f**, respectively. C-FOS has been reported to bind to the promoter and drive expression of the p60 AMOTL2 isoform. This isoform is primarily activated by severe hypoxia in ischemic tissues(25). However, no correlation between AMOTL2 and C-FOS mRNA expression could be detected (**Supplementary Fig. 5a**). YAP and TAZ are paralogs with overlapping function as transcriptional regulators of the Hippo pathway. However, there was no significant correlation between AMOTL2 and TAZ expression, suggesting that YAP is primarily driving AMOTL2 expression in the human aortic tissues (**Supplementary Fig. 5b**). In addition, there was no correlation of AmotL2 with gender (**Supplementary Fig. 5c-g**).

## Discussion

The vascular endothelium plays an important role in the biomechanical response to hemodynamic forces. Understanding the pathways involved in this response is of importance to comprehend the pathogenesis of vascular disease. In this report we show for the first time that the cellular junctions of arterial ECs are connected via AmotL2 and microfilaments to the nuclear lamina. Interference with this pathway impairs EC alignment in response to shear stress and abrogates nuclear positioning resulting in inflammation and formation of abdominal aortic aneurysms.

We have used an inducible mouse model to target AmotL2 in the EC lineage. In our previous publication, we could show that Amotl2 silencing in ECs *in utero* resulted in impaired aortic expansion and death at embryonic day 10. Silencing of AmotL2 in adult mice, however, did not affect overall survival or had any obvious negative effects up to six months after AmotL2 depletion. The defect in adult mice was clearly more subtle as it was restricted to ECs exposed to arterial flow. We demonstrated not only that AmotL2 is required for cellular alignment in areas of shear stress, but also provided novel insights into how VE-cadherin is mechanically coupled to the cytoskeleton and thereby controls cell shape. AmotL2 triggers the formation of radial actin filaments that mediate junctional tension between neighboring cells. These radial actin filaments were detected in arterial, but not or at least at lower levels in venous endothelium. AmotL2 is a scaffold protein and as such brings together protein complexes of different functions such as Par3, MAGI-1b, Merlin, Actin, VE-cadherin. Of interest is that, in HUVECs of the venous origin, AmotL2 is sufficient to induce radial actin filaments when the cells aligned under flow condition. Our data are consistent with the notion that VE-cadherin is part of a mechanosensory complex with VEGFR2 and PECAM-1 as previously described(38). The present data show that the VE-cadherin/AmotL2 protein complex is responsible for the actual cell shape modulation in arterial ECs. These most recent investigations together with our previous findings have shown that the VE-cadherin AmotL2 complex mediates mechanical forces between ECs, suggesting that AmotL2 may also relay mechanical forces not only from circulatory shear stress, but also transfers mechanical signals between cells. Of particular interest is the observation that AmotL2 is required for nuclear shape and subcellular positioning. Ingber and coworkers showed early on that there is a direct linkage between the cytoskeleton and the cell nucleus opening up the possibility of a novel mechanical signaling pathway from the exterior to the nucleus(39). Actin filaments are directly associated with the nuclear lamina by binding to Nesprin2, SUN-1 and -2, and Lamin A that form the LINC complex. The actin filaments are anchored to the nuclear lamina via the LINC complex to form linear punctae called TAN lines. In this report we present evidence for a novel mechanical pathway that mechanically links junctional proteins to the nuclear LINC complex. We show that concomitant with the loss of AmotL2 and radial actin filaments, nuclei lose their central position and translocate to a polarized position in the cell, downstream of the exerted flow direction. Arterial nuclei not only lose their sub-cellular positioning, but the integrity of the nuclear lamina is perturbed. Measurement of forces exerted on the LINC proteins suggest that nuclear positioning is a result of the dynamic interactions of the cytoskeleton where nuclei are exposed to constant actomyosin forces(40). Depletion of AmotL2 also had consequences for the integrity of the nuclear lamina. The nuclear lamina is an intermediate filament meshwork composed of two types of lamin proteins, the B type (Lamin B1 and B2) and the A type (Lamin A/C) and associated inner nuclear membrane proteins(41). This network determines the mechanical properties and morphology of the nucleus(42). In particular, Lamin A has been proposed to be responsive to mechanical cues from the extracellular matrix. Reduction of Lamin A in the nuclear membrane also results in a less rigid nuclear membrane.

The lack of alignment and the consequent irregular shapes of the arterial endothelium concomitant with the loss of nuclear positioning had direct consequences for the endothelial function. We could show that AmotL2 deficient cells acquire a pro-inflammatory phenotype with ensuing formation of areas of vascular inflammation characterized by the presence of CD45^+^ monocytes. Quantitative PCR-analysis indicated a gender-specific difference in inflammatory response. The underlying reason is unclear but may relate to levels of estrogen as shown by other reports. Of clinical importance is the formation of AAA in the *amotl2*^ec-/ec-^ male mice. This is of interest as AAA has a relatively high prevalence in males between 65-79 years of age and the rupture of AAA is the cause of over 15 000 deaths/year in the USA alone and 175 000 globally(2, 43, 44). Our data indicate a lower expression of AmotL2 in AAA from patients as compared to healthy aortae. Furthermore, the expression levels of AmotL2 were negatively correlated with the diameter of the aneurysms. Several mouse models of AAA have been established for developing therapies for AAA. However, so far these models have failed to reliably predict results in clinical trials. Here we suggest a model where AmotL2 deficiency activates pro-inflammatory markers such as VCAM-1 and ICAM-1 on the EC surface. This promotes the extravasation of CD45+ inflammatory cells. The presence of inflammatory cells in the tunica media promotes the degradation of the extracellular matrix such as elastin, and thus weakens the physical strength of the aortic wall. The weakened area of the artery then bulges out and poses an increased risk of blood vessel rupture and hemorrhage (as shown in the schematic **Fig. 7g**). The possibility that lower levels of AmotL2 could increase the risk of developing vascular inflammation, opens up the potential to therapeutically restore the AmotL2 mechano-transductory pathway and thereby increase the resilience of the arterial wall to shear stress.

## Methods

### Human material

Aortic samples from AAA patients were obtained from the surgery performed at Karolinska Hospital in Stockholm. Control samples were taken from the abdominal aorta of beating-heart, solid organ transplant donors. Ethical permit (2009/1098-32) was granted by Regionala etikprövningsnämnden (Regional ethical review boards) in Stockholm. RNA was extracted from both medial and adventitia layer of the aortic wall and subsequently sequenced on HTA 2.0 (Human Transcriptome Array 2.0 - Affymetrix) platform.

### Mice and tamoxifen injections

The *amotl2*^*flox/flox*^ mice, carrying a loxP-flanked amotl2 gene, were crossed to Cdh5(PAC)^CreERT2^ and ROSA26-EYFP double transgenic mice. To induce endothelial-specific amotl2 gene inactivation, tamoxifen was administered by intraperitoneal (IP) injection for 5 continuous days. For adult mice over 6 weeks old, 100µl of tamoxifen (20mg/ml) was administered at each injection site. Analysis of mice samples was performed four to six weeks after injections. All mice in this report were in C57BL/6 background, and both females and males were included. Ethical permits were obtained from North Stockholm Animal Ethical Committee and all experiments were carried out in accordance with the guidelines of the Swedish Board of Agriculture.

### Tissue preparation

Mice were euthanized using carbon dioxide. Thoracic cavity was rapidly opened, and the heart was exposed while still beating. Cold PBS was injected through a cannula for perfusion for 1 minute and then changed to 4% paraformaldehyde (PFA) for another 1-minute perfusion. Aorta from root to aortic-common iliac bifurcation was dissected followed by careful removal of the connective tissues. After 1h extra fixation in 4% PFA, the entire aorta was opened longitudinally. For the aortic arch part, using spring scissors, the inner curvature was cut along anteriorly, while the outer curvature of the aorta was opened from the aortic root through innominate, carotid, and subclavian arteries until the aortic arch resembled a Y shaped split. The whole flattened-out aorta was pinned onto the wax mold and prepared for immunostaining. For RNA isolation, the aorta was perfused with cold PBS for 2 minutes before careful dissection. Thoracic and abdominal aorta were taken out and frozen in -80°C for subsequent RNA extraction.

To isolate the urinary bladder, the mice were handled especially gently before sacrifice, which prevented urine leakage. Full bladders were immersed in fixative for 2 hours (2h) and pinned on the wax mold in an open flower shape for whole-mount staining. Mouse eyeballs were taken out at postnatal day 6 (P6) and month 6 (week 24). After a 2h fixation in 4% PFA, the retinas were dissected out and prepared for immunofluorescence staining for analysis of vasculature.

### Cell culture

MS1 cells were cultured in RPMI-1640 medium (GIBCO, 21875-034) supplemented with 10% FBS (GIBCO, 10270-106) and 1% penicillin/streptomycin (P/S, GIBCO, 15140-122). BAE cells (Bovine Aortic Endothelial cell) purchased from Sigma-Aldrich (B304-05) were cultured in Bovine Endothelial Cell Growth Medium (Sigma, B211-500). Human Umbilical Vein Endothelial Cells (HUVEC) purchased from ScienCell (#8000) were cultured in Endothelial cell Medium (ScienCell, #1001). Human Aortic Endothelial Cell (HAoEC) were purchased from PromoCell (C-12271) and cultured with Endothelial Cell Growth Medium MV (C-22020). To note, the batch of HAoEC used for this study was from a 55-year-old male donor with a Caucasian background.

### Lentiviral induced knock-down

For knockdown studies, HAoECs (PromoCell) were infected with customized AmotL2 shRNA Lentiviral particles (Sigma) or scrambled shRNA control virus in complete endothelial cell medium with 5 μg/mL polybrene (Vector Builder). The Lentivirus-containing medium was removed after overnight incubation and fresh medium was added. Further analyses of confluent cells (WB, IF, Ibidi flow) were performed at ≥ 72h after infection.

### Flow experiments

#### Ibidi flow system

Flow chamber slides (Ibidi μ**-**Slides VI 0,4 Ibidi**-**treated) with a volume of 30 μl per parallel channel were coated with fibronectin (Sigma). HAoECs/HUVECs were grown on the slides for 24h until 30-50% confluency followed by infection with Lentivirus or control treatment. 96h after that cells were subjected to 14 dyn/cm^2^ laminar flow using the Ibidi pump system and pump control software or kept at static conditions and cultured for 48h at 37°C 5% CO_2_. Cells were then harvested and processed for further analysis.

#### Orbital shaker

The Rotamax 120 (Heidolph) complies well with cell incubator and generates circular motion with the maximum speed of 300rpm. HUVECs on a 15cm culture dish with 16ml medium were placed on the shaker for 96h before the harvesting the cell lysates.

### Immunofluorescence (IF) staining

Immunofluorescence staining on MS1 cells, HAoEC, and HUVECs was performed as previously described(25). To stain open aorta pinned on wax, endothelium exposed on the top layer was carefully treated using the same protocol as for cell staining, with the exception that the aorta was permeabilized for 20 minutes with 0.1% Tween 20 in PBS. Retinas and bladders were blocked and permeabilized in 1% BSA and 0.3% Triton-X100 in PBS overnight. Pblec buffer (1.0% Triton X-100 plus 0.1M MgCl2, 0.1M CaCl2, 0.01M MnCl2 in PBS) was used for washing and incubating one or more primary antibodies. Then fluorophore-conjugated antibodies were added in blocking buffer, followed by 5 x 20-minutes wash with blocking buffer at 1:1 dilution in PBS. The cells/whole tissue was finally mounted with Fluoroshield™ with DAPI (Sigma, F6057). For a list of primary and secondary antibodies used see ***Supplementary Table 5***.

### Western blot (WB)

Cells were scraped directly from the culture dish in lysis buffer (50 mM Hepes buffer, 150 mM NaCl, 1.5 mM MgCl_2_, 1 mM EGTA, 10% Glycerol, 1% TritonX-100), with 1 x protease inhibitor (Roche, 04693159001) and optionally with Phosphatase Inhibitor Cocktail 1 (Sigma, P2850). Lysates were prepared with SDS sample buffer (Novex, 1225644) containing 10 % sample reducing agent (Novex, 1176192), separated in a polyacrylamide gel with 4-12% gradient (Novex, NP0322BOX), and transferred to a nitrocelluose membrane (Whatman, 10401396). The membrane was blocked in 5% non-fat milk PBS with 0.1% Tween 20 and sequentially incubated with the primary antibody at 4°C overnight. Horseradish peroxidase (HRP) conjugated secondary antibody was added in order for labelled proteins to be detected by Western Lightning Plus-ECL (PerkinElmer, 203-170071). Primary and secondary antibodies used are listed in ***Supplementary Table 5***.

### Co-immunoprecipitation (Co-IP) analysis

Mouse lung tissues were cut into small pieces before transferring into lysis buffer for western blot. Tissue homogenizer was gently applied for better protein extraction. Cell/tissue lysates were incubated with protein G sepharose beads (GE, 17-0618-01) for 1.5 hours at 4°C as pre-cleaning. Afterwards 2µg Amotl2, Lamin A/C, VE-cadherin antibody or control IgG were added in the lysates overnight at 4°C. The next morning, the immunocomplexes were enriched by protein G beads for 2h at 4°C, followed by five-time washing with lysis buffer. The final protein samples were fractionated by polyacrylamide gel and fractions were western blotted for evaluation of IP protein input level.

### Mass spectrometry (MS) analysis

IP samples were dissolved in 200µL lysis buffer (4 % SDS, 50 mM HEPES pH 7.6, 1 mM DTT). After heating at 95°C for 5 min and centrifugation at 14,000g for 15 min, the supernatant was transferred to new tubes. Protein digestion was performed using a modified SP3-protocol. In brief, each sample was alkylated with 40 mM chloroacetamide and Sera-Mag SP3 bead mix (20µl) was transferred into the protein sample together with 100 % acetonitrile to a final concentration of 70 %. The mix was incubated under rotation at room temperature for 20 min. The mixture was placed on the magnetic rack and the supernatant was discarded, followed by two washes with 70 % ethanol and one with 100 % acetonitrile. The beads-protein-complex was reconstituted in 100µl trypsin buffer (50 mM HEPES pH 7.6 and 0.8µg trypsin) and incubated overnight at 37°C. The eluted samples were dried in a SpeedVac. Approximately 10µg was suspended in LC mobile phase A and 1µg was injected on the LC-MS/MS system.

Online LC-MS was performed using a Dionex UltiMate™ 3000 RSLCnano System coupled to a Q-Exactive-HF mass spectrometer (Thermo Scientific). IP samples were trapped on a C18 guard-desalting column (Acclaim PepMap 100, 75μm x 2cm, nanoViper, C18, 5µm, 100 Å), and separated on a 50cm long C18 column (Easy spray PepMap RSLC, C18, 2μm, 100 Å, 75μm x 50cm). The nano capillary solvent A was 95% water, 5% DMSO, 0.1% formic acid; and solvent B was 5% water, 5% DMSO, 95% acetonitrile, 0.1% formic acid. At a constant flow of 0.25μl min^-1^, the curved gradient went from 2% B up to 40 % B in 240 min, followed by a steep increase to 100% B in 5 min.

FTMS master scans with 70,000 resolution (and mass range 300-1700 m/z) were followed by data-dependent MS/MS (35,000 resolution) on the top 5 ions using higher-energy collision dissociation (HCD) at 30-40 % normalized collision energy. Precursors were isolated with a 2m/z window. Automatic gain control (AGC) targets were 1e^6^ for MS1 and 1e^5^ for MS2. Maximum injection times were 100ms for MS1 and 150-200 ms for MS2. The entire duty cycle lasted ∼2.5s. Dynamic exclusion was used with 60s duration. Precursors with unassigned charge state or charge state 1 were excluded and an underfill ratio of 1% was used.

### Quantitative real-time RT-PCR (qRT-PCR)

To extract RNA from dissected aorta from *amotl2*^ec+/ec+^ (n=3) and *amotl2*^ec-/ec-^ mice (n=3) was immersed in TRIzol and homogenised by TissueLyser (Qiagen) in TRIzol. Chloroform addition allowed the homogenate to separate into the lower organic phase and the upper clear aqueous phase (containing RNA). MS1 cells were scraped directly from culture dish in RLT buffer. Total RNA purification was carried out using RNeasy Plus Mini Kit (Qiagen), after which RNA concentrations were measured by Nanodrop spectrophotometer (ThermoFisher Scientific). cDNA synthesis was performed using the High-Capacity cDNA Reverse Transcription Kit (Applied Biosystems). Quantitive real-time PCR was performed on a 7900HT Fast Real-Time PCR system (Perkin-Elmer Applied Biosystems) using TaqMan Assay-on-Demand (Applied Biosystems). Results were calculated as 2-ΔCT obtained by comparing the threshold cycle (CT) for the genes of interest with that obtained for housekeeping gene hypoxanthine guanine phosphoribosyltransferase (HPRT). All TaqMan probes involved were purchased from Thermo Fisher Scientific and detailed information can be found in ***Supplementary Table 5***.

### RNA-seq

Total RNA purified from full-length aorta (thoracic and abdominal aorta) of amotl2^ec+/ec+^ (n=3) and amotl2^ec-/ec-^ mice (n=3) were sent for RNA-seq analysis (Novogene, Beijing, China). Libraries were prepared from 4-5µg of total RNA. PolyA RNA was purified using the Dynabeads mRNA purification kit (Ambion) and fragmented using Fragmentation Reagent (Ambion). First strand cDNA was synthesized from polyA RNA using the SuperScript III Reverse Transcriptase Kit with random primers (Life Technologies). Second strand cDNA synthesis was performed using Second Strand Synthesis buffer, DNA Pol I, and RNase H (Life Technologies). cDNA libraries were prepared for sequencing using the mRNA TruSeq protocol (Illumina).

The genes with significantly differential expression were input to online Enrichr (Ma’ayan Laboratory, Computational systems biology) for GO term analysis (Biological process 2018)(45, 46).

### Statistical analysis

All statistical figures and analyses were performed using GraphPad Prism software, with the exception of the gene correlation graphs, which were generated using R (https://www.r-project.org/index.html). Statistical analysis of *in vivo* results was based on at least three animals per group. Comparisons between two groups with similar variances were analyzed by the standard unpaired two-tailed Student t-test, while comparisons between multiple groups were analyzed by Kruskal–Wallis test. The correlation between two genes were analyzed by Pearson Correlation and pearson correlation coefficient is referred to *r*. A value of *P*<0.05 was considered as statistically significant (*n*.*s*. not significant, ^*^*P* <0.05, ^**^*P* <0.01 and ^***^*P* <0.001). No statistical method was used to predetermine sample size in animal studies and the experiments were not randomized. The investigators were not blinded to allocation during experiments and outcome assessment.

## Acknowledgments

We are indebted to Dr. Ralf H Adams, University of Münster, who kindly provided the Cdh5(PAC)-CreER^T2^ and ROSA26-EYFP transgenic mice. Many thanks to Lynn Butler and Philip Dusart for letting us perform experiments using their Ibidi Flow System at the SciLifeLab, Stockholm, Sweden. The paraffin sections and IHC staining of the aortae were performed by Anna Malmerfelt in histology core facility at the Department of Oncology and Pathology, Karolinska Institute, Sweden. We are grateful for the polar bar chart coded by freelance programmer Andreas Gustafsson using Python.

This study was supported by grants from the Swedish Heart and Lung Foundation, Novo Nordisk Foundation (NNF15CC0018346) the Swedish Cancer Society, the Swedish Childhood Cancer Foundation, Cancer Society of Stockholm, the Swedish Research Council and Knut and Alice Wallenberg Foundation.

## Author contributions

Y.Z. performed *in vivo* experiments using the AmotL2 transgenic model, performed in vitro experiments, finalized all figures, and wrote the manuscript. E.H. performed Lenti-virus based transfection *in vitro* and Ibidi flow experiment with data analysis. S.H. performed *in vivo* experiments on the Amotl2 mouse model and summarized data. O.B. and P.E. performed bio-informatical analysis on human aortic samples. J.R. contributed with human samples from clinical surgery. D.K. designed the experiments to investigate inflammation and M.F. performed the TaqMan qPCR on RNA from murine aortic samples. Z.A. performed flow experiment with orbital shaker. L.H developed the theory, designed experiments and wrote the manuscript. All authors reviewed the manuscript prior to submission.

## Competing financial interests

The authors declare no competing financial interests.

## Notes

### Competing Interest Statement

The authors have declared no competing interest.

