## Supplementary figures and tables for "The VE-cadherin/AmotL2 mechanosensory pathway suppresses aortic inflammation and the formation of abdominal aortic aneurysms"

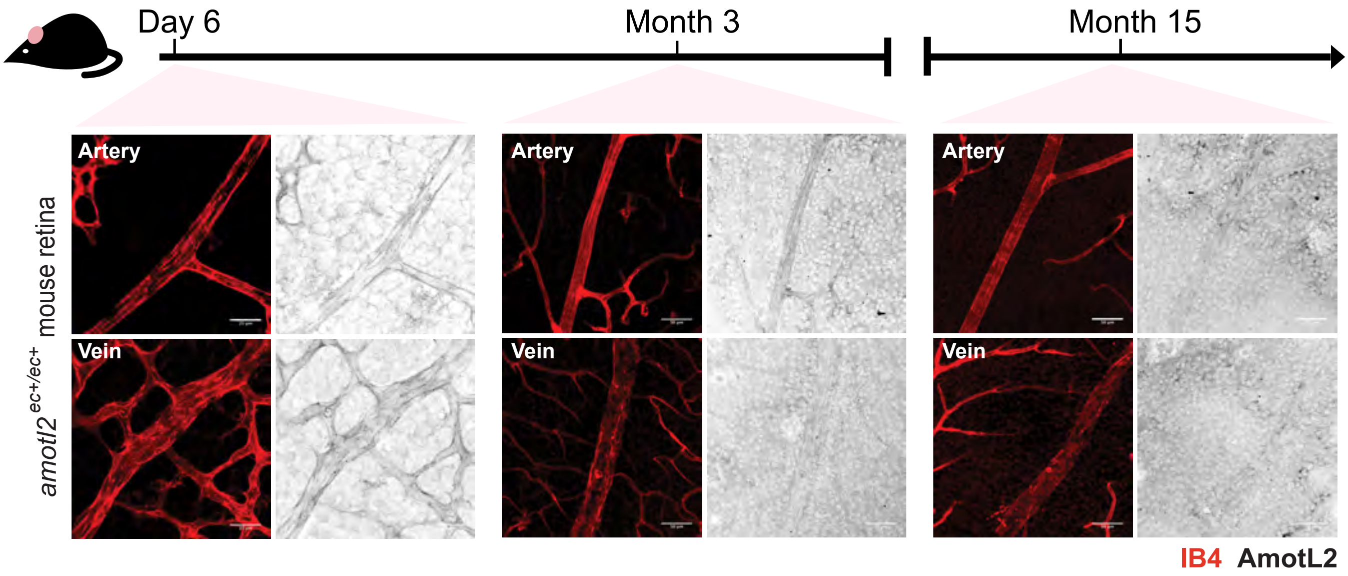


**Supplementary Figure 1**

**Supplementary Figure 1. AmotL2 expression pattern in mouse retina of different ages.** *amotl2*$^{ec+/ec+}$ mice retinas were stained with IB4 (in red) and AmotL2 (in gray) at the age of postnatal day 6, month 3 and month 15, which are placed in the left, middle and right panel, respectively. Both representative images of vasculature in arteries and veins are shown for each time point. Scale bars: 25 µm (left panel) and 50 µm (middle/right panel).

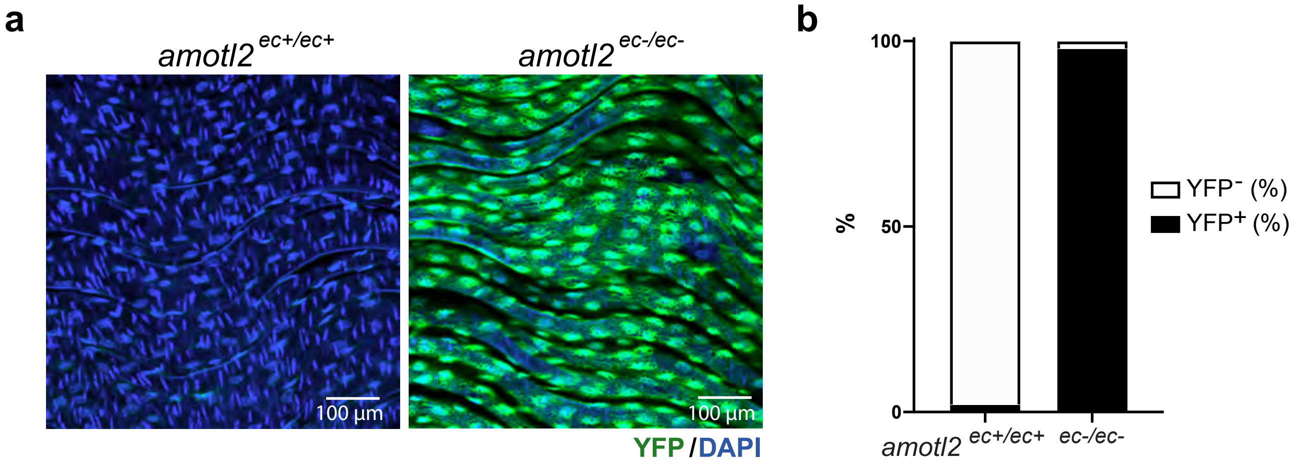


**Supplementary Figure 2**

**Supplementary Figure 2. Recombination efficiency in *amotl2*$^{flox/flox}$ Cdh5(PAC)$^{CreERT2}$ ROSA26-EYFP mice. a,** GFP immunofluorescence staining (in green) was performed to visualize the YFP reporter, as a marker for Cre-recombinase expression of DAs in *amotl2*$^{ec+/ec+}$ and *amotl2*$^{ec-/ec-}$ Cdh5(PAC)$^{CreERT2}$ ROSA26-EYFP mice. **b,** Quantification of the percentage $YFP^+/YFP^-$ of 3 representative *amotl2*$^{ec+/ec+}$ and *amotl2*$^{ec-/ec-}$ DAs. Size bars: 100 µm.

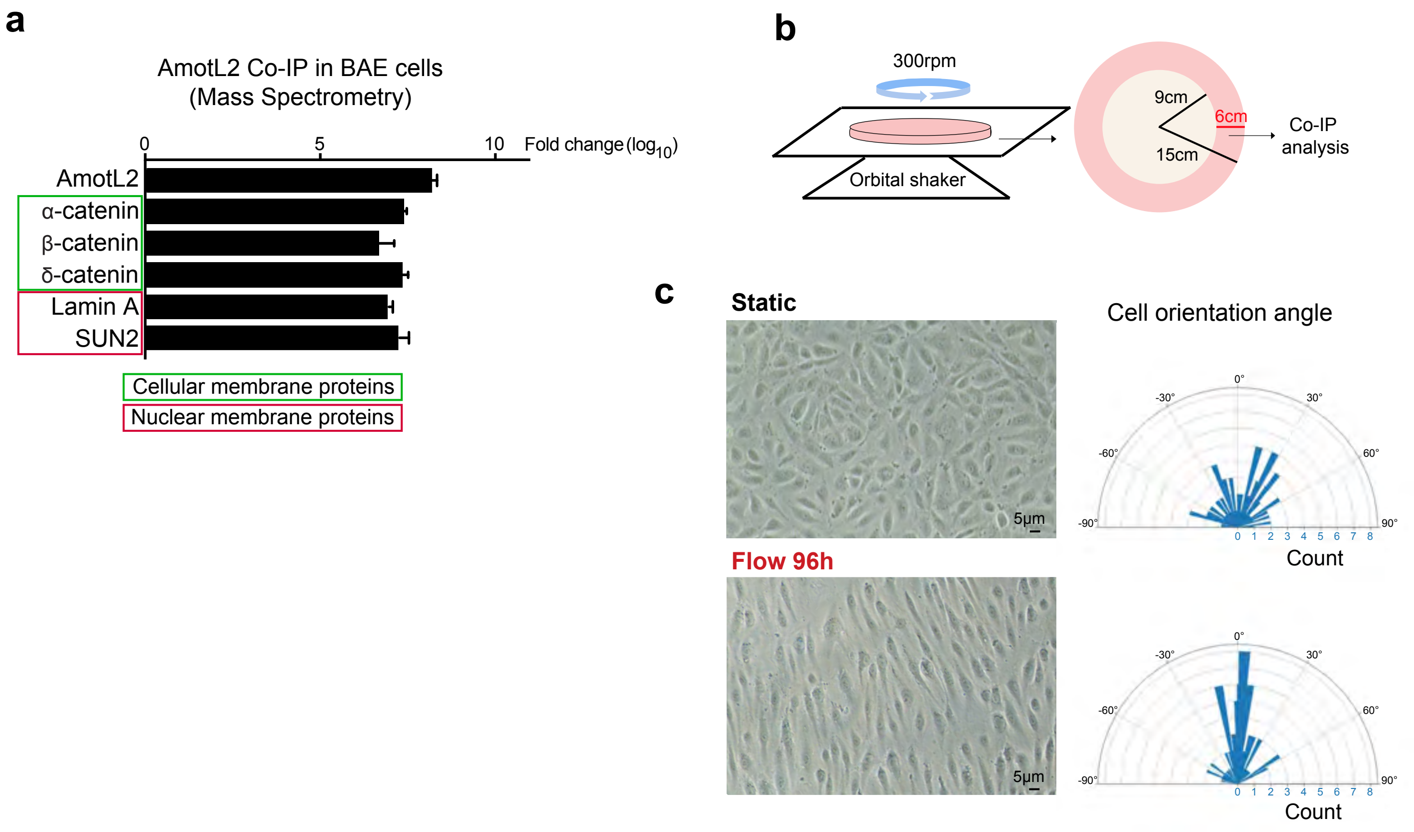


**Supplementary Figure 3**

**Supplementary Figure 3. AmotL2 links VE-cadherin to LINC complex through actin. a,** Catenins in cellular membrane proteins (framed in green box) and nuclear membrane proteins (framed in red box) were identified from AmotL2 immunoprecipitates from BAE cells analyzed by MS. The data was displayed with fold change ($log_{10}$) as compared to control IP samples. **b,** Schematic describing how circulatory flow was applied to HUVECs cultured in 15cm dish (300rpm, 96h). The peripheral area with a width of 6cm (pink area) is where the cells were harvested for CO-IP experiments. **c,** Bright field images of HUVECs located in the pink area indicated in **b**, illustrating the cell morphology in both static and one post-flow conditions (96h on orbital shaker). Polar bar chart depicting the angle of cell orientation are presented on the right side. The mean of the actual angular value was normalized to 0° in both conditions. Size bars: 5 μm.

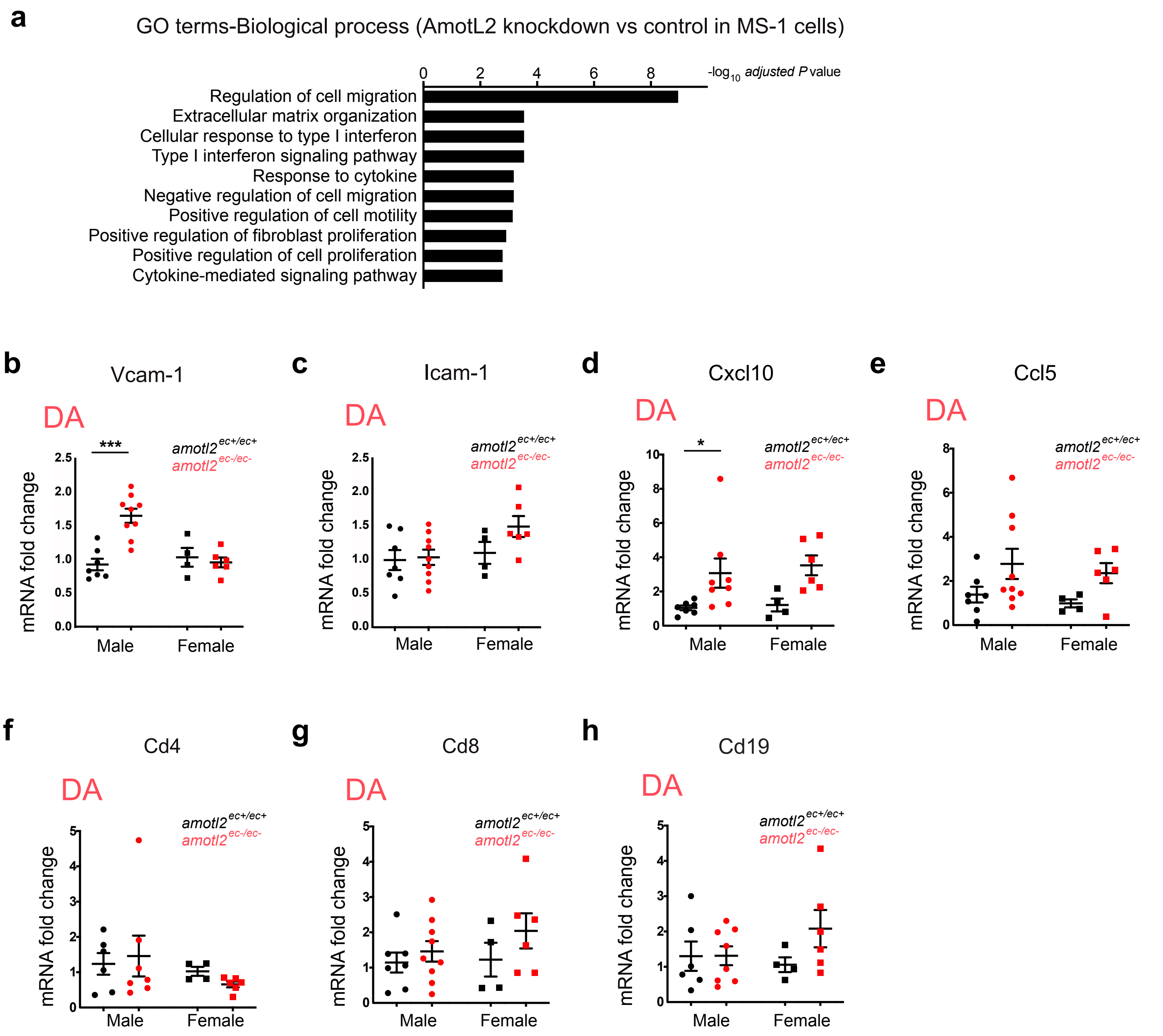


**Supplementary Figure 4**

**Supplementary Figure 4. Absence of AmotL2 provokes immune response in the aorta. a,** Enriched GO terms (Biological process 2018) analyzed by Enrichr (**Supplementary Table 4**) and presented in the graph ranking by $-\log_{10}$ *adjusted P* value. mRNA isolated from MS1 cells treated by control or AmotL2 siRNA (n=3) mice were sent for RNA-sequencing analysis. Top 500 differentially expressed genes with the lowest *adjusted-P* value were subjected to GO terms matching. **b-h,** mRNA was isolated separately from descending aorta (DA, in red), and analyzed by TaqMan qRT-PCR. Relative expression levels of *Vcam-1* (**b**), *Icam-1* (**c**), *Cxcl10* (**d**), *Ccl5* (**e**), *Cd4* (**f**), *Cd8* (**g**), and *Cd19* (**h**) were normalized versus *Hprt*. mRNA was obtained from about 10 *amotl2*$^{ec+/ec+}$ mice (in black dots, male n=5-7, female n=4-5) and 14 *amotl2*$^{ec-/ec-}$ mice (in red dots, male n=7-9 and female n=6). Fold-changes are quantified, and data shown are mean ± S.E. $^{*}P<0.05$. $^{***}P<0.001$. To note that, all *the p-values* unlabelled in the graphs indicate "not statistically significant".

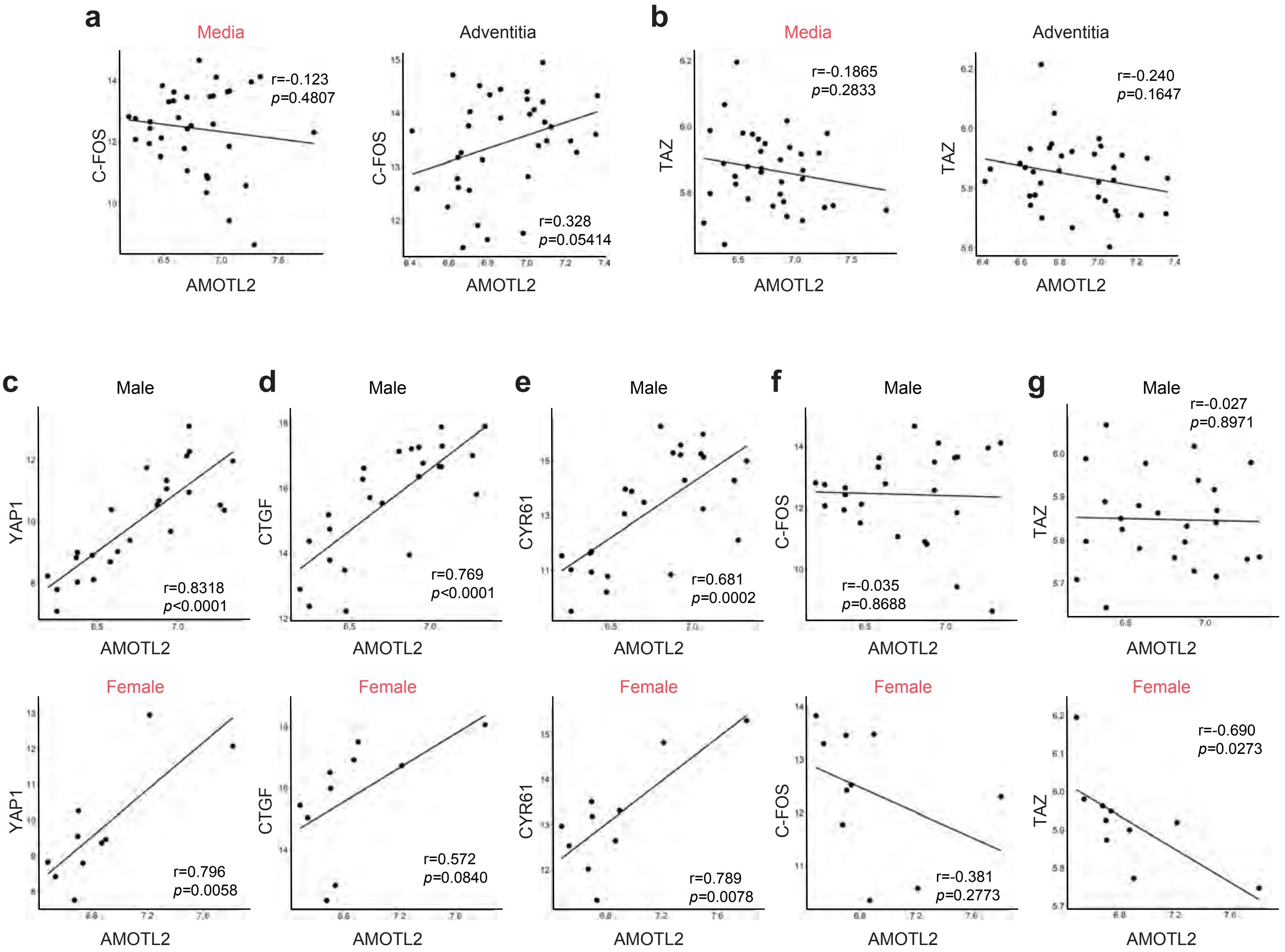


**Supplementary Figure 5**

**Supplementary Figure 5. The AmotL2 correlation with Hippo pathway in AAA samples from human patients.** mRNA expression level correlations of AmotL2 with C-FOS in **a,** and TAZ in **b** were analyzed separately in media (left panel) and adventitia (right panel) tissues. 25 male and 10 female patients were enrolled. Aortic media tissues from male and female of AAA patients were analyzed separately for AmotL2 mRNA expression correlation with YAP1 (**c**), CTGF (**d**), CYR61 (**e**), C-FOS (**f**) and TAZ (**g**). Correlation coefficient *r* and *p-value* are labelled in each individual figure.

### Figure 3a

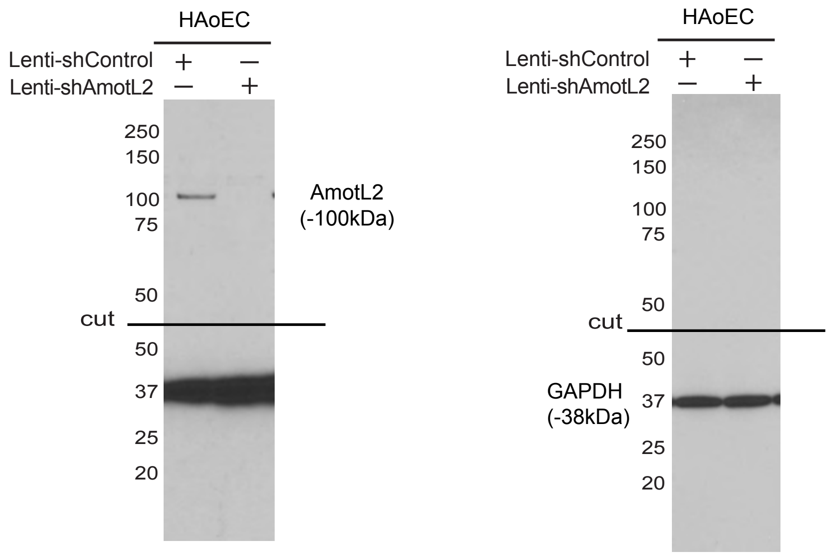


### Figure 4b

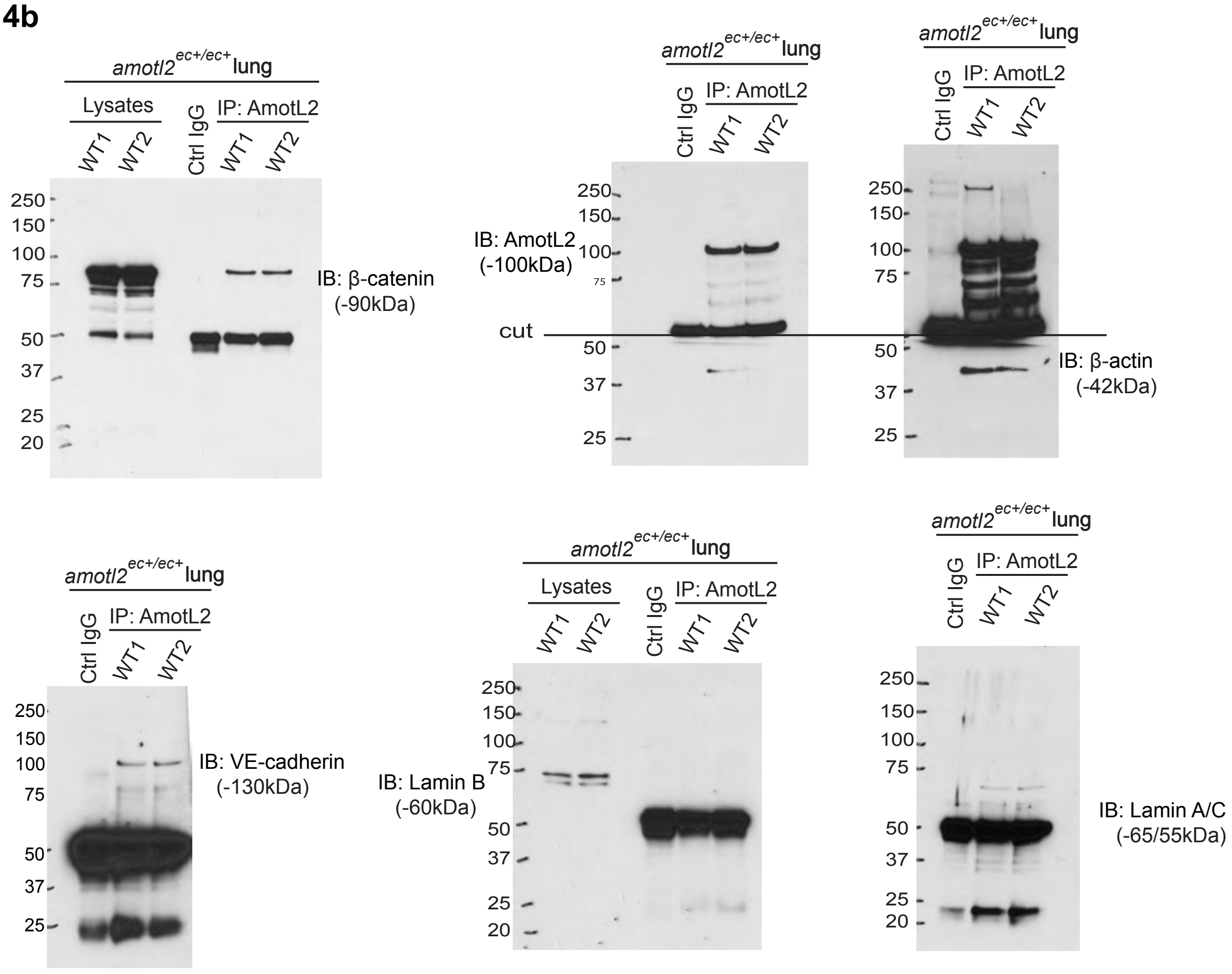


**Supplementary Figure 6.** Full length blots to Figure 3a and 4b.

### Figure 4c

All in MS1 cells

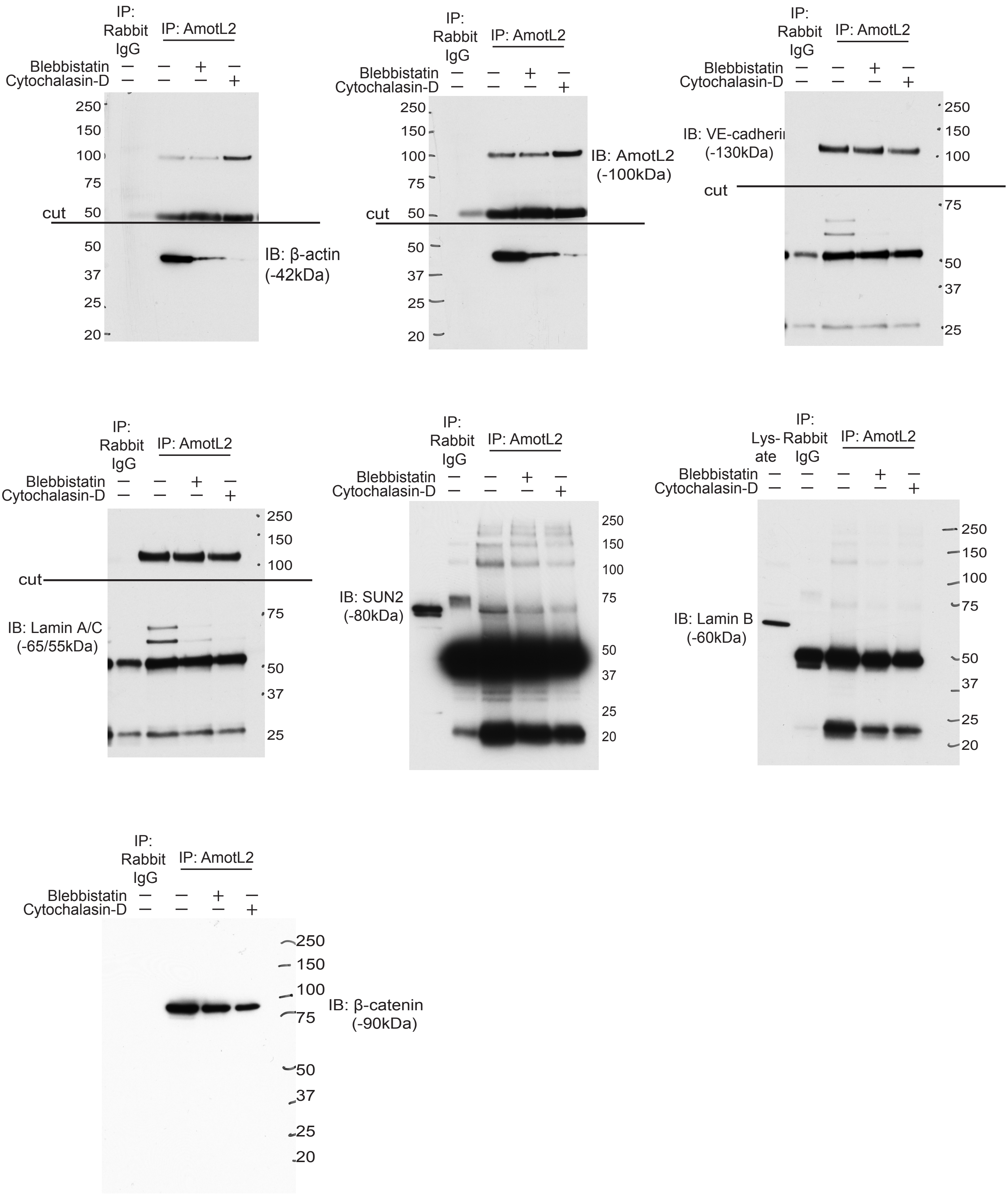


**Supplementary Figure 7.** Full length blots to Figure 4c.

**Figure 4d**

All in HAoEC

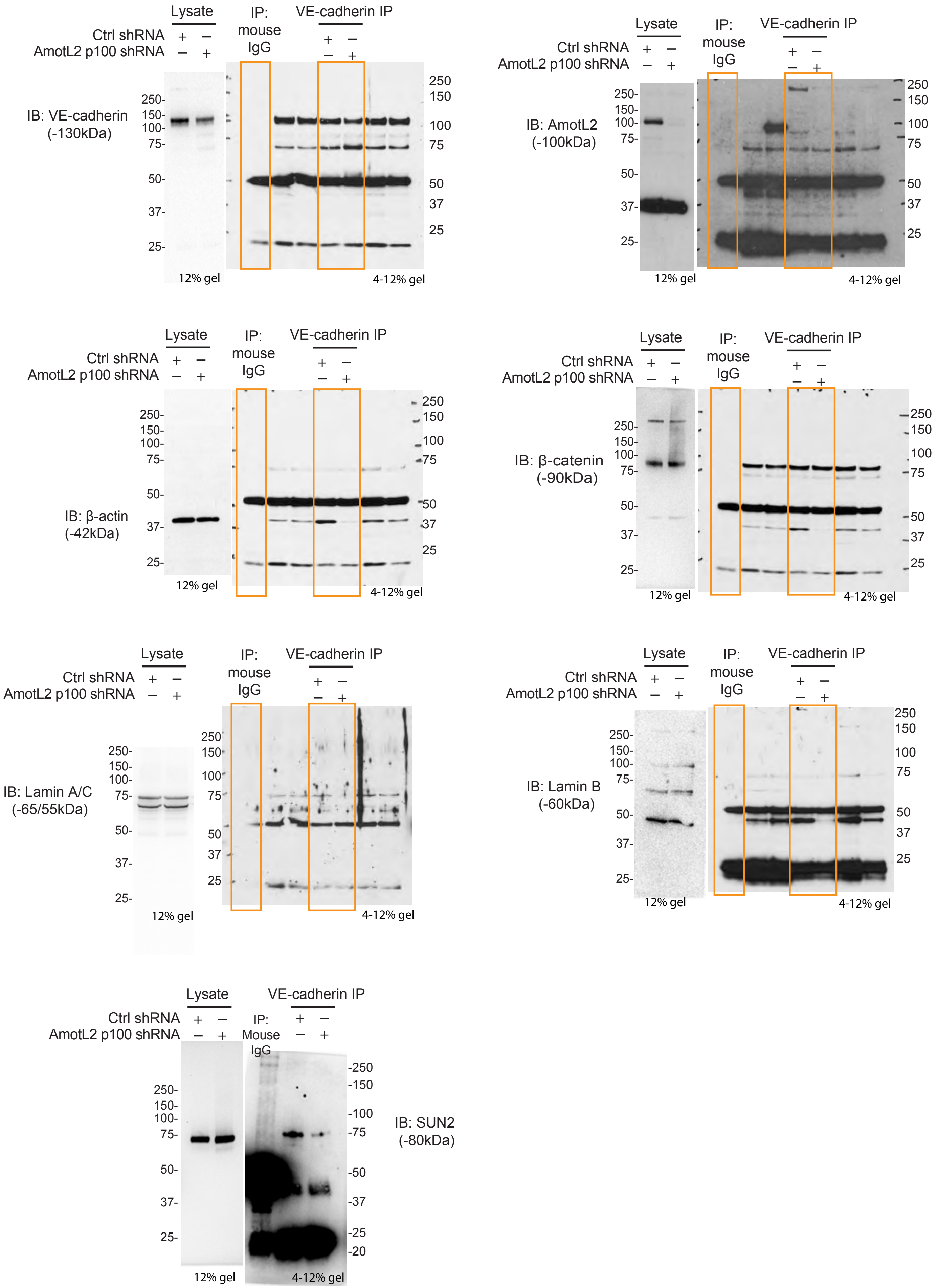


**Supplementary Figure 8.** Full length blots to Figure 4d.

### Figure 4e

All in HUVEC

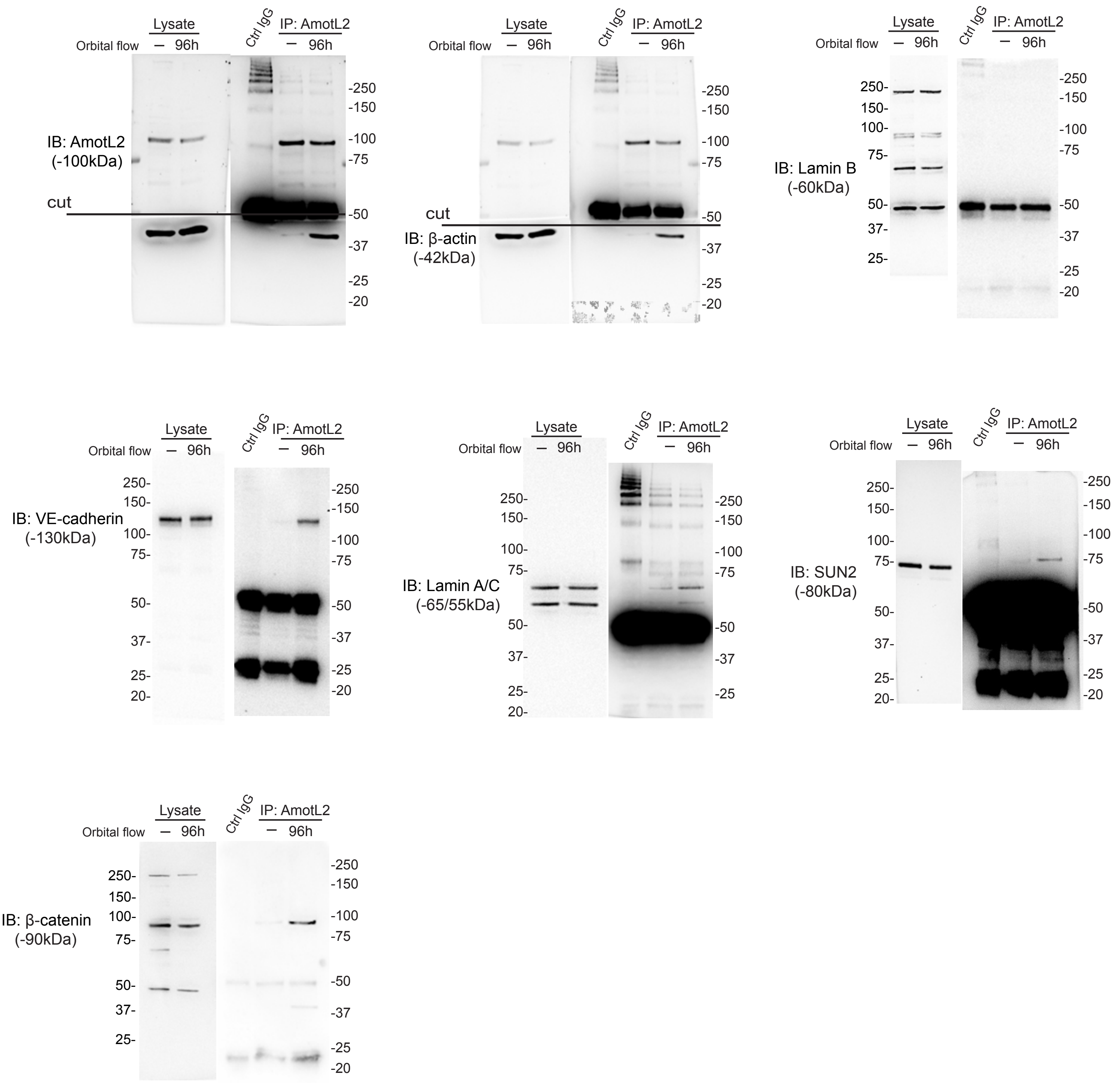


**Supplementary Figure 9.** Full length blots to Figure 4e.

**Supplementary Table 1. Mass-spec analysis of AmotL2 immunoprecipitation in MS1 cells**

| Accession | Description | Area_Ctrl_1 | Area_Ctrl_2 | Area_Ctrl_3 | Area_AmotL2_1 | Area_AmotL2_2 | Area_AmotL2_3 | Gene name |
|---|---|---|---|---|---|---|---|---|
| P99027 | 60S acidic ribosomal protein P2 OS=Mus musculus GN=Rplp2 PE=1 SV=3 - [RLA2_MOUSE] | 1,197E7 | 0,000E0 | 0,000E0 | 1,025E8 | 1,555E8 | 5,822E8 | Rplp2 |
| P62204 | Calmodulin OS=Mus musculus GN=Calm1 PE=1 SV=2 - [CALM_MOUSE] | 7,282E7 | 0,000E0 | 8,936E7 | 1,808E9 | 4,501E9 | 9,019E9 | Calm1 |
| P58771-2 | Isoform 2 of Tropomyosin alpha-1 chain OS=Mus musculus GN=Tpm1 - [TPM1_MOUSE] | 0,000E0 | 0,000E0 | 1,738E7 | 1,752E9 | 2,676E9 | 4,298E9 | Tpm1 |
| P60710 | Actin, cytoplasmic 1 OS=Mus musculus GN=Actb PE=1 SV=1 - [ACTB_MOUSE] | 9,315E8 | 8,508E5 | 2,323E8 | 1,154E10 | 2,141E10 | 1,591E10 | Actb |
| Q60605 | Myosin light polypeptide 6 OS=Mus musculus GN=Myl6 PE=1 SV=3 - [MYL6_MOUSE] | 3,418E8 | 0,000E0 | 1,038E8 | 7,357E9 | 1,214E10 | 1,374E10 | Myl6 |
| Q60605-2 | Isoform Smooth muscle of Myosin light polypeptide 6 OS=Mus musculus GN=Myl6 - [MYL6_MOUSE] | 3,354E8 | 0,000E0 | 1,038E8 | 7,357E9 | 1,214E10 | 1,374E10 | Myl6 |
| Q9JJU8 | SH3 domain-binding glutamic acid-rich-like protein OS=Mus musculus GN=Sh3bgrl PE=1 SV=1 - [SH3L1_MOUSE] | 0,000E0 | 0,000E0 | 0,000E0 | 2,505E6 | 1,384E7 | 7,620E7 | Sh3bgrl |
| P20152 | Vimentin OS=Mus musculus GN=Vim PE=1 SV=3 - [VIME_MOUSE] | 6,052E7 | 0,000E0 | 4,966E7 | 1,116E8 | 2,178E8 | 1,545E9 | Vim |
| P63168 | Dynein light chain 1, cytoplasmic OS=Mus musculus GN=Dynll1 PE=1 SV=1 - [DYL1_MOUSE] | 0,000E0 | 0,000E0 | 9,000E5 | 6,575E7 | 3,637E6 | 2,389E6 | Dynll1 |
| P62908 | 40S ribosomal protein S3 OS=Mus musculus GN=Rps3 PE=1 SV=1 - [RS3_MOUSE] | 2,511E7 | 0,000E0 | 1,247E7 | 5,806E8 | 1,105E9 | 2,353E9 | Rps3 |
| P18760 | Cofilin-1 OS=Mus musculus GN=Cfl1 PE=1 SV=3 - [COF1_MOUSE] | 0,000E0 | 0,000E0 | 1,219E7 | 2,350E8 | 6,076E8 | 5,269E8 | Cfl1 |
| Q6IRU2 | Tropomyosin alpha-4 chain OS=Mus musculus GN=Tpm4 PE=1 SV=3 - [TPM4_MOUSE] | 0,000E0 | 0,000E0 | 1,218E7 | 7,452E8 | 6,204E8 | 9,875E8 | Tpm4 |
| Q3THE2 | Myosin regulatory light chain 12B OS=Mus musculus GN=Myl12b PE=1 SV=2 - [ML12B_MOUSE] | 1,647E8 | 0,000E0 | 1,331E8 | 5,052E9 | 8,498E9 | 1,342E10 | Myl12b |
| Q8VDD5 | Myosin-9 OS=Mus musculus GN=Myh9 PE=1 SV=4 - [MYH9_MOUSE] | 7,800E8 | 0,000E0 | 3,751E8 | 1,395E10 | 2,268E10 | 1,618E10 | Myh9 |
| P63101 | 14-3-3 protein zeta/delta OS=Mus musculus GN=Ywhaz PE=1 SV=1 - [1433Z_MOUSE] | 0,000E0 | 0,000E0 | 5,426E6 | 1,197E8 | 2,006E8 | 3,279E8 | Ywhaz |
| Q9WTI7-3 | Isoform 3 of Unconventional myosin-Ic OS=Mus musculus GN=Myo1c - [MYO1C_MOUSE] | 5,544E7 | 0,000E0 | 4,243E7 | 1,796E9 | 4,341E9 | 1,165E10 | Myo1c |
| Q9WTI7 | Unconventional myosin-Ic OS=Mus musculus GN=Myo1c PE=1 SV=2 - [MYO1C_MOUSE] | 5,544E7 | 0,000E0 | 4,243E7 | 1,796E9 | 4,341E9 | 1,165E10 | Myo1c |
| Q9WVK4 | EH domain-containing protein 1 OS=Mus musculus GN=Ehd1 PE=1 SV=1 - [EHD1_MOUSE] | 0,000E0 | 0,000E0 | 0,000E0 | 1,740E8 | 2,243E8 | 3,573E8 | Ehd1 |
| P21107-2 | Isoform 2 of Tropomyosin alpha-3 chain OS=Mus musculus GN=Tpm3 - [TPM3_MOUSE] | 2,921E7 | 0,000E0 | 1,212E7 | 2,040E9 | 3,139E9 | 4,747E9 | Tpm3 |
| P62259 | 14-3-3 protein epsilon OS=Mus musculus GN=Ywhae PE=1 SV=1 - [1433E_MOUSE] | 0,000E0 | 0,000E0 | 4,263E6 | 1,606E8 | 3,051E8 | 5,462E8 | Ywhae |
| P47757-2 | Isoform 2 of F-actin-capping protein subunit beta OS=Mus musculus GN=Capzb - [CAPZB_MOUSE] | 0,000E0 | 0,000E0 | 0,000E0 | 2,392E8 | 5,788E8 | 8,469E8 | Capzb |
| P62702 | 40S ribosomal protein S4, X isoform OS=Mus musculus GN=Rps4x PE=1 SV=2 - [RS4X_MOUSE] | 0,000E0 | 0,000E0 | 0,000E0 | 4,817E8 | 8,993E8 | 1,160E9 | Rps4x |
| P59999 | Actin-related protein 2/3 complex subunit 4 OS=Mus musculus GN=Arpc4 PE=1 SV=3 - [ARPC4_MOUSE] | 0,000E0 | 0,000E0 | 0,000E0 | 1,485E8 | 3,024E8 | 4,275E8 | Arpc4 |
| P47754 | F-actin-capping protein subunit alpha-2 OS=Mus musculus GN=Capza2 PE=1 SV=3 - [CAZA2_MOUSE] | 0,000E0 | 0,000E0 | 2,214E5 | 1,560E8 | 3,100E8 | 2,515E8 | Capza2 |
| Q9D0M5 | Dynein light chain 2, cytoplasmic OS=Mus musculus GN=Dynll2 PE=1 SV=1 - [DYL2_MOUSE] | 0,000E0 | 0,000E0 | 1,134E6 | 3,076E7 | 4,492E6 | 2,203E6 | Dynll2 |
| Q61879 | Myosin-10 OS=Mus musculus GN=Myh10 PE=1 SV=2 - [MYH10_MOUSE] | 1,848E8 | 0,000E0 | 7,692E7 | 5,644E9 | 9,609E9 | 6,899E9 | Myh10 |
| P68040 | Receptor of activated protein C kinase 1 OS=Mus musculus GN=Rack1 PE=1 SV=3 - [RACK1_MOUSE] | 0,000E0 | 0,000E0 | 7,614E6 | 2,549E8 | 4,560E8 | 1,075E9 | Rack1 |
| P47757 | F-actin-capping protein subunit beta OS=Mus musculus GN=Capzb PE=1 SV=3 - [CAPZB_MOUSE] | 0,000E0 | 0,000E0 | 0,000E0 | 2,392E8 | 5,788E8 | 8,469E8 | Capzb |
| P57780 | Alpha-actinin-4 OS=Mus musculus GN=Actn4 PE=1 SV=1 - [ACTN4_MOUSE] | 0,000E0 | 0,000E0 | 3,995E7 | 6,351E7 | 2,203E8 | 3,475E8 | Actn4 |
| Q9CQV8-2 | Isoform Short of 14-3-3 protein beta/alpha OS=Mus musculus GN=Ywhab - [1433B_MOUSE] | 0,000E0 | 0,000E0 | 3,951E6 | 8,622E7 | 9,673E7 | 1,169E8 | Ywhab |
| P10854 | Histone H2B type 1-M OS=Mus musculus GN=Hist1h2bm PE=1 SV=2 - [H2B1M_MOUSE] | 0,000E0 | 0,000E0 | 3,713E6 | 5,988E7 | 6,547E7 | 2,447E8 | Hist1h2bm |
| P14131 | 40S ribosomal protein S16 OS=Mus musculus GN=Rps16 PE=1 SV=4 - [RS16_MOUSE] | 0,000E0 | 0,000E0 | 3,807E6 | 3,581E8 | 6,287E8 | 1,211E9 | Rps16 |
| P62889 | 60S ribosomal protein L30 OS=Mus musculus GN=Rpl30 PE=1 SV=2 - [RL30_MOUSE] | 0,000E0 | 0,000E0 | 3,622E6 | 1,217E8 | 3,463E8 | 4,716E8 | Rpl30 |
| P62962 | Profilin-1 OS=Mus musculus GN=Pfn1 PE=1 SV=2 - [PROF1_MOUSE] | 0,000E0 | 0,000E0 | 0,000E0 | 6,477E7 | 5,550E7 | 1,591E8 | Pfn1 |
| P26041 | Moesin OS=Mus musculus GN=Msn PE=1 SV=3 - [MOES_MOUSE] | 0,000E0 | 0,000E0 | 8,796E5 | 1,725E8 | 6,499E8 | 8,865E8 | Msn |
| Q9WTX5 | S-phase kinase-associated protein 1 OS=Mus musculus GN=Skp1 PE=1 SV=3 - [SKP1_MOUSE] | 0,000E0 | 0,000E0 | 3,377E6 | 1,064E7 | 4,061E7 | 7,292E7 | Skp1 |
| Q9QXS1-3 | Isoform PLEC-1A of Plectin OS=Mus musculus GN=Plec - [PLEC_MOUSE] | 0,000E0 | 0,000E0 | 3,986E6 | 3,329E8 | 7,844E8 | 1,021E9 | Plec |
| P39447 | Tight junction protein ZO-1 OS=Mus musculus GN=Tjp1 PE=1 SV=2 - [ZO1_MOUSE] | 0,000E0 | 0,000E0 | 1,395E5 | 5,906E8 | 1,169E9 | 7,026E8 | Tjp1 |
| Q9CPW4 | Actin-related protein 2/3 complex subunit 5 OS=Mus musculus GN=Arpc5 PE=1 SV=3 - [ARPC5_MOUSE] | 0,000E0 | 0,000E0 | 0,000E0 | 3,182E7 | 5,082E7 | 1,035E8 | Arpc5 |
| Q9WVA4 | Transgelin-2 OS=Mus musculus GN=Tagln2 PE=1 SV=4 - [TAGL2_MOUSE] | 0,000E0 | 0,000E0 | 0,000E0 | 1,844E7 | 5,034E7 | 9,225E7 | Tagln2 |
| Q9EQP2 | EH domain-containing protein 4 OS=Mus musculus GN=Ehd4 PE=1 SV=1 - [EHD4_MOUSE] | 0,000E0 | 0,000E0 | 1,218E6 | 2,279E8 | 3,490E8 | 4,719E8 | Ehd4 |
| P62830 | 60S ribosomal protein L23 OS=Mus musculus GN=Rpl23 PE=1 SV=1 - [RL23_MOUSE] | 0,000E0 | 0,000E0 | 1,581E7 | 1,267E8 | 2,247E8 | 2,290E8 | Rpl23 |
| P45591 | Cofilin-2 OS=Mus musculus GN=Cfl2 PE=1 SV=1 - [COF2_MOUSE] | 0,000E0 | 0,000E0 | 1,310E7 | 2,582E8 | 3,874E8 | 2,411E8 | Cfl2 |
| Q7TPR4 | Alpha-actinin-1 OS=Mus musculus GN=Actn1 PE=1 SV=1 - [ACTN1_MOUSE] | 0,000E0 | 0,000E0 | 3,995E7 | 6,662E7 | 2,352E8 | 3,356E8 | Actn1 |
| Q9QXS1-2 | Isoform PLEC-1 of Plectin OS=Mus musculus GN=Plec - [PLEC_MOUSE] | 0,000E0 | 0,000E0 | 3,986E6 | 3,329E8 | 7,844E8 | 1,021E9 | Plec |
| P30999 | Catenin delta-1 OS=Mus musculus GN=Ctnnd1 PE=1 SV=2 - [CTND1_MOUSE] | 0,000E0 | 0,000E0 | 3,219E6 | 1,240E8 | 3,330E8 | 4,168E8 | Ctnnd1 |
| Q8BH64 | EH domain-containing protein 2 OS=Mus musculus GN=Ehd2 PE=1 SV=1 - [EHD2_MOUSE] | 0,000E0 | 0,000E0 | 0,000E0 | 9,033E7 | 1,842E8 | 2,833E8 | Ehd2 |
| P26231 | Catenin alpha-1 OS=Mus musculus GN=Ctnna1 PE=1 SV=1 - [CTNA1_MOUSE] | 0,000E0 | 0,000E0 | 0,000E0 | 2,894E8 | 4,583E8 | 5,492E8 | Ctnna1 |
| P51410 | 60S ribosomal protein L9 OS=Mus musculus GN=Rpl9 PE=2 SV=2 - [RL9_MOUSE] | 0,000E0 | 0,000E0 | 0,000E0 | 1,632E8 | 2,751E8 | 2,248E8 | Rpl9 |
| P62264 | 40S ribosomal protein S14 OS=Mus musculus GN=Rps14 PE=1 SV=3 - [RS14_MOUSE] | 0,000E0 | 0,000E0 | 8,814E6 | 3,426E8 | 1,848E8 | 1,001E9 | Rps14 |
| Q68FD5 | Clathrin heavy chain 1 OS=Mus musculus GN=Cltc PE=1 SV=3 - [CLH1_MOUSE] | 0,000E0 | 0,000E0 | 5,093E6 | 9,943E8 | 1,302E9 | 1,498E9 | Cltc |
| P35979 | 60S ribosomal protein L12 OS=Mus musculus GN=Rpl12 PE=1 SV=2 - [RL12_MOUSE] | 1,348E7 | 0,000E0 | 3,911E6 | 2,635E8 | 5,822E8 | 9,416E8 | Rpl12 |
| P60843 | Eukaryotic initiation factor 4A-I OS=Mus musculus GN=Eif4a1 PE=1 SV=1 - [IF4A1_MOUSE] | 0,000E0 | 0,000E0 | 3,565E5 | 7,572E7 | 1,578E8 | 2,085E8 | Eif4a1 |
| Q91WK0 | Leucine-rich repeat flightless-interacting protein 2 OS=Mus musculus GN=Lrrfip2 PE=1 SV=1 - [LRRF2_MOUSE] | 0,000E0 | 0,000E0 | 1,370E6 | 1,962E8 | 3,153E8 | 2,418E8 | Lrrfip2 |
| P17182 | Alpha-enolase OS=Mus musculus GN=Eno1 PE=1 SV=3 - [ENOA_MOUSE] | 0,000E0 | 0,000E0 | 3,821E6 | 4,461E7 | 1,095E8 | 1,119E8 | Eno1 |
| P84228 | Histone H3.2 OS=Mus musculus GN=Hist1h3b PE=1 SV=2 - [H32_MOUSE] | 0,000E0 | 0,000E0 | 6,923E6 | 5,171E7 | 9,432E7 | 5,068E8 | Hist1h3b |
| P63242 | Eukaryotic translation initiation factor 5A-1 OS=Mus musculus GN=Eif5a PE=1 SV=2 - [IF5A1_MOUSE] | 0,000E0 | 0,000E0 | 0,000E0 | 1,968E7 | 8,325E7 | 9,981E7 | Eif5a |
| P21107 | Tropomyosin alpha-3 chain OS=Mus musculus GN=Tpm3 PE=1 SV=3 - [TPM3_MOUSE] | 1,721E7 | 0,000E0 | 8,255E6 | 1,895E9 | 2,603E9 | 4,298E9 | Tpm3 |
| Q99JY9 | Actin-related protein 3 OS=Mus musculus GN=Actr3 PE=1 SV=3 - [ARP3_MOUSE] | 0,000E0 | 0,000E0 | 2,151E6 | 9,174E7 | 2,808E8 | 3,414E8 | Actr3 |
| Q8BQ30 | Phostensin OS=Mus musculus GN=Ppp1r18 PE=1 SV=1 - [PPR18_MOUSE] | 0,000E0 | 0,000E0 | 0,000E0 | 8,519E7 | 2,448E8 | 5,048E8 | Ppp1r18 |
| E9Q634 | Unconventional myosin-Ie OS=Mus musculus GN=Myo1e PE=1 SV=1 - [MYO1E_MOUSE] | 0,000E0 | 0,000E0 | 2,601E6 | 2,534E8 | 4,588E8 | 8,539E8 | Myo1e |
| P97351 | 40S ribosomal protein S3a OS=Mus musculus GN=Rps3a PE=1 SV=3 - [RS3A_MOUSE] | 0,000E0 | 0,000E0 | 3,947E6 | 2,516E8 | 4,596E8 | 9,725E8 | Rps3a |
| P62245 | 40S ribosomal protein S15a OS=Mus musculus GN=Rps15a PE=1 SV=2 - [RS15A_MOUSE] | 0,000E0 | 0,000E0 | 0,000E0 | 2,970E8 | 5,802E8 | 7,716E8 | Rps15a |
| P23506 | Protein-L-isoaspartate(D-aspartate) O-methyltransferase OS=Mus musculus GN=Pcmt1 PE=1 SV=3 - [PIMT_MOUSE] | 0,000E0 | 0,000E0 | 3,447E5 | 4,734E7 | 5,273E7 | 4,213E7 | Pcmt1 |
| P16546-2 | Isoform 2 of Spectrin alpha chain, non-erythrocytic 1 OS=Mus musculus GN=Sptan1 - [SPTN1_MOUSE] | 0,000E0 | 0,000E0 | 0,000E0 | 7,210E7 | 1,351E8 | 1,963E8 | Sptan1 |
| Q9CVB6 | Actin-related protein 2/3 complex subunit 2 OS=Mus musculus GN=Arpc2 PE=1 SV=3 - [ARPC2_MOUSE] | 0,000E0 | 0,000E0 | 0,000E0 | 9,702E7 | 3,457E8 | 5,493E8 | Arpc2 |
| Q9JKB3-2 | Isoform 2 of Y-box-binding protein 3 OS=Mus musculus GN=Ybx3 - [YBOX3_MOUSE] | 0,000E0 | 0,000E0 | 0,000E0 | 1,701E8 | 3,202E8 | 7,389E8 | Ybx3 |
| P16546 | Spectrin alpha chain, non-erythrocytic 1 OS=Mus musculus GN=Sptan1 PE=1 SV=4 - [SPTN1_MOUSE] | 0,000E0 | 0,000E0 | 0,000E0 | 7,210E7 | 1,351E8 | 1,963E8 | Sptan1 |
| P63017 | Heat shock cognate 71 kDa protein OS=Mus musculus GN=Hspa8 PE=1 SV=1 - [HSP7C_MOUSE] | 3,744E7 | 0,000E0 | 1,198E7 | 6,088E8 | 1,368E9 | 2,139E9 | Hspa8 |
| P61750 | ADP-ribosylation factor 4 OS=Mus musculus GN=Arf4 PE=1 SV=2 - [ARF4_MOUSE] | 0,000E0 | 0,000E0 | 7,116E5 | 2,373E7 | 5,966E7 | 9,349E7 | Arf4 |
| Q9Z0U1 | Tight junction protein ZO-2 OS=Mus musculus GN=Tjp2 PE=1 SV=2 - [ZO2_MOUSE] | 0,000E0 | 0,000E0 | 2,474E6 | 5,557E8 | 1,315E9 | 1,020E9 | Tjp2 |
| O54962 | Barrier-to-autointegration factor OS=Mus musculus GN=Banf1 PE=1 SV=1 - [BAF_MOUSE] | 0,000E0 | 0,000E0 | 0,000E0 | 4,798E6 | 7,358E7 | 1,607E8 | Banf1 |
| Q9JHJ0 | Tropomodulin-3 OS=Mus musculus GN=Tmod3 PE=1 SV=1 - [TMOD3_MOUSE] | 0,000E0 | 0,000E0 | 1,356E6 | 3,738E8 | 7,196E8 | 9,605E8 | Tmod3 |
| P14869 | 60S acidic ribosomal protein P0 OS=Mus musculus GN=Rplp0 PE=1 SV=3 - [RLA0_MOUSE] | 4,904E7 | 0,000E0 | 0,000E0 | 1,401E8 | 3,082E8 | 8,169E8 | Rplp0 |
| Q9CZX8 | 40S ribosomal protein S19 OS=Mus musculus GN=Rps19 PE=1 SV=3 - [RS19_MOUSE] | 0,000E0 | 0,000E0 | 1,201E7 | 2,159E8 | 3,565E8 | 7,484E8 | Rps19 |
| Q64337 | Sequestosome-1 OS=Mus musculus GN=Sqstm1 PE=1 SV=1 - [SQSTM_MOUSE] | 0,000E0 | 0,000E0 | 0,000E0 | 1,741E7 | 1,819E8 | 1,250E8 | Sqstm1 |
| P62806 | Histone H4 OS=Mus musculus GN=Hist1h4a PE=1 SV=2 - [H4_MOUSE] | 3,548E7 | 0,000E0 | 2,100E6 | 4,491E7 | 1,256E8 | 1,525E9 | Hist1h4a |
| P14148 | 60S ribosomal protein L7 OS=Mus musculus GN=Rpl7 PE=1 SV=2 - [RL7_MOUSE] | 0,000E0 | 0,000E0 | 8,463E5 | 1,918E8 | 3,841E8 | 1,158E9 | Rpl7 |
| Q62261 | Spectrin beta chain, non-erythrocytic 1 OS=Mus musculus GN=Sptbn1 PE=1 SV=2 - [SPTB2_MOUSE] | 0,000E0 | 0,000E0 | 2,966E6 | 5,610E7 | 1,701E8 | 2,578E8 | Sptbn1 |
| Q61937 | Nucleophosmin OS=Mus musculus GN=Npm1 PE=1 SV=1 - [NPM_MOUSE] | 0,000E0 | 0,000E0 | 1,406E7 | 4,148E8 | 3,164E8 | 3,566E8 | Npm1 |
| P68254 | 14-3-3 protein theta OS=Mus musculus GN=Ywhaq PE=1 SV=1 - [1433T_MOUSE] | 0,000E0 | 0,000E0 | 5,736E6 | 6,738E7 | 8,293E7 | 1,954E8 | Ywhaq |
| Q9WU78 | Programmed cell death 6-interacting protein OS=Mus musculus GN=Pdcd6ip PE=1 SV=3 - [PDC6I_MOUSE] | 0,000E0 | 0,000E0 | 2,031E5 | 1,063E8 | 2,196E8 | 3,132E8 | Pdcd6ip |
| Q62261-2 | Isoform 2 of Spectrin beta chain, non-erythrocytic 1 OS=Mus musculus GN=Sptbn1 - [SPTB2_MOUSE] | 0,000E0 | 0,000E0 | 2,966E6 | 5,538E7 | 1,701E8 | 2,578E8 | Sptbn1 |
| P61255 | 60S ribosomal protein L26 OS=Mus musculus GN=Rpl26 PE=1 SV=1 - [RL26_MOUSE] | 0,000E0 | 0,000E0 | 0,000E0 | 2,152E8 | 2,489E8 | 7,882E8 | Rpl26 |
| P61982 | 14-3-3 protein gamma OS=Mus musculus GN=Ywhag PE=1 SV=2 - [1433G_MOUSE] | 0,000E0 | 0,000E0 | 0,000E0 | 1,573E8 | 1,548E8 | 1,471E8 | Ywhag |
| P97434-2 | Isoform 2 of Myosin phosphatase Rho-interacting protein OS=Mus musculus GN=Mprip - [MPRIP_MOUSE] | 0,000E0 | 0,000E0 | 0,000E0 | 1,158E8 | 1,846E8 | 2,425E8 | Mprip |
| P62082 | 40S ribosomal protein S7 OS=Mus musculus GN=Rps7 PE=2 SV=1 - [RS7_MOUSE] | 0,000E0 | 0,000E0 | 1,346E7 | 1,913E8 | 1,777E8 | 3,225E8 | Rps7 |
| P62270 | 40S ribosomal protein S18 OS=Mus musculus GN=Rps18 PE=1 SV=3 - [RS18_MOUSE] | 0,000E0 | 0,000E0 | 5,188E6 | 2,852E8 | 2,371E8 | 9,919E8 | Rps18 |
| Q9EP71 | Ankycorbin OS=Mus musculus GN=Rai14 PE=1 SV=1 - [RAI14_MOUSE] | 0,000E0 | 0,000E0 | 0,000E0 | 1,024E8 | 1,644E8 | 2,120E8 | Rai14 |
| Q6ZWV3 | 60S ribosomal protein L10 OS=Mus musculus GN=Rpl10 PE=1 SV=3 - [RL10_MOUSE] | 0,000E0 | 0,000E0 | 3,303E6 | 1,397E8 | 3,848E8 | 7,309E8 | Rpl10 |

| | | | | | | | | |
|---|---|---|---|---|---|---|---|---|
| Q62167 | ATP-dependent RNA helicase DDX3X OS=Mus musculus GN=Ddx3x PE=1 SV=3 - [DDX3X_MOUSE] | 1,704E6 | 0,000E0 | 2,060E6 | 1,659E8 | 3,520E8 | 4,797E8 | Ddx3x |
| Q9CPR4 | 60S ribosomal protein L17 OS=Mus musculus GN=Rpl17 PE=1 SV=3 - [RL17_MOUSE] | 0,000E0 | 0,000E0 | 0,000E0 | 2,699E8 | 5,729E8 | 7,844E8 | Rpl17 |
| Q8CI43 | Myosin light chain 6B OS=Mus musculus GN=Myl6b PE=1 SV=1 - [MYL6B_MOUSE] | 7,368E8 | 0,000E0 | 1,897E8 | 4,242E9 | 5,319E9 | 5,089E9 | Myl6b |
| Q2KN98 | Cytospin-A OS=Mus musculus GN=Specc1l PE=1 SV=1 - [CYTSA_MOUSE] | 0,000E0 | 0,000E0 | 0,000E0 | 1,569E8 | 2,118E8 | 1,865E8 | Specc1l |
| P16858 | Glyceraldehyde-3-phosphate dehydrogenase OS=Mus musculus GN=Gapdh PE=1 SV=2 - [G3P_MOUSE] | 8,286E7 | 0,000E0 | 3,409E6 | 4,568E8 | 9,609E8 | 1,120E9 | Gapdh |
| Q9QYB1 | Chloride intracellular channel protein 4 OS=Mus musculus GN=Clic4 PE=1 SV=3 - [CLIC4_MOUSE] | 0,000E0 | 0,000E0 | 2,973E6 | 1,351E7 | 5,759E7 | 1,109E8 | Clic4 |
| Q8CGP6 | Histone H2A type 1-H OS=Mus musculus GN=Hist1h2ah PE=1 SV=3 - [H2A1H_MOUSE] | 1,458E7 | 9,648E6 | 2,742E7 | 5,055E7 | 9,659E7 | 7,366E8 | Hist1h2ah |
| P47753 | F-actin-capping protein subunit alpha-1 OS=Mus musculus GN=Capza1 PE=1 SV=4 - [CAZA1_MOUSE] | 0,000E0 | 0,000E0 | 0,000E0 | 1,417E8 | 2,442E8 | 2,328E8 | Capza1 |
| P35700 | Peroxiredoxin-1 OS=Mus musculus GN=Prdx1 PE=1 SV=1 - [PRDX1_MOUSE] | 0,000E0 | 0,000E0 | 0,000E0 | 1,399E8 | 2,938E8 | 4,122E8 | Prdx1 |
| P52480 | Pyruvate kinase PKM OS=Mus musculus GN=Pkm PE=1 SV=4 - [KPYM_MOUSE] | 0,000E0 | 0,000E0 | 1,563E7 | 3,242E7 | 6,608E7 | 8,367E7 | Pkm |
| Q9JHL1-2 | Isoform 2 of Na(+)/H(+) exchange regulatory cofactor NHE-RF2 OS=Mus musculus GN=Slc9a3r2 - [NHRF2_MOUSE] | 0,000E0 | 0,000E0 | 0,000E0 | 2,469E7 | 7,123E7 | 9,701E7 | Slc9a3r2 |
| P62137 | Serine/threonine-protein phosphatase PP1-alpha catalytic subunit OS=Mus musculus GN=Ppp1ca PE=1 SV=1 - [PP1A_MOUSE] | 0,000E0 | 0,000E0 | 2,353E6 | 8,782E7 | 1,568E8 | 3,225E8 | Ppp1ca |
| P63276 | 40S ribosomal protein S17 OS=Mus musculus GN=Rps17 PE=1 SV=2 - [RS17_MOUSE] | 0,000E0 | 0,000E0 | 0,000E0 | 1,737E8 | 1,680E8 | 1,554E8 | Rps17 |
| Q9R1K9 | Centrin-2 OS=Mus musculus GN=Cetn2 PE=1 SV=1 - [CETN2_MOUSE] | 0,000E0 | 0,000E0 | 0,000E0 | 5,634E6 | 1,292E7 | 1,475E7 | Cetn2 |
| Q9DBJ1 | Phosphoglycerate mutase 1 OS=Mus musculus GN=Pgam1 PE=1 SV=3 - [PGAM1_MOUSE] | 0,000E0 | 0,000E0 | 0,000E0 | 2,791E7 | 3,348E7 | 7,829E7 | Pgam1 |
| P27659 | 60S ribosomal protein L3 OS=Mus musculus GN=Rpl3 PE=1 SV=3 - [RL3_MOUSE] | 0,000E0 | 0,000E0 | 3,066E6 | 9,362E7 | 2,469E8 | 8,229E8 | Rpl3 |
| P38647 | Stress-70 protein, mitochondrial OS=Mus musculus GN=Hspa9 PE=1 SV=3 - [GRP75_MOUSE] | 6,017E7 | 0,000E0 | 2,130E7 | 2,372E8 | 2,735E8 | 3,258E8 | Hspa9 |
| P68510 | 14-3-3 protein eta OS=Mus musculus GN=Ywhah PE=1 SV=2 - [1433F_MOUSE] | 0,000E0 | 0,000E0 | 7,196E5 | 1,335E8 | 1,491E8 | 1,981E8 | Ywhah |
| P62242 | 40S ribosomal protein S8 OS=Mus musculus GN=Rps8 PE=1 SV=2 - [RS8_MOUSE] | 0,000E0 | 0,000E0 | 1,222E6 | 3,501E8 | 6,842E8 | 3,991E8 | Rps8 |
| P61358 | 60S ribosomal protein L27 OS=Mus musculus GN=Rpl27 PE=1 SV=2 - [RL27_MOUSE] | 0,000E0 | 0,000E0 | 0,000E0 | 3,338E8 | 7,779E8 | 1,012E9 | Rpl27 |
| P08752 | Guanine nucleotide-binding protein G(i) subunit alpha-2 OS=Mus musculus GN=Gnai2 PE=1 SV=5 - [GNAI2_MOUSE] | 0,000E0 | 0,000E0 | 7,794E4 | 5,374E7 | 2,147E8 | 3,711E8 | Gnai2 |
| P62869 | Elongin-B OS=Mus musculus GN=Elob PE=1 SV=1 - [ELOB_MOUSE] | 0,000E0 | 0,000E0 | 0,000E0 | 1,002E7 | 1,568E7 | 3,827E7 | Elob |
| P54823 | Probable ATP-dependent RNA helicase DDX6 OS=Mus musculus GN=Ddx6 PE=1 SV=1 - [DDX6_MOUSE] | 0,000E0 | 0,000E0 | 0,000E0 | 4,259E7 | 5,714E7 | 7,585E7 | Ddx6 |
| P62918 | 60S ribosomal protein L8 OS=Mus musculus GN=Rpl8 PE=1 SV=2 - [RL8_MOUSE] | 0,000E0 | 0,000E0 | 0,000E0 | 1,063E8 | 3,480E8 | 8,066E8 | Rpl8 |
| Q8BKX1 | Brain-specific angiogenesis inhibitor 1-associated protein 2 OS=Mus musculus GN=Baiap2 PE=1 SV=2 - [BAIP2_MOUSE] | 0,000E0 | 0,000E0 | 0,000E0 | 1,514E7 | 3,729E7 | 7,650E7 | Baiap2 |
| O88569 | Heterogeneous nuclear ribonucleoproteins A2/B1 OS=Mus musculus GN=Hnrnpa2b1 PE=1 SV=2 - [ROA2_MOUSE] | 0,000E0 | 0,000E0 | 0,000E0 | 4,976E7 | 4,955E7 | 1,864E8 | Hnrnpa2b1 |
| O09044 | Synaptosomal-associated protein 23 OS=Mus musculus GN=Snap23 PE=1 SV=1 - [SNP23_MOUSE] | 0,000E0 | 0,000E0 | 0,000E0 | 1,280E7 | 4,024E7 | 8,525E7 | Snap23 |
| P61205 | ADP-ribosylation factor 3 OS=Mus musculus GN=Arf3 PE=2 SV=2 - [ARF3_MOUSE] | 0,000E0 | 0,000E0 | 0,000E0 | 5,665E7 | 1,137E8 | 9,176E7 | Arf3 |
| Q02248 | Catenin beta-1 OS=Mus musculus GN=Ctnnb1 PE=1 SV=1 - [CTNB1_MOUSE] | 0,000E0 | 0,000E0 | 3,246E6 | 3,082E8 | 4,762E8 | 5,489E8 | Ctnnb1 |
| Q5SYD0 | Unconventional myosin-Id OS=Mus musculus GN=Myo1d PE=1 SV=1 - [MYO1D_MOUSE] | 2,735E6 | 0,000E0 | 6,818E6 | 1,427E8 | 3,484E8 | 8,994E8 | Myo1d |
| P62281 | 40S ribosomal protein S11 OS=Mus musculus GN=Rps11 PE=1 SV=3 - [RS11_MOUSE] | 0,000E0 | 0,000E0 | 4,261E6 | 1,330E8 | 3,233E8 | 4,583E8 | Rps11 |
| Q64523 | Histone H2A type 2-C OS=Mus musculus GN=Hist2h2ac PE=1 SV=3 - [H2A2C_MOUSE] | 0,000E0 | 0,000E0 | 1,828E7 | 4,055E7 | 8,789E7 | 7,541E8 | Hist2h2ac |
| P83940 | Elongin-C OS=Mus musculus GN=Eloc PE=1 SV=1 - [ELOC_MOUSE] | 0,000E0 | 0,000E0 | 0,000E0 | 1,346E7 | 2,038E7 | 2,708E7 | Eloc |
| P29391 | Ferritin light chain 1 OS=Mus musculus GN=Ftl1 PE=1 SV=2 - [FRIL1_MOUSE] | 0,000E0 | 0,000E0 | 0,000E0 | 2,343E7 | 2,971E7 | 5,841E7 | Ftl1 |
| P97855 | Ras GTPase-activating protein-binding protein 1 OS=Mus musculus GN=G3bp1 PE=1 SV=1 - [G3BP1_MOUSE] | 8,322E6 | 0,000E0 | 0,000E0 | 1,364E8 | 2,011E8 | 6,557E8 | G3bp1 |
| P68037 | Ubiquitin-conjugating enzyme E2 L3 OS=Mus musculus GN=Ube2l3 PE=1 SV=1 - [UB2L3_MOUSE] | 0,000E0 | 0,000E0 | 0,000E0 | 8,416E6 | 7,511E6 | 5,615E6 | Ube2l3 |
| Q9D8E6 | 60S ribosomal protein L4 OS=Mus musculus GN=Rpl4 PE=1 SV=3 - [RL4_MOUSE] | 0,000E0 | 0,000E0 | 1,238E6 | 2,018E8 | 3,842E8 | 8,158E8 | Rpl4 |
| P68372 | Tubulin beta-4B chain OS=Mus musculus GN=Tubb4b PE=1 SV=1 - [TBB4B_MOUSE] | 7,924E6 | 0,000E0 | 5,484E6 | 1,362E8 | 2,937E8 | 3,976E8 | Tubb4b |
| Q8BT07 | Centrosomal protein of 55 kDa OS=Mus musculus GN=Cep55 PE=1 SV=2 - [CEP55_MOUSE] | 0,000E0 | 0,000E0 | 0,000E0 | 7,690E7 | 1,616E8 | 2,430E8 | Cep55 |
| P17426-2 | Isoform B of AP-2 complex subunit alpha-1 OS=Mus musculus GN=Ap2a1 - [AP2A1_MOUSE] | 0,000E0 | 0,000E0 | 0,000E0 | 1,427E8 | 2,388E8 | 3,233E8 | Ap2a1 |
| P98078 | Disabled homolog 2 OS=Mus musculus GN=Dab2 PE=1 SV=2 - [DAB2_MOUSE] | 0,000E0 | 0,000E0 | 0,000E0 | 2,629E8 | 3,975E8 | 6,565E8 | Dab2 |
| P12970 | 60S ribosomal protein L7a OS=Mus musculus GN=Rpl7a PE=1 SV=2 - [RL7A_MOUSE] | 0,000E0 | 0,000E0 | 9,615E5 | 2,438E8 | 4,138E8 | 1,418E9 | Rpl7a |
| P62983 | Ubiquitin-40S ribosomal protein S27a OS=Mus musculus GN=Rps27a PE=1 SV=2 - [RS27A_MOUSE] | 2,631E7 | 0,000E0 | 2,882E7 | 7,431E8 | 9,040E8 | 9,846E7 | Rps27a |
| Q8BFZ3 | Beta-actin-like protein 2 OS=Mus musculus GN=Actbl2 PE=1 SV=1 - [ACTBL_MOUSE] | 6,609E8 | 0,000E0 | 1,502E8 | 5,101E9 | 1,354E10 | 1,337E10 | Actbl2 |
| Q9CY58-2 | Isoform 2 of Plasminogen activator inhibitor 1 RNA-binding protein OS=Mus musculus GN=Serbp1 - [PAIRB_MOUSE] | 0,000E0 | 0,000E0 | 1,047E7 | 1,214E8 | 2,512E8 | 5,693E8 | Serbp1 |
| P20029 | 78 kDa glucose-regulated protein OS=Mus musculus GN=Hspa5 PE=1 SV=3 - [GRP78_MOUSE] | 6,100E7 | 0,000E0 | 1,220E7 | 1,944E8 | 2,001E8 | 4,070E8 | Hspa5 |
| Q61584-3 | Isoform B of Fragile X mental retardation syndrome-related protein 1 OS=Mus musculus GN=Fxr1 - [FXR1_MOUSE] | 0,000E0 | 0,000E0 | 3,780E5 | 9,430E7 | 1,716E8 | 3,795E8 | Fxr1 |
| P62960 | Nuclease-sensitive element-binding protein 1 OS=Mus musculus GN=Ybx1 PE=1 SV=3 - [YBOX1_MOUSE] | 1,609E7 | 0,000E0 | 8,242E5 | 2,920E8 | 6,613E8 | 1,151E9 | Ybx1 |
| P61161 | Actin-related protein 2 OS=Mus musculus GN=Actr2 PE=1 SV=1 - [ARP2_MOUSE] | 0,000E0 | 0,000E0 | 2,837E6 | 1,061E8 | 2,855E8 | 5,562E8 | Actr2 |
| P68033 | Actin, alpha cardiac muscle 1 OS=Mus musculus GN=Actc1 PE=1 SV=1 - [ACTC_MOUSE] | 7,491E8 | 0,000E0 | 1,763E8 | 9,328E9 | 2,030E10 | 1,500E10 | Actc1 |
| Q9CQ19 | Myosin regulatory light polypeptide 9 OS=Mus musculus GN=Myl9 PE=1 SV=3 - [MYL9_MOUSE] | 1,440E8 | 0,000E0 | 1,331E8 | 5,016E9 | 8,498E9 | 1,152E10 | Myl9 |
| P67871 | Casein kinase II subunit beta OS=Mus musculus GN=Csnk2b PE=1 SV=1 - [CSK2B_MOUSE] | 0,000E0 | 0,000E0 | 0,000E0 | 8,717E6 | 2,380E7 | 3,732E7 | Csnk2b |
| P80318 | T-complex protein 1 subunit gamma OS=Mus musculus GN=Cct3 PE=1 SV=1 - [TCPG_MOUSE] | 0,000E0 | 0,000E0 | 0,000E0 | 2,080E7 | 4,257E7 | 4,939E7 | Cct3 |
| P14206 | 40S ribosomal protein SA OS=Mus musculus GN=Rpsa PE=1 SV=4 - [RSSA_MOUSE] | 0,000E0 | 0,000E0 | 0,000E0 | 4,752E8 | 8,327E8 | 1,754E9 | Rpsa |
| P29341 | Polyadenylate-binding protein 1 OS=Mus musculus GN=Pabpc1 PE=1 SV=2 - [PABP1_MOUSE] | 0,000E0 | 0,000E0 | 0,000E0 | 1,375E8 | 2,779E8 | 5,690E8 | Pabpc1 |
| Q9JHL1 | Na(+)/H(+) exchange regulatory cofactor NHE-RF2 OS=Mus musculus GN=Slc9a3r2 PE=1 SV=2 - [NHRF2_MOUSE] | 0,000E0 | 0,000E0 | 0,000E0 | 2,469E7 | 7,123E7 | 9,769E7 | Slc9a3r2 |
| P62751 | 60S ribosomal protein L23a OS=Mus musculus GN=Rpl23a PE=1 SV=1 - [RL23A_MOUSE] | 0,000E0 | 0,000E0 | 1,154E7 | 2,928E8 | 5,583E8 | 1,436E9 | Rpl23a |
| P51881 | ADP/ATP translocase 2 OS=Mus musculus GN=Slc25a5 PE=1 SV=3 - [ADT2_MOUSE] | 3,742E5 | 0,000E0 | 1,161E7 | 1,170E8 | 2,168E8 | 2,044E8 | Slc25a5 |
| Q61033 | Lamina-associated polypeptide 2, isoforms alpha/zeta OS=Mus musculus GN=Tmpo PE=1 SV=4 - [LAP2A_MOUSE] | 0,000E0 | 0,000E0 | 7,435E5 | 8,448E7 | 2,430E8 | 4,506E8 | Tmpo |
| Q9D8B3 | Charged multivesicular body protein 4b OS=Mus musculus GN=Chmp4b PE=1 SV=2 - [CHM4B_MOUSE] | 0,000E0 | 0,000E0 | 0,000E0 | 1,016E8 | 2,077E8 | 3,188E8 | Chmp4b |
| P62301 | 40S ribosomal protein S13 OS=Mus musculus GN=Rps13 PE=1 SV=2 - [RS13_MOUSE] | 0,000E0 | 0,000E0 | 3,325E6 | 1,528E8 | 3,612E8 | 7,062E8 | Rps13 |
| P99024 | Tubulin beta-5 chain OS=Mus musculus GN=Tubb5 PE=1 SV=1 - [TBB5_MOUSE] | 7,924E6 | 0,000E0 | 5,484E6 | 1,705E8 | 4,630E8 | 3,976E8 | Tubb5 |
| P84104-2 | Isoform Short of Serine/arginine-rich splicing factor 3 OS=Mus musculus GN=Srsf3 - [SRSF3_MOUSE] | 0,000E0 | 0,000E0 | 0,000E0 | 6,099E7 | 7,007E7 | 1,265E8 | Srsf3 |
| Q99K01 | Pyridoxal-dependent decarboxylase domain-containing protein 1 OS=Mus musculus GN=Pdxdc1 PE=1 SV=2 - [PDXD1_MOUSE] | 0,000E0 | 0,000E0 | 0,000E0 | 1,205E8 | 9,430E7 | 9,452E7 | Pdxdc1 |
| P63325 | 40S ribosomal protein S10 OS=Mus musculus GN=Rps10 PE=1 SV=1 - [RS10_MOUSE] | 0,000E0 | 0,000E0 | 0,000E0 | 1,503E8 | 1,726E8 | 3,338E8 | Rps10 |
| P62141 | Serine/threonine-protein phosphatase PP1-beta catalytic subunit OS=Mus musculus GN=Ppp1cb PE=1 SV=3 - [PP1B_MOUSE] | 0,000E0 | 0,000E0 | 2,353E6 | 5,939E7 | 1,204E8 | 2,192E8 | Ppp1cb |
| P47911 | 60S ribosomal protein L6 OS=Mus musculus GN=Rpl6 PE=1 SV=3 - [RL6_MOUSE] | 0,000E0 | 0,000E0 | 2,927E6 | 1,822E8 | 4,127E8 | 9,566E8 | Rpl6 |
| Q99K01-2 | Isoform 2 of Pyridoxal-dependent decarboxylase domain-containing protein 1 OS=Mus musculus GN=Pdxdc1 - [PDXD1_MOUSE] | 0,000E0 | 0,000E0 | 0,000E0 | 1,547E8 | 9,430E7 | 1,446E8 | Pdxdc1 |
| P17742 | Peptidyl-prolyl cis-trans isomerase A OS=Mus musculus GN=Ppia PE=1 SV=2 - [PPIA_MOUSE] | 0,000E0 | 0,000E0 | 9,673E5 | 1,240E8 | 2,384E8 | 2,885E8 | Ppia |
| Q6ZWN5 | 40S ribosomal protein S9 OS=Mus musculus GN=Rps9 PE=1 SV=3 - [RS9_MOUSE] | 0,000E0 | 0,000E0 | 4,221E6 | 1,742E8 | 2,236E8 | 4,800E8 | Rps9 |
| Q3U0S6 | Ras-interacting protein 1 OS=Mus musculus GN=Rasip1 PE=1 SV=3 - [RAIN_MOUSE] | 5,784E6 | 0,000E0 | 1,974E6 | 2,222E8 | 5,526E8 | 1,186E9 | Rasip1 |
| P68373 | Tubulin alpha-1C chain OS=Mus musculus GN=Tuba1c PE=1 SV=1 - [TBA1C_MOUSE] | 1,425E7 | 0,000E0 | 7,585E6 | 7,937E7 | 4,307E8 | 4,357E8 | Tuba1c |
| P63037 | DnaJ homolog subfamily A member 1 OS=Mus musculus GN=Dnaja1 PE=1 SV=1 - [DNJA1_MOUSE] | 0,000E0 | 0,000E0 | 0,000E0 | 2,998E7 | 4,294E7 | 8,657E7 | Dnaja1 |
| Q9JKB3 | Y-box-binding protein 3 OS=Mus musculus GN=Ybx3 PE=1 SV=2 - [YBOX3_MOUSE] | 0,000E0 | 0,000E0 | 0,000E0 | 1,701E8 | 3,202E8 | 7,389E8 | Ybx3 |
| Q60598 | Src substrate cortactin OS=Mus musculus GN=Cttn PE=1 SV=2 - [SRC8_MOUSE] | 0,000E0 | 0,000E0 | 0,000E0 | 9,379E7 | 1,714E8 | 2,807E8 | Cttn |
| P07356 | Annexin A2 OS=Mus musculus GN=Anxa2 PE=1 SV=2 - [ANXA2_MOUSE] | 0,000E0 | 0,000E0 | 1,418E6 | 4,466E7 | 1,027E8 | 1,429E8 | Anxa2 |
| Q9JJI8 | 60S ribosomal protein L38 OS=Mus musculus GN=Rpl38 PE=1 SV=3 - [RL38_MOUSE] | 0,000E0 | 0,000E0 | 0,000E0 | 6,069E7 | 1,067E8 | 1,839E8 | Rpl38 |
| **Q8K371** | **Angiomotin-like protein 2 OS=Mus musculus GN=Amotl2 PE=1 SV=1 - [AMOL2_MOUSE]** | **0,000E0** | **0,000E0** | **0,000E0** | **2,745E8** | **2,477E8** | **2,037E8** | **Amotl2** |
| P10126 | Elongation factor 1-alpha 1 OS=Mus musculus GN=Eef1a1 PE=1 SV=3 - [EF1A1_MOUSE] | 4,552E6 | 0,000E0 | 2,176E6 | 4,168E8 | 7,622E8 | 9,864E8 | Eef1a1 |
| Q9D1G1 | Ras-related protein Rab-1B OS=Mus musculus GN=Rab1b PE=1 SV=1 - [RAB1B_MOUSE] | 0,000E0 | 0,000E0 | 3,456E5 | 6,625E7 | 1,481E8 | 2,129E8 | Rab1b |
| P42208 | Septin-2 OS=Mus musculus GN=Sept2 PE=1 SV=2 - [SEPT2_MOUSE] | 0,000E0 | 0,000E0 | 0,000E0 | 2,179E7 | 4,498E7 | 2,781E7 | SEPT2 |
| Q9JJ28 | Protein flightless-1 homolog OS=Mus musculus GN=Flii PE=1 SV=1 - [FLII_MOUSE] | 0,000E0 | 0,000E0 | 0,000E0 | 1,469E8 | 2,788E8 | 3,289E8 | Flii |
| Q99104 | Unconventional myosin-Va OS=Mus musculus GN=Myo5a PE=1 SV=2 - [MYO5A_MOUSE] | 2,735E6 | 0,000E0 | 6,683E5 | 8,479E7 | 1,885E8 | 5,972E8 | Myo5a |
| Q9DBG3 | AP-2 complex subunit beta OS=Mus musculus GN=Ap2b1 PE=1 SV=1 - [AP2B1_MOUSE] | 0,000E0 | 0,000E0 | 0,000E0 | 2,544E8 | 4,341E8 | 4,493E8 | Ap2b1 |
| P63323 | 40S ribosomal protein S12 OS=Mus musculus GN=Rps12 PE=1 SV=2 - [RS12_MOUSE] | 0,000E0 | 0,000E0 | 1,701E5 | 1,409E8 | 3,121E8 | 7,002E8 | Rps12 |
| Q8K4L3-3 | Isoform 2 of Supervillin OS=Mus musculus GN=Svil - [SVIL_MOUSE] | 0,000E0 | 0,000E0 | 4,058E5 | 9,909E7 | 1,704E8 | 2,919E8 | Svil |
| P62858 | 40S ribosomal protein S28 OS=Mus musculus GN=Rps28 PE=1 SV=1 - [RS28_MOUSE] | 1,014E7 | 0,000E0 | 9,284E6 | 5,536E7 | 3,534E7 | 2,064E7 | Rps28 |
| Q4ACU6 | SH3 and multiple ankyrin repeat domains protein 3 OS=Mus musculus GN=Shank3 PE=1 SV=3 - [SHAN3_MOUSE] | 0,000E0 | 0,000E0 | 0,000E0 | 7,759E7 | 1,981E8 | 3,177E8 | Shank3 |
| Q80X90 | Filamin-B OS=Mus musculus GN=Flnb PE=1 SV=3 - [FLNB_MOUSE] | 0,000E0 | 0,000E0 | 1,416E6 | 3,471E7 | 1,033E8 | 1,361E8 | Flnb |
| Q99LF4 | tRNA-splicing ligase RtcB homolog OS=Mus musculus GN=Rtcb PE=1 SV=1 - [RTCB_MOUSE] | 0,000E0 | 0,000E0 | 0,000E0 | 2,398E7 | 5,221E7 | 5,902E7 | Rtcb |
| Q8JZQ9 | Eukaryotic translation initiation factor 3 subunit B OS=Mus musculus GN=Eif3b PE=1 SV=1 - [EIF3B_MOUSE] | 0,000E0 | 0,000E0 | 0,000E0 | 3,702E7 | 6,414E7 | 1,069E8 | Eif3b |
| P68134 | Actin, alpha skeletal muscle OS=Mus musculus GN=Acta1 PE=1 SV=1 - [ACTS_MOUSE] | 7,491E8 | 0,000E0 | 1,507E8 | 9,328E9 | 2,030E10 | 1,500E10 | Acta1 |
| Q60737 | Casein kinase II subunit alpha OS=Mus musculus GN=Csnk2a1 PE=1 SV=2 - [CSK21_MOUSE] | 0,000E0 | 0,000E0 | 0,000E0 | 3,069E7 | 3,592E7 | 4,701E7 | Csnk2a1 |

| | | | | | | | | |
|---|---|---|---|---|---|---|---|---|
| P53026 | 60S ribosomal protein L10a OS=Mus musculus GN=Rpl10a PE=1 SV=3 - [RL10A_MOUSE] | 0,000E0 | 0,000E0 | 1,121E6 | 2,194E8 | 4,647E8 | 7,238E8 | Rpl10a |
| Q9CZM2 | 60S ribosomal protein L15 OS=Mus musculus GN=Rpl15 PE=2 SV=4 - [RL15_MOUSE] | 0,000E0 | 0,000E0 | 0,000E0 | 1,410E8 | 2,356E8 | 6,112E8 | Rpl15 |
| Q9JMH9 | Unconventional myosin-XVIIIa OS=Mus musculus GN=Myo18a PE=1 SV=2 - [MY18A_MOUSE] | 0,000E0 | 0,000E0 | 0,000E0 | 6,613E7 | 1,364E8 | 1,946E8 | Myo18a |
| Q04736 | Tyrosine-protein kinase Yes OS=Mus musculus GN=Yes1 PE=1 SV=3 - [YES_MOUSE] | 0,000E0 | 0,000E0 | 2,114E5 | 2,150E7 | 6,377E7 | 7,664E7 | Yes1 |
| P60867 | 40S ribosomal protein S20 OS=Mus musculus GN=Rps20 PE=1 SV=1 - [RS20_MOUSE] | 0,000E0 | 0,000E0 | 1,652E6 | 4,106E8 | 8,724E8 | 1,511E9 | Rps20 |
| Q3UZ39 | Leucine-rich repeat flightless-interacting protein 1 OS=Mus musculus GN=Lrrfip1 PE=1 SV=2 - [LRRF1_MOUSE] | 0,000E0 | 0,000E0 | 0,000E0 | 2,787E7 | 7,631E7 | 9,605E7 | Lrrfip1 |
| P62843 | 40S ribosomal protein S15 OS=Mus musculus GN=Rps15 PE=1 SV=2 - [RS15_MOUSE] | 0,000E0 | 0,000E0 | 1,279E6 | 2,102E8 | 3,951E8 | 1,292E9 | Rps15 |
| Q8BP67 | 60S ribosomal protein L24 OS=Mus musculus GN=Rpl24 PE=1 SV=2 - [RL24_MOUSE] | 0,000E0 | 0,000E0 | 0,000E0 | 1,450E8 | 3,722E8 | 2,636E8 | Rpl24 |
| O55142 | 60S ribosomal protein L35a OS=Mus musculus GN=Rpl35a PE=1 SV=2 - [RL35A_MOUSE] | 0,000E0 | 0,000E0 | 6,178E6 | 1,554E8 | 1,967E8 | 3,439E8 | Rpl35a |
| Q9Z0F4 | Calcium and integrin-binding protein 1 OS=Mus musculus GN=Cib1 PE=1 SV=3 - [CIB1_MOUSE] | 0,000E0 | 0,000E0 | 0,000E0 | 7,585E6 | 1,559E7 | 2,096E7 | Cib1 |
| Q64331 | Unconventional myosin-VI OS=Mus musculus GN=Myo6 PE=1 SV=1 - [MYO6_MOUSE] | 0,000E0 | 0,000E0 | 3,053E6 | 3,372E8 | 4,575E8 | 5,465E8 | Myo6 |
| Q61584-7 | Isoform G of Fragile X mental retardation syndrome-related protein 1 OS=Mus musculus GN=Fxr1 - [FXR1_MOUSE] | 0,000E0 | 0,000E0 | 3,780E5 | 8,816E7 | 1,716E8 | 3,689E8 | Fxr1 |
| P84084 | ADP-ribosylation factor 5 OS=Mus musculus GN=Arf5 PE=1 SV=2 - [ARF5_MOUSE] | 0,000E0 | 0,000E0 | 0,000E0 | 2,539E7 | 3,693E7 | 5,111E7 | Arf5 |
| Q99KW3-1 | Isoform 1 of TRIO and F-actin-binding protein OS=Mus musculus GN=Triobp - [TARA_MOUSE] | 0,000E0 | 0,000E0 | 0,000E0 | 2,693E7 | 5,169E7 | 6,085E7 | Triobp |
| P25444 | 40S ribosomal protein S2 OS=Mus musculus GN=Rps2 PE=1 SV=3 - [RS2_MOUSE] | 0,000E0 | 0,000E0 | 8,204E6 | 1,941E8 | 4,589E8 | 1,303E9 | Rps2 |
| Q9JIW9 | Ras-related protein Ral-B OS=Mus musculus GN=Ralb PE=1 SV=1 - [RALB_MOUSE] | 0,000E0 | 0,000E0 | 0,000E0 | 1,527E7 | 3,698E7 | 7,532E7 | Ralb |
| P11499 | Heat shock protein HSP 90-beta OS=Mus musculus GN=Hsp90ab1 PE=1 SV=3 - [HS90B_MOUSE] | 0,000E0 | 0,000E0 | 1,205E7 | 1,597E8 | 3,718E8 | 5,243E8 | Hsp90ab1 |
| P62821 | Ras-related protein Rab-1A OS=Mus musculus GN=Rab1A PE=1 SV=3 - [RAB1A_MOUSE] | 0,000E0 | 0,000E0 | 3,456E5 | 6,625E7 | 1,481E8 | 2,129E8 | Rab1A |
| P09528 | Ferritin heavy chain OS=Mus musculus GN=Fth1 PE=1 SV=2 - [FRIH_MOUSE] | 0,000E0 | 0,000E0 | 0,000E0 | 2,738E7 | 3,928E7 | 7,222E7 | Fth1 |
| P68368 | Tubulin alpha-4A chain OS=Mus musculus GN=Tuba4a PE=1 SV=1 - [TBA4A_MOUSE] | 1,425E7 | 0,000E0 | 1,199E6 | 7,937E7 | 3,125E8 | 2,375E8 | Tuba4a |
| Q61599 | Rho GDP-dissociation inhibitor 2 OS=Mus musculus GN=Arhgdib PE=1 SV=3 - [GDIR2_MOUSE] | 0,000E0 | 0,000E0 | 0,000E0 | 1,438E7 | 1,430E7 | 2,190E7 | Arhgdib |
| Q9CQR2 | 40S ribosomal protein S21 OS=Mus musculus GN=Rps21 PE=1 SV=1 - [RS21_MOUSE] | 0,000E0 | 0,000E0 | 0,000E0 | 8,197E7 | 1,348E8 | 2,985E8 | Rps21 |
| P19253 | 60S ribosomal protein L13a OS=Mus musculus GN=Rpl13a PE=1 SV=4 - [RL13A_MOUSE] | 0,000E0 | 0,000E0 | 0,000E0 | 1,611E8 | 2,253E8 | 7,323E8 | Rpl13a |
| Q9WUM4 | Coronin-1C OS=Mus musculus GN=Coro1c PE=1 SV=2 - [COR1C_MOUSE] | 0,000E0 | 0,000E0 | 1,502E6 | 1,097E8 | 1,440E8 | 1,489E8 | Coro1c |
| P80317 | T-complex protein 1 subunit zeta OS=Mus musculus GN=Cct6a PE=1 SV=3 - [TCPZ_MOUSE] | 0,000E0 | 0,000E0 | 0,000E0 | 3,232E7 | 5,697E7 | 4,656E7 | Cct6a |
| Q5FWH6 | Rho guanine nucleotide exchange factor 15 OS=Mus musculus GN=Arhgef15 PE=1 SV=1 - [ARHGF_MOUSE] | 0,000E0 | 0,000E0 | 7,429E5 | 4,489E7 | 1,085E8 | 1,937E8 | Arhgef15 |
| P99029-2 | Isoform Cytoplasmic+peroxisomal of Peroxiredoxin-5, mitochondrial OS=Mus musculus GN=Prdx5 - [PRDX5_MOUSE] | 0,000E0 | 0,000E0 | 0,000E0 | 2,649E6 | 1,180E7 | 1,535E7 | Prdx5 |
| Q8BMK4 | Cytoskeleton-associated protein 4 OS=Mus musculus GN=Ckap4 PE=1 SV=2 - [CKAP4_MOUSE] | 0,000E0 | 0,000E0 | 0,000E0 | 3,904E7 | 8,581E7 | 1,947E8 | Ckap4 |
| Q4ACU6-10 | Isoform 9 of SH3 and multiple ankyrin repeat domains protein 3 OS=Mus musculus GN=Shank3 - [SHAN3_MOUSE] | 0,000E0 | 0,000E0 | 0,000E0 | 6,440E7 | 1,816E8 | 2,818E8 | Shank3 |
| Q922F4 | Tubulin beta-6 chain OS=Mus musculus GN=Tubb6 PE=1 SV=1 - [TBB6_MOUSE] | 0,000E0 | 0,000E0 | 4,329E6 | 2,256E7 | 9,625E7 | 1,470E8 | Tubb6 |
| Q7TPV4 | Myb-binding protein 1A OS=Mus musculus GN=Mybbp1a PE=1 SV=2 - [MBB1A_MOUSE] | 0,000E0 | 0,000E0 | 8,014E5 | 5,727E7 | 1,472E8 | 2,763E8 | Mybbp1a |
| P09411 | Phosphoglycerate kinase 1 OS=Mus musculus GN=Pgk1 PE=1 SV=4 - [PGK1_MOUSE] | 0,000E0 | 0,000E0 | 4,676E6 | 3,787E7 | 9,668E7 | 1,492E8 | Pgk1 |
| O09167 | 60S ribosomal protein L21 OS=Mus musculus GN=Rpl21 PE=1 SV=3 - [RL21_MOUSE] | 0,000E0 | 0,000E0 | 4,561E5 | 1,193E8 | 2,589E8 | 2,638E7 | Rpl21 |
| Q02257 | Junction plakoglobin OS=Mus musculus GN=Jup PE=1 SV=3 - [PLAK_MOUSE] | 0,000E0 | 0,000E0 | 6,011E6 | 7,841E7 | 8,065E7 | 6,165E7 | Jup |
| Q8BX10 | Serine/threonine-protein phosphatase PGAM5, mitochondrial OS=Mus musculus GN=Pgam5 PE=1 SV=1 - [PGAM5_MOUSE] | 0,000E0 | 0,000E0 | 0,000E0 | 1,269E7 | 3,446E7 | 5,267E7 | Pgam5 |
| Q63918 | Serum deprivation-response protein OS=Mus musculus GN=Sdpr PE=1 SV=3 - [SDPR_MOUSE] | 0,000E0 | 0,000E0 | 0,000E0 | 1,519E7 | 3,899E7 | 3,275E7 | Sdpr |
| Q8BX10-2 | Isoform 2 of Serine/threonine-protein phosphatase PGAM5, mitochondrial OS=Mus musculus GN=Pgam5 - [PGAM5_MOUSE] | 0,000E0 | 0,000E0 | 0,000E0 | 1,269E7 | 3,446E7 | 5,267E7 | Pgam5 |
| Q99PL5 | Ribosome-binding protein 1 OS=Mus musculus GN=Rrbp1 PE=1 SV=2 - [RRBP1_MOUSE] | 0,000E0 | 0,000E0 | 7,835E5 | 3,909E7 | 7,199E7 | 1,697E8 | Rrbp1 |
| Q6PDM2 | Serine/arginine-rich splicing factor 1 OS=Mus musculus GN=Srsf1 PE=1 SV=3 - [SRSF1_MOUSE] | 0,000E0 | 0,000E0 | 0,000E0 | 5,573E7 | 1,008E8 | 1,991E8 | Srsf1 |
| Q9CQ69 | Cytochrome b-c1 complex subunit 8 OS=Mus musculus GN=Uqcrq PE=1 SV=3 - [QCR8_MOUSE] | 0,000E0 | 0,000E0 | 0,000E0 | 1,865E7 | 2,823E7 | 7,016E7 | Uqcrq |
| Q64152 | Transcription factor BTF3 OS=Mus musculus GN=Btf3 PE=1 SV=3 - [BTF3_MOUSE] | 0,000E0 | 0,000E0 | 0,000E0 | 4,684E6 | 2,488E7 | 7,617E7 | Btf3 |
| Q9R0P3 | S-formylglutathione hydrolase OS=Mus musculus GN=Esd PE=1 SV=1 - [ESTD_MOUSE] | 0,000E0 | 0,000E0 | 0,000E0 | 6,329E6 | 2,115E7 | 3,230E7 | Esd |
| Q01853 | Transitional endoplasmic reticulum ATPase OS=Mus musculus GN=Vcp PE=1 SV=4 - [TERA_MOUSE] | 0,000E0 | 0,000E0 | 9,011E6 | 2,354E7 | 4,093E7 | 7,549E7 | Vcp |
| Q9JJL8 | Serine--tRNA ligase, mitochondrial OS=Mus musculus GN=Sars2 PE=1 SV=2 - [SYSM_MOUSE] | 0,000E0 | 0,000E0 | 0,000E0 | 8,537E7 | 1,164E8 | 6,457E7 | Sars2 |
| P06151 | L-lactate dehydrogenase A chain OS=Mus musculus GN=Ldha PE=1 SV=3 - [LDHA_MOUSE] | 0,000E0 | 0,000E0 | 1,621E6 | 1,657E7 | 2,224E7 | 3,302E7 | Ldha |
| Q7TSJ2-3 | Isoform 3 of Microtubule-associated protein 6 OS=Mus musculus GN=Map6 - [MAP6_MOUSE] | 0,000E0 | 0,000E0 | 0,000E0 | 4,323E6 | 1,328E7 | 1,689E7 | Map6 |
| Q9ERG0 | LIM domain and actin-binding protein 1 OS=Mus musculus GN=Lima1 PE=1 SV=3 - [LIMA1_MOUSE] | 0,000E0 | 0,000E0 | 2,209E6 | 5,855E7 | 1,735E8 | 3,073E8 | Lima1 |
| Q7TMM9 | Tubulin beta-2A chain OS=Mus musculus GN=Tubb2a PE=1 SV=1 - [TBB2A_MOUSE] | 7,924E6 | 0,000E0 | 5,484E6 | 1,259E8 | 3,918E8 | 3,976E8 | Tubb2a |
| O54724 | Polymerase I and transcript release factor OS=Mus musculus GN=Ptrf PE=1 SV=1 - [PTRF_MOUSE] | 0,000E0 | 0,000E0 | 0,000E0 | 9,998E7 | 2,089E8 | 4,354E8 | Ptrf |
| P62880 | Guanine nucleotide-binding protein G(I)/G(S)/G(T) subunit beta-2 OS=Mus musculus GN=Gnb2 PE=1 SV=3 - [GBB2_MOUSE] | 0,000E0 | 0,000E0 | 2,238E6 | 8,151E7 | 1,019E8 | 1,458E8 | Gnb2 |
| O54931-2 | Isoform 2 of A-kinase anchor protein 2 OS=Mus musculus GN=Akap2 - [AKAP2_MOUSE] | 0,000E0 | 0,000E0 | 0,000E0 | 7,483E7 | 1,794E8 | 2,695E8 | Akap2 |
| P05064 | Fructose-bisphosphate aldolase A OS=Mus musculus GN=Aldoa PE=1 SV=2 - [ALDOA_MOUSE] | 0,000E0 | 0,000E0 | 3,110E6 | 1,479E7 | 4,467E7 | 6,221E7 | Aldoa |
| Q8K4L3 | Supervillin OS=Mus musculus GN=Svil PE=1 SV=1 - [SVIL_MOUSE] | 0,000E0 | 0,000E0 | 4,058E5 | 9,909E7 | 1,704E8 | 2,919E8 | Svil |
| Q9JKF1 | Ras GTPase-activating-like protein IQGAP1 OS=Mus musculus GN=Iqgap1 PE=1 SV=2 - [IQGA1_MOUSE] | 0,000E0 | 0,000E0 | 0,000E0 | 3,865E7 | 1,035E8 | 1,009E8 | Iqgap1 |
| Q8K0D0 | Cyclin-dependent kinase 17 OS=Mus musculus GN=Cdk17 PE=1 SV=2 - [CDK17_MOUSE] | 0,000E0 | 0,000E0 | 0,000E0 | 2,189E7 | 5,107E7 | 7,426E7 | Cdk17 |
| P70372 | ELAV-like protein 1 OS=Mus musculus GN=Elavl1 PE=1 SV=2 - [ELAV1_MOUSE] | 0,000E0 | 0,000E0 | 0,000E0 | 2,757E7 | 3,517E7 | 7,062E7 | Elavl1 |
| F8VQB6 | Unconventional myosin-X OS=Mus musculus GN=Myo10 PE=1 SV=1 - [MYO10_MOUSE] | 0,000E0 | 0,000E0 | 9,589E5 | 7,241E7 | 1,535E8 | 3,570E8 | Myo10 |
| Q91YR1 | Twinfilin-1 OS=Mus musculus GN=Twf1 PE=1 SV=2 - [TWF1_MOUSE] | 0,000E0 | 0,000E0 | 0,000E0 | 2,734E7 | 5,632E7 | 5,271E7 | Twf1 |
| P58774-2 | Isoform 2 of Tropomyosin beta chain OS=Mus musculus GN=Tpm2 - [TPM2_MOUSE] | 0,000E0 | 0,000E0 | 1,208E7 | 1,533E9 | 2,676E9 | 4,035E9 | Tpm2 |
| P17427 | AP-2 complex subunit alpha-2 OS=Mus musculus GN=Ap2a2 PE=1 SV=2 - [AP2A2_MOUSE] | 0,000E0 | 0,000E0 | 0,000E0 | 8,722E7 | 1,468E8 | 1,521E8 | Ap2a2 |
| Q61553 | Fascin OS=Mus musculus GN=Fscn1 PE=1 SV=4 - [FSCN1_MOUSE] | 0,000E0 | 0,000E0 | 0,000E0 | 2,980E7 | 1,009E8 | 1,233E8 | Fscn1 |
| Q8VD75 | Huntingtin-interacting protein 1 OS=Mus musculus GN=Hip1 PE=1 SV=2 - [HIP1_MOUSE] | 0,000E0 | 0,000E0 | 0,000E0 | 9,588E7 | 1,233E8 | 1,739E8 | Hip1 |
| Q9CY58 | Plasminogen activator inhibitor 1 RNA-binding protein OS=Mus musculus GN=Serbp1 PE=1 SV=2 - [PAIRB_MOUSE] | 0,000E0 | 0,000E0 | 1,047E7 | 1,214E8 | 2,632E8 | 5,693E8 | Serbp1 |
| Q62036 | Centrosomal protein of 131 kDa OS=Mus musculus GN=Cep131 PE=1 SV=2 - [CP131_MOUSE] | 0,000E0 | 0,000E0 | 0,000E0 | 2,516E7 | 3,654E7 | 4,793E7 | Cep131 |
| Q8CFI0-3 | Isoform 3 of E3 ubiquitin-protein ligase NEDD4-like OS=Mus musculus GN=Nedd4l - [NED4L_MOUSE] | 0,000E0 | 0,000E0 | 0,000E0 | 3,146E7 | 6,150E7 | 8,335E7 | Nedd4l |
| Q9D0E1-2 | Isoform 2 of Heterogeneous nuclear ribonucleoprotein M OS=Mus musculus GN=Hnrnpm - [HNRPM_MOUSE] | 0,000E0 | 0,000E0 | 2,753E6 | 3,202E7 | 7,252E7 | 8,884E7 | Hnrnpm |
| P84096 | Rho-related GTP-binding protein RhoG OS=Mus musculus GN=Rhog PE=1 SV=1 - [RHOG_MOUSE] | 0,000E0 | 0,000E0 | 0,000E0 | 1,246E7 | 3,793E7 | 2,480E7 | Rhog |
| Q62419 | Endophilin-A2 OS=Mus musculus GN=Sh3gl1 PE=1 SV=1 - [SH3G1_MOUSE] | 0,000E0 | 0,000E0 | 0,000E0 | 1,292E7 | 3,729E7 | 5,374E7 | Sh3gl1 |
| O08583 | THO complex subunit 4 OS=Mus musculus GN=Alyref PE=1 SV=3 - [THOC4_MOUSE] | 0,000E0 | 0,000E0 | 1,665E6 | 2,366E7 | 5,814E7 | 7,797E7 | Alyref |
| Q9DAS9 | Guanine nucleotide-binding protein G(I)/G(S)/G(O) subunit gamma-12 OS=Mus musculus GN=Gng12 PE=1 SV=3 - [GBG12_MOUSE] | 0,000E0 | 0,000E0 | 0,000E0 | 1,868E7 | 2,397E7 | 4,152E7 | Gng12 |
| P47962 | 60S ribosomal protein L5 OS=Mus musculus GN=Rpl5 PE=1 SV=3 - [RL5_MOUSE] | 0,000E0 | 0,000E0 | 7,551E6 | 2,499E8 | 5,103E8 | 1,052E9 | Rpl5 |
| O35887 | Calumenin OS=Mus musculus GN=Calu PE=1 SV=1 - [CALU_MOUSE] | 0,000E0 | 0,000E0 | 2,708E5 | 1,610E7 | 1,348E7 | 3,112E7 | Calu |
| P48962 | ADP/ATP translocase 1 OS=Mus musculus GN=Slc25a4 PE=1 SV=4 - [ADT1_MOUSE] | 3,742E5 | 0,000E0 | 1,161E7 | 4,647E7 | 8,723E7 | 9,796E7 | Slc25a4 |
| P62900 | 60S ribosomal protein L31 OS=Mus musculus GN=Rpl31 PE=1 SV=1 - [RL31_MOUSE] | 0,000E0 | 0,000E0 | 0,000E0 | 2,845E8 | 2,459E8 | 2,062E8 | Rpl31 |
| P47963 | 60S ribosomal protein L13 OS=Mus musculus GN=Rpl13 PE=1 SV=3 - [RL13_MOUSE] | 0,000E0 | 0,000E0 | 3,828E5 | 1,670E8 | 3,233E8 | 9,822E8 | Rpl13 |
| Q9CYL5 | Golgi-associated plant pathogenesis-related protein 1 OS=Mus musculus GN=Glipr2 PE=1 SV=3 - [GAPR1_MOUSE] | 0,000E0 | 0,000E0 | 0,000E0 | 1,357E7 | 7,178E6 | 7,037E7 | Glipr2 |
| Q9D7S7-2 | Isoform 2 of 60S ribosomal protein L22-like 1 OS=Mus musculus GN=Rpl22l1 - [RL22L_MOUSE] | 0,000E0 | 0,000E0 | 3,909E6 | 1,340E8 | 1,938E8 | 5,866E8 | Rpl22l1 |
| P70333 | Heterogeneous nuclear ribonucleoprotein H2 OS=Mus musculus GN=Hnrnph2 PE=1 SV=1 - [HNRH2_MOUSE] | 0,000E0 | 0,000E0 | 0,000E0 | 3,742E7 | 1,034E8 | 1,349E8 | Hnrnph2 |
| Q9QYB5-2 | Isoform 1 of Gamma-adducin OS=Mus musculus GN=Add3 - [ADDG_MOUSE] | 0,000E0 | 0,000E0 | 0,000E0 | 4,894E7 | 8,614E7 | 8,715E7 | Add3 |
| P80315 | T-complex protein 1 subunit delta OS=Mus musculus GN=Cct4 PE=1 SV=3 - [TCPD_MOUSE] | 0,000E0 | 0,000E0 | 0,000E0 | 2,245E7 | 4,479E7 | 3,445E7 | Cct4 |
| P16045 | Galectin-1 OS=Mus musculus GN=Lgals1 PE=1 SV=3 - [LEG1_MOUSE] | 0,000E0 | 0,000E0 | 0,000E0 | 2,196E7 | 1,144E8 | 1,585E8 | Lgals1 |
| Q6ZWX6 | Eukaryotic translation initiation factor 2 subunit 1 OS=Mus musculus GN=Eif2s1 PE=1 SV=3 - [IF2A_MOUSE] | 0,000E0 | 0,000E0 | 0,000E0 | 2,257E7 | 1,515E7 | 5,969E7 | Eif2s1 |
| Q8BTM8 | Filamin-A OS=Mus musculus GN=Flna PE=1 SV=5 - [FLNA_MOUSE] | 0,000E0 | 0,000E0 | 6,534E6 | 4,699E7 | 1,089E8 | 1,788E8 | Flna |
| Q8VHX6-2 | Isoform 2 of Filamin-C OS=Mus musculus GN=Flnc - [FLNC_MOUSE] | 0,000E0 | 0,000E0 | 0,000E0 | 3,769E7 | 8,105E7 | 1,164E8 | Flnc |
| O35658 | Complement component 1 Q subcomponent-binding protein, mitochondrial OS=Mus musculus GN=C1qbp PE=1 SV=1 - [C1QBP_MOUSE] | 0,000E0 | 0,000E0 | 1,830E7 | 1,081E7 | 8,232E7 | 1,538E8 | C1qbp |
| Q91VN6 | Probable ATP-dependent RNA helicase DDX41 OS=Mus musculus GN=Ddx41 PE=1 SV=2 - [DDX41_MOUSE] | 0,000E0 | 0,000E0 | 0,000E0 | 1,932E7 | 3,780E7 | 1,113E8 | Ddx41 |
| Q6ZQ58 | La-related protein 1 OS=Mus musculus GN=Larp1 PE=1 SV=3 - [LARP1_MOUSE] | 0,000E0 | 0,000E0 | 0,000E0 | 1,993E7 | 3,441E7 | 1,043E8 | Larp1 |
| O88342 | WD repeat-containing protein 1 OS=Mus musculus GN=Wdr1 PE=1 SV=3 - [WDR1_MOUSE] | 0,000E0 | 0,000E0 | 0,000E0 | 3,821E7 | 8,776E7 | 1,533E8 | Wdr1 |
| Q9JKK1 | Syntaxin-6 OS=Mus musculus GN=Stx6 PE=1 SV=1 - [STX6_MOUSE] | 0,000E0 | 0,000E0 | 0,000E0 | 5,035E6 | 5,487E6 | 1,000E7 | Stx6 |
| P09405 | Nucleolin OS=Mus musculus GN=Ncl PE=1 SV=2 - [NUCL_MOUSE] | 0,000E0 | 0,000E0 | 1,477E6 | 9,349E7 | 1,382E8 | 4,549E8 | Ncl |
| P32883-2 | Isoform 2B of GTPase KRas OS=Mus musculus GN=Kras - [RASK_MOUSE] | 0,000E0 | 0,000E0 | 0,000E0 | 5,661E6 | 2,507E7 | 3,621E7 | Kras |
| Q9WV32 | Actin-related protein 2/3 complex subunit 1B OS=Mus musculus GN=Arpc1b PE=1 SV=4 - [ARC1B_MOUSE] | 0,000E0 | 0,000E0 | 0,000E0 | 1,711E8 | 3,299E8 | 4,938E8 | Arpc1b |
| Q8BJL1 | F-box only protein 30 OS=Mus musculus GN=Fbxo30 PE=1 SV=2 - [FBX30_MOUSE] | 0,000E0 | 0,000E0 | 0,000E0 | 7,762E7 | 6,969E7 | 7,297E7 | Fbxo30 |

| | | | | | | | | |
|---|---|---|---|---|---|---|---|---|
| P17751 | Triosephosphate isomerase OS=Mus musculus GN=Tpi1 PE=1 SV=4 - [TPIS_MOUSE] | 0,000E0 | 0,000E0 | 0,000E0 | 4,244E7 | 1,087E8 | 9,983E7 | Tpi1 |
| Q61029 | Lamina-associated polypeptide 2, isoforms beta/delta/epsilon/gamma OS=Mus musculus GN=Tmpo PE=1 SV=4 - [LAP2B_MOUSE] | 0,000E0 | 0,000E0 | 9,723E5 | 3,714E7 | 2,430E8 | 4,506E8 | Tmpo |
| P08556 | GTPase NRas OS=Mus musculus GN=Nras PE=1 SV=1 - [RASN_MOUSE] | 0,000E0 | 0,000E0 | 0,000E0 | 5,964E6 | 1,781E7 | 2,239E7 | Nras |
| Q6PCP5-2 | Isoform 2 of Mitochondrial fission factor OS=Mus musculus GN=Mff - [MFF_MOUSE] | 0,000E0 | 0,000E0 | 0,000E0 | 1,152E7 | 3,571E7 | 5,104E7 | Mff |
| Q61029-3 | Isoform Epsilon of Lamina-associated polypeptide 2, isoforms beta/delta/epsilon/gamma OS=Mus musculus GN=Tmpo - [LAP2B_MOUSE] | 0,000E0 | 0,000E0 | 9,723E5 | 3,714E7 | 2,430E8 | 4,506E8 | Tmpo |
| P62071 | Ras-related protein R-Ras2 OS=Mus musculus GN=Rras2 PE=1 SV=1 - [RRAS2_MOUSE] | 0,000E0 | 0,000E0 | 0,000E0 | 7,704E7 | 1,674E8 | 2,174E8 | Rras2 |
| Q6PCP5-3 | Isoform 3 of Mitochondrial fission factor OS=Mus musculus GN=Mff - [MFF_MOUSE] | 0,000E0 | 0,000E0 | 0,000E0 | 1,152E7 | 2,788E7 | 5,104E7 | Mff |
| P50580-2 | Isoform 2 of Proliferation-associated protein 2G4 OS=Mus musculus GN=Pa2g4 - [PA2G4_MOUSE] | 0,000E0 | 0,000E0 | 0,000E0 | 1,663E7 | 9,169E7 | 2,356E8 | Pa2g4 |
| Q9QZD9 | Eukaryotic translation initiation factor 3 subunit I OS=Mus musculus GN=Eif3i PE=1 SV=1 - [EIF3I_MOUSE] | 0,000E0 | 0,000E0 | 0,000E0 | 3,220E7 | 5,741E7 | 6,133E7 | Eif3i |
| P80314 | T-complex protein 1 subunit beta OS=Mus musculus GN=Cct2 PE=1 SV=4 - [TCPB_MOUSE] | 0,000E0 | 0,000E0 | 0,000E0 | 2,422E7 | 4,273E7 | 6,022E7 | Cct2 |
| P48678 | Prelamin-A/C OS=Mus musculus GN=Lmna PE=1 SV=2 - [LMNA_MOUSE] | 0,000E0 | 0,000E0 | 0,000E0 | 6,761E6 | 9,536E6 | 4,185E7 | Lmna |
| Q9WVC3 | Caveolin-2 OS=Mus musculus GN=Cav2 PE=1 SV=1 - [CAV2_MOUSE] | 0,000E0 | 0,000E0 | 0,000E0 | 1,301E7 | 5,749E6 | 2,362E8 | Cav2 |
| P62717 | 60S ribosomal protein L18a OS=Mus musculus GN=Rpl18a PE=1 SV=1 - [RL18A_MOUSE] | 0,000E0 | 0,000E0 | 0,000E0 | 1,130E8 | 2,777E8 | 6,537E8 | Rpl18a |
| Q91V41 | Ras-related protein Rab-14 OS=Mus musculus GN=Rab14 PE=1 SV=3 - [RAB14_MOUSE] | 0,000E0 | 0,000E0 | 3,456E5 | 1,027E8 | 8,942E7 | 1,245E8 | Rab14 |
| Q61656 | Probable ATP-dependent RNA helicase DDX5 OS=Mus musculus GN=Ddx5 PE=1 SV=2 - [DDX5_MOUSE] | 0,000E0 | 0,000E0 | 0,000E0 | 1,638E8 | 3,091E8 | 4,592E8 | Ddx5 |
| Q08509 | Epidermal growth factor receptor kinase substrate 8 OS=Mus musculus GN=Eps8 PE=1 SV=2 - [EPS8_MOUSE] | 0,000E0 | 0,000E0 | 0,000E0 | 5,863E7 | 1,229E8 | 2,265E8 | Eps8 |
| Q9Z1Q5 | Chloride intracellular channel protein 1 OS=Mus musculus GN=Clic1 PE=1 SV=3 - [CLIC1_MOUSE] | 0,000E0 | 0,000E0 | 0,000E0 | 7,145E7 | 1,471E8 | 1,639E8 | Clic1 |
| Q9DBR7 | Protein phosphatase 1 regulatory subunit 12A OS=Mus musculus GN=Ppp1r12a PE=1 SV=2 - [MYPT1_MOUSE] | 0,000E0 | 0,000E0 | 0,000E0 | 7,218E7 | 1,219E8 | 1,762E8 | Ppp1r12a |
| Q9CYR0 | Single-stranded DNA-binding protein, mitochondrial OS=Mus musculus GN=Ssbp1 PE=1 SV=1 - [SSBP_MOUSE] | 0,000E0 | 0,000E0 | 9,309E5 | 6,492E6 | 1,036E7 | 1,504E7 | Ssbp1 |
| Q9WUD1 | STIP1 homology and U box-containing protein 1 OS=Mus musculus GN=Stub1 PE=1 SV=1 - [CHIP_MOUSE] | 0,000E0 | 0,000E0 | 0,000E0 | 1,338E7 | 2,392E7 | 3,369E7 | Stub1 |
| Q61194 | Phosphatidylinositol 4-phosphate 3-kinase C2 domain-containing subunit alpha OS=Mus musculus GN=Pik3c2a PE=1 SV=2 - [P3C2A_MOUSE] | 0,000E0 | 0,000E0 | 0,000E0 | 1,325E8 | 2,152E8 | 2,174E8 | Pik3c2a |
| Q9WUK4 | Replication factor C subunit 2 OS=Mus musculus GN=Rfc2 PE=1 SV=1 - [RFC2_MOUSE] | 0,000E0 | 0,000E0 | 0,000E0 | 1,233E7 | 2,693E7 | 7,828E7 | Rfc2 |
| P70460 | Vasodilator-stimulated phosphoprotein OS=Mus musculus GN=Vasp PE=1 SV=4 - [VASP_MOUSE] | 0,000E0 | 0,000E0 | 0,000E0 | 2,338E7 | 4,614E7 | 5,160E7 | Vasp |
| P35980 | 60S ribosomal protein L18 OS=Mus musculus GN=Rpl18 PE=1 SV=3 - [RL18_MOUSE] | 0,000E0 | 0,000E0 | 9,509E5 | 1,395E8 | 2,471E8 | 4,872E8 | Rpl18 |
| Q6R891 | Neurabin-2 OS=Mus musculus GN=Ppp1r9b PE=1 SV=1 - [NEB2_MOUSE] | 0,000E0 | 0,000E0 | 0,000E0 | 2,384E7 | 5,639E7 | 4,791E7 | Ppp1r9b |
| Q8BGS2 | BolA-like protein 2 OS=Mus musculus GN=Bola2 PE=1 SV=1 - [BOLA2_MOUSE] | 0,000E0 | 0,000E0 | 0,000E0 | 8,013E6 | 1,473E7 | 1,713E7 | Bola2 |
| Q61699-2 | Isoform HSP105-beta of Heat shock protein 105 kDa OS=Mus musculus GN=Hsph1 - [HS105_MOUSE] | 0,000E0 | 0,000E0 | 0,000E0 | 2,281E7 | 4,287E7 | 6,089E7 | Hsph1 |
| P51150 | Ras-related protein Rab-7a OS=Mus musculus GN=Rab7a PE=1 SV=2 - [RAB7A_MOUSE] | 0,000E0 | 0,000E0 | 0,000E0 | 1,686E7 | 2,573E7 | 4,079E7 | Rab7a |
| Q9D1C8 | Vacuolar protein sorting-associated protein 28 homolog OS=Mus musculus GN=Vps28 PE=1 SV=1 - [VPS28_MOUSE] | 0,000E0 | 0,000E0 | 0,000E0 | 9,550E6 | 1,693E7 | 2,457E7 | Vps28 |
| Q6AW69-5 | Isoform 5 of Cingulin-like protein 1 OS=Mus musculus GN=Cgnl1 - [CGNL1_MOUSE] | 0,000E0 | 0,000E0 | 0,000E0 | 1,233E7 | 2,225E7 | 4,436E7 | Cgnl1 |
| P0C0S6 | Histone H2A.Z OS=Mus musculus GN=H2afz PE=1 SV=2 - [H2AZ_MOUSE] | 0,000E0 | 0,000E0 | 1,349E7 | 3,580E7 | 1,246E8 | 7,278E8 | H2afz |
| Q61598 | Rab GDP dissociation inhibitor beta OS=Mus musculus GN=Gdi2 PE=1 SV=1 - [GDIB_MOUSE] | 0,000E0 | 0,000E0 | 0,000E0 | 1,071E7 | 1,674E7 | 2,722E7 | Gdi2 |
| O08709 | Peroxiredoxin-6 OS=Mus musculus GN=Prdx6 PE=1 SV=3 - [PRDX6_MOUSE] | 0,000E0 | 0,000E0 | 0,000E0 | 5,746E6 | 8,958E6 | 8,522E6 | Prdx6 |
| Q921U8-2 | Isoform L2 of Smoothelin OS=Mus musculus GN=Smtn - [SMTN_MOUSE] | 0,000E0 | 0,000E0 | 0,000E0 | 1,817E7 | 1,588E7 | 2,681E7 | Smtn |
| P11440 | Cyclin-dependent kinase 1 OS=Mus musculus GN=Cdk1 PE=1 SV=3 - [CDK1_MOUSE] | 0,000E0 | 0,000E0 | 0,000E0 | 1,447E7 | 3,150E7 | 4,251E7 | Cdk1 |
| P97461 | 40S ribosomal protein S5 OS=Mus musculus GN=Rps5 PE=1 SV=3 - [RS5_MOUSE] | 0,000E0 | 0,000E0 | 0,000E0 | 5,890E7 | 1,996E8 | 2,289E8 | Rps5 |
| Q91WK2 | Eukaryotic translation initiation factor 3 subunit H OS=Mus musculus GN=Eif3h PE=1 SV=1 - [EIF3H_MOUSE] | 0,000E0 | 0,000E0 | 0,000E0 | 1,755E7 | 1,304E7 | 4,705E7 | Eif3h |
| P11031 | Activated RNA polymerase II transcriptional coactivator p15 OS=Mus musculus GN=Sub1 PE=1 SV=3 - [TCP4_MOUSE] | 0,000E0 | 0,000E0 | 7,455E5 | 8,276E6 | 3,919E7 | 1,363E8 | Sub1 |
| O88746 | Target of Myb protein 1 OS=Mus musculus GN=Tom1 PE=1 SV=1 - [TOM1_MOUSE] | 0,000E0 | 0,000E0 | 0,000E0 | 6,818E6 | 9,103E6 | 2,581E7 | Tom1 |
| Q99JP6 | Homer protein homolog 3 OS=Mus musculus GN=Homer3 PE=1 SV=2 - [HOME3_MOUSE] | 0,000E0 | 0,000E0 | 0,000E0 | 8,433E6 | 2,793E7 | 3,148E7 | Homer3 |
| P61979-3 | Isoform 3 of Heterogeneous nuclear ribonucleoprotein K OS=Mus musculus GN=Hnrnpk - [HNRPK_MOUSE] | 0,000E0 | 0,000E0 | 5,493E6 | 7,494E7 | 1,383E8 | 1,478E8 | Hnrnpk |
| O88952 | Protein lin-7 homolog C OS=Mus musculus GN=Lin7c PE=1 SV=2 - [LIN7C_MOUSE] | 0,000E0 | 0,000E0 | 0,000E0 | 1,219E7 | 1,148E7 | 2,671E7 | Lin7c |
| P63094 | Guanine nucleotide-binding protein G(s) subunit alpha isoforms short OS=Mus musculus GN=Gnas PE=1 SV=1 - [GNAS2_MOUSE] | 0,000E0 | 0,000E0 | 9,807E4 | 4,359E7 | 1,460E8 | 1,327E8 | Gnas |
| O35737 | Heterogeneous nuclear ribonucleoprotein H OS=Mus musculus GN=Hnrnph1 PE=1 SV=3 - [HNRH1_MOUSE] | 0,000E0 | 0,000E0 | 0,000E0 | 6,189E7 | 1,034E8 | 1,349E8 | Hnrnph1 |
| P46062 | Signal-induced proliferation-associated protein 1 OS=Mus musculus GN=Sipa1 PE=1 SV=2 - [SIPA1_MOUSE] | 0,000E0 | 0,000E0 | 0,000E0 | 9,330E7 | 2,101E8 | 2,001E8 | Sipa1 |
| Q03141-2 | Isoform 2 of MAP/microtubule affinity-regulating kinase 3 OS=Mus musculus GN=Mark3 - [MARK3_MOUSE] | 0,000E0 | 0,000E0 | 0,000E0 | 7,830E6 | 1,999E7 | 3,415E7 | Mark3 |
| Q8BFR5 | Elongation factor Tu, mitochondrial OS=Mus musculus GN=Tufm PE=1 SV=1 - [EFTU_MOUSE] | 0,000E0 | 0,000E0 | 0,000E0 | 1,445E7 | 4,087E7 | 3,940E7 | Tufm |
| Q80WJ7 | Protein LYRIC OS=Mus musculus GN=Mtdh PE=1 SV=1 - [LYRIC_MOUSE] | 0,000E0 | 0,000E0 | 0,000E0 | 1,880E7 | 2,869E7 | 5,753E7 | Mtdh |
| Q9CQH7 | Transcription factor BTF3 homolog 4 OS=Mus musculus GN=Btf3l4 PE=1 SV=1 - [BT3L4_MOUSE] | 0,000E0 | 0,000E0 | 0,000E0 | 8,981E6 | 5,634E6 | 2,250E7 | Btf3l4 |
| Q9R0L6 | Pericentriolar material 1 protein OS=Mus musculus GN=Pcm1 PE=1 SV=2 - [PCM1_MOUSE] | 0,000E0 | 0,000E0 | 0,000E0 | 3,804E7 | 5,684E7 | 7,729E7 | Pcm1 |
| Q9WTM5 | RuvB-like 2 OS=Mus musculus GN=Ruvbl2 PE=1 SV=3 - [RUVB2_MOUSE] | 0,000E0 | 0,000E0 | 7,090E6 | 5,668E6 | 1,560E7 | 1,195E7 | Ruvbl2 |
| Q9ESZ8-5 | Isoform 5 of General transcription factor II-I OS=Mus musculus GN=Gtf2i - [GTF2I_MOUSE] | 0,000E0 | 0,000E0 | 0,000E0 | 2,469E7 | 4,671E7 | 5,957E7 | Gtf2i |
| Q8QZY1 | Eukaryotic translation initiation factor 3 subunit L OS=Mus musculus GN=Eif3l PE=1 SV=1 - [EIF3L_MOUSE] | 0,000E0 | 0,000E0 | 0,000E0 | 1,678E7 | 4,144E7 | 7,162E7 | Eif3l |
| Q6ZWV7 | 60S ribosomal protein L35 OS=Mus musculus GN=Rpl35 PE=1 SV=1 - [RL35_MOUSE] | 0,000E0 | 0,000E0 | 0,000E0 | 1,505E8 | 3,069E8 | 3,144E8 | Rpl35 |
| P23116 | Eukaryotic translation initiation factor 3 subunit A OS=Mus musculus GN=Eif3a PE=1 SV=5 - [EIF3A_MOUSE] | 0,000E0 | 0,000E0 | 0,000E0 | 3,221E7 | 9,439E7 | 1,025E8 | Eif3a |
| Q9JLQ0 | CD2-associated protein OS=Mus musculus GN=Cd2ap PE=1 SV=3 - [CD2AP_MOUSE] | 0,000E0 | 0,000E0 | 0,000E0 | 9,052E6 | 2,508E7 | 3,529E7 | Cd2ap |
| Q922Y1 | UBX domain-containing protein 1 OS=Mus musculus GN=Ubxn1 PE=1 SV=1 - [UBXN1_MOUSE] | 0,000E0 | 0,000E0 | 0,000E0 | 6,038E6 | 5,523E6 | 1,906E7 | Ubxn1 |
| Q8BL97-3 | Isoform 3 of Serine/arginine-rich splicing factor 7 OS=Mus musculus GN=Srsf7 - [SRSF7_MOUSE] | 0,000E0 | 0,000E0 | 0,000E0 | 3,855E7 | 8,333E7 | 7,202E7 | Srsf7 |
| P84099 | 60S ribosomal protein L19 OS=Mus musculus GN=Rpl19 PE=1 SV=1 - [RL19_MOUSE] | 0,000E0 | 0,000E0 | 0,000E0 | 1,113E8 | 1,560E8 | 4,269E8 | Rpl19 |
| P60335 | Poly(rC)-binding protein 1 OS=Mus musculus GN=Pcbp1 PE=1 SV=1 - [PCBP1_MOUSE] | 0,000E0 | 0,000E0 | 6,208E5 | 1,724E7 | 4,403E7 | 8,903E7 | Pcbp1 |
| P46638 | Ras-related protein Rab-11B OS=Mus musculus GN=Rab11b PE=1 SV=3 - [RB11B_MOUSE] | 0,000E0 | 0,000E0 | 0,000E0 | 1,056E7 | 1,009E7 | 3,102E7 | Rab11b |
| P58252 | Elongation factor 2 OS=Mus musculus GN=Eef2 PE=1 SV=2 - [EF2_MOUSE] | 0,000E0 | 0,000E0 | 0,000E0 | 4,746E7 | 1,351E8 | 1,998E8 | Eef2 |
| Q6P1F6 | Serine/threonine-protein phosphatase 2A 55 kDa regulatory subunit B alpha isoform OS=Mus musculus GN=Ppp2r2a PE=1 SV=1 - [2ABA_MOUSE] | 0,000E0 | 0,000E0 | 0,000E0 | 1,087E7 | 2,245E7 | 3,175E7 | Ppp2r2a |
| Q9QYC0 | Alpha-adducin OS=Mus musculus GN=Add1 PE=1 SV=2 - [ADDA_MOUSE] | 0,000E0 | 0,000E0 | 0,000E0 | 1,140E8 | 1,436E8 | 1,348E8 | Add1 |
| P62274 | 40S ribosomal protein S29 OS=Mus musculus GN=Rps29 PE=3 SV=2 - [RS29_MOUSE] | 0,000E0 | 0,000E0 | 0,000E0 | 1,264E7 | 1,214E8 | 3,232E8 | Rps29 |
| Q8BU31 | Ras-related protein Rap-2c OS=Mus musculus GN=Rap2c PE=1 SV=1 - [RAP2C_MOUSE] | 0,000E0 | 0,000E0 | 0,000E0 | 5,651E6 | 1,716E7 | 1,418E7 | Rap2c |
| P63087-2 | Isoform Gamma-2 of Serine/threonine-protein phosphatase PP1-gamma catalytic subunit OS=Mus musculus GN=Ppp1cc - [PP1G_MOUSE] | 0,000E0 | 0,000E0 | 2,353E6 | 5,878E7 | 1,101E8 | 2,064E8 | Ppp1cc |
| P84091 | AP-2 complex subunit mu OS=Mus musculus GN=Ap2m1 PE=1 SV=1 - [AP2M1_MOUSE] | 0,000E0 | 0,000E0 | 0,000E0 | 2,087E8 | 2,792E8 | 3,114E8 | Ap2m1 |
| Q61187 | Tumor susceptibility gene 101 protein OS=Mus musculus GN=Tsg101 PE=1 SV=2 - [TS101_MOUSE] | 0,000E0 | 0,000E0 | 0,000E0 | 1,464E7 | 2,686E7 | 6,469E7 | Tsg101 |
| P42932 | T-complex protein 1 subunit theta OS=Mus musculus GN=Cct8 PE=1 SV=3 - [TCPQ_MOUSE] | 0,000E0 | 0,000E0 | 6,390E5 | 2,318E7 | 4,541E7 | 5,123E7 | Cct8 |
| P62849-2 | Isoform 2 of 40S ribosomal protein S24 OS=Mus musculus GN=Rps24 - [RS24_MOUSE] | 0,000E0 | 0,000E0 | 0,000E0 | 1,181E8 | 1,024E8 | 1,444E8 | Rps24 |
| P97379-2 | Isoform B of Ras GTPase-activating protein-binding protein 2 OS=Mus musculus GN=G3bp2 - [G3BP2_MOUSE] | 0,000E0 | 0,000E0 | 5,026E6 | 3,650E7 | 7,730E7 | 1,353E8 | G3bp2 |
| Q3TN34 | MICAL-like protein 2 OS=Mus musculus GN=Micall2 PE=1 SV=1 - [MILK2_MOUSE] | 0,000E0 | 0,000E0 | 0,000E0 | 1,204E7 | 2,860E7 | 4,201E7 | Micall2 |
| Q9DBC7 | cAMP-dependent protein kinase type I-alpha regulatory subunit OS=Mus musculus GN=Prkar1a PE=1 SV=3 - [KAP0_MOUSE] | 0,000E0 | 0,000E0 | 0,000E0 | 2,393E7 | 5,342E7 | 6,425E7 | Prkar1a |
| P67984 | 60S ribosomal protein L22 OS=Mus musculus GN=Rpl22 PE=1 SV=2 - [RL22_MOUSE] | 0,000E0 | 0,000E0 | 2,391E6 | 1,809E8 | 3,362E8 | 7,719E8 | Rpl22 |
| Q8BG05 | Heterogeneous nuclear ribonucleoprotein A3 OS=Mus musculus GN=Hnrnpa3 PE=1 SV=1 - [ROA3_MOUSE] | 0,000E0 | 0,000E0 | 6,529E5 | 9,406E7 | 1,311E8 | 2,374E8 | Hnrnpa3 |
| Q6ZWY3 | 40S ribosomal protein S27-like OS=Mus musculus GN=Rps27l PE=1 SV=3 - [RS27L_MOUSE] | 0,000E0 | 0,000E0 | 1,094E7 | 2,132E8 | 2,594E8 | 8,226E8 | Rps27l |
| Q6ZWU9 | 40S ribosomal protein S27 OS=Mus musculus GN=Rps27 PE=1 SV=3 - [RS27_MOUSE] | 0,000E0 | 0,000E0 | 1,094E7 | 2,519E8 | 4,230E8 | 8,883E8 | Rps27 |
| P05480 | Neuronal proto-oncogene tyrosine-protein kinase Src OS=Mus musculus GN=Src PE=1 SV=4 - [SRC_MOUSE] | 0,000E0 | 0,000E0 | 2,114E5 | 1,340E7 | 5,183E7 | 6,219E7 | Src |
| Q8BG81 | Polymerase delta-interacting protein 3 OS=Mus musculus GN=Poldip3 PE=1 SV=1 - [PDIP3_MOUSE] | 0,000E0 | 0,000E0 | 0,000E0 | 1,115E7 | 2,284E7 | 3,877E7 | Poldip3 |
| Q8BSD5 | SH2 domain-containing adapter protein E OS=Mus musculus GN=She PE=1 SV=1 - [SHE_MOUSE] | 0,000E0 | 0,000E0 | 0,000E0 | 1,152E7 | 3,268E7 | 1,414E7 | She |
| Q01320 | DNA topoisomerase 2-alpha OS=Mus musculus GN=Top2a PE=1 SV=2 - [TOP2A_MOUSE] | 0,000E0 | 0,000E0 | 0,000E0 | 2,889E7 | 6,027E7 | 1,207E8 | Top2a |
| Q8CC35 | Synaptopodin OS=Mus musculus GN=Synpo PE=1 SV=2 - [SYNPO_MOUSE] | 0,000E0 | 0,000E0 | 0,000E0 | 8,920E6 | 3,404E7 | 5,640E7 | Synpo |
| Q9CQE8 | UPF0568 protein C14orf166 homolog OS=Mus musculus PE=1 SV=1 - [CN166_MOUSE] | 0,000E0 | 0,000E0 | 0,000E0 | 1,244E7 | 2,785E7 | 4,335E7 | CN166 |
| **Q8VHG2** | **Angiomotin OS=Mus musculus GN=Amot PE=1 SV=3 - [AMOT_MOUSE]** | **0,000E0** | **0,000E0** | **0,000E0** | **1,728E7** | **9,856E6** | **1,533E7** | Amot |
| Q78PY7 | Staphylococcal nuclease domain-containing protein 1 OS=Mus musculus GN=Snd1 PE=1 SV=1 - [SND1_MOUSE] | 0,000E0 | 0,000E0 | 0,000E0 | 2,632E7 | 4,073E7 | 8,755E7 | Snd1 |
| P62911 | 60S ribosomal protein L32 OS=Mus musculus GN=Rpl32 PE=1 SV=2 - [RL32_MOUSE] | 0,000E0 | 0,000E0 | 0,000E0 | 2,833E7 | 7,406E7 | 1,433E8 | Rpl32 |
| P62855 | 40S ribosomal protein S26 OS=Mus musculus GN=Rps26 PE=1 SV=3 - [RS26_MOUSE] | 0,000E0 | 0,000E0 | 0,000E0 | 1,540E8 | 4,089E8 | 4,789E8 | Rps26 |
| Q99K48 | Non-POU domain-containing octamer-binding protein OS=Mus musculus GN=Nono PE=1 SV=3 - [NONO_MOUSE] | 0,000E0 | 0,000E0 | 0,000E0 | 3,121E7 | 4,880E7 | 1,454E8 | Nono |
| Q9Z2U0 | Proteasome subunit alpha type-7 OS=Mus musculus GN=Psma7 PE=1 SV=1 - [PSA7_MOUSE] | 0,000E0 | 0,000E0 | 0,000E0 | 5,100E6 | 5,638E6 | 9,853E6 | Psma7 |
| P27661 | Histone H2AX OS=Mus musculus GN=H2afx PE=1 SV=2 - [H2AX_MOUSE] | 1,458E7 | 9,648E6 | 2,742E7 | 5,055E7 | 9,659E7 | 7,366E8 | H2afx |
| P48036 | Annexin A5 OS=Mus musculus GN=Anxa5 PE=1 SV=1 - [ANXA5_MOUSE] | 0,000E0 | 0,000E0 | 3,591E6 | 8,024E6 | 6,314E6 | 9,419E6 | Anxa5 |
| Q3V038 | Tetratricopeptide repeat protein 9A OS=Mus musculus GN=Ttc9 PE=1 SV=1 - [TTC9A_MOUSE] | 0,000E0 | 0,000E0 | 0,000E0 | 8,639E6 | 2,000E7 | 3,373E7 | Ttc9 |

| | | | | | | | | |
|---|---|---|---|---|---|---|---|---|
| Q9QZQ1-2 | Isoform 1 of Afadin OS=Mus musculus GN=Afdn - [AFAD_MOUSE] | 0,000E0 | 0,000E0 | 0,000E0 | 3,317E7 | 5,627E7 | 7,192E7 | Afdn |
| Q3UKC1 | Tax1-binding protein 1 homolog OS=Mus musculus GN=Tax1bp1 PE=1 SV=2 - [TAXB1_MOUSE] | 0,000E0 | 0,000E0 | 0,000E0 | 2,523E7 | 4,905E7 | 7,367E7 | Tax1bp1 |
| Q8BHE0 | Proline-rich protein 11 OS=Mus musculus GN=Prr11 PE=2 SV=1 - [PRR11_MOUSE] | 0,000E0 | 0,000E0 | 0,000E0 | 1,001E7 | 2,125E7 | 5,087E7 | Prr11 |
| P63085 | Mitogen-activated protein kinase 1 OS=Mus musculus GN=Mapk1 PE=1 SV=3 - [MK01_MOUSE] | 0,000E0 | 0,000E0 | 0,000E0 | 1,495E7 | 2,924E7 | 2,626E7 | Mapk1 |
| Q9EPU0-2 | Isoform 2 of Regulator of nonsense transcripts 1 OS=Mus musculus GN=Upf1 - [RENT1_MOUSE] | 0,000E0 | 0,000E0 | 0,000E0 | 1,840E7 | 3,985E7 | 6,251E7 | Upf1 |
| P26043 | Radixin OS=Mus musculus GN=Rdx PE=1 SV=3 - [RADI_MOUSE] | 0,000E0 | 0,000E0 | 2,157E5 | 1,246E8 | 4,352E8 | 7,129E8 | Rdx |
| Q99LE6 | ATP-binding cassette sub-family F member 2 OS=Mus musculus GN=Abcf2 PE=1 SV=1 - [ABCF2_MOUSE] | 0,000E0 | 0,000E0 | 0,000E0 | 4,174E7 | 7,807E7 | 1,309E8 | Abcf2 |
| O70194 | Eukaryotic translation initiation factor 3 subunit D OS=Mus musculus GN=Eif3d PE=1 SV=2 - [EIF3D_MOUSE] | 0,000E0 | 0,000E0 | 0,000E0 | 1,504E7 | 4,491E7 | 7,832E7 | Eif3d |
| P35278 | Ras-related protein Rab-5C OS=Mus musculus GN=Rab5c PE=1 SV=2 - [RAB5C_MOUSE] | 0,000E0 | 0,000E0 | 0,000E0 | 4,074E6 | 2,691E7 | 1,939E7 | Rab5c |
| Q8R323 | Replication factor C subunit 3 OS=Mus musculus GN=Rfc3 PE=1 SV=1 - [RFC3_MOUSE] | 0,000E0 | 0,000E0 | 2,006E6 | 1,036E7 | 8,776E6 | 3,592E7 | Rfc3 |
| Q8BFW7 | Lipoma-preferred partner homolog OS=Mus musculus GN=Lpp PE=1 SV=1 - [LPP_MOUSE] | 0,000E0 | 0,000E0 | 0,000E0 | 1,323E7 | 2,344E7 | 6,516E7 | Lpp |
| P07901 | Heat shock protein HSP 90-alpha OS=Mus musculus GN=Hsp90aa1 PE=1 SV=4 - [HS90A_MOUSE] | 1,255E7 | 0,000E0 | 5,594E6 | 1,376E8 | 3,170E8 | 4,339E8 | Hsp90aa1 |
| Q69ZN7 | Myoferlin OS=Mus musculus GN=Myof PE=1 SV=2 - [MYOF_MOUSE] | 0,000E0 | 0,000E0 | 0,000E0 | 2,787E7 | 5,833E7 | 9,810E7 | Myof |
| P62317 | Small nuclear ribonucleoprotein Sm D2 OS=Mus musculus GN=Snrpd2 PE=1 SV=1 - [SMD2_MOUSE] | 0,000E0 | 0,000E0 | 0,000E0 | 2,805E6 | 4,126E6 | 8,124E6 | Snrpd2 |
| P32233 | Developmentally-regulated GTP-binding protein 1 OS=Mus musculus GN=Drg1 PE=1 SV=1 - [DRG1_MOUSE] | 0,000E0 | 0,000E0 | 0,000E0 | 6,982E6 | 7,431E6 | 1,208E7 | Drg1 |
| Q8R105 | Vacuolar protein sorting-associated protein 37C OS=Mus musculus GN=Vps37c PE=1 SV=1 - [VP37C_MOUSE] | 0,000E0 | 0,000E0 | 0,000E0 | 1,138E7 | 1,580E7 | 6,983E6 | Vps37c |
| P62331 | ADP-ribosylation factor 6 OS=Mus musculus GN=Arf6 PE=1 SV=2 - [ARF6_MOUSE] | 0,000E0 | 0,000E0 | 0,000E0 | 1,985E7 | 1,674E7 | 5,163E7 | Arf6 |
| Q61235 | Beta-2-syntrophin OS=Mus musculus GN=Sntb2 PE=1 SV=2 - [SNTB2_MOUSE] | 0,000E0 | 0,000E0 | 0,000E0 | 1,425E7 | 3,096E7 | 6,413E7 | Sntb2 |
| Q99JY3 | GTPase IMAP family member 4 OS=Mus musculus GN=Gimap4 PE=1 SV=1 - [GIMA4_MOUSE] | 0,000E0 | 0,000E0 | 0,000E0 | 1,139E7 | 2,714E7 | 1,689E7 | Gimap4 |
| Q3UEB3-2 | Isoform 2 of Poly(U)-binding-splicing factor PUF60 OS=Mus musculus GN=Puf60 - [PUF60_MOUSE] | 0,000E0 | 0,000E0 | 0,000E0 | 1,578E7 | 3,409E7 | 4,270E7 | Puf60 |
| Q61316 | Heat shock 70 kDa protein 4 OS=Mus musculus GN=Hspa4 PE=1 SV=1 - [HSP74_MOUSE] | 0,000E0 | 0,000E0 | 0,000E0 | 1,679E7 | 3,326E7 | 5,921E7 | Hspa4 |
| Q922Q4 | Pyrroline-5-carboxylate reductase 2 OS=Mus musculus GN=Pycr2 PE=1 SV=1 - [P5CR2_MOUSE] | 0,000E0 | 0,000E0 | 0,000E0 | 4,318E7 | 5,612E7 | 1,792E8 | Pycr2 |
| O70551 | SRSF protein kinase 1 OS=Mus musculus GN=Srpk1 PE=1 SV=2 - [SRPK1_MOUSE] | 0,000E0 | 0,000E0 | 0,000E0 | 8,180E6 | 1,890E7 | 4,935E7 | Srpk1 |
| P35922-9 | Isoform 9 of Synaptic functional regulator FMR1 OS=Mus musculus GN=Fmr1 - [FMR1_MOUSE] | 0,000E0 | 0,000E0 | 1,048E6 | 6,566E7 | 1,602E8 | 3,140E8 | Fmr1 |
| P63001 | Ras-related C3 botulinum toxin substrate 1 OS=Mus musculus GN=Rac1 PE=1 SV=1 - [RAC1_MOUSE] | 0,000E0 | 0,000E0 | 0,000E0 | 3,428E7 | 8,787E7 | 1,288E8 | Rac1 |
| P80316 | T-complex protein 1 subunit epsilon OS=Mus musculus GN=Cct5 PE=1 SV=1 - [TCPE_MOUSE] | 0,000E0 | 0,000E0 | 2,663E5 | 6,466E7 | 3,368E7 | 2,674E7 | Cct5 |
| P26040 | Ezrin OS=Mus musculus GN=Ezr PE=1 SV=3 - [EZRI_MOUSE] | 0,000E0 | 0,000E0 | 6,577E5 | 1,246E8 | 4,352E8 | 7,129E8 | Ezr |
| Q91VR2 | ATP synthase subunit gamma, mitochondrial OS=Mus musculus GN=Atp5c1 PE=1 SV=1 - [ATPG_MOUSE] | 0,000E0 | 0,000E0 | 0,000E0 | 1,589E7 | 2,744E7 | 3,222E7 | Atp5c1 |
| Q60902 | Epidermal growth factor receptor substrate 15-like 1 OS=Mus musculus GN=Eps15l1 PE=1 SV=3 - [EP15R_MOUSE] | 0,000E0 | 0,000E0 | 0,000E0 | 1,358E7 | 2,052E7 | 2,776E7 | Eps15l1 |
| Q05512-3 | Isoform 3 of Serine/threonine-protein kinase MARK2 OS=Mus musculus GN=Mark2 - [MARK2_MOUSE] | 0,000E0 | 0,000E0 | 0,000E0 | 1,491E7 | 3,665E7 | 5,667E7 | Mark2 |
| P60229 | Eukaryotic translation initiation factor 3 subunit E OS=Mus musculus GN=Eif3e PE=1 SV=1 - [EIF3E_MOUSE] | 0,000E0 | 0,000E0 | 0,000E0 | 1,019E7 | 4,321E7 | 5,216E7 | Eif3e |
| P14069 | Protein S100-A6 OS=Mus musculus GN=S100a6 PE=1 SV=3 - [S10A6_MOUSE] | 0,000E0 | 0,000E0 | 0,000E0 | 1,257E6 | 2,975E7 | 4,743E7 | S100a6 |
| P49817 | Caveolin-1 OS=Mus musculus GN=Cav1 PE=1 SV=1 - [CAV1_MOUSE] | 0,000E0 | 0,000E0 | 0,000E0 | 1,334E8 | 2,111E8 | 3,995E8 | Cav1 |
| Q5SF07 | Insulin-like growth factor 2 mRNA-binding protein 2 OS=Mus musculus GN=Igf2bp2 PE=1 SV=1 - [IF2B2_MOUSE] | 0,000E0 | 0,000E0 | 0,000E0 | 2,192E7 | 4,628E7 | 7,923E7 | Igf2bp2 |
| P17225 | Polypyrimidine tract-binding protein 1 OS=Mus musculus GN=Ptbp1 PE=1 SV=2 - [PTBP1_MOUSE] | 0,000E0 | 0,000E0 | 5,409E4 | 7,832E7 | 1,331E8 | 2,944E8 | Ptbp1 |
| Q91VR5 | ATP-dependent RNA helicase DDX1 OS=Mus musculus GN=Ddx1 PE=1 SV=1 - [DDX1_MOUSE] | 0,000E0 | 0,000E0 | 0,000E0 | 2,905E7 | 4,032E7 | 9,545E7 | Ddx1 |
| Q60854 | Serpin B6 OS=Mus musculus GN=Serpinb6 PE=1 SV=1 - [SPB6_MOUSE] | 0,000E0 | 0,000E0 | 0,000E0 | 3,046E7 | 3,197E7 | 6,054E7 | Serpinb6 |
| P57776 | Elongation factor 1-delta OS=Mus musculus GN=Eef1d PE=1 SV=3 - [EF1D_MOUSE] | 0,000E0 | 0,000E0 | 0,000E0 | 7,344E7 | 4,998E7 | 1,104E8 | Eef1d |
| Q99JI6 | Ras-related protein Rap-1b OS=Mus musculus GN=Rap1b PE=1 SV=2 - [RAP1B_MOUSE] | 0,000E0 | 0,000E0 | 0,000E0 | 1,082E7 | 4,667E7 | 7,742E7 | Rap1b |
| P35279 | Ras-related protein Rab-6A OS=Mus musculus GN=Rab6a PE=1 SV=4 - [RAB6A_MOUSE] | 0,000E0 | 0,000E0 | 3,456E5 | 3,992E7 | 1,305E8 | 1,415E8 | Rab6a |
| **Q9D4H4-2** | **Isoform 2 of Angiomotin-like protein 1 OS=Mus musculus GN=Amotl1 - [AMOL1_MOUSE]** | **0,000E0** | **0,000E0** | **0,000E0** | **3,518E7** | **3,733E7** | **4,278E7** | Amotl1 |
| Q9CWK8 | Sorting nexin-2 OS=Mus musculus GN=Snx2 PE=1 SV=2 - [SNX2_MOUSE] | 0,000E0 | 0,000E0 | 2,776E6 | 1,623E6 | 5,251E6 | 1,081E7 | Snx2 |
| P23242 | Gap junction alpha-1 protein OS=Mus musculus GN=Gja1 PE=1 SV=2 - [CXA1_MOUSE] | 0,000E0 | 0,000E0 | 0,000E0 | 7,979E7 | 1,158E8 | 1,664E8 | Gja1 |
| Q9D898 | Actin-related protein 2/3 complex subunit 5-like protein OS=Mus musculus GN=Arpc5l PE=1 SV=1 - [ARP5L_MOUSE] | 0,000E0 | 0,000E0 | 0,000E0 | 2,921E7 | 8,950E7 | 1,197E8 | Arpc5l |
| P63321 | Ras-related protein Ral-A OS=Mus musculus GN=Rala PE=1 SV=1 - [RALA_MOUSE] | 0,000E0 | 0,000E0 | 0,000E0 | 9,094E6 | 9,536E6 | 2,875E7 | Rala |
| P11983 | T-complex protein 1 subunit alpha OS=Mus musculus GN=Tcp1 PE=1 SV=3 - [TCPA_MOUSE] | 0,000E0 | 0,000E0 | 2,616E6 | 3,239E7 | 5,712E7 | 6,535E7 | Tcp1 |
| Q60875-5 | Isoform 5 of Rho guanine nucleotide exchange factor 2 OS=Mus musculus GN=Arhgef2 - [ARHG2_MOUSE] | 0,000E0 | 0,000E0 | 0,000E0 | 1,999E7 | 3,828E7 | 3,560E7 | Arhgef2 |
| Q99PT1 | Rho GDP-dissociation inhibitor 1 OS=Mus musculus GN=Arhgdia PE=1 SV=3 - [GDIR1_MOUSE] | 0,000E0 | 0,000E0 | 0,000E0 | 2,653E7 | 5,288E7 | 1,434E8 | Arhgdia |
| P61027 | Ras-related protein Rab-10 OS=Mus musculus GN=Rab10 PE=1 SV=1 - [RAB10_MOUSE] | 0,000E0 | 0,000E0 | 3,456E5 | 5,919E7 | 1,322E8 | 1,924E8 | Rab10 |
| Q9DCT8 | Cysteine-rich protein 2 OS=Mus musculus GN=Crip2 PE=1 SV=1 - [CRIP2_MOUSE] | 0,000E0 | 0,000E0 | 0,000E0 | 6,090E7 | 5,476E7 | 1,094E6 | Crip2 |
| Q5SRX1 | TOM1-like protein 2 OS=Mus musculus GN=Tom1l2 PE=1 SV=1 - [TM1L2_MOUSE] | 0,000E0 | 0,000E0 | 0,000E0 | 6,885E6 | 2,288E7 | 3,616E7 | Tom1l2 |
| Q9Z2X1-2 | Isoform 2 of Heterogeneous nuclear ribonucleoprotein F OS=Mus musculus GN=Hnrnpf - [HNRPF_MOUSE] | 0,000E0 | 0,000E0 | 1,777E6 | 5,494E7 | 8,067E7 | 4,935E7 | Hnrnpf |
| E9Q5G3 | Kinesin-like protein KIF23 OS=Mus musculus GN=Kif23 PE=1 SV=1 - [KIF23_MOUSE] | 0,000E0 | 0,000E0 | 0,000E0 | 1,194E7 | 3,136E7 | 9,652E7 | Kif23 |
| O08529 | Calpain-2 catalytic subunit OS=Mus musculus GN=Capn2 PE=1 SV=4 - [CAN2_MOUSE] | 0,000E0 | 0,000E0 | 0,000E0 | 1,254E7 | 2,816E7 | 4,059E7 | Capn2 |
| Q8C1B7-3 | Isoform 3 of Septin-11 OS=Mus musculus GN=Sept11 - [SEP11_MOUSE] | 0,000E0 | 0,000E0 | 4,406E6 | 1,720E7 | 3,815E7 | 1,994E7 | SEP11 |
| P42669 | Transcriptional activator protein Pur-alpha OS=Mus musculus GN=Pura PE=1 SV=1 - [PURA_MOUSE] | 0,000E0 | 0,000E0 | 0,000E0 | 8,450E6 | 2,327E7 | 4,283E7 | Pura |
| P80313 | T-complex protein 1 subunit eta OS=Mus musculus GN=Cct7 PE=1 SV=1 - [TCPH_MOUSE] | 0,000E0 | 0,000E0 | 8,568E6 | 2,270E7 | 4,153E7 | 5,233E7 | Cct7 |
| P60122 | RuvB-like 1 OS=Mus musculus GN=Ruvbl1 PE=1 SV=1 - [RUVB1_MOUSE] | 0,000E0 | 0,000E0 | 0,000E0 | 6,639E6 | 1,183E7 | 2,234E7 | Ruvbl1 |
| Q5F2E7 | Nuclear fragile X mental retardation-interacting protein 2 OS=Mus musculus GN=Nufip2 PE=1 SV=1 - [NUFP2_MOUSE] | 0,000E0 | 0,000E0 | 1,339E6 | 1,442E7 | 4,056E7 | 8,343E7 | Nufip2 |
| Q9D1I6 | 39S ribosomal protein L14, mitochondrial OS=Mus musculus GN=Mrpl14 PE=1 SV=1 - [RM14_MOUSE] | 0,000E0 | 0,000E0 | 0,000E0 | 8,072E6 | 2,486E7 | 1,484E7 | Mrpl14 |
| P70698 | CTP synthase 1 OS=Mus musculus GN=Ctps1 PE=1 SV=2 - [PYRG1_MOUSE] | 0,000E0 | 0,000E0 | 0,000E0 | 1,140E7 | 1,289E7 | 4,524E7 | Ctps1 |
| Q8BK63-2 | Isoform 2 of Casein kinase I isoform alpha OS=Mus musculus GN=Csnk1a1 - [KC1A_MOUSE] | 0,000E0 | 0,000E0 | 0,000E0 | 5,485E6 | 1,552E7 | 1,259E7 | Csnk1a1 |
| Q99L45 | Eukaryotic translation initiation factor 2 subunit 2 OS=Mus musculus GN=Eif2s2 PE=1 SV=1 - [IF2B_MOUSE] | 0,000E0 | 0,000E0 | 0,000E0 | 1,279E7 | 1,732E7 | 1,940E7 | Eif2s2 |
| P62315 | Small nuclear ribonucleoprotein Sm D1 OS=Mus musculus GN=Snrpd1 PE=1 SV=1 - [SMD1_MOUSE] | 0,000E0 | 0,000E0 | 2,089E6 | 9,435E6 | 1,590E7 | 2,582E7 | Snrpd1 |
| P62192 | 26S protease regulatory subunit 4 OS=Mus musculus GN=Psmc1 PE=1 SV=1 - [PRS4_MOUSE] | 0,000E0 | 0,000E0 | 0,000E0 | 5,738E6 | 1,003E7 | 1,440E7 | Psmc1 |
| P17918 | Proliferating cell nuclear antigen OS=Mus musculus GN=Pcna PE=1 SV=2 - [PCNA_MOUSE] | 0,000E0 | 0,000E0 | 8,188E5 | 1,183E7 | 1,534E7 | 2,127E7 | Pcna |
| Q3UQN2 | F-BAR domain only protein 2 OS=Mus musculus GN=Fcho2 PE=1 SV=1 - [FCHO2_MOUSE] | 0,000E0 | 0,000E0 | 0,000E0 | 3,931E7 | 6,535E7 | 6,284E7 | Fcho2 |
| Q99K51 | Plastin-3 OS=Mus musculus GN=Pls3 PE=1 SV=3 - [PLST_MOUSE] | 0,000E0 | 0,000E0 | 0,000E0 | 8,897E6 | 2,724E7 | 3,385E7 | Pls3 |
| Q9DB05 | Alpha-soluble NSF attachment protein OS=Mus musculus GN=Napa PE=1 SV=1 - [SNAA_MOUSE] | 0,000E0 | 0,000E0 | 0,000E0 | 7,134E6 | 8,585E6 | 1,656E7 | Napa |
| P57776-2 | Isoform 2 of Elongation factor 1-delta OS=Mus musculus GN=Eef1d - [EF1D_MOUSE] | 0,000E0 | 0,000E0 | 0,000E0 | 2,851E7 | 2,366E7 | 9,933E7 | Eef1d |
| Q9DCH4 | Eukaryotic translation initiation factor 3 subunit F OS=Mus musculus GN=Eif3f PE=1 SV=2 - [EIF3F_MOUSE] | 0,000E0 | 0,000E0 | 0,000E0 | 2,204E7 | 3,839E7 | 9,736E7 | Eif3f |
| Q8C7U1 | NEDD4-binding protein 3 OS=Mus musculus GN=N4bp3 PE=1 SV=1 - [N4BP3_MOUSE] | 0,000E0 | 0,000E0 | 0,000E0 | 1,985E7 | 4,326E7 | 4,841E7 | N4bp3 |
| P40124 | Adenylyl cyclase-associated protein 1 OS=Mus musculus GN=Cap1 PE=1 SV=4 - [CAP1_MOUSE] | 0,000E0 | 0,000E0 | 0,000E0 | 3,900E6 | 8,752E6 | 9,536E6 | Cap1 |
| P14115 | 60S ribosomal protein L27a OS=Mus musculus GN=Rpl27a PE=1 SV=5 - [RL27A_MOUSE] | 0,000E0 | 0,000E0 | 3,543E5 | 1,074E8 | 2,647E8 | 4,072E8 | Rpl27a |
| Q6A065 | Centrosomal protein of 170 kDa OS=Mus musculus GN=Cep170 PE=1 SV=2 - [CE170_MOUSE] | 0,000E0 | 0,000E0 | 2,926E5 | 3,143E7 | 3,321E7 | 3,629E7 | Cep170 |
| P25911-2 | Isoform 2 of Tyrosine-protein kinase Lyn OS=Mus musculus GN=Lyn - [LYN_MOUSE] | 0,000E0 | 0,000E0 | 0,000E0 | 1,185E7 | 2,666E7 | 6,034E7 | Lyn |
| P35293 | Ras-related protein Rab-18 OS=Mus musculus GN=Rab18 PE=1 SV=2 - [RAB18_MOUSE] | 0,000E0 | 0,000E0 | 0,000E0 | 3,896E6 | 2,097E7 | 1,646E7 | Rab18 |
| P26039 | Talin-1 OS=Mus musculus GN=Tln1 PE=1 SV=2 - [TLN1_MOUSE] | 0,000E0 | 0,000E0 | 0,000E0 | 1,185E7 | 3,566E7 | 6,755E7 | Tln1 |
| Q9D6Z1 | Nucleolar protein 56 OS=Mus musculus GN=Nop56 PE=1 SV=2 - [NOP56_MOUSE] | 0,000E0 | 0,000E0 | 0,000E0 | 5,334E6 | 1,353E7 | 3,920E7 | Nop56 |
| Q9R0P5 | Destrin OS=Mus musculus GN=Dstn PE=1 SV=3 - [DEST_MOUSE] | 0,000E0 | 0,000E0 | 0,000E0 | 7,401E6 | 4,813E7 | 5,375E7 | Dstn |
| Q922B2 | Aspartate--tRNA ligase, cytoplasmic OS=Mus musculus GN=Dars PE=1 SV=2 - [SYDC_MOUSE] | 0,000E0 | 0,000E0 | 0,000E0 | 8,969E6 | 2,554E7 | 5,697E7 | Dars |
| Q7TMK9-2 | Isoform 2 of Heterogeneous nuclear ribonucleoprotein Q OS=Mus musculus GN=Syncrip - [HNRPQ_MOUSE] | 0,000E0 | 0,000E0 | 0,000E0 | 3,354E7 | 6,045E7 | 1,287E8 | Syncrip |
| Q8R361 | Rab11 family-interacting protein 5 OS=Mus musculus GN=Rab11fip5 PE=1 SV=2 - [RFIP5_MOUSE] | 0,000E0 | 0,000E0 | 0,000E0 | 1,404E7 | 3,032E7 | 5,150E7 | Rab11fip5 |
| P46935 | E3 ubiquitin-protein ligase NEDD4 OS=Mus musculus GN=Nedd4 PE=1 SV=3 - [NEDD4_MOUSE] | 0,000E0 | 0,000E0 | 0,000E0 | 1,561E7 | 2,262E7 | 2,872E7 | Nedd4 |
| Q61166 | Microtubule-associated protein RP/EB family member 1 OS=Mus musculus GN=Mapre1 PE=1 SV=3 - [MARE1_MOUSE] | 0,000E0 | 0,000E0 | 0,000E0 | 4,351E6 | 1,225E7 | 1,624E7 | Mapre1 |
| Q8R411 | Myc target protein 1 OS=Mus musculus GN=Myct1 PE=1 SV=1 - [MYCT1_MOUSE] | 0,000E0 | 0,000E0 | 1,124E6 | 1,514E8 | 3,331E8 | 2,796E8 | Myct1 |
| Q9QYJ0 | DnaJ homolog subfamily A member 2 OS=Mus musculus GN=Dnaja2 PE=1 SV=1 - [DNJA2_MOUSE] | 0,000E0 | 0,000E0 | 0,000E0 | 2,477E7 | 5,809E7 | 8,467E7 | Dnaja2 |
| P35922-3 | Isoform 3 of Synaptic functional regulator FMR1 OS=Mus musculus GN=Fmr1 - [FMR1_MOUSE] | 0,000E0 | 0,000E0 | 1,048E6 | 6,566E7 | 1,602E8 | 3,140E8 | Fmr1 |
| Q9D8N0 | Elongation factor 1-gamma OS=Mus musculus GN=Eef1g PE=1 SV=3 - [EF1G_MOUSE] | 0,000E0 | 0,000E0 | 1,369E6 | 5,285E7 | 9,319E7 | 1,575E8 | Eef1g |
| Q9DC51 | Guanine nucleotide-binding protein G(k) subunit alpha OS=Mus musculus GN=Gnai3 PE=1 SV=3 - [GNAI3_MOUSE] | 0,000E0 | 0,000E0 | 7,794E4 | 3,831E7 | 1,634E8 | 7,500E7 | Gnai3 |
| P49312 | Heterogeneous nuclear ribonucleoprotein A1 OS=Mus musculus GN=Hnrnpa1 PE=1 SV=2 - [ROA1_MOUSE] | 0,000E0 | 0,000E0 | 2,262E6 | 6,352E7 | 6,110E7 | 1,429E8 | Hnrnpa1 |
| Q91W92 | Cdc42 effector protein 1 OS=Mus musculus GN=Cdc42ep1 PE=1 SV=1 - [BORG5_MOUSE] | 0,000E0 | 0,000E0 | 0,000E0 | 1,837E7 | 3,988E7 | 4,361E7 | Cdc42ep1 |
| P97379 | Ras GTPase-activating protein-binding protein 2 OS=Mus musculus GN=G3bp2 PE=1 SV=2 - [G3BP2_MOUSE] | 0,000E0 | 0,000E0 | 5,026E6 | 3,650E7 | 7,730E7 | 1,353E8 | G3bp2 |

| | | | | | | | | |
|---|---|---|---|---|---|---|---|---|
| P62874 | Guanine nucleotide-binding protein G(I)/G(S)/G(T) subunit beta-1 OS=Mus musculus GN=Gnb1 PE=1 SV=3 - [GBB1_MOUSE] | 0,000E0 | 0,000E0 | 2,238E6 | 8,151E7 | 1,097E8 | 1,052E8 | Gnb1 |
| Q9QZ06 | Toll-interacting protein OS=Mus musculus GN=Tollip PE=1 SV=1 - [TOLIP_MOUSE] | 0,000E0 | 0,000E0 | 0,000E0 | 8,920E6 | 3,551E7 | 4,536E7 | Tollip |
| Q501J6 | Probable ATP-dependent RNA helicase DDX17 OS=Mus musculus GN=Ddx17 PE=1 SV=1 - [DDX17_MOUSE] | 0,000E0 | 0,000E0 | 0,000E0 | 1,227E8 | 3,091E8 | 3,813E8 | Ddx17 |
| P56873 | Sjoegren syndrome/scleroderma autoantigen 1 homolog OS=Mus musculus GN=Sssca1 PE=1 SV=1 - [SSA27_MOUSE] | 0,000E0 | 0,000E0 | 0,000E0 | 4,624E6 | 8,224E6 | 8,401E6 | Sssca1 |
| Q91Z67 | SLIT-ROBO Rho GTPase-activating protein 2 OS=Mus musculus GN=Srgap2 PE=1 SV=2 - [SRGP2_MOUSE] | 0,000E0 | 0,000E0 | 3,662E5 | 3,400E7 | 5,991E7 | 5,075E7 | Srgap2 |
| Q6PGB6-3 | Isoform 3 of N-alpha-acetyltransferase 50 OS=Mus musculus GN=Naa50 - [NAA50_MOUSE] | 0,000E0 | 0,000E0 | 0,000E0 | 6,866E5 | 2,613E6 | 5,084E6 | Naa50 |
| Q60865 | Caprin-1 OS=Mus musculus GN=Caprin1 PE=1 SV=2 - [CAPR1_MOUSE] | 0,000E0 | 0,000E0 | 6,825E6 | 2,103E8 | 4,260E8 | 9,731E8 | Caprin1 |
| Q8CHP5-2 | Isoform 2 of Partner of Y14 and mago OS=Mus musculus GN=Pym1 - [PYM1_MOUSE] | 0,000E0 | 0,000E0 | 1,961E5 | 2,149E6 | 4,501E6 | 1,344E7 | Pym1 |
| Q9QXS6-3 | Isoform E2 of Drebrin OS=Mus musculus GN=Dbn1 - [DREB_MOUSE] | 0,000E0 | 0,000E0 | 0,000E0 | 5,168E7 | 1,054E8 | 2,197E8 | Dbn1 |
| Q8R1A4-2 | Isoform 2 of Dedicator of cytokinesis protein 7 OS=Mus musculus GN=Dock7 - [DOCK7_MOUSE] | 0,000E0 | 0,000E0 | 0,000E0 | 3,671E7 | 8,172E7 | 8,590E7 | Dock7 |
| P48024 | Eukaryotic translation initiation factor 1 OS=Mus musculus GN=Eif1 PE=1 SV=2 - [EIF1_MOUSE] | 0,000E0 | 0,000E0 | 0,000E0 | 3,642E5 | 1,994E6 | 8,159E6 | Eif1 |
| Q9D0B6 | Protein PBDC1 OS=Mus musculus GN=Pbdc1 PE=1 SV=1 - [PBDC1_MOUSE] | 0,000E0 | 0,000E0 | 0,000E0 | 4,026E5 | 3,951E6 | 7,323E6 | Pbdc1 |
| P62754 | 40S ribosomal protein S6 OS=Mus musculus GN=Rps6 PE=1 SV=1 - [RS6_MOUSE] | 0,000E0 | 0,000E0 | 2,070E6 | 6,530E7 | 2,832E8 | 7,990E8 | Rps6 |
| O35295 | Transcriptional activator protein Pur-beta OS=Mus musculus GN=Purb PE=1 SV=3 - [PURB_MOUSE] | 0,000E0 | 0,000E0 | 0,000E0 | 4,627E6 | 1,351E7 | 5,019E7 | Purb |
| P25911 | Tyrosine-protein kinase Lyn OS=Mus musculus GN=Lyn PE=1 SV=4 - [LYN_MOUSE] | 0,000E0 | 0,000E0 | 0,000E0 | 1,185E7 | 2,666E7 | 6,034E7 | Lyn |
| P58468 | Protein FAM207A OS=Mus musculus GN=Fam207a PE=1 SV=1 - [F207A_MOUSE] | 0,000E0 | 0,000E0 | 0,000E0 | 6,126E6 | 7,327E6 | 3,762E6 | Fam207a |
| O08992 | Syntenin-1 OS=Mus musculus GN=Sdcbp PE=1 SV=1 - [SDCB1_MOUSE] | 0,000E0 | 0,000E0 | 0,000E0 | 4,975E7 | 1,798E7 | 8,151E7 | Sdcbp |
| Q8K1N2 | Pleckstrin homology-like domain family B member 2 OS=Mus musculus GN=Phldb2 PE=1 SV=2 - [PHLB2_MOUSE] | 0,000E0 | 0,000E0 | 0,000E0 | 1,612E7 | 1,964E7 | 3,291E7 | Phldb2 |
| P63330 | Serine/threonine-protein phosphatase 2A catalytic subunit alpha isoform OS=Mus musculus GN=Ppp2ca PE=1 SV=1 - [PP2AA_MOUSE] | 0,000E0 | 0,000E0 | 0,000E0 | 1,422E7 | 1,728E7 | 1,129E7 | Ppp2ca |
| P83882 | 60S ribosomal protein L36a OS=Mus musculus GN=Rpl36a PE=1 SV=2 - [RL36A_MOUSE] | 0,000E0 | 0,000E0 | 3,401E6 | 1,254E8 | 2,063E8 | 5,486E8 | Rpl36a |
| O54824-2 | Isoform 2 of Pro-interleukin-16 OS=Mus musculus GN=Il16 - [IL16_MOUSE] | 0,000E0 | 0,000E0 | 0,000E0 | 2,722E7 | 5,224E7 | 5,908E7 | Il16 |
| O70325-2 | Isoform Cytoplasmic of Phospholipid hydroperoxide glutathione peroxidase, mitochondrial OS=Mus musculus GN=Gpx4 - [GPX41_MOUSE] | 0,000E0 | 0,000E0 | 0,000E0 | 5,742E6 | 1,710E6 | 1,312E7 | Gpx4 |
| O89086 | RNA-binding protein 3 OS=Mus musculus GN=Rbm3 PE=1 SV=1 - [RBM3_MOUSE] | 0,000E0 | 0,000E0 | 0,000E0 | 2,435E7 | 4,328E7 | 7,787E7 | Rbm3 |
| Q8CC35-2 | Isoform 2 of Synaptopodin OS=Mus musculus GN=Synpo - [SYNPO_MOUSE] | 0,000E0 | 0,000E0 | 0,000E0 | 8,920E6 | 1,585E7 | 3,134E7 | Synpo |
| Q9CQ65 | S-methyl-5'-thioadenosine phosphorylase OS=Mus musculus GN=Mtap PE=1 SV=1 - [MTAP_MOUSE] | 0,000E0 | 0,000E0 | 0,000E0 | 1,504E7 | 2,064E7 | 1,153E7 | Mtap |
| Q9DBY8 | Nuclear valosin-containing protein-like OS=Mus musculus GN=Nvl PE=1 SV=1 - [NVL_MOUSE] | 0,000E0 | 0,000E0 | 0,000E0 | 7,359E6 | 1,504E7 | 3,411E7 | Nvl |
| O54833 | Casein kinase II subunit alpha' OS=Mus musculus GN=Csnk2a2 PE=1 SV=1 - [CSK22_MOUSE] | 0,000E0 | 0,000E0 | 0,000E0 | 8,999E6 | 7,068E6 | 2,742E7 | Csnk2a2 |
| P55284 | Cadherin-5 OS=Mus musculus GN=Cdh5 PE=1 SV=2 - [CADH5_MOUSE] | 0,000E0 | 0,000E0 | 0,000E0 | 1,798E8 | 2,365E8 | 1,645E8 | Cdh5 |
| Q9QUR7 | Peptidyl-prolyl cis-trans isomerase NIMA-interacting 1 OS=Mus musculus GN=Pin1 PE=1 SV=1 - [PIN1_MOUSE] | 0,000E0 | 0,000E0 | 0,000E0 | 1,972E6 | 1,882E6 | 1,479E6 | Pin1 |
| Q9Z108 | Double-stranded RNA-binding protein Staufen homolog 1 OS=Mus musculus GN=Stau1 PE=1 SV=1 - [STAU1_MOUSE] | 0,000E0 | 0,000E0 | 0,000E0 | 1,767E7 | 3,046E7 | 5,957E7 | Stau1 |
| P27546-3 | Isoform 3 of Microtubule-associated protein 4 OS=Mus musculus GN=Map4 - [MAP4_MOUSE] | 0,000E0 | 0,000E0 | 0,000E0 | 6,365E6 | 2,015E7 | 2,844E7 | Map4 |
| Q9JJA9 | General receptor for phosphoinositides 1-associated scaffold protein OS=Mus musculus GN=Grasp PE=1 SV=2 - [GRASP_MOUSE] | 0,000E0 | 0,000E0 | 0,000E0 | 5,810E6 | 8,331E6 | 2,913E7 | Grasp |
| Q3UTJ2-3 | Isoform 3 of Sorbin and SH3 domain-containing protein 2 OS=Mus musculus GN=Sorbs2 - [SRBS2_MOUSE] | 0,000E0 | 0,000E0 | 0,000E0 | 3,852E6 | 1,294E7 | 1,957E7 | Sorbs2 |
| Q9D0T1 | NHP2-like protein 1 OS=Mus musculus GN=Snu13 PE=1 SV=4 - [NH2L1_MOUSE] | 0,000E0 | 0,000E0 | 0,000E0 | 3,638E6 | 5,437E6 | 1,946E7 | Snu13 |
| P12399 | Protein CTLA-2-alpha OS=Mus musculus GN=Ctla2a PE=2 SV=2 - [CTL2A_MOUSE] | 0,000E0 | 0,000E0 | 0,000E0 | 1,371E7 | 2,130E7 | 2,974E6 | Ctla2a |
| Q9WTX6 | Cullin-1 OS=Mus musculus GN=Cul1 PE=1 SV=1 - [CUL1_MOUSE] | 0,000E0 | 0,000E0 | 0,000E0 | 2,746E7 | 2,275E7 | 3,225E7 | Cul1 |
| P62267 | 40S ribosomal protein S23 OS=Mus musculus GN=Rps23 PE=1 SV=3 - [RS23_MOUSE] | 0,000E0 | 0,000E0 | 0,000E0 | 1,824E8 | 3,247E8 | 6,382E8 | Rps23 |
| Q921M7 | Protein FAM49B OS=Mus musculus GN=Fam49b PE=1 SV=1 - [FA49B_MOUSE] | 0,000E0 | 0,000E0 | 0,000E0 | 5,147E6 | 1,096E7 | 1,774E7 | Fam49b |
| Q8BFY7 | Protein FAM64A OS=Mus musculus GN=Fam64a PE=1 SV=1 - [FA64A_MOUSE] | 0,000E0 | 0,000E0 | 0,000E0 | 8,115E6 | 1,556E7 | 1,863E7 | Fam64a |
| Q9WUM3 | Coronin-1B OS=Mus musculus GN=Coro1b PE=1 SV=1 - [COR1B_MOUSE] | 0,000E0 | 0,000E0 | 0,000E0 | 7,533E6 | 3,962E7 | 7,708E7 | Coro1b |
| Q91Z96 | BMP-2-inducible protein kinase OS=Mus musculus GN=Bmp2k PE=1 SV=1 - [BMP2K_MOUSE] | 0,000E0 | 0,000E0 | 0,000E0 | 4,933E7 | 9,379E7 | 8,089E7 | Bmp2k |
| Q8CGB3 | Uveal autoantigen with coiled-coil domains and ankyrin repeats OS=Mus musculus GN=Uaca PE=1 SV=2 - [UACA_MOUSE] | 0,000E0 | 0,000E0 | 0,000E0 | 3,024E7 | 5,798E7 | 6,492E7 | Uaca |
| Q9CQD1 | Ras-related protein Rab-5A OS=Mus musculus GN=Rab5a PE=1 SV=1 - [RAB5A_MOUSE] | 0,000E0 | 0,000E0 | 0,000E0 | 1,978E6 | 2,563E7 | 3,682E7 | Rab5a |
| P62867 | 40S ribosomal protein S30 OS=Mus spicilegus GN=Fau PE=3 SV=1 - [RS30_MUSSI] | 0,000E0 | 0,000E0 | 0,000E0 | 3,120E7 | 2,086E8 | 2,241E8 | Fau |
| P70313 | Nitric oxide synthase, endothelial OS=Mus musculus GN=Nos3 PE=1 SV=4 - [NOS3_MOUSE] | 0,000E0 | 0,000E0 | 0,000E0 | 5,605E7 | 1,240E8 | 1,950E8 | Nos3 |
| Q6PHN9 | Ras-related protein Rab-35 OS=Mus musculus GN=Rab35 PE=1 SV=1 - [RAB35_MOUSE] | 0,000E0 | 0,000E0 | 3,456E5 | 6,493E7 | 1,449E8 | 2,096E8 | Rab35 |
| Q8C863 | E3 ubiquitin-protein ligase Itchy OS=Mus musculus GN=Itch PE=1 SV=2 - [ITCH_MOUSE] | 0,000E0 | 0,000E0 | 0,000E0 | 1,676E7 | 5,431E7 | 5,098E7 | Itch |
| Q9JLB2 | MAGUK p55 subfamily member 5 OS=Mus musculus GN=Mpp5 PE=1 SV=1 - [MPP5_MOUSE] | 0,000E0 | 0,000E0 | 0,000E0 | 1,155E7 | 1,314E7 | 6,369E6 | Mpp5 |
| Q6PAM1 | Alpha-taxilin OS=Mus musculus GN=Txlna PE=1 SV=1 - [TXLNA_MOUSE] | 0,000E0 | 0,000E0 | 0,000E0 | 1,945E7 | 2,437E7 | 4,189E7 | Txlna |
| Q99020 | Heterogeneous nuclear ribonucleoprotein A/B OS=Mus musculus GN=Hnrnpab PE=1 SV=1 - [ROAA_MOUSE] | 0,000E0 | 0,000E0 | 1,457E6 | 1,603E7 | 6,814E7 | 1,107E8 | Hnrnpab |
| Q925I1 | ATPase family AAA domain-containing protein 3 OS=Mus musculus GN=Atad3 PE=1 SV=1 - [ATAD3_MOUSE] | 0,000E0 | 0,000E0 | 0,000E0 | 1,707E7 | 3,857E7 | 4,622E7 | Atad3 |
| P15864 | Histone H1.2 OS=Mus musculus GN=Hist1h1c PE=1 SV=2 - [H12_MOUSE] | 0,000E0 | 0,000E0 | 6,982E5 | 6,443E7 | 1,356E8 | 3,936E8 | Hist1h1c |
| Q80X85 | 28S ribosomal protein S7, mitochondrial OS=Mus musculus GN=Mrps7 PE=1 SV=1 - [RT07_MOUSE] | 0,000E0 | 0,000E0 | 0,000E0 | 2,532E6 | 4,189E6 | 6,744E6 | Mrps7 |
| Q9D824-4 | Isoform 4 of Pre-mRNA 3'-end-processing factor FIP1 OS=Mus musculus GN=Fip1l1 - [FIP1_MOUSE] | 0,000E0 | 0,000E0 | 0,000E0 | 5,832E6 | 1,672E7 | 8,311E6 | Fip1l1 |
| Q3UZ39-2 | Isoform 2 of Leucine-rich repeat flightless-interacting protein 1 OS=Mus musculus GN=Lrrfip1 - [LRRF1_MOUSE] | 0,000E0 | 0,000E0 | 0,000E0 | 4,588E7 | 9,723E7 | 1,245E8 | Lrrfip1 |
| O35643 | AP-1 complex subunit beta-1 OS=Mus musculus GN=Ap1b1 PE=1 SV=2 - [AP1B1_MOUSE] | 0,000E0 | 0,000E0 | 0,000E0 | 2,272E8 | 3,054E8 | 3,824E8 | Ap1b1 |
| Q9CSN1 | SNW domain-containing protein 1 OS=Mus musculus GN=Snw1 PE=1 SV=3 - [SNW1_MOUSE] | 0,000E0 | 0,000E0 | 0,000E0 | 3,256E6 | 5,869E6 | 1,096E7 | Snw1 |
| Q8CI11 | Guanine nucleotide-binding protein-like 3 OS=Mus musculus GN=Gnl3 PE=1 SV=2 - [GNL3_MOUSE] | 0,000E0 | 0,000E0 | 0,000E0 | 1,329E7 | 2,000E7 | 4,750E7 | Gnl3 |
| Q8VEK3 | Heterogeneous nuclear ribonucleoprotein U OS=Mus musculus GN=Hnrnpu PE=1 SV=1 - [HNRPU_MOUSE] | 0,000E0 | 0,000E0 | 0,000E0 | 5,206E7 | 1,019E8 | 9,049E7 | Hnrnpu |
| P34022 | Ran-specific GTPase-activating protein OS=Mus musculus GN=Ranbp1 PE=1 SV=2 - [RANG_MOUSE] | 0,000E0 | 0,000E0 | 0,000E0 | 5,821E6 | 2,511E7 | 1,554E7 | Ranbp1 |
| Q8CBW3-5 | Isoform 5 of Abl interactor 1 OS=Mus musculus GN=Abi1 - [ABI1_MOUSE] | 0,000E0 | 0,000E0 | 0,000E0 | 1,595E7 | 5,288E7 | 6,136E7 | Abi1 |
| Q9Z315 | U4/U6.U5 tri-snRNP-associated protein 1 OS=Mus musculus GN=Sart1 PE=1 SV=1 - [SNUT1_MOUSE] | 0,000E0 | 0,000E0 | 0,000E0 | 4,632E6 | 7,366E6 | 1,288E7 | Sart1 |
| Q3UJB9-2 | Isoform 2 of Enhancer of mRNA-decapping protein 4 OS=Mus musculus GN=Edc4 - [EDC4_MOUSE] | 0,000E0 | 0,000E0 | 0,000E0 | 1,246E7 | 3,479E7 | 5,171E7 | Edc4 |
| Q60631 | Growth factor receptor-bound protein 2 OS=Mus musculus GN=Grb2 PE=1 SV=1 - [GRB2_MOUSE] | 0,000E0 | 0,000E0 | 0,000E0 | 6,084E6 | 2,165E7 | 6,661E6 | Grb2 |
| Q91VH2 | Sorting nexin-9 OS=Mus musculus GN=Snx9 PE=1 SV=1 - [SNX9_MOUSE] | 0,000E0 | 0,000E0 | 0,000E0 | 4,403E7 | 6,422E7 | 1,053E8 | Snx9 |
| Q9QYI3 | DnaJ homolog subfamily C member 7 OS=Mus musculus GN=Dnajc7 PE=1 SV=2 - [DNJC7_MOUSE] | 0,000E0 | 0,000E0 | 0,000E0 | 3,884E6 | 1,192E7 | 9,746E6 | Dnajc7 |
| Q9JM76 | Actin-related protein 2/3 complex subunit 3 OS=Mus musculus GN=Arpc3 PE=1 SV=3 - [ARPC3_MOUSE] | 0,000E0 | 0,000E0 | 7,380E5 | 9,633E7 | 1,244E8 | 4,962E7 | Arpc3 |
| Q8VDN2 | Sodium/potassium-transporting ATPase subunit alpha-1 OS=Mus musculus GN=Atp1a1 PE=1 SV=1 - [AT1A1_MOUSE] | 0,000E0 | 0,000E0 | 0,000E0 | 1,542E7 | 3,870E7 | 5,302E7 | Atp1a1 |
| Q5SV80 | Unconventional myosin-XIX OS=Mus musculus GN=Myo19 PE=2 SV=1 - [MYO19_MOUSE] | 0,000E0 | 0,000E0 | 0,000E0 | 1,546E7 | 4,680E7 | 1,137E8 | Myo19 |
| Q8BP92 | Reticulocalbin-2 OS=Mus musculus GN=Rcn2 PE=1 SV=1 - [RCN2_MOUSE] | 0,000E0 | 0,000E0 | 0,000E0 | 6,868E6 | 7,837E6 | 9,801E6 | Rcn2 |
| Q3UPF5 | Zinc finger CCCH-type antiviral protein 1 OS=Mus musculus GN=Zc3hav1 PE=1 SV=1 - [ZCCHV_MOUSE] | 0,000E0 | 0,000E0 | 0,000E0 | 1,162E7 | 3,385E7 | 5,499E7 | Zc3hav1 |
| Q9D7X3 | Dual specificity protein phosphatase 3 OS=Mus musculus GN=Dusp3 PE=1 SV=1 - [DUS3_MOUSE] | 0,000E0 | 0,000E0 | 0,000E0 | 5,026E6 | 1,652E7 | 1,630E7 | Dusp3 |
| Q8BGD9 | Eukaryotic translation initiation factor 4B OS=Mus musculus GN=Eif4b PE=1 SV=1 - [IF4B_MOUSE] | 0,000E0 | 0,000E0 | 0,000E0 | 2,896E7 | 7,821E7 | 1,342E8 | Eif4b |
| P19324 | Serpin H1 OS=Mus musculus GN=Serpinh1 PE=1 SV=3 - [SERPH_MOUSE] | 0,000E0 | 0,000E0 | 0,000E0 | 1,283E7 | 2,961E7 | 6,453E7 | Serpinh1 |
| Q99JX4 | Eukaryotic translation initiation factor 3 subunit M OS=Mus musculus GN=Eif3m PE=1 SV=1 - [EIF3M_MOUSE] | 0,000E0 | 0,000E0 | 0,000E0 | 2,294E6 | 1,269E7 | 2,249E7 | Eif3m |
| Q99K43 | Protein regulator of cytokinesis 1 OS=Mus musculus GN=Prc1 PE=1 SV=2 - [PRC1_MOUSE] | 0,000E0 | 0,000E0 | 0,000E0 | 9,965E6 | 2,177E7 | 5,021E7 | Prc1 |
| Q61510 | E3 ubiquitin/ISG15 ligase TRIM25 OS=Mus musculus GN=Trim25 PE=1 SV=2 - [TRI25_MOUSE] | 0,000E0 | 0,000E0 | 0,000E0 | 3,692E6 | 9,474E6 | 1,924E7 | Trim25 |
| P27773 | Protein disulfide-isomerase A3 OS=Mus musculus GN=Pdia3 PE=1 SV=2 - [PDIA3_MOUSE] | 0,000E0 | 0,000E0 | 0,000E0 | 7,711E6 | 1,758E7 | 2,700E7 | Pdia3 |
| Q9JMB0 | G kinase-anchoring protein 1 OS=Mus musculus GN=Gkap1 PE=1 SV=1 - [GKAP1_MOUSE] | 0,000E0 | 0,000E0 | 0,000E0 | 4,392E6 | 8,445E6 | 2,065E7 | Gkap1 |
| Q9CQW9 | Interferon-induced transmembrane protein 3 OS=Mus musculus GN=Ifitm3 PE=1 SV=1 - [IFM3_MOUSE] | 0,000E0 | 0,000E0 | 0,000E0 | 5,875E7 | 1,817E8 | 3,251E8 | Ifitm3 |
| P49710 | Hematopoietic lineage cell-specific protein OS=Mus musculus GN=Hcls1 PE=1 SV=2 - [HCLS1_MOUSE] | 0,000E0 | 0,000E0 | 0,000E0 | 5,149E6 | 3,039E7 | 4,934E7 | Hcls1 |
| P63024 | Vesicle-associated membrane protein 3 OS=Mus musculus GN=Vamp3 PE=1 SV=1 - [VAMP3_MOUSE] | 0,000E0 | 0,000E0 | 0,000E0 | 8,104E6 | 2,040E7 | 3,192E7 | Vamp3 |
| Q61990-2 | Isoform 2 of Poly(rC)-binding protein 2 OS=Mus musculus GN=Pcbp2 - [PCBP2_MOUSE] | 0,000E0 | 0,000E0 | 0,000E0 | 2,185E7 | 4,015E7 | 1,195E8 | Pcbp2 |
| Q99KN9 | Clathrin interactor 1 OS=Mus musculus GN=Clint1 PE=1 SV=2 - [EPN4_MOUSE] | 0,000E0 | 0,000E0 | 0,000E0 | 5,761E7 | 1,243E8 | 1,776E8 | Clint1 |
| Q9WV95 | Pleckstrin homology-like domain family A member 3 OS=Mus musculus GN=Phlda3 PE=1 SV=1 - [PHLA3_MOUSE] | 0,000E0 | 0,000E0 | 0,000E0 | 8,774E6 | 9,073E6 | 2,023E7 | Phlda3 |
| P98083-2 | Isoform p52Shc of SHC-transforming protein 1 OS=Mus musculus GN=Shc1 - [SHC1_MOUSE] | 0,000E0 | 0,000E0 | 0,000E0 | 2,970E5 | 9,397E6 | 8,370E6 | Shc1 |
| Q6A0D4 | Raftlin OS=Mus musculus GN=Rftn1 PE=1 SV=4 - [RFTN1_MOUSE] | 0,000E0 | 0,000E0 | 2,147E5 | 3,394E6 | 1,906E7 | 2,461E7 | Rftn1 |
| P24369 | Peptidyl-prolyl cis-trans isomerase B OS=Mus musculus GN=Ppib PE=1 SV=2 - [PPIB_MOUSE] | 0,000E0 | 0,000E0 | 0,000E0 | 2,723E6 | 4,032E6 | 9,577E6 | Ppib |
| Q7TMB8 | Cytoplasmic FMR1-interacting protein 1 OS=Mus musculus GN=Cyfip1 PE=1 SV=1 - [CYFP1_MOUSE] | 0,000E0 | 0,000E0 | 1,056E6 | 3,183E7 | 7,211E7 | 1,079E8 | Cyfip1 |
| Q8CGB6-2 | Isoform 2 of Tensin-2 OS=Mus musculus GN=Tns2 - [TNS2_MOUSE] | 0,000E0 | 0,000E0 | 0,000E0 | 1,830E7 | 2,899E7 | 6,879E7 | Tns2 |
| Q80UG5-3 | Isoform 3 of Septin-9 OS=Mus musculus GN=Sept9 - [SEPT9_MOUSE] | 0,000E0 | 0,000E0 | 0,000E0 | 9,735E6 | 3,039E7 | 4,188E7 | SEPT9 |
| Q8BIE6 | FERM domain-containing protein 4A OS=Mus musculus GN=Frmd4a PE=1 SV=2 - [FRM4A_MOUSE] | 0,000E0 | 0,000E0 | 0,000E0 | 1,217E7 | 2,155E7 | 3,558E7 | Frmd4a |
| P21279 | Guanine nucleotide-binding protein G(q) subunit alpha OS=Mus musculus GN=Gnaq PE=1 SV=4 - [GNAQ_MOUSE] | 0,000E0 | 0,000E0 | 0,000E0 | 7,952E6 | 1,498E7 | 6,855E7 | Gnaq |

| | | | | | | | | |
|---|---|---|---|---|---|---|---|---|
| Q9CR26 | Vacuolar protein sorting-associated protein VTA1 homolog OS=Mus musculus GN=Vta1 PE=1 SV=1 - [VTA1_MOUSE] | 0,000E0 | 0,000E0 | 0,000E0 | 7,016E6 | 2,514E7 | 2,132E7 | Vta1 |
| Q8CGC4 | Protein LSM14 homolog B OS=Mus musculus GN=Lsm14b PE=1 SV=3 - [LS14B_MOUSE] | 0,000E0 | 0,000E0 | 0,000E0 | 5,806E6 | 1,099E7 | 2,024E7 | Lsm14b |
| O09131 | Glutathione S-transferase omega-1 OS=Mus musculus GN=Gsto1 PE=1 SV=2 - [GSTO1_MOUSE] | 0,000E0 | 0,000E0 | 0,000E0 | 2,292E6 | 6,208E6 | 3,593E6 | Gsto1 |
| Q8C8U0-3 | Isoform 3 of Liprin-beta-1 OS=Mus musculus GN=Ppfibp1 - [LIPB1_MOUSE] | 0,000E0 | 0,000E0 | 0,000E0 | 6,686E6 | 2,453E7 | 2,451E7 | Ppfibp1 |
| Q9Z0N1 | Eukaryotic translation initiation factor 2 subunit 3, X-linked OS=Mus musculus GN=Eif2s3x PE=1 SV=2 - [IF2G_MOUSE] | 0,000E0 | 0,000E0 | 0,000E0 | 2,187E7 | 4,219E7 | 1,079E8 | Eif2s3x |
| Q3TWW8 | Serine/arginine-rich splicing factor 6 OS=Mus musculus GN=Srsf6 PE=1 SV=1 - [SRSF6_MOUSE] | 0,000E0 | 0,000E0 | 0,000E0 | 6,749E7 | 9,168E7 | 1,236E8 | Srsf6 |
| Q9DC28-2 | Isoform 2 of Casein kinase I isoform delta OS=Mus musculus GN=Csnk1d - [KC1D_MOUSE] | 0,000E0 | 0,000E0 | 0,000E0 | 1,802E7 | 1,701E7 | 5,040E7 | Csnk1d |
| Q9Z1Z2 | Serine-threonine kinase receptor-associated protein OS=Mus musculus GN=Strap PE=1 SV=2 - [STRAP_MOUSE] | 0,000E0 | 0,000E0 | 0,000E0 | 2,811E6 | 9,091E6 | 1,280E7 | Strap |
| Q9D0I9 | Arginine--tRNA ligase, cytoplasmic OS=Mus musculus GN=Rars PE=1 SV=2 - [SYRC_MOUSE] | 0,000E0 | 0,000E0 | 0,000E0 | 1,698E7 | 4,400E7 | 4,834E7 | Rars |
| Q7TSC1 | Protein PRRC2A OS=Mus musculus GN=Prrc2a PE=1 SV=1 - [PRC2A_MOUSE] | 0,000E0 | 0,000E0 | 0,000E0 | 1,548E7 | 2,932E7 | 5,129E7 | Prrc2a |
| Q9Z1X4-2 | Isoform 2 of Interleukin enhancer-binding factor 3 OS=Mus musculus GN=Ilf3 - [ILF3_MOUSE] | 0,000E0 | 0,000E0 | 0,000E0 | 1,067E7 | 2,008E7 | 3,246E7 | Ilf3 |
| P08032 | Spectrin alpha chain, erythrocytic 1 OS=Mus musculus GN=Spta1 PE=1 SV=3 - [SPTA1_MOUSE] | 0,000E0 | 0,000E0 | 0,000E0 | 5,728E6 | 3,475E7 | 1,557E7 | Spta1 |
| P59325 | Eukaryotic translation initiation factor 5 OS=Mus musculus GN=Eif5 PE=1 SV=1 - [IF5_MOUSE] | 0,000E0 | 0,000E0 | 0,000E0 | 1,703E7 | 1,619E7 | 1,360E7 | Eif5 |
| Q9QXX4 | Calcium-binding mitochondrial carrier protein Aralar2 OS=Mus musculus GN=Slc25a13 PE=1 SV=1 - [CMC2_MOUSE] | 0,000E0 | 0,000E0 | 0,000E0 | 4,422E6 | 6,795E6 | 1,449E7 | Slc25a13 |
| P70452 | Syntaxin-4 OS=Mus musculus GN=Stx4 PE=1 SV=1 - [STX4_MOUSE] | 0,000E0 | 0,000E0 | 0,000E0 | 1,853E7 | 3,916E7 | 6,333E7 | Stx4 |
| Q9D735 | Uncharacterized protein C19orf43 homolog OS=Mus musculus PE=1 SV=1 - [CS043_MOUSE] | 0,000E0 | 0,000E0 | 0,000E0 | 2,063E6 | 1,013E7 | 2,284E6 | CS043 |
| Q91VC3 | Eukaryotic initiation factor 4A-III OS=Mus musculus GN=Eif4a3 PE=1 SV=3 - [IF4A3_MOUSE] | 0,000E0 | 0,000E0 | 0,000E0 | 1,511E7 | 9,340E7 | 6,129E7 | Eif4a3 |
| Q7M6Y3-6 | Isoform 6 of Phosphatidylinositol-binding clathrin assembly protein OS=Mus musculus GN=Picalm - [PICAL_MOUSE] | 0,000E0 | 0,000E0 | 0,000E0 | 5,366E7 | 5,766E7 | 8,626E7 | Picalm |
| O08553 | Dihydropyrimidinase-related protein 2 OS=Mus musculus GN=Dpysl2 PE=1 SV=2 - [DPYL2_MOUSE] | 0,000E0 | 0,000E0 | 0,000E0 | 3,094E6 | 5,656E6 | 2,043E7 | Dpysl2 |
| Q99J72-4 | Isoform 4 of DNA dC->dU-editing enzyme APOBEC-3 OS=Mus musculus GN=Apobec3 - [ABEC3_MOUSE] | 0,000E0 | 0,000E0 | 0,000E0 | 1,687E7 | 3,229E7 | 6,607E7 | Apobec3 |
| P62835 | Ras-related protein Rap-1A OS=Mus musculus GN=Rap1a PE=1 SV=1 - [RAP1A_MOUSE] | 0,000E0 | 0,000E0 | 0,000E0 | 1,164E7 | 4,687E7 | 6,988E7 | Rap1a |
| Q6PHZ2-2 | Isoform 2 of Calcium/calmodulin-dependent protein kinase type II subunit delta OS=Mus musculus GN=Camk2d - [KCC2D_MOUSE] | 0,000E0 | 0,000E0 | 0,000E0 | 1,122E7 | 2,205E7 | 5,066E7 | Camk2d |
| Q3UMF0-3 | Isoform 3 of Cordon-bleu protein-like 1 OS=Mus musculus GN=Cobll1 - [COBL1_MOUSE] | 0,000E0 | 0,000E0 | 0,000E0 | 5,329E6 | 8,762E6 | 1,537E7 | Cobll1 |
| P46938-2 | Isoform 2 of Transcriptional coactivator YAP1 OS=Mus musculus GN=Yap1 - [YAP1_MOUSE] | 0,000E0 | 0,000E0 | 0,000E0 | 2,125E7 | 2,034E7 | 2,711E7 | Yap1 |
| O08664 | B-cell CLL/lymphoma 7 protein family member C OS=Mus musculus GN=Bcl7c PE=1 SV=1 - [BCL7C_MOUSE] | 0,000E0 | 0,000E0 | 0,000E0 | 4,896E6 | 1,098E7 | 2,133E7 | Bcl7c |
| Q99LJ0 | CTTNBP2 N-terminal-like protein OS=Mus musculus GN=Cttnbp2nl PE=1 SV=1 - [CT2NL_MOUSE] | 0,000E0 | 0,000E0 | 0,000E0 | 4,181E6 | 1,498E7 | 1,145E7 | Cttnbp2nl |
| Q9CZT6 | Protein CMSS1 OS=Mus musculus GN=Cmss1 PE=2 SV=1 - [CMS1_MOUSE] | 0,000E0 | 0,000E0 | 0,000E0 | 4,121E6 | 6,794E6 | 6,962E6 | Cmss1 |
| O55131 | Septin-7 OS=Mus musculus GN=Sept7 PE=1 SV=1 - [SEPT7_MOUSE] | 0,000E0 | 0,000E0 | 0,000E0 | 2,813E7 | 6,631E7 | 6,870E7 | SEPT7 |
| Q8CAQ8-2 | Isoform 2 of MICOS complex subunit Mic60 OS=Mus musculus GN=Immt - [MIC60_MOUSE] | 0,000E0 | 0,000E0 | 1,006E6 | 7,826E6 | 2,905E7 | 4,285E7 | Immt |
| O35608 | Angiopoietin-2 OS=Mus musculus GN=Angpt2 PE=2 SV=2 - [ANGP2_MOUSE] | 0,000E0 | 0,000E0 | 0,000E0 | 1,818E7 | 1,822E7 | 4,407E7 | Angpt2 |
| Q8BHL4 | Retinoic acid-induced protein 3 OS=Mus musculus GN=Gprc5a PE=1 SV=1 - [RAI3_MOUSE] | 0,000E0 | 0,000E0 | 0,000E0 | 3,834E7 | 9,301E7 | 1,499E8 | Gprc5a |
| Q9CQ22 | Ragulator complex protein LAMTOR1 OS=Mus musculus GN=Lamtor1 PE=1 SV=1 - [LTOR1_MOUSE] | 0,000E0 | 0,000E0 | 0,000E0 | 5,700E6 | 2,655E6 | 1,469E7 | Lamtor1 |
| Q8R1B4 | Eukaryotic translation initiation factor 3 subunit C OS=Mus musculus GN=Eif3c PE=1 SV=1 - [EIF3C_MOUSE] | 0,000E0 | 0,000E0 | 1,214E6 | 3,060E7 | 4,183E7 | 8,884E7 | Eif3c |
| Q9CR57 | 60S ribosomal protein L14 OS=Mus musculus GN=Rpl14 PE=1 SV=3 - [RL14_MOUSE] | 0,000E0 | 0,000E0 | 2,375E5 | 9,775E7 | 1,658E8 | 3,495E8 | Rpl14 |
| Q9R0Y5 | Adenylate kinase isoenzyme 1 OS=Mus musculus GN=Ak1 PE=1 SV=1 - [KAD1_MOUSE] | 0,000E0 | 0,000E0 | 0,000E0 | 1,613E6 | 2,610E6 | 3,376E6 | Ak1 |
| Q99PU8-2 | Isoform 2 of Putative ATP-dependent RNA helicase DHX30 OS=Mus musculus GN=Dhx30 - [DHX30_MOUSE] | 0,000E0 | 0,000E0 | 0,000E0 | 6,139E6 | 1,251E7 | 2,627E7 | Dhx30 |
| Q8BK67 | Protein RCC2 OS=Mus musculus GN=Rcc2 PE=1 SV=1 - [RCC2_MOUSE] | 0,000E0 | 0,000E0 | 0,000E0 | 1,435E6 | 7,118E6 | 1,014E7 | Rcc2 |
| Q9DC28 | Casein kinase I isoform delta OS=Mus musculus GN=Csnk1d PE=1 SV=2 - [KC1D_MOUSE] | 0,000E0 | 0,000E0 | 0,000E0 | 1,802E7 | 1,827E7 | 5,035E7 | Csnk1d |
| P21278 | Guanine nucleotide-binding protein subunit alpha-11 OS=Mus musculus GN=Gna11 PE=1 SV=1 - [GNA11_MOUSE] | 0,000E0 | 0,000E0 | 0,000E0 | 1,313E7 | 2,392E7 | 5,427E7 | Gna11 |
| P35601-2 | Isoform 2 of Replication factor C subunit 1 OS=Mus musculus GN=Rfc1 - [RFC1_MOUSE] | 0,000E0 | 0,000E0 | 5,241E5 | 1,702E7 | 3,416E7 | 4,237E7 | Rfc1 |
| Q08481 | Platelet endothelial cell adhesion molecule OS=Mus musculus GN=Pecam1 PE=1 SV=1 - [PECA1_MOUSE] | 0,000E0 | 0,000E0 | 0,000E0 | 1,680E7 | 1,080E8 | 1,578E8 | Pecam1 |
| Q9WVM1 | Rac GTPase-activating protein 1 OS=Mus musculus GN=Racgap1 PE=1 SV=1 - [RGAP1_MOUSE] | 0,000E0 | 0,000E0 | 0,000E0 | 6,677E6 | 1,981E7 | 7,133E7 | Racgap1 |
| P13020-2 | Isoform 2 of Gelsolin OS=Mus musculus GN=Gsn - [GELS_MOUSE] | 0,000E0 | 0,000E0 | 9,668E6 | 1,955E7 | 4,489E7 | 8,300E7 | Gsn |
| Q7M6Y3-5 | Isoform 5 of Phosphatidylinositol-binding clathrin assembly protein OS=Mus musculus GN=Picalm - [PICAL_MOUSE] | 0,000E0 | 0,000E0 | 0,000E0 | 6,471E7 | 1,028E8 | 1,098E8 | Picalm |
| P63028 | Translationally-controlled tumor protein OS=Mus musculus GN=Tpt1 PE=1 SV=1 - [TCTP_MOUSE] | 0,000E0 | 0,000E0 | 0,000E0 | 6,326E6 | 3,679E7 | 9,994E6 | Tpt1 |
| Q99J62 | Replication factor C subunit 4 OS=Mus musculus GN=Rfc4 PE=1 SV=1 - [RFC4_MOUSE] | 0,000E0 | 0,000E0 | 0,000E0 | 3,254E6 | 2,363E7 | 4,087E7 | Rfc4 |
| Q6P542 | ATP-binding cassette sub-family F member 1 OS=Mus musculus GN=Abcf1 PE=1 SV=1 - [ABCF1_MOUSE] | 0,000E0 | 0,000E0 | 0,000E0 | 2,163E7 | 3,402E7 | 6,852E7 | Abcf1 |
| Q9WTX2 | Interferon-inducible double-stranded RNA-dependent protein kinase activator A OS=Mus musculus GN=Prkra PE=1 SV=1 - [PRKRA_MOUSE] | 0,000E0 | 0,000E0 | 0,000E0 | 8,709E6 | 7,977E6 | 3,095E7 | Prkra |
| P19157 | Glutathione S-transferase P 1 OS=Mus musculus GN=Gstp1 PE=1 SV=2 - [GSTP1_MOUSE] | 0,000E0 | 0,000E0 | 0,000E0 | 6,291E6 | 2,846E7 | 1,988E7 | Gstp1 |
| Q1W617 | Protein Shroom4 OS=Mus musculus GN=Shroom4 PE=1 SV=1 - [SHRM4_MOUSE] | 0,000E0 | 0,000E0 | 0,000E0 | 1,488E7 | 4,474E7 | 7,397E7 | Shroom4 |
| Q8K298 | Anillin OS=Mus musculus GN=Anln PE=1 SV=2 - [ANLN_MOUSE] | 0,000E0 | 0,000E0 | 0,000E0 | 6,823E6 | 1,148E7 | 1,416E7 | Anln |
| Q8VH51-2 | Isoform 2 of RNA-binding protein 39 OS=Mus musculus GN=Rbm39 - [RBM39_MOUSE] | 1,010E7 | 0,000E0 | 2,877E6 | 6,717E7 | 1,269E8 | 2,173E8 | Rbm39 |
| Q8VDR9 | Dedicator of cytokinesis protein 6 OS=Mus musculus GN=Dock6 PE=1 SV=4 - [DOCK6_MOUSE] | 0,000E0 | 0,000E0 | 0,000E0 | 2,811E7 | 5,121E7 | 6,504E7 | Dock6 |
| Q6A028 | Switch-associated protein 70 OS=Mus musculus GN=Swap70 PE=1 SV=2 - [SWP70_MOUSE] | 0,000E0 | 0,000E0 | 0,000E0 | 7,522E6 | 1,159E7 | 2,236E7 | Swap70 |
| Q9D1R9 | 60S ribosomal protein L34 OS=Mus musculus GN=Rpl34 PE=1 SV=2 - [RL34_MOUSE] | 0,000E0 | 0,000E0 | 0,000E0 | 9,653E7 | 3,124E8 | 6,352E8 | Rpl34 |
| Q9ESD7-2 | Isoform 2 of Dysferlin OS=Mus musculus GN=Dysf - [DYSF_MOUSE] | 0,000E0 | 0,000E0 | 0,000E0 | 1,472E7 | 2,247E7 | 1,128E8 | Dysf |
| P54775 | 26S protease regulatory subunit 6B OS=Mus musculus GN=Psmc4 PE=1 SV=2 - [PRS6B_MOUSE] | 0,000E0 | 0,000E0 | 9,752E5 | 3,711E6 | 6,879E6 | 2,076E6 | Psmc4 |
| Q64010-2 | Isoform Crk-I of Adapter molecule crk OS=Mus musculus GN=Crk - [CRK_MOUSE] | 0,000E0 | 0,000E0 | 0,000E0 | 1,024E6 | 3,410E6 | 6,294E6 | Crk |
| Q60864 | Stress-induced-phosphoprotein 1 OS=Mus musculus GN=Stip1 PE=1 SV=1 - [STIP1_MOUSE] | 0,000E0 | 0,000E0 | 0,000E0 | 5,908E6 | 1,515E7 | 1,678E7 | Stip1 |
| Q62465 | Synaptic vesicle membrane protein VAT-1 homolog OS=Mus musculus GN=Vat1 PE=1 SV=3 - [VAT1_MOUSE] | 0,000E0 | 0,000E0 | 0,000E0 | 1,740E7 | 3,133E7 | 3,327E7 | Vat1 |
| Q9D0F6 | Replication factor C subunit 5 OS=Mus musculus GN=Rfc5 PE=1 SV=1 - [RFC5_MOUSE] | 0,000E0 | 0,000E0 | 0,000E0 | 6,434E6 | 1,869E7 | 3,372E7 | Rfc5 |
| Q8BMA3 | Connector enhancer of kinase suppressor of ras 3 OS=Mus musculus GN=Cnksr3 PE=1 SV=1 - [CNKR3_MOUSE] | 0,000E0 | 0,000E0 | 0,000E0 | 3,575E6 | 3,502E7 | 1,434E7 | Cnksr3 |
| Q76MZ3 | Serine/threonine-protein phosphatase 2A 65 kDa regulatory subunit A alpha isoform OS=Mus musculus GN=Ppp2r1a PE=1 SV=3 - [2AAA_MOUSE] | 0,000E0 | 0,000E0 | 0,000E0 | 2,808E7 | 3,379E7 | 6,792E7 | Ppp2r1a |
| Q9ESD7 | Dysferlin OS=Mus musculus GN=Dysf PE=1 SV=3 - [DYSF_MOUSE] | 0,000E0 | 0,000E0 | 0,000E0 | 1,472E7 | 2,225E7 | 1,128E8 | Dysf |
| P52927 | High mobility group protein HMGI-C OS=Mus musculus GN=Hmga2 PE=1 SV=1 - [HMGA2_MOUSE] | 0,000E0 | 0,000E0 | 0,000E0 | 4,944E7 | 8,353E7 | 2,038E8 | Hmga2 |
| O70492 | Sorting nexin-3 OS=Mus musculus GN=Snx3 PE=1 SV=3 - [SNX3_MOUSE] | 0,000E0 | 0,000E0 | 0,000E0 | 7,197E5 | 1,315E7 | 2,166E7 | Snx3 |
| P26883 | Peptidyl-prolyl cis-trans isomerase FKBP1A OS=Mus musculus GN=Fkbp1a PE=1 SV=2 - [FKB1A_MOUSE] | 0,000E0 | 0,000E0 | 0,000E0 | 1,798E5 | 2,116E7 | 4,437E7 | Fkbp1a |
| P62852 | 40S ribosomal protein S25 OS=Mus musculus GN=Rps25 PE=1 SV=1 - [RS25_MOUSE] | 0,000E0 | 0,000E0 | 0,000E0 | 4,270E8 | 4,491E8 | 9,905E8 | Rps25 |
| P62743 | AP-2 complex subunit sigma OS=Mus musculus GN=Ap2s1 PE=1 SV=1 - [AP2S1_MOUSE] | 0,000E0 | 0,000E0 | 0,000E0 | 1,235E8 | 9,021E7 | 8,276E7 | Ap2s1 |
| Q6NZK5 | Protein hinderin OS=Mus musculus GN=Kiaa1328 PE=1 SV=2 - [K1328_MOUSE] | 0,000E0 | 0,000E0 | 0,000E0 | 5,606E6 | 4,169E6 | 6,911E6 | Kiaa1328 |
| Q8R4U7 | Leucine zipper protein 1 OS=Mus musculus GN=Luzp1 PE=1 SV=2 - [LUZP1_MOUSE] | 0,000E0 | 0,000E0 | 0,000E0 | 5,260E6 | 1,013E7 | 1,647E7 | Luzp1 |
| Q99K85 | Phosphoserine aminotransferase OS=Mus musculus GN=Psat1 PE=1 SV=1 - [SERC_MOUSE] | 0,000E0 | 0,000E0 | 0,000E0 | 7,764E6 | 2,652E7 | 4,732E7 | Psat1 |
| Q9ES28-4 | Isoform D of Rho guanine nucleotide exchange factor 7 OS=Mus musculus GN=Arhgef7 - [ARHG7_MOUSE] | 0,000E0 | 0,000E0 | 0,000E0 | 7,933E6 | 1,967E7 | 2,425E7 | Arhgef7 |
| Q9Z1Y4 | Thyroid receptor-interacting protein 6 OS=Mus musculus GN=Trip6 PE=1 SV=1 - [TRIP6_MOUSE] | 0,000E0 | 0,000E0 | 0,000E0 | 4,312E6 | 1,240E7 | 5,904E6 | Trip6 |
| Q9D0I8 | mRNA turnover protein 4 homolog OS=Mus musculus GN=Mrto4 PE=1 SV=1 - [MRT4_MOUSE] | 0,000E0 | 0,000E0 | 0,000E0 | 2,490E6 | 1,134E7 | 2,526E7 | Mrto4 |
| Q9D937 | Uncharacterized protein C11orf98 homolog OS=Mus musculus PE=1 SV=1 - [CK098_MOUSE] | 0,000E0 | 0,000E0 | 0,000E0 | 2,049E7 | 1,145E7 | 7,374E7 | CK098 |
| Q9CZY3 | Ubiquitin-conjugating enzyme E2 variant 1 OS=Mus musculus GN=Ube2v1 PE=1 SV=1 - [UB2V1_MOUSE] | 0,000E0 | 0,000E0 | 0,000E0 | 5,127E6 | 1,410E7 | 2,524E7 | Ube2v1 |
| Q99JB8 | Protein kinase C and casein kinase II substrate protein 3 OS=Mus musculus GN=Pacsin3 PE=1 SV=1 - [PACN3_MOUSE] | 0,000E0 | 0,000E0 | 0,000E0 | 1,013E7 | 2,183E7 | 1,543E7 | Pacsin3 |
| P43274 | Histone H1.4 OS=Mus musculus GN=Hist1h1e PE=1 SV=2 - [H14_MOUSE] | 0,000E0 | 0,000E0 | 6,982E5 | 5,100E7 | 1,359E8 | 3,077E8 | Hist1h1e |
| Q99J93 | Interferon-induced transmembrane protein 2 OS=Mus musculus GN=Ifitm2 PE=1 SV=1 - [IFM2_MOUSE] | 0,000E0 | 0,000E0 | 0,000E0 | 3,591E6 | 1,890E8 | 2,051E8 | Ifitm2 |
| Q62188 | Dihydropyrimidinase-related protein 3 OS=Mus musculus GN=Dpysl3 PE=1 SV=1 - [DPYL3_MOUSE] | 0,000E0 | 0,000E0 | 0,000E0 | 4,926E6 | 1,162E7 | 2,964E7 | Dpysl3 |
| Q04750 | DNA topoisomerase 1 OS=Mus musculus GN=Top1 PE=1 SV=2 - [TOP1_MOUSE] | 0,000E0 | 0,000E0 | 0,000E0 | 1,463E7 | 1,640E7 | 8,076E7 | Top1 |
| Q7TQH0-2 | Isoform 2 of Ataxin-2-like protein OS=Mus musculus GN=Atxn2l - [ATX2L_MOUSE] | 0,000E0 | 0,000E0 | 2,775E5 | 2,831E7 | 4,559E7 | 1,007E8 | Atxn2l |
| Q99KY4 | Cyclin-G-associated kinase OS=Mus musculus GN=Gak PE=1 SV=2 - [GAK_MOUSE] | 0,000E0 | 0,000E0 | 0,000E0 | 3,046E7 | 2,869E7 | 4,032E7 | Gak |
| Q62448-2 | Isoform 2 of Eukaryotic translation initiation factor 4 gamma 2 OS=Mus musculus GN=Eif4g2 - [IF4G2_MOUSE] | 0,000E0 | 0,000E0 | 0,000E0 | 1,365E7 | 2,158E7 | 2,428E7 | Eif4g2 |
| Q60634-3 | Isoform 3 of Flotillin-2 OS=Mus musculus GN=Flot2 - [FLOT2_MOUSE] | 0,000E0 | 0,000E0 | 0,000E0 | 7,771E5 | 6,085E6 | 3,056E7 | Flot2 |
| P70441 | Na(+)/H(+) exchange regulatory cofactor NHE-RF1 OS=Mus musculus GN=Slc9a3r1 PE=1 SV=3 - [NHRF1_MOUSE] | 0,000E0 | 0,000E0 | 0,000E0 | 3,953E6 | 1,152E7 | 2,282E7 | Slc9a3r1 |
| P47955 | 60S acidic ribosomal protein P1 OS=Mus musculus GN=Rplp1 PE=1 SV=1 - [RLA1_MOUSE] | 0,000E0 | 0,000E0 | 0,000E0 | 8,722E7 | 6,223E7 | 2,263E8 | Rplp1 |
| P97479-2 | Isoform 2 of Unconventional myosin-VIIa OS=Mus musculus GN=Myo7a - [MYO7A_MOUSE] | 2,735E6 | 0,000E0 | 0,000E0 | 1,253E7 | 2,377E7 | 4,692E7 | Myo7a |
| Q9WV55 | Vesicle-associated membrane protein-associated protein A OS=Mus musculus GN=Vapa PE=1 SV=2 - [VAPA_MOUSE] | 0,000E0 | 0,000E0 | 0,000E0 | 1,038E7 | 1,461E7 | 2,391E7 | Vapa |
| Q64727 | Vinculin OS=Mus musculus GN=Vcl PE=1 SV=4 - [VINC_MOUSE] | 0,000E0 | 0,000E0 | 6,575E6 | 6,419E6 | 9,594E6 | 1,489E7 | Vcl |
| Q8C310 | Roundabout homolog 4 OS=Mus musculus GN=Robo4 PE=1 SV=2 - [ROBO4_MOUSE] | 0,000E0 | 0,000E0 | 0,000E0 | 1,506E7 | 3,265E7 | 4,558E7 | Robo4 |
| Q60817 | Nascent polypeptide-associated complex subunit alpha OS=Mus musculus GN=Naca PE=1 SV=1 - [NACA_MOUSE] | 0,000E0 | 0,000E0 | 0,000E0 | 1,286E8 | 1,393E8 | 3,863E8 | Naca |

| | | | | | | | | |
|---|---|---|---|---|---|---|---|---|
| Q9WVR4 | Fragile X mental retardation syndrome-related protein 2 OS=Mus musculus GN=Fxr2 PE=1 SV=1 - [FXR2_MOUSE] | 0,000E0 | 0,000E0 | 0,000E0 | 6,711E7 | 1,504E8 | 3,104E8 | Fxr2 |
| P45377 | Aldose reductase-related protein 2 OS=Mus musculus GN=Akr1b8 PE=1 SV=2 - [ALD2_MOUSE] | 0,000E0 | 0,000E0 | 0,000E0 | 6,907E6 | 8,907E6 | 1,044E7 | Akr1b8 |
| Q9CQW1 | Synaptobrevin homolog YKT6 OS=Mus musculus GN=Ykt6 PE=1 SV=1 - [YKT6_MOUSE] | 0,000E0 | 0,000E0 | 0,000E0 | 2,513E6 | 6,558E6 | 1,345E7 | Ykt6 |
| Q8C1Q6 | Small integral membrane protein 4 OS=Mus musculus GN=Smim4 PE=1 SV=2 - [SMIM4_MOUSE] | 0,000E0 | 0,000E0 | 0,000E0 | 5,398E6 | 4,131E6 | 2,890E6 | Smim4 |
| Q8BVU0 | Leucine-rich repeat and calponin homology domain-containing protein 3 OS=Mus musculus GN=Lrch3 PE=1 SV=3 - [LRCH3_MOUSE] | 0,000E0 | 0,000E0 | 0,000E0 | 2,790E7 | 6,295E7 | 5,882E7 | Lrch3 |
| P11103 | Poly [ADP-ribose] polymerase 1 OS=Mus musculus GN=Parp1 PE=1 SV=3 - [PARP1_MOUSE] | 0,000E0 | 0,000E0 | 0,000E0 | 3,326E7 | 5,602E7 | 1,078E8 | Parp1 |
| A2A7S8-2 | Isoform 2 of Uncharacterized protein KIAA1522 OS=Mus musculus GN=Kiaa1522 - [K1522_MOUSE] | 0,000E0 | 0,000E0 | 0,000E0 | 4,083E6 | 8,124E6 | 1,459E7 | Kiaa1522 |
| O08967 | Cytohesin-3 OS=Mus musculus GN=Cyth3 PE=1 SV=1 - [CYH3_MOUSE] | 0,000E0 | 0,000E0 | 0,000E0 | 9,632E6 | 1,203E7 | 1,007E7 | Cyth3 |
| Q0GNC1 | Inverted formin-2 OS=Mus musculus GN=Inf2 PE=1 SV=1 - [INF2_MOUSE] | 0,000E0 | 0,000E0 | 0,000E0 | 1,073E7 | 2,623E7 | 3,561E7 | Inf2 |
| P55258 | Ras-related protein Rab-8A OS=Mus musculus GN=Rab8a PE=1 SV=2 - [RAB8A_MOUSE] | 0,000E0 | 0,000E0 | 3,456E5 | 8,538E7 | 1,285E8 | 1,853E8 | Rab8a |
| P61028 | Ras-related protein Rab-8B OS=Mus musculus GN=Rab8b PE=1 SV=1 - [RAB8B_MOUSE] | 0,000E0 | 0,000E0 | 3,456E5 | 8,538E7 | 1,285E8 | 1,853E8 | Rab8b |
| P10833 | Ras-related protein R-Ras OS=Mus musculus GN=Rras PE=1 SV=1 - [RRAS_MOUSE] | 0,000E0 | 0,000E0 | 0,000E0 | 5,125E7 | 1,417E8 | 2,087E8 | Rras |
| Q99KP6 | Pre-mRNA-processing factor 19 OS=Mus musculus GN=Prpf19 PE=1 SV=1 - [PRP19_MOUSE] | 0,000E0 | 0,000E0 | 0,000E0 | 1,126E7 | 1,887E7 | 4,198E7 | Prpf19 |
| Q8C310-2 | Isoform 2 of Roundabout homolog 4 OS=Mus musculus GN=Robo4 - [ROBO4_MOUSE] | 0,000E0 | 0,000E0 | 0,000E0 | 1,233E7 | 3,040E7 | 3,519E7 | Robo4 |
| Q99JF8 | PC4 and SFRS1-interacting protein OS=Mus musculus GN=Psip1 PE=1 SV=1 - [PSIP1_MOUSE] | 0,000E0 | 0,000E0 | 0,000E0 | 4,153E6 | 1,615E7 | 3,209E7 | Psip1 |
| O70318 | Band 4.1-like protein 2 OS=Mus musculus GN=Epb41l2 PE=1 SV=2 - [E41L2_MOUSE] | 0,000E0 | 0,000E0 | 0,000E0 | 1,023E7 | 3,059E7 | 3,963E7 | Epb41l2 |
| Q9Z1D1 | Eukaryotic translation initiation factor 3 subunit G OS=Mus musculus GN=Eif3g PE=1 SV=2 - [EIF3G_MOUSE] | 0,000E0 | 0,000E0 | 0,000E0 | 1,971E7 | 3,817E7 | 6,295E7 | Eif3g |
| Q9JIK5 | Nucleolar RNA helicase 2 OS=Mus musculus GN=Ddx21 PE=1 SV=3 - [DDX21_MOUSE] | 0,000E0 | 0,000E0 | 0,000E0 | 3,983E6 | 8,756E6 | 3,872E7 | Ddx21 |
| Q8R0J7 | Vacuolar protein sorting-associated protein 37B OS=Mus musculus GN=Vps37b PE=1 SV=1 - [VP37B_MOUSE] | 0,000E0 | 0,000E0 | 0,000E0 | 6,840E6 | 8,200E6 | 2,561E7 | Vps37b |
| Q62425 | Cytochrome c oxidase subunit NDUFA4 OS=Mus musculus GN=Ndufa4 PE=1 SV=2 - [NDUA4_MOUSE] | 0,000E0 | 0,000E0 | 0,000E0 | 1,639E7 | 2,127E7 | 4,603E7 | Ndufa4 |
| Q5DTX6 | Junctional protein associated with coronary artery disease OS=Mus musculus GN=Jcad PE=1 SV=2 - [JCAD_MOUSE] | 0,000E0 | 0,000E0 | 0,000E0 | 2,165E6 | 1,951E7 | 4,623E7 | Jcad |
| Q91ZU6-3 | Isoform 3 of Dystonin OS=Mus musculus GN=Dst - [DYST_MOUSE] | 0,000E0 | 0,000E0 | 0,000E0 | 1,372E7 | 2,283E7 | 1,736E7 | Dst |
| Q9D8Y0 | EF-hand domain-containing protein D2 OS=Mus musculus GN=Efhd2 PE=1 SV=1 - [EFHD2_MOUSE] | 0,000E0 | 0,000E0 | 0,000E0 | 1,699E6 | 2,151E7 | 4,016E7 | Efhd2 |
| Q80X50-2 | Isoform 2 of Ubiquitin-associated protein 2-like OS=Mus musculus GN=Ubap2l - [UBP2L_MOUSE] | 0,000E0 | 0,000E0 | 0,000E0 | 2,248E7 | 2,929E7 | 5,600E7 | Ubap2l |
| D0QMC3 | Myeloid cell nuclear differentiation antigen-like protein OS=Mus musculus GN=Mndal PE=1 SV=1 - [MNDAL_MOUSE] | 0,000E0 | 0,000E0 | 1,867E5 | 9,720E6 | 1,238E7 | 3,916E7 | Mndal |
| G3X9J0 | Signal-induced proliferation-associated 1-like protein 3 OS=Mus musculus GN=Sipa1l3 PE=1 SV=1 - [SI1L3_MOUSE] | 0,000E0 | 0,000E0 | 0,000E0 | 1,902E7 | 1,497E7 | 5,078E7 | Sipa1l3 |
| P08207 | Protein S100-A10 OS=Mus musculus GN=S100a10 PE=1 SV=2 - [S10AA_MOUSE] | 0,000E0 | 0,000E0 | 0,000E0 | 9,947E6 | 7,128E6 | 1,307E7 | S100a10 |
| Q9CR16 | Peptidyl-prolyl cis-trans isomerase D OS=Mus musculus GN=Ppid PE=1 SV=3 - [PPID_MOUSE] | 0,000E0 | 0,000E0 | 0,000E0 | 2,557E6 | 2,588E6 | 5,581E6 | Ppid |
| P35550 | rRNA 2'-O-methyltransferase fibrillarin OS=Mus musculus GN=Fbl PE=1 SV=2 - [FBRL_MOUSE] | 0,000E0 | 0,000E0 | 0,000E0 | 9,037E6 | 2,221E7 | 5,299E7 | Fbl |
| P56480 | ATP synthase subunit beta, mitochondrial OS=Mus musculus GN=Atp5b PE=1 SV=2 - [ATPB_MOUSE] | 0,000E0 | 0,000E0 | 0,000E0 | 2,061E6 | 6,927E6 | 4,275E6 | Atp5b |
| Q8R1F1 | Niban-like protein 1 OS=Mus musculus GN=Fam129b PE=1 SV=2 - [NIBL1_MOUSE] | 0,000E0 | 0,000E0 | 0,000E0 | 7,346E6 | 1,495E7 | 2,254E7 | Fam129b |
| P10630 | Eukaryotic initiation factor 4A-II OS=Mus musculus GN=Eif4a2 PE=1 SV=2 - [IF4A2_MOUSE] | 0,000E0 | 0,000E0 | 3,565E5 | 3,511E7 | 9,717E7 | 7,502E7 | Eif4a2 |
| P97863 | Nuclear factor 1 B-type OS=Mus musculus GN=Nfib PE=1 SV=2 - [NFIB_MOUSE] | 0,000E0 | 0,000E0 | 0,000E0 | 1,739E7 | 4,045E7 | 4,062E7 | Nfib |
| O08585 | Clathrin light chain A OS=Mus musculus GN=Clta PE=1 SV=2 - [CLCA_MOUSE] | 0,000E0 | 0,000E0 | 0,000E0 | 1,618E8 | 2,524E8 | 1,907E8 | Clta |
| P61089 | Ubiquitin-conjugating enzyme E2 N OS=Mus musculus GN=Ube2n PE=1 SV=1 - [UBE2N_MOUSE] | 0,000E0 | 0,000E0 | 0,000E0 | 4,490E6 | 8,080E6 | 1,842E7 | Ube2n |
| A2ALU4 | Protein Shroom2 OS=Mus musculus GN=Shroom2 PE=1 SV=1 - [SHRM2_MOUSE] | 0,000E0 | 0,000E0 | 0,000E0 | 3,672E6 | 1,142E7 | 1,142E7 | Shroom2 |
| P52293 | Importin subunit alpha-1 OS=Mus musculus GN=Kpna2 PE=1 SV=2 - [IMA1_MOUSE] | 0,000E0 | 0,000E0 | 0,000E0 | 6,294E6 | 1,788E7 | 5,973E7 | Kpna2 |
| Q9DB77 | Cytochrome b-c1 complex subunit 2, mitochondrial OS=Mus musculus GN=Uqcrc2 PE=1 SV=1 - [QCR2_MOUSE] | 0,000E0 | 0,000E0 | 0,000E0 | 1,047E7 | 1,787E7 | 2,301E7 | Uqcrc2 |
| Q7TT50 | Serine/threonine-protein kinase MRCK beta OS=Mus musculus GN=Cdc42bpb PE=1 SV=2 - [MRCKB_MOUSE] | 0,000E0 | 0,000E0 | 0,000E0 | 2,322E7 | 3,905E7 | 5,367E7 | Cdc42bpb |
| P14685 | 26S proteasome non-ATPase regulatory subunit 3 OS=Mus musculus GN=Psmd3 PE=1 SV=3 - [PSMD3_MOUSE] | 0,000E0 | 0,000E0 | 0,000E0 | 3,528E6 | 6,222E6 | 2,676E6 | Psmd3 |
| P16460 | Argininosuccinate synthase OS=Mus musculus GN=Ass1 PE=1 SV=1 - [ASSY_MOUSE] | 0,000E0 | 0,000E0 | 0,000E0 | 1,045E6 | 7,049E6 | 7,550E6 | Ass1 |
| P56389 | Cytidine deaminase OS=Mus musculus GN=Cda PE=1 SV=2 - [CDD_MOUSE] | 0,000E0 | 0,000E0 | 0,000E0 | 1,768E6 | 3,275E6 | 5,216E6 | Cda |
| P25976-2 | Isoform UBF2 of Nucleolar transcription factor 1 OS=Mus musculus GN=Ubtf - [UBF1_MOUSE] | 0,000E0 | 0,000E0 | 0,000E0 | 2,807E7 | 4,107E7 | 7,156E7 | Ubtf |
| P61226 | Ras-related protein Rap-2b OS=Mus musculus GN=Rap2b PE=1 SV=1 - [RAP2B_MOUSE] | 0,000E0 | 0,000E0 | 0,000E0 | 6,647E6 | 1,636E7 | 1,418E7 | Rap2b |
| Q9CQF3 | Cleavage and polyadenylation specificity factor subunit 5 OS=Mus musculus GN=Nudt21 PE=1 SV=1 - [CPSF5_MOUSE] | 0,000E0 | 0,000E0 | 0,000E0 | 8,429E6 | 4,930E6 | 8,752E6 | Nudt21 |
| Q8CI32 | BAG family molecular chaperone regulator 5 OS=Mus musculus GN=Bag5 PE=1 SV=1 - [BAG5_MOUSE] | 0,000E0 | 0,000E0 | 0,000E0 | 1,632E6 | 5,031E6 | 7,844E6 | Bag5 |
| Q8VI36 | Paxillin OS=Mus musculus GN=Pxn PE=1 SV=1 - [PAXI_MOUSE] | 0,000E0 | 0,000E0 | 0,000E0 | 8,372E6 | 1,208E7 | 2,223E7 | Pxn |
| Q9JL15 | Galectin-8 OS=Mus musculus GN=Lgals8 PE=1 SV=1 - [LEG8_MOUSE] | 0,000E0 | 0,000E0 | 0,000E0 | 8,521E6 | 6,596E6 | 1,544E7 | Lgals8 |
| P42567 | Epidermal growth factor receptor substrate 15 OS=Mus musculus GN=Eps15 PE=1 SV=1 - [EPS15_MOUSE] | 0,000E0 | 0,000E0 | 0,000E0 | 5,288E6 | 1,221E7 | 2,108E6 | Eps15 |
| Q9ES28-3 | Isoform C of Rho guanine nucleotide exchange factor 7 OS=Mus musculus GN=Arhgef7 - [ARHG7_MOUSE] | 0,000E0 | 0,000E0 | 0,000E0 | 7,933E6 | 1,967E7 | 2,425E7 | Arhgef7 |
| Q03963 | Interferon-induced, double-stranded RNA-activated protein kinase OS=Mus musculus GN=Eif2ak2 PE=1 SV=2 - [E2AK2_MOUSE] | 0,000E0 | 0,000E0 | 0,000E0 | 1,825E7 | 1,189E7 | 4,821E7 | Eif2ak2 |
| Q3TJZ6 | Protein FAM98A OS=Mus musculus GN=Fam98a PE=1 SV=1 - [FA98A_MOUSE] | 0,000E0 | 0,000E0 | 0,000E0 | 1,311E7 | 3,298E7 | 5,471E7 | Fam98a |
| Q9CPQ8 | ATP synthase subunit g, mitochondrial OS=Mus musculus GN=Atp5l PE=1 SV=1 - [ATP5L_MOUSE] | 0,000E0 | 0,000E0 | 0,000E0 | 7,494E6 | 1,746E6 | 2,020E6 | Atp5l |
| Q9DBR1 | 5'-3' exoribonuclease 2 OS=Mus musculus GN=Xrn2 PE=1 SV=1 - [XRN2_MOUSE] | 0,000E0 | 0,000E0 | 0,000E0 | 1,271E7 | 2,706E7 | 1,111E7 | Xrn2 |
| Q8BMJ3 | Eukaryotic translation initiation factor 1A, X-chromosomal OS=Mus musculus GN=Eif1ax PE=2 SV=3 - [IF1AX_MOUSE] | 0,000E0 | 0,000E0 | 0,000E0 | 5,414E6 | 2,123E7 | 5,076E7 | Eif1ax |
| Q5F2E8 | Serine/threonine-protein kinase TAO1 OS=Mus musculus GN=Taok1 PE=1 SV=1 - [TAOK1_MOUSE] | 0,000E0 | 0,000E0 | 8,146E5 | 7,558E6 | 3,267E7 | 9,395E7 | Taok1 |
| Q921F4 | Heterogeneous nuclear ribonucleoprotein L-like OS=Mus musculus GN=Hnrnpll PE=1 SV=3 - [HNRLL_MOUSE] | 0,000E0 | 0,000E0 | 0,000E0 | 7,243E6 | 1,012E7 | 1,466E7 | Hnrnpll |
| Q9DB34 | Charged multivesicular body protein 2a OS=Mus musculus GN=Chmp2a PE=1 SV=1 - [CHM2A_MOUSE] | 0,000E0 | 0,000E0 | 0,000E0 | 1,985E7 | 4,101E7 | 5,862E7 | Chmp2a |
| P49718 | DNA replication licensing factor MCM5 OS=Mus musculus GN=Mcm5 PE=1 SV=1 - [MCM5_MOUSE] | 0,000E0 | 0,000E0 | 0,000E0 | 1,051E7 | 2,035E7 | 2,451E7 | Mcm5 |
| P25206 | DNA replication licensing factor MCM3 OS=Mus musculus GN=Mcm3 PE=1 SV=2 - [MCM3_MOUSE] | 0,000E0 | 0,000E0 | 0,000E0 | 6,610E6 | 1,312E7 | 1,015E7 | Mcm3 |
| O55135 | Eukaryotic translation initiation factor 6 OS=Mus musculus GN=Eif6 PE=1 SV=2 - [IF6_MOUSE] | 0,000E0 | 0,000E0 | 0,000E0 | 1,485E7 | 2,684E7 | 1,696E7 | Eif6 |
| P28660-2 | Isoform 2 of Nck-associated protein 1 OS=Mus musculus GN=Nckap1 - [NCKP1_MOUSE] | 0,000E0 | 0,000E0 | 6,061E5 | 2,268E7 | 5,553E7 | 5,784E7 | Nckap1 |
| Q8VIJ6 | Splicing factor, proline- and glutamine-rich OS=Mus musculus GN=Sfpq PE=1 SV=1 - [SFPQ_MOUSE] | 0,000E0 | 0,000E0 | 0,000E0 | 3,941E7 | 7,024E7 | 1,793E8 | Sfpq |
| Q99KJ8 | Dynactin subunit 2 OS=Mus musculus GN=Dctn2 PE=1 SV=3 - [DCTN2_MOUSE] | 0,000E0 | 0,000E0 | 0,000E0 | 1,506E6 | 5,075E6 | 3,515E6 | Dctn2 |
| Q8CBY8-2 | Isoform 2 of Dynactin subunit 4 OS=Mus musculus GN=Dctn4 - [DCTN4_MOUSE] | 0,000E0 | 0,000E0 | 0,000E0 | 3,898E6 | 4,745E6 | 1,342E7 | Dctn4 |
| Q64012-2 | Isoform 1 of RNA-binding protein Raly OS=Mus musculus GN=Raly - [RALY_MOUSE] | 0,000E0 | 0,000E0 | 0,000E0 | 1,885E6 | 1,563E7 | 2,233E7 | Raly |
| Q91VR8 | Protein BRICK1 OS=Mus musculus GN=Brk1 PE=1 SV=1 - [BRK1_MOUSE] | 0,000E0 | 0,000E0 | 0,000E0 | 5,373E6 | 1,025E7 | 1,071E7 | Brk1 |
| Q9Z204-2 | Isoform C1 of Heterogeneous nuclear ribonucleoproteins C1/C2 OS=Mus musculus GN=Hnrnpc - [HNRPC_MOUSE] | 0,000E0 | 0,000E0 | 0,000E0 | 5,070E6 | 9,824E5 | 1,648E7 | Hnrnpc |
| Q8VEE4 | Replication protein A 70 kDa DNA-binding subunit OS=Mus musculus GN=Rpa1 PE=1 SV=1 - [RFA1_MOUSE] | 0,000E0 | 0,000E0 | 0,000E0 | 4,744E6 | 8,574E6 | 3,929E6 | Rpa1 |
| Q8R080 | G2 and S phase-expressed protein 1 OS=Mus musculus GN=Gtse1 PE=1 SV=2 - [GTSE1_MOUSE] | 0,000E0 | 0,000E0 | 0,000E0 | 8,248E6 | 1,316E7 | 2,286E7 | Gtse1 |
| Q9CX86 | Heterogeneous nuclear ribonucleoprotein A0 OS=Mus musculus GN=Hnrnpa0 PE=1 SV=1 - [ROA0_MOUSE] | 0,000E0 | 0,000E0 | 0,000E0 | 1,633E7 | 3,586E7 | 5,517E7 | Hnrnpa0 |
| P58137 | Acyl-coenzyme A thioesterase 8 OS=Mus musculus GN=Acot8 PE=1 SV=1 - [ACOT8_MOUSE] | 0,000E0 | 0,000E0 | 0,000E0 | 4,770E6 | 3,566E6 | 1,936E7 | Acot8 |
| P27601 | Guanine nucleotide-binding protein subunit alpha-13 OS=Mus musculus GN=Gna13 PE=1 SV=1 - [GNA13_MOUSE] | 0,000E0 | 0,000E0 | 0,000E0 | 2,341E7 | 1,140E8 | 4,388E6 | Gna13 |
| Q80XC3 | USP6 N-terminal-like protein OS=Mus musculus GN=Usp6nl PE=1 SV=2 - [US6NL_MOUSE] | 0,000E0 | 0,000E0 | 0,000E0 | 4,142E6 | 7,592E6 | 3,751E7 | Usp6nl |
| Q8VDJ3 | Vigilin OS=Mus musculus GN=Hdlbp PE=1 SV=1 - [VIGLN_MOUSE] | 0,000E0 | 0,000E0 | 0,000E0 | 5,970E6 | 7,004E6 | 2,054E7 | Hdlbp |
| Q8CGU1 | Calcium-binding and coiled-coil domain-containing protein 1 OS=Mus musculus GN=Calcoco1 PE=1 SV=2 - [CACO1_MOUSE] | 0,000E0 | 0,000E0 | 0,000E0 | 1,143E7 | 1,697E7 | 2,881E7 | Calcoco1 |
| Q66GT5 | Phosphatidylglycerophosphatase and protein-tyrosine phosphatase 1 OS=Mus musculus GN=Ptpmt1 PE=1 SV=1 - [PTPM1_MOUSE] | 0,000E0 | 0,000E0 | 0,000E0 | 3,818E6 | 7,785E6 | 9,755E6 | Ptpmt1 |
| G5E829 | Plasma membrane calcium-transporting ATPase 1 OS=Mus musculus GN=Atp2b1 PE=1 SV=1 - [AT2B1_MOUSE] | 0,000E0 | 0,000E0 | 0,000E0 | 8,901E6 | 2,316E7 | 2,315E7 | Atp2b1 |
| Q60872 | Eukaryotic translation initiation factor 1A OS=Mus musculus GN=Eif1a PE=2 SV=3 - [IF1A_MOUSE] | 0,000E0 | 0,000E0 | 0,000E0 | 8,348E6 | 2,123E7 | 5,076E7 | Eif1a |
| Q9DB20 | ATP synthase subunit O, mitochondrial OS=Mus musculus GN=Atp5o PE=1 SV=1 - [ATPO_MOUSE] | 0,000E0 | 0,000E0 | 0,000E0 | 1,631E6 | 4,673E6 | 4,143E6 | Atp5o |
| Q8K124 | Pleckstrin homology domain-containing family O member 2 OS=Mus musculus GN=Plekho2 PE=1 SV=1 - [PKHO2_MOUSE] | 0,000E0 | 0,000E0 | 0,000E0 | 2,216E6 | 5,138E6 | 6,130E6 | Plekho2 |
| Q03265 | ATP synthase subunit alpha, mitochondrial OS=Mus musculus GN=Atp5a1 PE=1 SV=1 - [ATPA_MOUSE] | 0,000E0 | 0,000E0 | 1,105E6 | 8,822E6 | 6,280E6 | 1,502E7 | Atp5a1 |
| Q60766-2 | Isoform 2 of Immunity-related GTPase family M protein 1 OS=Mus musculus GN=Irgm1 - [IRGM1_MOUSE] | 0,000E0 | 0,000E0 | 0,000E0 | 9,248E6 | 2,004E7 | 3,084E7 | Irgm1 |
| P56135 | ATP synthase subunit f, mitochondrial OS=Mus musculus GN=Atp5j2 PE=1 SV=3 - [ATPK_MOUSE] | 0,000E0 | 0,000E0 | 0,000E0 | 5,203E6 | 2,964E7 | 4,474E7 | Atp5j2 |
| Q9WVA3 | Mitotic checkpoint protein BUB3 OS=Mus musculus GN=Bub3 PE=1 SV=2 - [BUB3_MOUSE] | 0,000E0 | 0,000E0 | 0,000E0 | 6,979E6 | 1,265E7 | 2,698E7 | Bub3 |
| P54116 | Erythrocyte band 7 integral membrane protein OS=Mus musculus GN=Stom PE=1 SV=3 - [STOM_MOUSE] | 0,000E0 | 0,000E0 | 3,789E5 | 2,246E6 | 5,342E6 | 1,535E7 | Stom |
| P10107 | Annexin A1 OS=Mus musculus GN=Anxa1 PE=1 SV=2 - [ANXA1_MOUSE] | 0,000E0 | 0,000E0 | 0,000E0 | 1,019E7 | 1,799E7 | 1,162E7 | Anxa1 |
| P25976 | Nucleolar transcription factor 1 OS=Mus musculus GN=Ubtf PE=1 SV=1 - [UBF1_MOUSE] | 0,000E0 | 0,000E0 | 0,000E0 | 2,807E7 | 4,107E7 | 7,156E7 | Ubtf |
| Q8BYK6-3 | Isoform 3 of YTH domain-containing family protein 3 OS=Mus musculus GN=Ythdf3 - [YTHD3_MOUSE] | 0,000E0 | 0,000E0 | 5,369E6 | 1,804E7 | 1,525E7 | 1,924E7 | Ythdf3 |
| Q9CRB9 | MICOS complex subunit Mic19 OS=Mus musculus GN=Chchd3 PE=1 SV=1 - [MIC19_MOUSE] | 0,000E0 | 0,000E0 | 5,868E5 | 8,755E6 | 1,533E7 | 2,979E7 | Chchd3 |
| Q9Z0R6 | Intersectin-2 OS=Mus musculus GN=Itsn2 PE=1 SV=2 - [ITSN2_MOUSE] | 0,000E0 | 0,000E0 | 0,000E0 | 1,868E7 | 2,768E7 | 5,085E7 | Itsn2 |
| Q9D1C2 | Protein chibby homolog 1 OS=Mus musculus GN=Cby1 PE=1 SV=1 - [CBY1_MOUSE] | 0,000E0 | 0,000E0 | 0,000E0 | 1,215E6 | 3,119E6 | 4,995E6 | Cby1 |
| Q91W59-2 | Isoform 2 of RNA-binding motif, single-stranded-interacting protein 1 OS=Mus musculus GN=Rbms1 - [RBMS1_MOUSE] | 0,000E0 | 0,000E0 | 0,000E0 | 2,295E6 | 1,371E7 | 5,239E7 | Rbms1 |

| | | | | | | | | |
|---|---|---|---|---|---|---|---|---|
| Q8BPB0 | MOB kinase activator 1B OS=Mus musculus GN=Mob1b PE=1 SV=3 - [MOB1B_MOUSE] | 0,000E0 | 0,000E0 | 0,000E0 | 1,913E7 | 4,068E7 | 3,950E7 | Mob1b |
| P62827 | GTP-binding nuclear protein Ran OS=Mus musculus GN=Ran PE=1 SV=3 - [RAN_MOUSE] | 0,000E0 | 0,000E0 | 0,000E0 | 3,064E7 | 1,239E9 | 1,129E8 | Ran |
| P26645 | Myristoylated alanine-rich C-kinase substrate OS=Mus musculus GN=Marcks PE=1 SV=2 - [MARCS_MOUSE] | 0,000E0 | 0,000E0 | 0,000E0 | 4,771E6 | 6,669E6 | 9,817E6 | Marcks |
| O08579 | Emerin OS=Mus musculus GN=Emd PE=1 SV=1 - [EMD_MOUSE] | 0,000E0 | 0,000E0 | 0,000E0 | 3,630E6 | 9,661E6 | 1,267E7 | Emd |
| Q8VDZ4 | Palmitoyltransferase ZDHHC5 OS=Mus musculus GN=Zdhhc5 PE=1 SV=1 - [ZDHC5_MOUSE] | 0,000E0 | 0,000E0 | 0,000E0 | 3,212E6 | 6,478E6 | 1,889E7 | Zdhhc5 |
| Q8C2Q3 | RNA-binding protein 14 OS=Mus musculus GN=Rbm14 PE=1 SV=1 - [RBM14_MOUSE] | 0,000E0 | 0,000E0 | 0,000E0 | 1,986E7 | 3,539E7 | 5,818E7 | Rbm14 |
| Q4VAA2-2 | Isoform 2 of Protein CDV3 OS=Mus musculus GN=Cdv3 - [CDV3_MOUSE] | 0,000E0 | 0,000E0 | 0,000E0 | 1,485E6 | 1,685E6 | 6,074E6 | Cdv3 |
| P97477 | Aurora kinase A OS=Mus musculus GN=Aurka PE=1 SV=1 - [AURKA_MOUSE] | 0,000E0 | 0,000E0 | 0,000E0 | 1,045E6 | 4,874E6 | 4,838E6 | Aurka |
| Q9CQM9 | Glutaredoxin-3 OS=Mus musculus GN=Glrx3 PE=1 SV=1 - [GLRX3_MOUSE] | 0,000E0 | 0,000E0 | 0,000E0 | 9,684E6 | 2,899E7 | 2,885E7 | Glrx3 |
| P35831 | Tyrosine-protein phosphatase non-receptor type 12 OS=Mus musculus GN=Ptpn12 PE=1 SV=3 - [PTN12_MOUSE] | 0,000E0 | 0,000E0 | 0,000E0 | 3,444E6 | 1,364E7 | 1,241E7 | Ptpn12 |
| Q62418-3 | Isoform 3 of Drebrin-like protein OS=Mus musculus GN=Dbnl - [DBNL_MOUSE] | 0,000E0 | 0,000E0 | 0,000E0 | 8,414E6 | 1,922E7 | 3,395E7 | Dbnl |
| O54916 | RalBP1-associated Eps domain-containing protein 1 OS=Mus musculus GN=Reps1 PE=1 SV=2 - [REPS1_MOUSE] | 0,000E0 | 0,000E0 | 0,000E0 | 1,497E7 | 4,671E7 | 3,297E7 | Reps1 |
| Q91VI7 | Ribonuclease inhibitor OS=Mus musculus GN=Rnh1 PE=1 SV=1 - [RINI_MOUSE] | 0,000E0 | 0,000E0 | 0,000E0 | 1,471E7 | 3,332E7 | 3,525E7 | Rnh1 |
| P47915 | 60S ribosomal protein L29 OS=Mus musculus GN=Rpl29 PE=2 SV=2 - [RL29_MOUSE] | 5,786E6 | 0,000E0 | 0,000E0 | 1,387E8 | 2,903E8 | 3,218E8 | Rpl29 |
| P32067 | Lupus La protein homolog OS=Mus musculus GN=Ssb PE=1 SV=1 - [LA_MOUSE] | 0,000E0 | 0,000E0 | 0,000E0 | 7,627E6 | 1,431E7 | 3,148E7 | Ssb |
| Q9QWT9 | Kinesin-like protein KIFC1 OS=Mus musculus GN=Kifc1 PE=1 SV=2 - [KIFC1_MOUSE] | 0,000E0 | 0,000E0 | 0,000E0 | 6,150E6 | 1,388E7 | 2,053E7 | Kifc1 |
| Q02053 | Ubiquitin-like modifier-activating enzyme 1 OS=Mus musculus GN=Uba1 PE=1 SV=1 - [UBA1_MOUSE] | 0,000E0 | 0,000E0 | 0,000E0 | 1,124E7 | 2,029E7 | 2,900E7 | Uba1 |
| Q8BX02 | KN motif and ankyrin repeat domain-containing protein 2 OS=Mus musculus GN=Kank2 PE=1 SV=1 - [KANK2_MOUSE] | 0,000E0 | 0,000E0 | 0,000E0 | 3,546E6 | 1,385E7 | 2,069E7 | Kank2 |
| E9Q6B2 | Coiled-coil domain-containing protein 85C OS=Mus musculus GN=Ccdc85c PE=1 SV=1 - [CC85C_MOUSE] | 0,000E0 | 0,000E0 | 0,000E0 | 1,008E7 | 1,893E7 | 2,081E7 | Ccdc85c |
| Q69Z38 | Pseudopodium-enriched atypical kinase 1 OS=Mus musculus GN=Peak1 PE=1 SV=4 - [PEAK1_MOUSE] | 0,000E0 | 0,000E0 | 0,000E0 | 8,664E6 | 2,644E7 | 3,621E7 | Peak1 |
| P01942 | Hemoglobin subunit alpha OS=Mus musculus GN=Hba PE=1 SV=2 - [HBA_MOUSE] | 8,540E7 | 0,000E0 | 0,000E0 | 3,162E6 | 1,324E6 | 1,836E6 | Hba |
| Q99KD5 | Protein unc-45 homolog A OS=Mus musculus GN=Unc45a PE=1 SV=2 - [UN45A_MOUSE] | 0,000E0 | 0,000E0 | 0,000E0 | 1,333E7 | 3,343E7 | 4,808E7 | Unc45a |
| O35326 | Serine/arginine-rich splicing factor 5 OS=Mus musculus GN=Srsf5 PE=1 SV=2 - [SRSF5_MOUSE] | 0,000E0 | 0,000E0 | 0,000E0 | 5,935E7 | 1,001E8 | 1,434E8 | Srsf5 |
| Q8R081 | Heterogeneous nuclear ribonucleoprotein L OS=Mus musculus GN=Hnrnpl PE=1 SV=2 - [HNRPL_MOUSE] | 0,000E0 | 0,000E0 | 0,000E0 | 5,301E6 | 6,586E6 | 3,423E7 | Hnrnpl |
| Q8K4G5-2 | Isoform 2 of Actin-binding LIM protein 1 OS=Mus musculus GN=Ablim1 - [ABLM1_MOUSE] | 0,000E0 | 0,000E0 | 0,000E0 | 1,459E7 | 2,145E7 | 2,120E7 | Ablim1 |
| Q9D6T0 | Nitric oxide synthase-interacting protein OS=Mus musculus GN=Nosip PE=1 SV=1 - [NOSIP_MOUSE] | 0,000E0 | 0,000E0 | 0,000E0 | 2,716E6 | 1,921E6 | 3,403E6 | Nosip |
| Q3UHK8 | Trinucleotide repeat-containing gene 6A protein OS=Mus musculus GN=Tnrc6a PE=2 SV=1 - [TNR6A_MOUSE] | 0,000E0 | 0,000E0 | 0,000E0 | 1,113E7 | 1,839E7 | 2,219E7 | Tnrc6a |
| Q5NBX1-3 | Isoform 3 of Protein cordon-bleu OS=Mus musculus GN=Cobl - [COBL_MOUSE] | 0,000E0 | 0,000E0 | 0,000E0 | 3,065E6 | 1,138E7 | 1,383E7 | Cobl |
| Q9Z0P5-2 | Isoform 2 of Twinfilin-2 OS=Mus musculus GN=Twf2 - [TWF2_MOUSE] | 0,000E0 | 0,000E0 | 0,000E0 | 2,270E6 | 1,763E7 | 2,147E7 | Twf2 |
| P81270-4 | Isoform 5 of Transcriptional regulator ERG OS=Mus musculus GN=Erg - [ERG_MOUSE] | 0,000E0 | 0,000E0 | 0,000E0 | 1,590E7 | 3,929E7 | 4,297E7 | Erg |
| Q9DCJ5 | NADH dehydrogenase [ubiquinone] 1 alpha subcomplex subunit 8 OS=Mus musculus GN=Ndufa8 PE=1 SV=3 - [NDUA8_MOUSE] | 0,000E0 | 0,000E0 | 0,000E0 | 5,153E6 | 1,979E7 | 3,327E7 | Ndufa8 |
| Q9CX84 | Regulator of G-protein signaling 19 OS=Mus musculus GN=Rgs19 PE=1 SV=1 - [RGS19_MOUSE] | 0,000E0 | 0,000E0 | 0,000E0 | 2,998E6 | 5,032E6 | 1,203E7 | Rgs19 |
| P24452 | Macrophage-capping protein OS=Mus musculus GN=Capg PE=1 SV=2 - [CAPG_MOUSE] | 0,000E0 | 0,000E0 | 0,000E0 | 1,727E7 | 4,524E7 | 5,665E7 | Capg |
| P17156 | Heat shock-related 70 kDa protein 2 OS=Mus musculus GN=Hspa2 PE=1 SV=2 - [HSP72_MOUSE] | 3,628E7 | 0,000E0 | 9,603E6 | 5,026E8 | 1,340E9 | 1,698E9 | Hspa2 |
| Q8BTI8-3 | Isoform 3 of Serine/arginine repetitive matrix protein 2 OS=Mus musculus GN=Srrm2 - [SRRM2_MOUSE] | 0,000E0 | 0,000E0 | 0,000E0 | 6,293E7 | 1,057E8 | 3,756E7 | Srrm2 |
| Q99JR5 | Tubulointerstitial nephritis antigen-like OS=Mus musculus GN=Tinagl1 PE=1 SV=1 - [TINAL_MOUSE] | 0,000E0 | 0,000E0 | 0,000E0 | 2,047E7 | 3,463E7 | 6,051E7 | Tinagl1 |
| Q8CHU3-2 | Isoform 2 of Epsin-2 OS=Mus musculus GN=Epn2 - [EPN2_MOUSE] | 0,000E0 | 0,000E0 | 0,000E0 | 1,593E7 | 3,361E7 | 4,295E7 | Epn2 |
| Q9R0P4 | Small acidic protein OS=Mus musculus GN=Smap PE=1 SV=1 - [SMAP_MOUSE] | 0,000E0 | 0,000E0 | 0,000E0 | 1,453E7 | 2,317E7 | 2,234E7 | Smap |
| Q8K2Q7 | BRO1 domain-containing protein BROX OS=Mus musculus GN=Brox PE=1 SV=1 - [BROX_MOUSE] | 0,000E0 | 0,000E0 | 0,000E0 | 1,442E7 | 2,271E7 | 1,951E7 | Brox |
| Q8CIN4 | Serine/threonine-protein kinase PAK 2 OS=Mus musculus GN=Pak2 PE=1 SV=1 - [PAK2_MOUSE] | 0,000E0 | 0,000E0 | 0,000E0 | 2,749E6 | 6,116E6 | 6,080E6 | Pak2 |
| O35286 | Pre-mRNA-splicing factor ATP-dependent RNA helicase DHX15 OS=Mus musculus GN=Dhx15 PE=1 SV=2 - [DHX15_MOUSE] | 0,000E0 | 0,000E0 | 0,000E0 | 1,157E7 | 2,387E7 | 4,372E7 | Dhx15 |
| Q63844 | Mitogen-activated protein kinase 3 OS=Mus musculus GN=Mapk3 PE=1 SV=5 - [MK03_MOUSE] | 0,000E0 | 0,000E0 | 0,000E0 | 7,220E6 | 2,158E7 | 1,576E7 | Mapk3 |
| Q3UL36 | Arginine and glutamate-rich protein 1 OS=Mus musculus GN=Arglu1 PE=1 SV=2 - [ARGL1_MOUSE] | 0,000E0 | 0,000E0 | 0,000E0 | 2,188E7 | 5,139E7 | 1,013E8 | Arglu1 |
| Q9WVE8 | Protein kinase C and casein kinase substrate in neurons protein 2 OS=Mus musculus GN=Pacsin2 PE=1 SV=1 - [PACN2_MOUSE] | 0,000E0 | 0,000E0 | 0,000E0 | 8,708E6 | 1,935E7 | 2,479E7 | Pacsin2 |
| O35682 | Myeloid-associated differentiation marker OS=Mus musculus GN=Myadm PE=1 SV=2 - [MYADM_MOUSE] | 0,000E0 | 0,000E0 | 0,000E0 | 1,847E7 | 3,261E7 | 6,739E7 | Myadm |
| O70251 | Elongation factor 1-beta OS=Mus musculus GN=Eef1b PE=1 SV=5 - [EF1B_MOUSE] | 0,000E0 | 0,000E0 | 0,000E0 | 8,634E6 | 1,486E7 | 2,198E7 | Eef1b |
| Q922Q8 | Leucine-rich repeat-containing protein 59 OS=Mus musculus GN=Lrrc59 PE=1 SV=1 - [LRC59_MOUSE] | 0,000E0 | 0,000E0 | 0,000E0 | 1,407E7 | 1,216E7 | 1,421E7 | Lrrc59 |
| Q8CIB5 | Fermitin family homolog 2 OS=Mus musculus GN=Fermt2 PE=1 SV=1 - [FERM2_MOUSE] | 0,000E0 | 0,000E0 | 0,000E0 | 5,421E6 | 1,288E7 | 1,880E7 | Fermt2 |
| Q8BRV5 | Uncharacterized protein KIAA1671 OS=Mus musculus GN=Kiaa1671 PE=2 SV=1 - [K1671_MOUSE] | 0,000E0 | 0,000E0 | 0,000E0 | 3,164E6 | 9,524E6 | 1,937E7 | Kiaa1671 |
| Q3UV16 | Inositol 1,4,5-trisphosphate receptor-interacting protein-like 2 OS=Mus musculus GN=Itpripl2 PE=2 SV=1 - [IPIL2_MOUSE] | 0,000E0 | 0,000E0 | 0,000E0 | 7,271E6 | 1,156E7 | 1,591E7 | Itpripl2 |
| Q8CD15 | Bifunctional lysine-specific demethylase and histidyl-hydroxylase MINA OS=Mus musculus GN=Mina PE=1 SV=2 - [MINA_MOUSE] | 0,000E0 | 0,000E0 | 0,000E0 | 6,348E6 | 1,140E7 | 2,259E7 | Mina |
| Q99M87-2 | Isoform 2 of DnaJ homolog subfamily A member 3, mitochondrial OS=Mus musculus GN=Dnaja3 - [DNJA3_MOUSE] | 0,000E0 | 0,000E0 | 0,000E0 | 4,330E6 | 4,381E6 | 2,074E6 | Dnaja3 |
| Q3TLH4 | Protein PRRC2C OS=Mus musculus GN=Prrc2c PE=1 SV=3 - [PRC2C_MOUSE] | 0,000E0 | 0,000E0 | 0,000E0 | 2,502E7 | 4,852E7 | 1,051E8 | Prrc2c |
| Q6PGC1 | ATP-dependent RNA helicase DHX29 OS=Mus musculus GN=Dhx29 PE=1 SV=1 - [DHX29_MOUSE] | 0,000E0 | 0,000E0 | 0,000E0 | 7,366E6 | 1,401E7 | 3,320E7 | Dhx29 |
| Q8BZI0 | Actin filament-associated protein 1-like 1 OS=Mus musculus GN=Afap1l1 PE=1 SV=1 - [AF1L1_MOUSE] | 0,000E0 | 0,000E0 | 0,000E0 | 2,343E6 | 8,594E6 | 1,284E7 | Afap1l1 |
| Q91W53 | Golgin subfamily A member 7 OS=Mus musculus GN=Golga7 PE=1 SV=1 - [GOGA7_MOUSE] | 0,000E0 | 0,000E0 | 0,000E0 | 5,104E6 | 3,380E6 | 3,987E6 | Golga7 |
| Q811D0-2 | Isoform 2 of Disks large homolog 1 OS=Mus musculus GN=Dlg1 - [DLG1_MOUSE] | 0,000E0 | 0,000E0 | 0,000E0 | 5,706E6 | 1,062E7 | 1,638E7 | Dlg1 |
| O08917 | Flotillin-1 OS=Mus musculus GN=Flot1 PE=1 SV=1 - [FLOT1_MOUSE] | 0,000E0 | 0,000E0 | 0,000E0 | 6,363E6 | 1,408E7 | 3,919E7 | Flot1 |
| Q05793 | Basement membrane-specific heparan sulfate proteoglycan core protein OS=Mus musculus GN=Hspg2 PE=1 SV=1 - [PGBM_MOUSE] | 0,000E0 | 0,000E0 | 2,843E5 | 4,050E7 | 4,860E7 | 1,022E8 | Hspg2 |
| Q04899 | Cyclin-dependent kinase 18 OS=Mus musculus GN=Cdk18 PE=1 SV=1 - [CDK18_MOUSE] | 0,000E0 | 0,000E0 | 0,000E0 | 1,843E7 | 4,135E7 | 6,772E7 | Cdk18 |
| Q9R1P4 | Proteasome subunit alpha type-1 OS=Mus musculus GN=Psma1 PE=1 SV=1 - [PSA1_MOUSE] | 0,000E0 | 0,000E0 | 0,000E0 | 2,894E6 | 1,458E6 | 3,993E6 | Psma1 |
| P97762 | Retinitis pigmentosa 9 protein homolog OS=Mus musculus GN=rp9 PE=1 SV=1 - [RP9_MOUSE] | 0,000E0 | 0,000E0 | 0,000E0 | 1,814E7 | 6,423E6 | 8,916E7 | rp9 |
| Q61024 | Asparagine synthetase [glutamine-hydrolyzing] OS=Mus musculus GN=Asns PE=1 SV=3 - [ASNS_MOUSE] | 0,000E0 | 0,000E0 | 0,000E0 | 4,585E6 | 1,146E7 | 1,541E7 | Asns |
| Q60780-2 | Isoform 2 of Growth arrest-specific protein 7 OS=Mus musculus GN=Gas7 - [GAS7_MOUSE] | 0,000E0 | 0,000E0 | 0,000E0 | 4,506E6 | 1,638E7 | 1,713E7 | Gas7 |
| P47964 | 60S ribosomal protein L36 OS=Mus musculus GN=Rpl36 PE=3 SV=2 - [RL36_MOUSE] | 0,000E0 | 0,000E0 | 0,000E0 | 5,638E7 | 2,324E8 | 8,175E8 | Rpl36 |
| Q9EQK5 | Major vault protein OS=Mus musculus GN=Mvp PE=1 SV=4 - [MVP_MOUSE] | 0,000E0 | 0,000E0 | 0,000E0 | 5,282E6 | 8,100E6 | 1,092E7 | Mvp |
| P23249 | Putative helicase MOV-10 OS=Mus musculus GN=Mov10 PE=1 SV=2 - [MOV10_MOUSE] | 0,000E0 | 0,000E0 | 0,000E0 | 9,086E6 | 1,693E7 | 3,938E7 | Mov10 |
| O08784 | Treacle protein OS=Mus musculus GN=Tcof1 PE=1 SV=1 - [TCOF_MOUSE] | 0,000E0 | 0,000E0 | 0,000E0 | 9,476E6 | 1,466E7 | 7,132E7 | Tcof1 |
| O88456 | Calpain small subunit 1 OS=Mus musculus GN=Capns1 PE=1 SV=1 - [CPNS1_MOUSE] | 0,000E0 | 0,000E0 | 0,000E0 | 1,073E7 | 3,470E7 | 4,177E7 | Capns1 |
| Q9QZS3-4 | Isoform 4 of Protein numb homolog OS=Mus musculus GN=Numb - [NUMB_MOUSE] | 0,000E0 | 0,000E0 | 0,000E0 | 1,424E7 | 2,309E7 | 3,875E7 | Numb |
| Q6NZJ6 | Eukaryotic translation initiation factor 4 gamma 1 OS=Mus musculus GN=Eif4g1 PE=1 SV=1 - [IF4G1_MOUSE] | 0,000E0 | 0,000E0 | 0,000E0 | 2,752E7 | 6,198E7 | 9,263E7 | Eif4g1 |
| Q80X41-2 | Isoform 2 of Serine/threonine-protein kinase VRK1 OS=Mus musculus GN=Vrk1 - [VRK1_MOUSE] | 0,000E0 | 0,000E0 | 0,000E0 | 3,237E6 | 9,328E5 | 1,020E7 | Vrk1 |
| Q62130 | Tyrosine-protein phosphatase non-receptor type 14 OS=Mus musculus GN=Ptpn14 PE=1 SV=2 - [PTN14_MOUSE] | 0,000E0 | 0,000E0 | 0,000E0 | 4,449E6 | 1,381E7 | 1,150E7 | Ptpn14 |
| O55222 | Integrin-linked protein kinase OS=Mus musculus GN=Ilk PE=1 SV=2 - [ILK_MOUSE] | 0,000E0 | 0,000E0 | 0,000E0 | 6,128E6 | 2,165E7 | 2,143E7 | Ilk |
| Q9JM52-3 | Isoform 3 of Misshapen-like kinase 1 OS=Mus musculus GN=Mink1 - [MINK1_MOUSE] | 0,000E0 | 0,000E0 | 0,000E0 | 1,194E7 | 2,380E7 | 2,645E7 | Mink1 |
| Q05D44 | Eukaryotic translation initiation factor 5B OS=Mus musculus GN=Eif5b PE=1 SV=2 - [IF2P_MOUSE] | 0,000E0 | 0,000E0 | 0,000E0 | 1,555E7 | 1,733E7 | 3,988E7 | Eif5b |
| Q61792 | LIM and SH3 domain protein 1 OS=Mus musculus GN=Lasp1 PE=1 SV=1 - [LASP1_MOUSE] | 0,000E0 | 0,000E0 | 1,221E6 | 1,556E7 | 3,632E7 | 4,679E7 | Lasp1 |
| P03975 | IgE-binding protein OS=Mus musculus GN=Iap PE=2 SV=1 - [IGEB_MOUSE] | 0,000E0 | 0,000E0 | 0,000E0 | 2,446E7 | 5,838E7 | 3,430E7 | Iap |
| Q9Z0P4 | Paralemmin-1 OS=Mus musculus GN=Palm PE=1 SV=1 - [PALM_MOUSE] | 0,000E0 | 0,000E0 | 0,000E0 | 1,697E6 | 4,314E6 | 1,275E7 | Palm |
| P0C0A3 | Charged multivesicular body protein 6 OS=Mus musculus GN=Chmp6 PE=1 SV=2 - [CHMP6_MOUSE] | 0,000E0 | 0,000E0 | 0,000E0 | 3,477E6 | 8,921E6 | 7,572E6 | Chmp6 |
| P06745 | Glucose-6-phosphate isomerase OS=Mus musculus GN=Gpi PE=1 SV=4 - [G6PI_MOUSE] | 0,000E0 | 0,000E0 | 0,000E0 | 1,591E6 | 5,944E6 | 1,402E7 | Gpi |
| O70152 | Dolichol-phosphate mannosyltransferase subunit 1 OS=Mus musculus GN=Dpm1 PE=1 SV=1 - [DPM1_MOUSE] | 0,000E0 | 0,000E0 | 0,000E0 | 3,099E6 | 7,989E6 | 8,095E6 | Dpm1 |
| P46467 | Vacuolar protein sorting-associated protein 4B OS=Mus musculus GN=Vps4b PE=1 SV=2 - [VPS4B_MOUSE] | 0,000E0 | 0,000E0 | 0,000E0 | 1,535E7 | 3,574E7 | 2,822E7 | Vps4b |
| Q9WVJ2 | 26S proteasome non-ATPase regulatory subunit 13 OS=Mus musculus GN=Psmd13 PE=1 SV=1 - [PSD13_MOUSE] | 0,000E0 | 0,000E0 | 0,000E0 | 1,557E6 | 4,578E6 | 6,930E6 | Psmd13 |
| P58871 | 182 kDa tankyrase-1-binding protein OS=Mus musculus GN=Tnks1bp1 PE=1 SV=2 - [TB182_MOUSE] | 0,000E0 | 0,000E0 | 0,000E0 | 8,133E6 | 1,472E7 | 2,318E7 | Tnks1bp1 |
| Q8R1S4-2 | Isoform 2 of Metastasis suppressor protein 1 OS=Mus musculus GN=Mtss1 - [MTSS1_MOUSE] | 0,000E0 | 0,000E0 | 0,000E0 | 5,823E6 | 5,977E6 | 1,087E7 | Mtss1 |
| Q9D083-3 | Isoform 3 of Kinetochore protein Spc24 OS=Mus musculus GN=Spc24 - [SPC24_MOUSE] | 0,000E0 | 0,000E0 | 0,000E0 | 8,734E5 | 2,966E6 | 1,288E6 | Spc24 |
| P17095 | High mobility group protein HMG-I/HMG-Y OS=Mus musculus GN=Hmga1 PE=1 SV=4 - [HMGA1_MOUSE] | 0,000E0 | 0,000E0 | 0,000E0 | 7,761E6 | 7,456E6 | 5,975E7 | Hmga1 |
| P70168 | Importin subunit beta-1 OS=Mus musculus GN=Kpnb1 PE=1 SV=2 - [IMB1_MOUSE] | 0,000E0 | 0,000E0 | 0,000E0 | 2,376E7 | 2,902E7 | 7,291E7 | Kpnb1 |
| Q8BWW4 | La-related protein 4 OS=Mus musculus GN=Larp4 PE=1 SV=2 - [LARP4_MOUSE] | 0,000E0 | 0,000E0 | 0,000E0 | 2,113E7 | 4,057E7 | 6,761E7 | Larp4 |
| Q8VI94 | 2'-5'-oligoadenylate synthase-like protein 1 OS=Mus musculus GN=Oasl1 PE=2 SV=1 - [OASL1_MOUSE] | 0,000E0 | 0,000E0 | 0,000E0 | 7,767E6 | 3,928E7 | 6,882E7 | Oasl1 |
| Q8BJS4-3 | Isoform 3 of SUN domain-containing protein 2 OS=Mus musculus GN=Sun2 - [SUN2_MOUSE] | 0,000E0 | 0,000E0 | 0,000E0 | 2,530E6 | 7,310E6 | 1,358E7 | Sun2 |
| O70493 | Sorting nexin-12 OS=Mus musculus GN=Snx12 PE=1 SV=1 - [SNX12_MOUSE] | 0,000E0 | 0,000E0 | 0,000E0 | 1,689E6 | 9,900E6 | 1,219E7 | Snx12 |

| | | | | | | | | |
|---|---|---|---|---|---|---|---|---|
| Q80UM3 | N-alpha-acetyltransferase 15, NatA auxiliary subunit OS=Mus musculus GN=Naa15 PE=1 SV=1 - [NAA15_MOUSE] | 0,000E0 | 0,000E0 | 0,000E0 | 9,395E6 | 1,095E7 | 2,746E7 | Naa15 |
| A2A5Z6 | E3 ubiquitin-protein ligase SMURF2 OS=Mus musculus GN=Smurf2 PE=1 SV=1 - [SMUF2_MOUSE] | 0,000E0 | 0,000E0 | 0,000E0 | 9,907E6 | 2,289E7 | 2,449E7 | Smurf2 |
| Q8K1N4 | Spermatogenesis-associated serine-rich protein 2 OS=Mus musculus GN=Spats2 PE=1 SV=1 - [SPAS2_MOUSE] | 0,000E0 | 0,000E0 | 0,000E0 | 4,632E6 | 3,335E6 | 1,913E7 | Spats2 |
| Q76N33 | AMSH-like protease OS=Mus musculus GN=Stambpl1 PE=1 SV=1 - [STALP_MOUSE] | 0,000E0 | 0,000E0 | 0,000E0 | 4,477E6 | 1,332E7 | 1,445E7 | Stambpl1 |
| P62196 | 26S protease regulatory subunit 8 OS=Mus musculus GN=Psmc5 PE=1 SV=1 - [PRS8_MOUSE] | 0,000E0 | 0,000E0 | 0,000E0 | 6,374E5 | 3,117E6 | 4,484E6 | Psmc5 |
| P11352 | Glutathione peroxidase 1 OS=Mus musculus GN=Gpx1 PE=1 SV=2 - [GPX1_MOUSE] | 0,000E0 | 0,000E0 | 0,000E0 | 1,390E6 | 2,417E6 | 7,535E6 | Gpx1 |
| Q61753 | D-3-phosphoglycerate dehydrogenase OS=Mus musculus GN=Phgdh PE=1 SV=3 - [SERA_MOUSE] | 0,000E0 | 0,000E0 | 0,000E0 | 1,120E7 | 2,966E7 | 3,892E7 | Phgdh |
| Q9JII7 | AN1-type zinc finger protein 2A OS=Mus musculus GN=Zfand2a PE=2 SV=1 - [ZFN2A_MOUSE] | 0,000E0 | 0,000E0 | 0,000E0 | 2,063E6 | 1,942E7 | 1,404E7 | Zfand2a |
| P97820 | Mitogen-activated protein kinase kinase kinase kinase 4 OS=Mus musculus GN=Map4k4 PE=1 SV=1 - [M4K4_MOUSE] | 0,000E0 | 0,000E0 | 7,335E5 | 1,709E7 | 3,924E7 | 5,588E7 | Map4k4 |
| P63280 | SUMO-conjugating enzyme UBC9 OS=Mus musculus GN=Ube2i PE=1 SV=1 - [UBC9_MOUSE] | 0,000E0 | 0,000E0 | 0,000E0 | 3,191E5 | 5,271E6 | 5,223E6 | Ube2i |
| Q9JI10-2 | Isoform 2 of Serine/threonine-protein kinase 3 OS=Mus musculus GN=Stk3 - [STK3_MOUSE] | 0,000E0 | 0,000E0 | 0,000E0 | 2,199E6 | 3,381E6 | 2,556E6 | Stk3 |
| Q3UHX2 | 28 kDa heat- and acid-stable phosphoprotein OS=Mus musculus GN=Pdap1 PE=1 SV=1 - [HAP28_MOUSE] | 0,000E0 | 0,000E0 | 0,000E0 | 3,005E7 | 4,895E7 | 7,068E7 | Pdap1 |
| P61211 | ADP-ribosylation factor-like protein 1 OS=Mus musculus GN=Arl1 PE=1 SV=1 - [ARL1_MOUSE] | 0,000E0 | 0,000E0 | 0,000E0 | 5,809E6 | 8,103E6 | 1,017E7 | Arl1 |
| P46061 | Ran GTPase-activating protein 1 OS=Mus musculus GN=Rangap1 PE=1 SV=2 - [RAGP1_MOUSE] | 0,000E0 | 0,000E0 | 0,000E0 | 8,310E5 | 9,599E6 | 2,210E7 | Rangap1 |
| A2A699-2 | Isoform 2 of Protein FAM171A2 OS=Mus musculus GN=Fam171a2 - [F1712_MOUSE] | 0,000E0 | 0,000E0 | 0,000E0 | 3,642E6 | 6,221E6 | 7,218E6 | Fam171a2 |
| Q3UH60 | Disco-interacting protein 2 homolog B OS=Mus musculus GN=Dip2b PE=1 SV=1 - [DIP2B_MOUSE] | 0,000E0 | 0,000E0 | 0,000E0 | 1,078E7 | 2,206E7 | 3,748E7 | Dip2b |
| Q8VBX6-2 | Isoform 2 of Multiple PDZ domain protein OS=Mus musculus GN=Mpdz - [MPDZ_MOUSE] | 0,000E0 | 0,000E0 | 0,000E0 | 1,728E7 | 2,365E7 | 2,371E7 | Mpdz |
| Q8CFE2 | Histone PARylation factor 1 OS=Mus musculus GN=Hpf1 PE=1 SV=1 - [HPF1_MOUSE] | 0,000E0 | 0,000E0 | 0,000E0 | 2,220E6 | 2,362E6 | 4,225E6 | Hpf1 |
| Q8CHC4 | Synaptojanin-1 OS=Mus musculus GN=Synj1 PE=1 SV=3 - [SYNJ1_MOUSE] | 0,000E0 | 0,000E0 | 0,000E0 | 4,993E6 | 8,675E6 | 9,989E6 | Synj1 |
| Q6A0A9 | Constitutive coactivator of PPAR-gamma-like protein 1 OS=Mus musculus GN=FAM120A PE=1 SV=2 - [F120A_MOUSE] | 0,000E0 | 0,000E0 | 0,000E0 | 5,463E6 | 2,658E7 | 5,671E7 | FAM120A |
| O88384 | Vesicle transport through interaction with t-SNAREs homolog 1B OS=Mus musculus GN=Vti1b PE=1 SV=1 - [VTI1B_MOUSE] | 0,000E0 | 0,000E0 | 0,000E0 | 4,794E6 | 8,905E6 | 2,242E6 | Vti1b |
| P06869 | Urokinase-type plasminogen activator OS=Mus musculus GN=Plau PE=1 SV=1 - [UROK_MOUSE] | 0,000E0 | 0,000E0 | 0,000E0 | 5,666E6 | 8,101E6 | 3,151E7 | Plau |
| A2A6T1 | Cerebellar degeneration-related protein 2-like OS=Mus musculus GN=Cdr2l PE=1 SV=1 - [CDR2L_MOUSE] | 0,000E0 | 0,000E0 | 0,000E0 | 3,174E6 | 1,327E7 | 1,679E7 | Cdr2l |
| Q922S8 | Kinesin-like protein KIF2C OS=Mus musculus GN=Kif2c PE=1 SV=1 - [KIF2C_MOUSE] | 0,000E0 | 0,000E0 | 0,000E0 | 1,088E7 | 1,358E7 | 1,373E7 | Kif2c |
| Q3UHJ0-2 | Isoform 2 of AP2-associated protein kinase 1 OS=Mus musculus GN=Aak1 - [AAK1_MOUSE] | 0,000E0 | 0,000E0 | 0,000E0 | 6,965E6 | 1,404E7 | 1,393E7 | Aak1 |
| Q9CQ71 | Replication protein A 14 kDa subunit OS=Mus musculus GN=Rpa3 PE=1 SV=1 - [RFA3_MOUSE] | 0,000E0 | 0,000E0 | 0,000E0 | 3,372E6 | 3,016E6 | 4,829E6 | Rpa3 |
| Q6NXY1 | TBC1 domain family member 31 OS=Mus musculus GN=Tbc1d31 PE=1 SV=1 - [TBC31_MOUSE] | 0,000E0 | 0,000E0 | 0,000E0 | 2,719E6 | 4,502E6 | 6,390E6 | Tbc1d31 |
| Q921J2 | GTP-binding protein Rheb OS=Mus musculus GN=Rheb PE=1 SV=1 - [RHEB_MOUSE] | 0,000E0 | 0,000E0 | 0,000E0 | 1,711E6 | 7,163E6 | 9,167E6 | Rheb |
| Q80Z96 | Vang-like protein 1 OS=Mus musculus GN=Vangl1 PE=1 SV=2 - [VANG1_MOUSE] | 0,000E0 | 0,000E0 | 0,000E0 | 8,226E6 | 1,761E7 | 2,405E7 | Vangl1 |
| Q923T9-3 | Isoform 3 of Calcium/calmodulin-dependent protein kinase type II subunit gamma OS=Mus musculus GN=Camk2g - [KCC2G_MOUSE] | 0,000E0 | 0,000E0 | 0,000E0 | 2,093E7 | 1,749E7 | 4,232E7 | Camk2g |
| Q9Z0G0 | PDZ domain-containing protein GIPC1 OS=Mus musculus GN=Gipc1 PE=1 SV=1 - [GIPC1_MOUSE] | 0,000E0 | 0,000E0 | 0,000E0 | 2,849E6 | 2,383E6 | 3,256E6 | Gipc1 |
| Q9JLV1 | BAG family molecular chaperone regulator 3 OS=Mus musculus GN=Bag3 PE=1 SV=2 - [BAG3_MOUSE] | 0,000E0 | 0,000E0 | 0,000E0 | 2,944E6 | 5,236E6 | 6,464E6 | Bag3 |
| O88685 | 26S protease regulatory subunit 6A OS=Mus musculus GN=Psmc3 PE=1 SV=2 - [PRS6A_MOUSE] | 0,000E0 | 0,000E0 | 0,000E0 | 2,886E6 | 2,997E6 | 7,228E6 | Psmc3 |
| Q61081 | Hsp90 co-chaperone Cdc37 OS=Mus musculus GN=Cdc37 PE=1 SV=1 - [CDC37_MOUSE] | 0,000E0 | 0,000E0 | 0,000E0 | 9,211E6 | 6,098E6 | 2,902E7 | Cdc37 |
| Q9DCL9 | Multifunctional protein ADE2 OS=Mus musculus GN=Paics PE=1 SV=4 - [PUR6_MOUSE] | 0,000E0 | 0,000E0 | 0,000E0 | 9,520E6 | 9,746E6 | 2,651E7 | Paics |
| Q8VDF2-2 | Isoform 2 of E3 ubiquitin-protein ligase UHRF1 OS=Mus musculus GN=Uhrf1 - [UHRF1_MOUSE] | 0,000E0 | 0,000E0 | 0,000E0 | 5,050E6 | 6,417E6 | 1,313E7 | Uhrf1 |
| Q8C804 | Spindle and centriole-associated protein 1 OS=Mus musculus GN=Spice1 PE=1 SV=2 - [SPICE_MOUSE] | 0,000E0 | 0,000E0 | 0,000E0 | 2,627E6 | 4,336E6 | 1,290E6 | Spice1 |
| Q80SY4 | E3 ubiquitin-protein ligase MIB1 OS=Mus musculus GN=Mib1 PE=1 SV=1 - [MIB1_MOUSE] | 0,000E0 | 0,000E0 | 0,000E0 | 9,291E6 | 1,315E7 | 1,944E7 | Mib1 |
| Q9D0B0 | Serine/arginine-rich splicing factor 9 OS=Mus musculus GN=Srsf9 PE=1 SV=1 - [SRSF9_MOUSE] | 0,000E0 | 0,000E0 | 0,000E0 | 7,898E6 | 1,969E7 | 2,276E7 | Srsf9 |
| Q91YU6 | Leucine zipper putative tumor suppressor 2 OS=Mus musculus GN=Lzts2 PE=1 SV=3 - [LZTS2_MOUSE] | 0,000E0 | 0,000E0 | 0,000E0 | 5,866E6 | 8,866E6 | 1,170E7 | Lzts2 |
| Q6P9Q6 | FK506-binding protein 15 OS=Mus musculus GN=Fkbp15 PE=1 SV=2 - [FKB15_MOUSE] | 0,000E0 | 0,000E0 | 0,000E0 | 1,176E7 | 2,349E7 | 3,190E7 | Fkbp15 |
| P59326 | YTH domain-containing family protein 1 OS=Mus musculus GN=Ythdf1 PE=1 SV=1 - [YTHD1_MOUSE] | 0,000E0 | 0,000E0 | 5,369E6 | 1,748E6 | 8,562E6 | 2,171E7 | Ythdf1 |
| Q8CJG0 | Protein argonaute-2 OS=Mus musculus GN=Ago2 PE=1 SV=3 - [AGO2_MOUSE] | 0,000E0 | 0,000E0 | 0,000E0 | 9,572E6 | 2,013E7 | 2,308E7 | Ago2 |
| Q922J9 | Fatty acyl-CoA reductase 1 OS=Mus musculus GN=Far1 PE=1 SV=1 - [FACR1_MOUSE] | 0,000E0 | 0,000E0 | 0,000E0 | 4,808E6 | 1,573E7 | 2,175E7 | Far1 |
| P46471 | 26S protease regulatory subunit 7 OS=Mus musculus GN=Psmc2 PE=1 SV=5 - [PRS7_MOUSE] | 0,000E0 | 0,000E0 | 0,000E0 | 3,186E6 | 4,674E6 | 1,178E7 | Psmc2 |
| Q91VJ2 | Protein kinase C delta-binding protein OS=Mus musculus GN=Prkcdbp PE=1 SV=1 - [PRDBP_MOUSE] | 0,000E0 | 0,000E0 | 0,000E0 | 1,960E7 | 2,419E7 | 1,155E8 | Prkcdbp |
| Q9CXW4 | 60S ribosomal protein L11 OS=Mus musculus GN=Rpl11 PE=1 SV=4 - [RL11_MOUSE] | 3,381E7 | 0,000E0 | 1,574E7 | 2,628E8 | 4,519E8 | 8,359E8 | Rpl11 |
| P70336 | Rho-associated protein kinase 2 OS=Mus musculus GN=Rock2 PE=1 SV=1 - [ROCK2_MOUSE] | 0,000E0 | 0,000E0 | 1,352E5 | 8,759E6 | 1,528E7 | 2,314E7 | Rock2 |
| P63034-3 | Isoform 3 of Cytohesin-2 OS=Mus musculus GN=Cyth2 - [CYH2_MOUSE] | 0,000E0 | 0,000E0 | 0,000E0 | 6,002E6 | 1,902E7 | 1,211E7 | Cyth2 |
| P97427 | Dihydropyrimidinase-related protein 1 OS=Mus musculus GN=Crmp1 PE=1 SV=1 - [DPYL1_MOUSE] | 0,000E0 | 0,000E0 | 0,000E0 | 3,092E6 | 3,970E6 | 2,221E7 | Crmp1 |
| Q3UGC7 | Eukaryotic translation initiation factor 3 subunit J-A OS=Mus musculus GN=Eif3j1 PE=2 SV=1 - [EI3JA_MOUSE] | 0,000E0 | 0,000E0 | 0,000E0 | 2,092E6 | 1,031E7 | 1,266E7 | Eif3j1 |
| Q6DFW4 | Nucleolar protein 58 OS=Mus musculus GN=Nop58 PE=1 SV=1 - [NOP58_MOUSE] | 0,000E0 | 0,000E0 | 0,000E0 | 2,212E6 | 7,028E6 | 2,353E7 | Nop58 |
| O70503 | Very-long-chain 3-oxoacyl-CoA reductase OS=Mus musculus GN=Hsd17b12 PE=1 SV=1 - [DHB12_MOUSE] | 0,000E0 | 0,000E0 | 0,000E0 | 6,565E6 | 3,142E7 | 3,408E7 | Hsd17b12 |
| Q9CXY6 | Interleukin enhancer-binding factor 2 OS=Mus musculus GN=Ilf2 PE=1 SV=1 - [ILF2_MOUSE] | 0,000E0 | 0,000E0 | 0,000E0 | 2,362E6 | 6,575E5 | 4,972E6 | Ilf2 |
| Q8K2J7-2 | Isoform 2 of RELT-like protein 1 OS=Mus musculus GN=Rell1 - [RELL1_MOUSE] | 0,000E0 | 0,000E0 | 0,000E0 | 2,517E6 | 4,497E6 | 1,202E7 | Rell1 |
| Q9DD03 | Ras-related protein Rab-13 OS=Mus musculus GN=Rab13 PE=1 SV=1 - [RAB13_MOUSE] | 0,000E0 | 0,000E0 | 0,000E0 | 3,615E7 | 6,702E7 | 6,617E7 | Rab13 |
| Q3UFY8 | Mitochondrial ribonuclease P protein 1 OS=Mus musculus GN=Trmt10c PE=1 SV=2 - [MRRP1_MOUSE] | 0,000E0 | 0,000E0 | 0,000E0 | 2,560E6 | 1,396E7 | 1,870E7 | Trmt10c |
| O70311 | Glycylpeptide N-tetradecanoyltransferase 2 OS=Mus musculus GN=Nmt2 PE=1 SV=1 - [NMT2_MOUSE] | 0,000E0 | 0,000E0 | 0,000E0 | 5,967E6 | 1,059E7 | 1,900E7 | Nmt2 |
| P11276 | Fibronectin OS=Mus musculus GN=Fn1 PE=1 SV=4 - [FINC_MOUSE] | 0,000E0 | 0,000E0 | 0,000E0 | 1,421E7 | 3,555E7 | 6,967E7 | Fn1 |
| Q07832 | Serine/threonine-protein kinase PLK1 OS=Mus musculus GN=Plk1 PE=1 SV=2 - [PLK1_MOUSE] | 0,000E0 | 0,000E0 | 0,000E0 | 3,303E6 | 6,468E6 | 1,424E7 | Plk1 |
| P63038 | 60 kDa heat shock protein, mitochondrial OS=Mus musculus GN=Hspd1 PE=1 SV=1 - [CH60_MOUSE] | 0,000E0 | 0,000E0 | 2,327E6 | 1,869E6 | 3,667E6 | 6,890E6 | Hspd1 |
| Q8JZX3-3 | Isoform 3 of POC1 centriolar protein homolog A OS=Mus musculus GN=Poc1a - [POC1A_MOUSE] | 0,000E0 | 0,000E0 | 0,000E0 | 1,943E6 | 2,016E6 | 3,526E6 | Poc1a |
| P00493 | Hypoxanthine-guanine phosphoribosyltransferase OS=Mus musculus GN=Hprt1 PE=1 SV=3 - [HPRT_MOUSE] | 0,000E0 | 0,000E0 | 0,000E0 | 1,010E6 | 3,314E6 | 4,216E6 | Hprt1 |
| Q9DBU6 | Serine/Arginine-related protein 53 OS=Mus musculus GN=Rsrc1 PE=1 SV=1 - [RSRC1_MOUSE] | 0,000E0 | 0,000E0 | 0,000E0 | 7,670E6 | 7,425E6 | 2,003E7 | Rsrc1 |
| Q0VGY8 | Protein TANC1 OS=Mus musculus GN=Tanc1 PE=1 SV=2 - [TANC1_MOUSE] | 0,000E0 | 0,000E0 | 0,000E0 | 4,532E6 | 1,065E7 | 9,495E6 | Tanc1 |
| O55029 | Coatomer subunit beta' OS=Mus musculus GN=Copb2 PE=1 SV=2 - [COPB2_MOUSE] | 0,000E0 | 0,000E0 | 0,000E0 | 3,313E6 | 5,361E6 | 5,242E6 | Copb2 |
| Q9EPK2-3 | Isoform 3 of Protein XRP2 OS=Mus musculus GN=Rp2 - [XRP2_MOUSE] | 0,000E0 | 0,000E0 | 0,000E0 | 6,839E6 | 2,917E6 | 4,658E7 | Rp2 |
| A2AJI0 | MAP7 domain-containing protein 1 OS=Mus musculus GN=Map7d1 PE=1 SV=1 - [MA7D1_MOUSE] | 0,000E0 | 0,000E0 | 0,000E0 | 2,432E6 | 9,492E6 | 3,072E7 | Map7d1 |
| B2RY56 | RNA-binding protein 25 OS=Mus musculus GN=Rbm25 PE=1 SV=2 - [RBM25_MOUSE] | 0,000E0 | 0,000E0 | 0,000E0 | 7,150E6 | 1,567E7 | 3,457E7 | Rbm25 |
| Q9JJK2 | LanC-like protein 2 OS=Mus musculus GN=Lancl2 PE=1 SV=1 - [LANC2_MOUSE] | 0,000E0 | 0,000E0 | 0,000E0 | 2,190E7 | 1,958E7 | 1,622E7 | Lancl2 |
| P14602-2 | Isoform B of Heat shock protein beta-1 OS=Mus musculus GN=Hspb1 - [HSPB1_MOUSE] | 0,000E0 | 0,000E0 | 6,467E6 | 7,573E6 | 1,754E7 | 8,848E6 | Hspb1 |
| Q3TW96 | UDP-N-acetylhexosamine pyrophosphorylase-like protein 1 OS=Mus musculus GN=Uap1l1 PE=1 SV=1 - [UAP1L_MOUSE] | 0,000E0 | 0,000E0 | 0,000E0 | 3,531E6 | 9,751E6 | 1,701E7 | Uap1l1 |
| Q3UIA2 | Rho GTPase-activating protein 17 OS=Mus musculus GN=Arhgap17 PE=1 SV=1 - [RHG17_MOUSE] | 0,000E0 | 0,000E0 | 0,000E0 | 2,402E6 | 7,691E6 | 6,211E6 | Arhgap17 |
| Q99JN2 | Kelch-like protein 22 OS=Mus musculus GN=Klhl22 PE=1 SV=1 - [KLH22_MOUSE] | 0,000E0 | 0,000E0 | 0,000E0 | 3,111E6 | 5,413E6 | 1,810E6 | Klhl22 |
| Q8BMA6 | Signal recognition particle subunit SRP68 OS=Mus musculus GN=Srp68 PE=1 SV=2 - [SRP68_MOUSE] | 0,000E0 | 0,000E0 | 0,000E0 | 4,619E6 | 4,176E6 | 9,298E6 | Srp68 |
| P30285 | Cyclin-dependent kinase 4 OS=Mus musculus GN=Cdk4 PE=1 SV=1 - [CDK4_MOUSE] | 0,000E0 | 0,000E0 | 0,000E0 | 9,391E6 | 2,088E7 | 3,528E7 | Cdk4 |
| Q91YD3 | mRNA-decapping enzyme 1A OS=Mus musculus GN=Dcp1a PE=1 SV=1 - [DCP1A_MOUSE] | 0,000E0 | 0,000E0 | 0,000E0 | 3,234E6 | 4,938E6 | 1,499E7 | Dcp1a |
| Q5SXY1-3 | Isoform 4 of Cytospin-B OS=Mus musculus GN=Specc1 - [CYTSB_MOUSE] | 0,000E0 | 0,000E0 | 0,000E0 | 3,509E6 | 1,006E7 | 1,905E7 | Specc1 |
| Q9ET54-4 | Isoform 4 of Palladin OS=Mus musculus GN=Palld - [PALLD_MOUSE] | 0,000E0 | 0,000E0 | 0,000E0 | 1,035E7 | 2,581E7 | 2,404E7 | Palld |
| P61963 | DDB1- and CUL4-associated factor 7 OS=Mus musculus GN=Dcaf7 PE=1 SV=1 - [DCAF7_MOUSE] | 0,000E0 | 0,000E0 | 0,000E0 | 2,564E6 | 5,953E6 | 8,447E6 | Dcaf7 |
| Q8VEH3 | ADP-ribosylation factor-like protein 8A OS=Mus musculus GN=Arl8a PE=1 SV=1 - [ARL8A_MOUSE] | 0,000E0 | 0,000E0 | 0,000E0 | 3,034E6 | 4,126E6 | 3,245E6 | Arl8a |
| P61759 | Prefoldin subunit 3 OS=Mus musculus GN=Vbp1 PE=1 SV=2 - [PFD3_MOUSE] | 0,000E0 | 0,000E0 | 0,000E0 | 1,069E6 | 2,875E6 | 2,854E6 | Vbp1 |
| Q9JMK2 | Casein kinase I isoform epsilon OS=Mus musculus GN=Csnk1e PE=1 SV=2 - [KC1E_MOUSE] | 0,000E0 | 0,000E0 | 0,000E0 | 1,802E7 | 1,414E7 | 3,542E7 | Csnk1e |
| Q6Y7W8 | PERQ amino acid-rich with GYF domain-containing protein 2 OS=Mus musculus GN=Gigyf2 PE=1 SV=2 - [PERQ2_MOUSE] | 0,000E0 | 0,000E0 | 0,000E0 | 6,806E6 | 1,734E7 | 5,043E7 | Gigyf2 |
| Q62318 | Transcription intermediary factor 1-beta OS=Mus musculus GN=Trim28 PE=1 SV=3 - [TIF1B_MOUSE] | 0,000E0 | 0,000E0 | 0,000E0 | 4,817E6 | 1,123E7 | 1,715E7 | Trim28 |
| P21981 | Protein-glutamine gamma-glutamyltransferase 2 OS=Mus musculus GN=Tgm2 PE=1 SV=4 - [TGM2_MOUSE] | 0,000E0 | 0,000E0 | 0,000E0 | 4,802E6 | 9,148E6 | 2,607E7 | Tgm2 |
| Q9ESX5 | H/ACA ribonucleoprotein complex subunit 4 OS=Mus musculus GN=Dkc1 PE=1 SV=4 - [DKC1_MOUSE] | 0,000E0 | 0,000E0 | 0,000E0 | 3,347E6 | 5,017E6 | 9,911E6 | Dkc1 |
| Q9CZ44 | NSFL1 cofactor p47 OS=Mus musculus GN=Nsfl1c PE=1 SV=1 - [NSF1C_MOUSE] | 0,000E0 | 0,000E0 | 0,000E0 | 1,622E6 | 5,938E6 | 9,790E6 | Nsfl1c |
| P47941 | Crk-like protein OS=Mus musculus GN=Crkl PE=1 SV=2 - [CRKL_MOUSE] | 0,000E0 | 0,000E0 | 0,000E0 | 1,295E6 | 1,866E6 | 4,185E6 | Crkl |
| P81270-3 | Isoform 1 of Transcriptional regulator ERG OS=Mus musculus GN=Erg - [ERG_MOUSE] | 0,000E0 | 0,000E0 | 0,000E0 | 1,590E7 | 3,929E7 | 4,297E7 | Erg |
| Q91VX2 | Ubiquitin-associated protein 2 OS=Mus musculus GN=Ubap2 PE=1 SV=1 - [UBAP2_MOUSE] | 0,000E0 | 0,000E0 | 0,000E0 | 2,538E7 | 3,871E7 | 5,953E7 | Ubap2 |
| P56959 | RNA-binding protein FUS OS=Mus musculus GN=Fus PE=1 SV=1 - [FUS_MOUSE] | 0,000E0 | 0,000E0 | 0,000E0 | 1,050E8 | 7,401E7 | 1,250E8 | Fus |
| Q8C570 | mRNA export factor OS=Mus musculus GN=Rae1 PE=1 SV=1 - [RAE1L_MOUSE] | 0,000E0 | 0,000E0 | 0,000E0 | 4,069E6 | 7,391E6 | 5,248E6 | Rae1 |

| | | | | | | | | |
|---|---|---|---|---|---|---|---|---|
| Q7TNC4-2 | Isoform 2 of Putative RNA-binding protein Luc7-like 2 OS=Mus musculus GN=Luc7l2 - [LC7L2_MOUSE] | 0,000E0 | 0,000E0 | 0,000E0 | 1,859E7 | 1,285E7 | 1,522E7 | Luc7l2 |
| Q8C650-2 | Isoform 2 of Septin-10 OS=Mus musculus GN=Sept10 - [SEP10_MOUSE] | 0,000E0 | 0,000E0 | 0,000E0 | 1,377E7 | 2,659E7 | 1,836E7 | SEP10 |
| O70479 | BTB/POZ domain-containing adapter for CUL3-mediated RhoA degradation protein 2 OS=Mus musculus GN=Tnfaip1 PE=2 SV=1 - [BACD2_MOUSE] | 0,000E0 | 0,000E0 | 0,000E0 | 1,470E6 | 1,089E6 | 5,919E6 | Tnfaip1 |
| P97376 | Protein FRG1 OS=Mus musculus GN=Frg1 PE=1 SV=2 - [FRG1_MOUSE] | 0,000E0 | 0,000E0 | 0,000E0 | 5,329E6 | 8,235E6 | 2,307E7 | Frg1 |
| Q8BH59 | Calcium-binding mitochondrial carrier protein Aralar1 OS=Mus musculus GN=Slc25a12 PE=1 SV=1 - [CMC1_MOUSE] | 0,000E0 | 0,000E0 | 0,000E0 | 3,337E6 | 6,330E6 | 1,401E7 | Slc25a12 |
| Q80VP1 | Epsin-1 OS=Mus musculus GN=Epn1 PE=1 SV=3 - [EPN1_MOUSE] | 0,000E0 | 0,000E0 | 0,000E0 | 1,231E7 | 1,602E7 | 1,741E7 | Epn1 |
| Q60770 | Syntaxin-binding protein 3 OS=Mus musculus GN=Stxbp3 PE=1 SV=1 - [STXB3_MOUSE] | 0,000E0 | 0,000E0 | 0,000E0 | 5,633E6 | 1,007E7 | 2,380E7 | Stxbp3 |
| Q9DBF7 | Pre-mRNA-splicing factor CWC25 homolog OS=Mus musculus GN=Cwc25 PE=2 SV=2 - [CWC25_MOUSE] | 0,000E0 | 0,000E0 | 0,000E0 | 8,290E6 | 2,534E7 | 5,619E7 | Cwc25 |
| Q6P9J9 | Anoctamin-6 OS=Mus musculus GN=Ano6 PE=1 SV=1 - [ANO6_MOUSE] | 0,000E0 | 0,000E0 | 0,000E0 | 1,782E7 | 7,361E7 | 5,032E7 | Ano6 |
| P55012 | Solute carrier family 12 member 2 OS=Mus musculus GN=Slc12a2 PE=1 SV=2 - [S12A2_MOUSE] | 0,000E0 | 0,000E0 | 0,000E0 | 1,428E7 | 4,314E7 | 7,206E7 | Slc12a2 |
| Q69ZH9 | Rho GTPase-activating protein 23 OS=Mus musculus GN=Arhgap23 PE=1 SV=2 - [RHG23_MOUSE] | 0,000E0 | 0,000E0 | 0,000E0 | 6,428E6 | 1,428E7 | 1,929E7 | Arhgap23 |
| O88983 | Syntaxin-8 OS=Mus musculus GN=Stx8 PE=1 SV=1 - [STX8_MOUSE] | 0,000E0 | 0,000E0 | 0,000E0 | 1,409E6 | 2,829E6 | 6,116E6 | Stx8 |
| P19096 | Fatty acid synthase OS=Mus musculus GN=Fasn PE=1 SV=2 - [FAS_MOUSE] | 0,000E0 | 0,000E0 | 0,000E0 | 1,005E7 | 1,049E8 | 3,513E7 | Fasn |
| Q61595-3 | Isoform 3 of Kinectin OS=Mus musculus GN=Ktn1 - [KTN1_MOUSE] | 0,000E0 | 0,000E0 | 0,000E0 | 3,072E6 | 5,064E6 | 1,429E7 | Ktn1 |
| Q61696 | Heat shock 70 kDa protein 1A OS=Mus musculus GN=Hspa1a PE=1 SV=2 - [HS71A_MOUSE] | 4,285E7 | 0,000E0 | 6,668E6 | 6,497E8 | 3,755E8 | 4,452E8 | Hspa1a |
| Q9WV60 | Glycogen synthase kinase-3 beta OS=Mus musculus GN=Gsk3b PE=1 SV=2 - [GSK3B_MOUSE] | 0,000E0 | 0,000E0 | 0,000E0 | 7,454E6 | 2,103E6 | 1,642E7 | Gsk3b |
| Q9EQH3 | Vacuolar protein sorting-associated protein 35 OS=Mus musculus GN=Vps35 PE=1 SV=1 - [VPS35_MOUSE] | 0,000E0 | 0,000E0 | 4,640E6 | 3,466E6 | 1,376E7 | 1,532E7 | Vps35 |
| Q8BP47 | Asparagine--tRNA ligase, cytoplasmic OS=Mus musculus GN=Nars PE=1 SV=2 - [SYNC_MOUSE] | 0,000E0 | 0,000E0 | 0,000E0 | 2,679E6 | 9,245E6 | 2,191E7 | Nars |
| Q80U72 | Protein scribble homolog OS=Mus musculus GN=Scrib PE=1 SV=2 - [SCRIB_MOUSE] | 0,000E0 | 0,000E0 | 0,000E0 | 4,114E6 | 1,275E7 | 1,263E7 | Scrib |
| O08807 | Peroxiredoxin-4 OS=Mus musculus GN=Prdx4 PE=1 SV=1 - [PRDX4_MOUSE] | 0,000E0 | 0,000E0 | 0,000E0 | 7,418E7 | 1,501E8 | 2,181E8 | Prdx4 |
| Q3UMC0 | Spermatogenesis-associated protein 5 OS=Mus musculus GN=Spata5 PE=1 SV=2 - [SPAT5_MOUSE] | 0,000E0 | 0,000E0 | 0,000E0 | 1,559E6 | 1,190E7 | 5,678E6 | Spata5 |
| O08638-2 | Isoform 2 of Myosin-11 OS=Mus musculus GN=Myh11 - [MYH11_MOUSE] | 2,336E8 | 0,000E0 | 2,340E8 | 3,928E9 | 7,665E9 | 5,658E9 | Myh11 |
| Q9CQ62 | 2,4-dienoyl-CoA reductase, mitochondrial OS=Mus musculus GN=Decr1 PE=1 SV=1 - [DECR_MOUSE] | 0,000E0 | 0,000E0 | 0,000E0 | 1,257E6 | 1,449E7 | 2,072E7 | Decr1 |
| P97473 | RISC-loading complex subunit TARBP2 OS=Mus musculus GN=Tarbp2 PE=1 SV=2 - [TRBP2_MOUSE] | 0,000E0 | 0,000E0 | 0,000E0 | 3,365E6 | 9,437E6 | 1,787E7 | Tarbp2 |
| Q78HU3 | Multivesicular body subunit 12A OS=Mus musculus GN=Mvb12a PE=1 SV=1 - [MB12A_MOUSE] | 0,000E0 | 0,000E0 | 0,000E0 | 8,758E6 | 1,521E7 | 2,166E7 | Mvb12a |
| Q9Z110-2 | Isoform Short of Delta-1-pyrroline-5-carboxylate synthase OS=Mus musculus GN=Aldh18a1 - [P5CS_MOUSE] | 0,000E0 | 0,000E0 | 0,000E0 | 5,133E6 | 7,406E6 | 6,006E6 | Aldh18a1 |
| Q62093 | Serine/arginine-rich splicing factor 2 OS=Mus musculus GN=Srsf2 PE=1 SV=4 - [SRSF2_MOUSE] | 0,000E0 | 0,000E0 | 0,000E0 | 5,248E7 | 1,203E8 | 1,941E8 | Srsf2 |
| Q99JB2 | Stomatin-like protein 2, mitochondrial OS=Mus musculus GN=Stoml2 PE=1 SV=1 - [STML2_MOUSE] | 0,000E0 | 0,000E0 | 0,000E0 | 4,798E6 | 3,895E6 | 7,812E6 | Stoml2 |
| Q99MI1 | ELKS/Rab6-interacting/CAST family member 1 OS=Mus musculus GN=Erc1 PE=1 SV=1 - [RB6I2_MOUSE] | 0,000E0 | 0,000E0 | 0,000E0 | 2,984E6 | 5,782E6 | 1,285E7 | Erc1 |
| P62334 | 26S protease regulatory subunit 10B OS=Mus musculus GN=Psmc6 PE=1 SV=1 - [PRS10_MOUSE] | 0,000E0 | 0,000E0 | 0,000E0 | 3,346E6 | 6,920E6 | 9,617E6 | Psmc6 |
| Q3UJV1 | Coiled-coil domain-containing protein 61 OS=Mus musculus GN=Ccdc61 PE=1 SV=1 - [CCD61_MOUSE] | 0,000E0 | 0,000E0 | 0,000E0 | 1,260E7 | 6,014E6 | 1,051E7 | Ccdc61 |
| P28798 | Granulins OS=Mus musculus GN=Grn PE=1 SV=2 - [GRN_MOUSE] | 0,000E0 | 0,000E0 | 3,374E6 | 3,539E7 | 2,659E7 | 1,343E7 | Grn |
| O08573-3 | Isoform 3 of Galectin-9 OS=Mus musculus GN=Lgals9 - [LEG9_MOUSE] | 0,000E0 | 0,000E0 | 0,000E0 | 4,338E7 | 8,750E7 | 1,411E8 | Lgals9 |
| Q6PDN3-2 | Isoform 2 of Myosin light chain kinase, smooth muscle OS=Mus musculus GN=Mylk - [MYLK_MOUSE] | 0,000E0 | 0,000E0 | 0,000E0 | 3,810E6 | 5,593E6 | 4,908E6 | Mylk |
| P11928 | 2'-5'-oligoadenylate synthase 1A OS=Mus musculus GN=Oas1a PE=1 SV=2 - [OAS1A_MOUSE] | 0,000E0 | 0,000E0 | 0,000E0 | 2,327E6 | 3,065E5 | 4,501E6 | Oas1a |
| P45376 | Aldose reductase OS=Mus musculus GN=Akr1b1 PE=1 SV=3 - [ALDR_MOUSE] | 0,000E0 | 0,000E0 | 0,000E0 | 4,119E6 | 4,037E6 | 2,748E6 | Akr1b1 |
| P62892 | 60S ribosomal protein L39 OS=Mus musculus GN=Rpl39 PE=1 SV=2 - [RL39_MOUSE] | 0,000E0 | 0,000E0 | 0,000E0 | 5,020E7 | 1,776E8 | 2,018E7 | Rpl39 |
| P18654 | Ribosomal protein S6 kinase alpha-3 OS=Mus musculus GN=Rps6ka3 PE=1 SV=2 - [KS6A3_MOUSE] | 0,000E0 | 0,000E0 | 0,000E0 | 8,356E6 | 8,702E6 | 9,966E6 | Rps6ka3 |
| Q9D0M3-2 | Isoform 2 of Cytochrome c1, heme protein, mitochondrial OS=Mus musculus GN=Cyc1 - [CY1_MOUSE] | 0,000E0 | 0,000E0 | 0,000E0 | 3,243E6 | 5,202E6 | 3,767E6 | Cyc1 |
| Q9JLQ2 | ARF GTPase-activating protein GIT2 OS=Mus musculus GN=Git2 PE=1 SV=2 - [GIT2_MOUSE] | 0,000E0 | 0,000E0 | 0,000E0 | 2,868E6 | 8,952E6 | 3,523E6 | Git2 |
| Q9DC53 | Copine-8 OS=Mus musculus GN=Cpne8 PE=2 SV=3 - [CPNE8_MOUSE] | 0,000E0 | 0,000E0 | 0,000E0 | 1,988E6 | 5,852E6 | 6,593E6 | Cpne8 |
| Q8K382 | DENN domain-containing protein 1A OS=Mus musculus GN=Dennd1a PE=1 SV=2 - [DEN1A_MOUSE] | 0,000E0 | 0,000E0 | 0,000E0 | 1,630E7 | 2,689E7 | 2,916E7 | Dennd1a |
| Q00547-2 | Isoform RHAMM1 of Hyaluronan mediated motility receptor OS=Mus musculus GN=Hmmr - [HMMR_MOUSE] | 0,000E0 | 0,000E0 | 0,000E0 | 4,188E6 | 5,031E6 | 1,273E7 | Hmmr |
| P26369 | Splicing factor U2AF 65 kDa subunit OS=Mus musculus GN=U2af2 PE=1 SV=3 - [U2AF2_MOUSE] | 0,000E0 | 0,000E0 | 0,000E0 | 3,688E6 | 1,219E7 | 2,900E7 | U2af2 |
| Q62433 | Protein NDRG1 OS=Mus musculus GN=Ndrg1 PE=1 SV=1 - [NDRG1_MOUSE] | 0,000E0 | 0,000E0 | 0,000E0 | 1,007E7 | 1,516E7 | 1,487E7 | Ndrg1 |
| P12382 | ATP-dependent 6-phosphofructokinase, liver type OS=Mus musculus GN=Pfkl PE=1 SV=4 - [PFKAL_MOUSE] | 0,000E0 | 0,000E0 | 0,000E0 | 7,220E6 | 1,677E7 | 9,026E6 | Pfkl |
| Q6PIU9 | Uncharacterized protein FLJ45252 homolog OS=Mus musculus PE=1 SV=2 - [YJ005_MOUSE] | 0,000E0 | 0,000E0 | 0,000E0 | 7,404E6 | 1,401E7 | 7,361E6 | YJ0051 |
| Q3UHD6-2 | Isoform 2 of Sorting nexin-27 OS=Mus musculus GN=Snx27 - [SNX27_MOUSE] | 0,000E0 | 0,000E0 | 0,000E0 | 3,634E6 | 7,773E6 | 1,341E7 | Snx27 |
| P61327 | Protein mago nashi homolog OS=Mus musculus GN=Magoh PE=2 SV=1 - [MGN_MOUSE] | 0,000E0 | 0,000E0 | 0,000E0 | 6,361E6 | 9,293E6 | 1,220E7 | Magoh |
| Q8VDM4 | 26S proteasome non-ATPase regulatory subunit 2 OS=Mus musculus GN=Psmd2 PE=1 SV=1 - [PSMD2_MOUSE] | 0,000E0 | 0,000E0 | 7,491E5 | 4,942E6 | 1,437E7 | 1,767E7 | Psmd2 |
| Q8BYC6 | Serine/threonine-protein kinase TAO3 OS=Mus musculus GN=Taok3 PE=1 SV=2 - [TAOK3_MOUSE] | 0,000E0 | 0,000E0 | 0,000E0 | 2,401E6 | 1,906E7 | 8,089E7 | Taok3 |
| Q8R1F0 | Leydig cell tumor 10 kDa protein homolog OS=Mus musculus GN=D8Ertd738e PE=3 SV=1 - [L10K_MOUSE] | 0,000E0 | 0,000E0 | 0,000E0 | 3,468E6 | 9,898E6 | 2,568E7 | D8Ertd738e |
| Q9DAW9 | Calponin-3 OS=Mus musculus GN=Cnn3 PE=1 SV=1 - [CNN3_MOUSE] | 0,000E0 | 0,000E0 | 0,000E0 | 1,946E6 | 4,101E6 | 2,096E7 | Cnn3 |
| Q9JHU4 | Cytoplasmic dynein 1 heavy chain 1 OS=Mus musculus GN=Dync1h1 PE=1 SV=2 - [DYHC1_MOUSE] | 0,000E0 | 0,000E0 | 2,944E6 | 5,116E6 | 7,967E6 | 1,125E7 | Dync1h1 |
| Q3UMT1 | Protein phosphatase 1 regulatory subunit 12C OS=Mus musculus GN=Ppp1r12c PE=1 SV=1 - [PP12C_MOUSE] | 0,000E0 | 0,000E0 | 0,000E0 | 4,169E6 | 7,718E6 | 7,432E6 | Ppp1r12c |
| Q569Z6 | Thyroid hormone receptor-associated protein 3 OS=Mus musculus GN=Thrap3 PE=1 SV=1 - [TR150_MOUSE] | 0,000E0 | 0,000E0 | 0,000E0 | 4,365E6 | 8,536E6 | 1,974E7 | Thrap3 |
| Q4VAA7 | Sorting nexin-33 OS=Mus musculus GN=Snx33 PE=1 SV=1 - [SNX33_MOUSE] | 0,000E0 | 0,000E0 | 0,000E0 | 9,825E6 | 1,609E7 | 3,495E7 | Snx33 |
| P70297 | Signal transducing adapter molecule 1 OS=Mus musculus GN=Stam PE=1 SV=3 - [STAM1_MOUSE] | 0,000E0 | 0,000E0 | 0,000E0 | 1,535E6 | 1,783E7 | 7,236E6 | Stam |
| O70133-2 | Isoform 2 of ATP-dependent RNA helicase A OS=Mus musculus GN=Dhx9 - [DHX9_MOUSE] | 0,000E0 | 0,000E0 | 0,000E0 | 1,772E6 | 4,093E6 | 1,239E7 | Dhx9 |
| Q8C147 | Dedicator of cytokinesis protein 8 OS=Mus musculus GN=Dock8 PE=1 SV=4 - [DOCK8_MOUSE] | 0,000E0 | 0,000E0 | 0,000E0 | 1,105E7 | 5,246E7 | 2,007E7 | Dock8 |
| O55143-2 | Isoform 2 of Sarcoplasmic/endoplasmic reticulum calcium ATPase 2 OS=Mus musculus GN=Atp2a2 - [AT2A2_MOUSE] | 0,000E0 | 0,000E0 | 0,000E0 | 7,180E6 | 6,207E6 | 2,255E7 | Atp2a2 |
| O88643 | Serine/threonine-protein kinase PAK 1 OS=Mus musculus GN=Pak1 PE=1 SV=1 - [PAK1_MOUSE] | 0,000E0 | 0,000E0 | 0,000E0 | 1,356E6 | 1,486E6 | 5,065E6 | Pak1 |
| Q9CWT6 | Probable ATP-dependent RNA helicase DDX28 OS=Mus musculus GN=Ddx28 PE=2 SV=2 - [DDX28_MOUSE] | 0,000E0 | 0,000E0 | 0,000E0 | 3,404E6 | 8,764E6 | 5,939E6 | Ddx28 |
| Q9EPL8 | Importin-7 OS=Mus musculus GN=Ipo7 PE=1 SV=2 - [IPO7_MOUSE] | 0,000E0 | 0,000E0 | 0,000E0 | 1,051E7 | 1,243E7 | 3,272E7 | Ipo7 |
| P12815 | Programmed cell death protein 6 OS=Mus musculus GN=Pdcd6 PE=1 SV=2 - [PDCD6_MOUSE] | 0,000E0 | 0,000E0 | 0,000E0 | 1,087E6 | 1,745E6 | 2,091E5 | Pdcd6 |
| Q9Z1Q9 | Valine--tRNA ligase OS=Mus musculus GN=Vars PE=1 SV=1 - [SYVC_MOUSE] | 0,000E0 | 0,000E0 | 0,000E0 | 1,242E7 | 2,355E7 | 4,246E7 | Vars |
| Q8BPU7 | Engulfment and cell motility protein 1 OS=Mus musculus GN=Elmo1 PE=1 SV=2 - [ELMO1_MOUSE] | 0,000E0 | 0,000E0 | 0,000E0 | 4,423E6 | 2,429E6 | 1,424E7 | Elmo1 |
| Q9Z1P7 | KN motif and ankyrin repeat domain-containing protein 3 OS=Mus musculus GN=Kank3 PE=1 SV=1 - [KANK3_MOUSE] | 0,000E0 | 0,000E0 | 0,000E0 | 7,252E6 | 8,195E6 | 2,766E7 | Kank3 |
| P52332 | Tyrosine-protein kinase JAK1 OS=Mus musculus GN=Jak1 PE=1 SV=1 - [JAK1_MOUSE] | 0,000E0 | 0,000E0 | 0,000E0 | 5,634E6 | 1,422E7 | 1,417E7 | Jak1 |
| Q99JX7 | Nuclear RNA export factor 1 OS=Mus musculus GN=Nxf1 PE=1 SV=3 - [NXF1_MOUSE] | 0,000E0 | 0,000E0 | 0,000E0 | 6,390E6 | 8,956E6 | 3,575E7 | Nxf1 |
| Q8CIE6 | Coatomer subunit alpha OS=Mus musculus GN=Copa PE=1 SV=2 - [COPA_MOUSE] | 0,000E0 | 0,000E0 | 0,000E0 | 7,174E6 | 1,016E7 | 1,350E7 | Copa |
| Q6A068 | Cell division cycle 5-like protein OS=Mus musculus GN=Cdc5l PE=1 SV=2 - [CDC5L_MOUSE] | 0,000E0 | 0,000E0 | 0,000E0 | 2,788E6 | 4,923E6 | 8,101E6 | Cdc5l |
| P24547 | Inosine-5'-monophosphate dehydrogenase 2 OS=Mus musculus GN=Impdh2 PE=1 SV=2 - [IMDH2_MOUSE] | 0,000E0 | 0,000E0 | 1,492E6 | 2,540E6 | 9,987E6 | 1,309E7 | Impdh2 |
| Q9JIF0-3 | Isoform 3 of Protein arginine N-methyltransferase 1 OS=Mus musculus GN=Prmt1 - [ANM1_MOUSE] | 0,000E0 | 0,000E0 | 0,000E0 | 2,328E6 | 1,527E6 | 8,366E6 | Prmt1 |
| Q810D6 | Glutamate-rich WD repeat-containing protein 1 OS=Mus musculus GN=Grwd1 PE=1 SV=2 - [GRWD1_MOUSE] | 0,000E0 | 0,000E0 | 0,000E0 | 2,322E6 | 6,257E6 | 1,152E7 | Grwd1 |
| O70310 | Glycylpeptide N-tetradecanoyltransferase 1 OS=Mus musculus GN=Nmt1 PE=1 SV=1 - [NMT1_MOUSE] | 0,000E0 | 0,000E0 | 0,000E0 | 5,233E6 | 1,449E7 | 1,426E7 | Nmt1 |
| Q9D0F4 | NF-kappa-B-activating protein OS=Mus musculus GN=Nkap PE=2 SV=1 - [NKAP_MOUSE] | 0,000E0 | 0,000E0 | 0,000E0 | 1,025E7 | 1,824E7 | 1,192E7 | Nkap |
| Q60668-4 | Isoform 4 of Heterogeneous nuclear ribonucleoprotein D0 OS=Mus musculus GN=Hnrnpd - [HNRPD_MOUSE] | 0,000E0 | 0,000E0 | 0,000E0 | 5,570E6 | 3,767E7 | 4,084E6 | Hnrnpd |
| Q8K045 | Serine/threonine-protein kinase N3 OS=Mus musculus GN=Pkn3 PE=1 SV=1 - [PKN3_MOUSE] | 0,000E0 | 0,000E0 | 0,000E0 | 2,128E6 | 5,524E6 | 3,647E6 | Pkn3 |
| Q3U1J4 | DNA damage-binding protein 1 OS=Mus musculus GN=Ddb1 PE=1 SV=2 - [DDB1_MOUSE] | 0,000E0 | 0,000E0 | 0,000E0 | 4,267E6 | 3,772E6 | 9,223E6 | Ddb1 |
| Q3UQ28 | Peroxidasin homolog OS=Mus musculus GN=Pxdn PE=1 SV=2 - [PXDN_MOUSE] | 0,000E0 | 0,000E0 | 0,000E0 | 9,280E6 | 1,365E7 | 2,619E7 | Pxdn |
| Q08288 | Cell growth-regulating nucleolar protein OS=Mus musculus GN=Lyar PE=1 SV=2 - [LYAR_MOUSE] | 0,000E0 | 0,000E0 | 0,000E0 | 2,423E6 | 3,167E6 | 4,356E6 | Lyar |
| Q4FK66 | Pre-mRNA-splicing factor 38A OS=Mus musculus GN=Prpf38a PE=1 SV=1 - [PR38A_MOUSE] | 0,000E0 | 0,000E0 | 0,000E0 | 2,106E6 | 1,084E7 | 3,121E7 | Prpf38a |
| Q07076 | Annexin A7 OS=Mus musculus GN=Anxa7 PE=1 SV=2 - [ANXA7_MOUSE] | 0,000E0 | 0,000E0 | 0,000E0 | 7,293E6 | 1,131E7 | 1,712E7 | Anxa7 |
| Q8CIW5 | Twinkle protein, mitochondrial OS=Mus musculus GN=Peo1 PE=2 SV=1 - [PEO1_MOUSE] | 0,000E0 | 0,000E0 | 0,000E0 | 1,675E6 | 2,124E6 | 6,791E6 | Peo1 |
| Q61074 | Protein phosphatase 1G OS=Mus musculus GN=Ppm1g PE=1 SV=3 - [PPM1G_MOUSE] | 0,000E0 | 0,000E0 | 0,000E0 | 1,792E6 | 4,290E6 | 2,796E6 | Ppm1g |
| Q5SW75 | Protein phosphatase Slingshot homolog 2 OS=Mus musculus GN=Ssh2 PE=1 SV=2 - [SSH2_MOUSE] | 0,000E0 | 0,000E0 | 0,000E0 | 1,604E6 | 2,964E6 | 7,512E6 | Ssh2 |
| Q01730 | Ras suppressor protein 1 OS=Mus musculus GN=Rsu1 PE=1 SV=3 - [RSU1_MOUSE] | 0,000E0 | 0,000E0 | 0,000E0 | 3,683E6 | 7,382E6 | 4,768E6 | Rsu1 |
| Q6IRU5-2 | Isoform 2 of Clathrin light chain B OS=Mus musculus GN=Cltb - [CLCB_MOUSE] | 0,000E0 | 0,000E0 | 0,000E0 | 2,662E7 | 5,010E7 | 8,397E7 | Cltb |
| Q9CS00 | Cactin OS=Mus musculus GN=Cactin PE=1 SV=2 - [CATIN_MOUSE] | 0,000E0 | 0,000E0 | 0,000E0 | 1,148E6 | 2,699E6 | 2,574E7 | Cactin |
| P97372 | Proteasome activator complex subunit 2 OS=Mus musculus GN=Psme2 PE=1 SV=4 - [PSME2_MOUSE] | 0,000E0 | 0,000E0 | 0,000E0 | 1,607E6 | 4,578E6 | 1,006E7 | Psme2 |
| Q61768 | Kinesin-1 heavy chain OS=Mus musculus GN=Kif5b PE=1 SV=3 - [KINH_MOUSE] | 0,000E0 | 0,000E0 | 0,000E0 | 3,844E6 | 7,054E6 | 7,687E6 | Kif5b |
| P68181 | cAMP-dependent protein kinase catalytic subunit beta OS=Mus musculus GN=Prkacb PE=1 SV=2 - [KAPCB_MOUSE] | 0,000E0 | 0,000E0 | 0,000E0 | 5,456E6 | 9,244E6 | 7,668E6 | Prkacb |

| | | | | | | | | |
|---|---|---|---|---|---|---|---|---|
| Q80TH2-1 | Isoform 1 of Erbin OS=Mus musculus GN=Erbin - [ERBIN_MOUSE] | 0,000E0 | 0,000E0 | 0,000E0 | 2,347E6 | 6,759E6 | 8,371E6 | Erbin |
| Q8VBT6 | Apolipoprotein B receptor OS=Mus musculus GN=Apobr PE=1 SV=1 - [APOBR_MOUSE] | 0,000E0 | 0,000E0 | 0,000E0 | 3,354E6 | 6,926E6 | 7,357E6 | Apobr |
| Q9CWJ9 | Bifunctional purine biosynthesis protein PURH OS=Mus musculus GN=Atic PE=1 SV=2 - [PUR9_MOUSE] | 0,000E0 | 0,000E0 | 0,000E0 | 2,409E6 | 6,303E6 | 4,578E6 | Atic |
| A2AQ19 | RNA polymerase-associated protein RTF1 homolog OS=Mus musculus GN=Rtf1 PE=1 SV=1 - [RTF1_MOUSE] | 0,000E0 | 0,000E0 | 0,000E0 | 5,034E6 | 3,960E6 | 1,019E7 | Rtf1 |
| P39688 | Tyrosine-protein kinase Fyn OS=Mus musculus GN=Fyn PE=1 SV=4 - [FYN_MOUSE] | 0,000E0 | 0,000E0 | 2,114E5 | 1,442E7 | 6,131E7 | 7,532E7 | Fyn |
| Q9CR62 | Mitochondrial 2-oxoglutarate/malate carrier protein OS=Mus musculus GN=Slc25a11 PE=1 SV=3 - [M2OM_MOUSE] | 0,000E0 | 0,000E0 | 0,000E0 | 6,059E6 | 1,770E7 | 1,029E7 | Slc25a11 |
| P63005 | Platelet-activating factor acetylhydrolase IB subunit alpha OS=Mus musculus GN=Pafah1b1 PE=1 SV=2 - [LIS1_MOUSE] | 0,000E0 | 0,000E0 | 0,000E0 | 8,818E6 | 6,403E6 | 2,972E7 | Pafah1b1 |
| Q9Z2X8 | Kelch-like ECH-associated protein 1 OS=Mus musculus GN=Keap1 PE=1 SV=1 - [KEAP1_MOUSE] | 0,000E0 | 0,000E0 | 0,000E0 | 2,604E6 | 6,488E6 | 1,298E7 | Keap1 |
| P97386-2 | Isoform Beta of DNA ligase 3 OS=Mus musculus GN=Lig3 - [DNLI3_MOUSE] | 0,000E0 | 0,000E0 | 0,000E0 | 3,950E6 | 2,136E6 | 1,202E7 | Lig3 |
| Q8VHK9 | ATP-dependent RNA helicase DHX36 OS=Mus musculus GN=Dhx36 PE=1 SV=2 - [DHX36_MOUSE] | 0,000E0 | 0,000E0 | 0,000E0 | 6,937E6 | 7,944E6 | 1,198E7 | Dhx36 |
| Q61846 | Maternal embryonic leucine zipper kinase OS=Mus musculus GN=Melk PE=1 SV=2 - [MELK_MOUSE] | 0,000E0 | 0,000E0 | 0,000E0 | 1,905E6 | 2,942E6 | 6,697E6 | Melk |
| Q8CHH9 | Septin-8 OS=Mus musculus GN=Sept8 PE=1 SV=4 - [SEPT8_MOUSE] | 0,000E0 | 0,000E0 | 0,000E0 | 1,802E7 | 3,045E7 | 1,040E7 | SEPT18 |
| Q80UP5 | Ankyrin repeat domain-containing protein 13A OS=Mus musculus GN=Ankrd13a PE=1 SV=2 - [AN13A_MOUSE] | 0,000E0 | 0,000E0 | 0,000E0 | 2,476E6 | 1,354E6 | 2,324E6 | Ankrd13a |
| Q8R570 | Synaptosomal-associated protein 47 OS=Mus musculus GN=Snap47 PE=1 SV=1 - [SNP47_MOUSE] | 0,000E0 | 0,000E0 | 0,000E0 | 9,854E5 | 4,840E6 | 6,747E6 | Snap47 |
| P50516 | V-type proton ATPase catalytic subunit A OS=Mus musculus GN=Atp6v1a PE=1 SV=2 - [VATA_MOUSE] | 0,000E0 | 0,000E0 | 0,000E0 | 2,553E6 | 1,379E6 | 5,067E6 | Atp6v1a |
| Q69ZB0-3 | Isoform 3 of Leucine-rich repeat and coiled-coil domain-containing protein 1 OS=Mus musculus GN=Lrrcc1 - [LRCC1_MOUSE] | 0,000E0 | 0,000E0 | 0,000E0 | 2,347E6 | 4,735E6 | 3,274E6 | Lrrcc1 |
| Q99KK2 | N-acylneuraminate cytidylyltransferase OS=Mus musculus GN=Cmas PE=1 SV=2 - [NEUA_MOUSE] | 0,000E0 | 0,000E0 | 0,000E0 | 1,756E6 | 5,798E6 | 7,022E6 | Cmas |
| Q62172 | RalA-binding protein 1 OS=Mus musculus GN=Ralbp1 PE=1 SV=4 - [RBP1_MOUSE] | 0,000E0 | 0,000E0 | 0,000E0 | 2,565E7 | 1,146E7 | 1,415E7 | Ralbp1 |
| B2RQC6 | CAD protein OS=Mus musculus GN=Cad PE=1 SV=1 - [PYR1_MOUSE] | 0,000E0 | 0,000E0 | 0,000E0 | 5,483E6 | 1,615E7 | 1,448E7 | Cad |
| Q1HFZ0 | tRNA (cytosine(34)-C(5))-methyltransferase OS=Mus musculus GN=Nsun2 PE=1 SV=2 - [NSUN2_MOUSE] | 0,000E0 | 0,000E0 | 0,000E0 | 2,205E6 | 2,475E6 | 7,144E6 | Nsun2 |
| Q68FH0 | Plakophilin-4 OS=Mus musculus GN=Pkp4 PE=1 SV=1 - [PKP4_MOUSE] | 0,000E0 | 0,000E0 | 0,000E0 | 2,776E6 | 7,408E6 | 5,875E6 | Pkp4 |
| Q3TGF2 | Protein FAM107B OS=Mus musculus GN=Fam107b PE=1 SV=2 - [F107B_MOUSE] | 0,000E0 | 0,000E0 | 0,000E0 | 2,030E6 | 9,268E6 | 1,621E7 | Fam107b |
| Q8BMB3 | Eukaryotic translation initiation factor 4E type 2 OS=Mus musculus GN=Eif4e2 PE=1 SV=1 - [IF4E2_MOUSE] | 0,000E0 | 0,000E0 | 0,000E0 | 8,189E6 | 6,987E6 | 3,439E7 | Eif4e2 |
| Q80XI4 | Phosphatidylinositol 5-phosphate 4-kinase type-2 beta OS=Mus musculus GN=Pip4k2b PE=1 SV=1 - [PI42B_MOUSE] | 0,000E0 | 0,000E0 | 0,000E0 | 2,082E6 | 3,995E6 | 1,697E7 | Pip4k2b |
| Q9D8W5 | 26S proteasome non-ATPase regulatory subunit 12 OS=Mus musculus GN=Psmd12 PE=1 SV=4 - [PSD12_MOUSE] | 0,000E0 | 0,000E0 | 0,000E0 | 1,880E6 | 5,001E6 | 3,952E6 | Psmd12 |
| Q8R2Q8 | Bone marrow stromal antigen 2 OS=Mus musculus GN=Bst2 PE=1 SV=1 - [BST2_MOUSE] | 0,000E0 | 0,000E0 | 0,000E0 | 4,800E7 | 1,182E8 | 3,273E8 | Bst2 |
| Q923D4 | Splicing factor 3B subunit 5 OS=Mus musculus GN=Sf3b5 PE=1 SV=1 - [SF3B5_MOUSE] | 0,000E0 | 0,000E0 | 0,000E0 | 3,545E6 | 8,978E6 | 1,730E7 | Sf3b5 |
| P97465 | Docking protein 1 OS=Mus musculus GN=Dok1 PE=1 SV=2 - [DOK1_MOUSE] | 0,000E0 | 0,000E0 | 0,000E0 | 3,292E6 | 6,291E6 | 1,275E7 | Dok1 |
| P28656 | Nucleosome assembly protein 1-like 1 OS=Mus musculus GN=Nap1l1 PE=1 SV=2 - [NP1L1_MOUSE] | 0,000E0 | 0,000E0 | 0,000E0 | 8,986E6 | 3,494E7 | 3,736E7 | Nap1l1 |
| Q9QX11-2 | Isoform 2 of Cytohesin-1 OS=Mus musculus GN=Cyth1 - [CYH1_MOUSE] | 0,000E0 | 0,000E0 | 0,000E0 | 4,744E6 | 2,455E7 | 4,210E6 | Cyth1 |
| Q64511 | DNA topoisomerase 2-beta OS=Mus musculus GN=Top2b PE=1 SV=2 - [TOP2B_MOUSE] | 0,000E0 | 0,000E0 | 0,000E0 | 1,330E7 | 4,832E7 | 9,462E7 | Top2b |
| P61164 | Alpha-centractin OS=Mus musculus GN=Actr1a PE=1 SV=1 - [ACTZ_MOUSE] | 0,000E0 | 0,000E0 | 0,000E0 | 5,285E6 | 3,488E7 | 2,330E7 | Actr1a |
| P17257 | Protein FAM167B OS=Mus musculus GN=Fam167b PE=2 SV=2 - [F167B_MOUSE] | 0,000E0 | 0,000E0 | 0,000E0 | 5,233E6 | 1,353E7 | 9,100E6 | Fam167b |
| Q9EST5 | Acidic leucine-rich nuclear phosphoprotein 32 family member B OS=Mus musculus GN=Anp32b PE=1 SV=1 - [AN32B_MOUSE] | 0,000E0 | 0,000E0 | 0,000E0 | 3,187E6 | 6,358E6 | 3,465E6 | Anp32b |
| P35235 | Tyrosine-protein phosphatase non-receptor type 11 OS=Mus musculus GN=Ptpn11 PE=1 SV=2 - [PTN11_MOUSE] | 0,000E0 | 0,000E0 | 0,000E0 | 2,902E6 | 3,135E6 | 5,808E6 | Ptpn11 |
| Q78IK2 | Up-regulated during skeletal muscle growth protein 5 OS=Mus musculus GN=Usmg5 PE=1 SV=1 - [USMG5_MOUSE] | 0,000E0 | 0,000E0 | 0,000E0 | 3,272E6 | 4,998E6 | 1,352E7 | Usmg5 |
| Q8VEM8 | Phosphate carrier protein, mitochondrial OS=Mus musculus GN=Slc25a3 PE=1 SV=1 - [MPCP_MOUSE] | 0,000E0 | 0,000E0 | 0,000E0 | 2,277E7 | 3,369E7 | 5,978E7 | Slc25a3 |
| Q9R0N0 | Galactokinase OS=Mus musculus GN=Galk1 PE=1 SV=2 - [GALK1_MOUSE] | 0,000E0 | 0,000E0 | 0,000E0 | 8,671E6 | 8,518E6 | 1,557E7 | Galk1 |
| Q99NH2-3 | Isoform 3 of Partitioning defective 3 homolog OS=Mus musculus GN=Pard3 - [PARD3_MOUSE] | 0,000E0 | 0,000E0 | 0,000E0 | 3,627E6 | 7,534E6 | 9,230E6 | Pard3 |
| Q6RHR9 | Membrane-associated guanylate kinase, WW and PDZ domain-containing protein 1 OS=Mus musculus GN=Magi1 PE=1 SV=1 - [MAGI1_MOUSE] | 0,000E0 | 0,000E0 | 0,000E0 | 9,352E6 | 2,147E7 | 3,627E7 | Magi1 |
| Q9JLV5 | Cullin-3 OS=Mus musculus GN=Cul3 PE=1 SV=1 - [CUL3_MOUSE] | 0,000E0 | 0,000E0 | 0,000E0 | 5,636E5 | 4,144E6 | 5,389E6 | Cul3 |
| Q64324 | Syntaxin-binding protein 2 OS=Mus musculus GN=Stxbp2 PE=1 SV=1 - [STXB2_MOUSE] | 0,000E0 | 0,000E0 | 0,000E0 | 4,569E5 | 4,727E6 | 5,283E6 | Stxbp2 |
| Q9QZ88 | Vacuolar protein sorting-associated protein 29 OS=Mus musculus GN=Vps29 PE=1 SV=1 - [VPS29_MOUSE] | 0,000E0 | 0,000E0 | 0,000E0 | 4,303E6 | 4,444E6 | 8,343E6 | Vps29 |
| Q9CYH6 | Ribosome biogenesis regulatory protein homolog OS=Mus musculus GN=Rrs1 PE=1 SV=1 - [RRS1_MOUSE] | 0,000E0 | 0,000E0 | 0,000E0 | 7,456E5 | 2,835E6 | 3,820E6 | Rrs1 |
| Q9WV02-2 | Isoform 2 of RNA-binding motif protein, X chromosome OS=Mus musculus GN=Rbmx - [RBMX_MOUSE] | 0,000E0 | 0,000E0 | 0,000E0 | 4,958E6 | 3,403E6 | 2,397E7 | Rbmx |
| Q924C6 | Lysyl oxidase homolog 4 OS=Mus musculus GN=Loxl4 PE=2 SV=2 - [LOXL4_MOUSE] | 0,000E0 | 0,000E0 | 0,000E0 | 7,417E6 | 1,448E7 | 3,724E7 | Loxl4 |
| P15379-2 | Isoform 13 of CD44 antigen OS=Mus musculus GN=Cd44 - [CD44_MOUSE] | 0,000E0 | 0,000E0 | 0,000E0 | 3,270E6 | 1,960E7 | 4,612E7 | Cd44 |
| Q9CQV6 | Microtubule-associated proteins 1A/1B light chain 3B OS=Mus musculus GN=Map1lc3b PE=1 SV=3 - [MLP3B_MOUSE] | 0,000E0 | 0,000E0 | 0,000E0 | 9,970E6 | 2,051E7 | 2,933E7 | Map1lc3b |
| Q9CZH8-2 | Isoform 2 of Coiled-coil domain-containing protein 77 OS=Mus musculus GN=Ccdc77 - [CCD77_MOUSE] | 0,000E0 | 0,000E0 | 0,000E0 | 2,014E6 | 8,513E6 | 5,174E6 | Ccdc77 |
| Q91WB7 | Ubiquitin domain-containing protein 1 OS=Mus musculus GN=Ubtd1 PE=2 SV=1 - [UBTD1_MOUSE] | 0,000E0 | 0,000E0 | 0,000E0 | 1,787E6 | 9,254E5 | 2,288E6 | Ubtd1 |
| Q99NB9 | Splicing factor 3B subunit 1 OS=Mus musculus GN=Sf3b1 PE=1 SV=1 - [SF3B1_MOUSE] | 0,000E0 | 0,000E0 | 0,000E0 | 2,034E6 | 1,116E7 | 1,570E7 | Sf3b1 |
| Q60848-2 | Isoform 2 of Lymphocyte-specific helicase OS=Mus musculus GN=Hells - [HELLS_MOUSE] | 0,000E0 | 0,000E0 | 0,000E0 | 2,855E6 | 4,705E6 | 8,351E6 | Hells |
| O54946 | DnaJ homolog subfamily B member 6 OS=Mus musculus GN=Dnajb6 PE=1 SV=4 - [DNJB6_MOUSE] | 0,000E0 | 0,000E0 | 0,000E0 | 9,339E6 | 9,220E6 | 8,046E6 | Dnajb6 |
| Q8K2X3 | CST complex subunit STN1 OS=Mus musculus GN=Stn1 PE=1 SV=2 - [STN1_MOUSE] | 0,000E0 | 0,000E0 | 0,000E0 | 4,694E6 | 4,647E6 | 5,395E6 | Stn1 |
| A2RSY6 | TRMT1-like protein OS=Mus musculus GN=Trmt1l PE=1 SV=1 - [TRM1L_MOUSE] | 0,000E0 | 0,000E0 | 0,000E0 | 5,755E6 | 7,540E6 | 1,548E7 | Trmt1l |
| Q3TKT4 | Transcription activator BRG1 OS=Mus musculus GN=Smarca4 PE=1 SV=1 - [SMCA4_MOUSE] | 0,000E0 | 0,000E0 | 0,000E0 | 5,388E6 | 4,917E6 | 6,777E6 | Smarca4 |
| P97432 | Next to BRCA1 gene 1 protein OS=Mus musculus GN=Nbr1 PE=1 SV=1 - [NBR1_MOUSE] | 0,000E0 | 0,000E0 | 0,000E0 | 1,301E7 | 2,664E7 | 2,224E7 | Nbr1 |
| Q91V92 | ATP-citrate synthase OS=Mus musculus GN=Acly PE=1 SV=1 - [ACLY_MOUSE] | 0,000E0 | 0,000E0 | 0,000E0 | 3,734E6 | 7,970E6 | 1,037E7 | Acly |
| Q9ET26 | E3 ubiquitin-protein ligase RNF114 OS=Mus musculus GN=Rnf114 PE=1 SV=2 - [RN114_MOUSE] | 0,000E0 | 0,000E0 | 0,000E0 | 1,559E6 | 3,166E6 | 4,096E6 | Rnf114 |
| Q62312-2 | Isoform RII-1 of TGF-beta receptor type-2 OS=Mus musculus GN=Tgfbr2 - [TGFR2_MOUSE] | 0,000E0 | 0,000E0 | 0,000E0 | 3,425E6 | 9,760E6 | 1,333E7 | Tgfbr2 |
| O08788-2 | Isoform 2 of Dynactin subunit 1 OS=Mus musculus GN=Dctn1 - [DCTN1_MOUSE] | 0,000E0 | 0,000E0 | 0,000E0 | 1,861E6 | 1,100E7 | 6,943E6 | Dctn1 |
| Q8C0C7 | Phenylalanine--tRNA ligase alpha subunit OS=Mus musculus GN=Farsa PE=1 SV=1 - [SYFA_MOUSE] | 0,000E0 | 0,000E0 | 0,000E0 | 2,208E6 | 4,154E6 | 8,435E6 | Farsa |
| Q9Z2D1 | Myotubularin-related protein 2 OS=Mus musculus GN=Mtmr2 PE=1 SV=3 - [MTMR2_MOUSE] | 0,000E0 | 0,000E0 | 0,000E0 | 8,915E5 | 4,390E6 | 6,080E6 | Mtmr2 |
| Q9CWX9 | Probable ATP-dependent RNA helicase DDX47 OS=Mus musculus GN=Ddx47 PE=2 SV=2 - [DDX47_MOUSE] | 0,000E0 | 0,000E0 | 0,000E0 | 3,773E6 | 4,408E6 | 7,407E6 | Ddx47 |
| Q9Z2B9 | Ribosomal protein S6 kinase alpha-4 OS=Mus musculus GN=Rps6ka4 PE=1 SV=2 - [KS6A4_MOUSE] | 0,000E0 | 0,000E0 | 0,000E0 | 2,213E6 | 4,829E6 | 4,433E6 | Rps6ka4 |
| Q9CQU1 | Microfibrillar-associated protein 1 OS=Mus musculus GN=Mfap1 PE=1 SV=1 - [MFAP1_MOUSE] | 0,000E0 | 0,000E0 | 0,000E0 | 6,425E5 | 4,287E6 | 3,922E6 | Mfap1 |
| Q08943 | FACT complex subunit SSRP1 OS=Mus musculus GN=Ssrp1 PE=1 SV=2 - [SSRP1_MOUSE] | 0,000E0 | 0,000E0 | 0,000E0 | 1,363E6 | 2,438E6 | 2,111E7 | Ssrp1 |
| P31324 | cAMP-dependent protein kinase type II-beta regulatory subunit OS=Mus musculus GN=Prkar2b PE=1 SV=3 - [KAP3_MOUSE] | 0,000E0 | 0,000E0 | 0,000E0 | 2,595E6 | 3,574E6 | 9,664E5 | Prkar2b |
| Q9DBE9 | pre-rRNA processing protein FTSJ3 OS=Mus musculus GN=Ftsj3 PE=1 SV=1 - [SPB1_MOUSE] | 0,000E0 | 0,000E0 | 0,000E0 | 1,958E6 | 3,378E6 | 9,315E6 | Ftsj3 |
| P62996 | Transformer-2 protein homolog beta OS=Mus musculus GN=Tra2b PE=1 SV=1 - [TRA2B_MOUSE] | 0,000E0 | 0,000E0 | 0,000E0 | 1,430E7 | 1,984E7 | 1,334E7 | Tra2b |
| Q80UM7 | Mannosyl-oligosaccharide glucosidase OS=Mus musculus GN=Mogs PE=1 SV=1 - [MOGS_MOUSE] | 0,000E0 | 0,000E0 | 0,000E0 | 2,944E6 | 1,768E6 | 5,937E6 | Mogs |
| P61021 | Ras-related protein Rab-5B OS=Mus musculus GN=Rab5b PE=1 SV=1 - [RAB5B_MOUSE] | 0,000E0 | 0,000E0 | 0,000E0 | 1,294E6 | 2,563E7 | 5,475E7 | Rab5b |
| Q91X43 | SH3 domain-containing protein 19 OS=Mus musculus GN=Sh3d19 PE=1 SV=2 - [SH319_MOUSE] | 0,000E0 | 0,000E0 | 0,000E0 | 2,247E6 | 6,127E6 | 3,582E6 | Sh3d19 |
| P28658 | Ataxin-10 OS=Mus musculus GN=Atxn10 PE=1 SV=2 - [ATX10_MOUSE] | 0,000E0 | 0,000E0 | 0,000E0 | 3,189E6 | 2,810E6 | 1,759E6 | Atxn10 |
| Q922W5 | Pyrroline-5-carboxylate reductase 1, mitochondrial OS=Mus musculus GN=Pycr1 PE=1 SV=1 - [P5CR1_MOUSE] | 0,000E0 | 0,000E0 | 0,000E0 | 3,839E6 | 8,170E6 | 6,841E7 | Pycr1 |
| Q9CWZ3-2 | Isoform 2 of RNA-binding protein 8A OS=Mus musculus GN=Rbm8a - [RBM8A_MOUSE] | 0,000E0 | 0,000E0 | 0,000E0 | 1,669E6 | 6,538E6 | 5,105E6 | Rbm8a |
| Q68FG3 | Protein SPT2 homolog OS=Mus musculus GN=Spty2d1 PE=2 SV=1 - [SPT2_MOUSE] | 0,000E0 | 0,000E0 | 0,000E0 | 2,451E6 | 1,018E7 | 9,001E6 | Spty2d1 |
| Q9JKX6 | ADP-sugar pyrophosphatase OS=Mus musculus GN=Nudt5 PE=1 SV=1 - [NUDT5_MOUSE] | 0,000E0 | 0,000E0 | 0,000E0 | 6,197E6 | 4,715E6 | 7,761E6 | Nudt5 |
| Q9D9Z1 | Small kinetochore-associated protein OS=Mus musculus GN=Knstrn PE=2 SV=2 - [SKAP_MOUSE] | 0,000E0 | 0,000E0 | 0,000E0 | 3,112E6 | 7,991E6 | 8,356E6 | Knstrn |
| B2RRE7 | OTU domain-containing protein 4 OS=Mus musculus GN=Otud4 PE=1 SV=1 - [OTUD4_MOUSE] | 0,000E0 | 0,000E0 | 0,000E0 | 7,442E6 | 6,910E6 | 1,338E7 | Otud4 |
| Q9D2Q3 | Uncharacterized protein C10orf88 homolog OS=Mus musculus PE=2 SV=1 - [CJ088_MOUSE] | 0,000E0 | 0,000E0 | 0,000E0 | 3,169E6 | 5,142E6 | 3,502E6 | SKT |
| A2AQ25-8 | Isoform 8 of Sickle tail protein OS=Mus musculus GN=Skt - [SKT_MOUSE] | 0,000E0 | 0,000E0 | 0,000E0 | 1,650E6 | 5,131E6 | 2,511E6 | Skt |
| Q7TMF2 | 3'-5' exoribonuclease 1 OS=Mus musculus GN=Eri1 PE=1 SV=2 - [ERI1_MOUSE] | 0,000E0 | 0,000E0 | 0,000E0 | 1,411E6 | 7,569E6 | 8,443E6 | Eri1 |
| Q8BG67 | Protein EFR3 homolog A OS=Mus musculus GN=Efr3a PE=1 SV=1 - [EFR3A_MOUSE] | 0,000E0 | 0,000E0 | 0,000E0 | 3,915E6 | 1,151E7 | 1,620E7 | Efr3a |
| Q62415 | Apoptosis-stimulating of p53 protein 1 OS=Mus musculus GN=Ppp1r13b PE=1 SV=2 - [ASPP1_MOUSE] | 0,000E0 | 0,000E0 | 0,000E0 | 7,092E6 | 1,296E7 | 2,250E7 | Ppp1r13b |
| P35441 | Thrombospondin-1 OS=Mus musculus GN=Thbs1 PE=1 SV=1 - [TSP1_MOUSE] | 0,000E0 | 0,000E0 | 0,000E0 | 8,753E6 | 1,171E7 | 2,563E7 | Thbs1 |
| Q60749 | KH domain-containing, RNA-binding, signal transduction-associated protein 1 OS=Mus musculus GN=Khdrbs1 PE=1 SV=2 - [KHDR1_MOUSE] | 0,000E0 | 0,000E0 | 0,000E0 | 1,327E7 | 1,459E7 | 3,045E7 | Khdrbs1 |
| Q91W50 | Cold shock domain-containing protein E1 OS=Mus musculus GN=Csde1 PE=1 SV=1 - [CSDE1_MOUSE] | 0,000E0 | 0,000E0 | 0,000E0 | 3,768E6 | 7,555E6 | 1,671E7 | Csde1 |
| Q9CR67 | Transmembrane protein 33 OS=Mus musculus GN=Tmem33 PE=1 SV=1 - [TMM33_MOUSE] | 0,000E0 | 0,000E0 | 0,000E0 | 1,557E6 | 1,401E6 | 7,004E6 | Tmem33 |
| Q62523 | Zyxin OS=Mus musculus GN=Zyx PE=1 SV=2 - [ZYX_MOUSE] | 0,000E0 | 0,000E0 | 0,000E0 | 5,576E6 | 1,209E7 | 1,979E7 | Zyx |
| Q03958 | Prefoldin subunit 6 OS=Mus musculus GN=Pfdn6 PE=1 SV=1 - [PFD6_MOUSE] | 0,000E0 | 0,000E0 | 0,000E0 | 1,502E6 | 2,888E6 | 4,143E6 | Pfdn6 |
| P54822 | Adenylosuccinate lyase OS=Mus musculus GN=Adsl PE=1 SV=2 - [PUR8_MOUSE] | 0,000E0 | 0,000E0 | 0,000E0 | 1,150E6 | 2,581E6 | 5,568E6 | Adsl |
| Q64213-2 | Isoform CW17E of Splicing factor 1 OS=Mus musculus GN=Sf1 - [SF01_MOUSE] | 0,000E0 | 0,000E0 | 0,000E0 | 4,830E6 | 6,298E6 | 3,930E7 | Sf1 |

| | | | | | | | | |
|---|---|---|---|---|---|---|---|---|
| Q80VD1 | Protein FAM98B OS=Mus musculus GN=Fam98b PE=1 SV=1 - [FA98B_MOUSE] | 0,000E0 | 0,000E0 | 0,000E0 | 1,435E6 | 5,079E6 | 2,171E6 | Fam98b |
| Q9WUA2 | Phenylalanine--tRNA ligase beta subunit OS=Mus musculus GN=Farsb PE=1 SV=2 - [SYFB_MOUSE] | 0,000E0 | 0,000E0 | 0,000E0 | 3,032E6 | 3,172E6 | 1,251E7 | Farsb |
| Q3TZZ7 | Extended synaptotagmin-2 OS=Mus musculus GN=Esyt2 PE=1 SV=1 - [ESYT2_MOUSE] | 0,000E0 | 0,000E0 | 0,000E0 | 4,744E6 | 3,524E6 | 7,689E6 | Esyt2 |
| Q6NVF9 | Cleavage and polyadenylation specificity factor subunit 6 OS=Mus musculus GN=Cpsf6 PE=1 SV=1 - [CPSF6_MOUSE] | 0,000E0 | 0,000E0 | 0,000E0 | 5,040E6 | 1,384E7 | 1,272E7 | Cpsf6 |
| P40336 | Vacuolar protein sorting-associated protein 26A OS=Mus musculus GN=Vps26a PE=1 SV=1 - [VP26A_MOUSE] | 0,000E0 | 0,000E0 | 0,000E0 | 1,322E6 | 3,810E6 | 5,425E6 | Vps26a |
| Q99M28-2 | Isoform 2 of RNA-binding protein with serine-rich domain 1 OS=Mus musculus GN=Rnps1 - [RNPS1_MOUSE] | 0,000E0 | 0,000E0 | 0,000E0 | 2,831E6 | 1,237E7 | 1,851E7 | Rnps1 |
| P27048 | Small nuclear ribonucleoprotein-associated protein B OS=Mus musculus GN=Snrpb PE=1 SV=1 - [RSMB_MOUSE] | 0,000E0 | 0,000E0 | 0,000E0 | 2,612E6 | 8,608E6 | 1,963E7 | Snrpb |
| Q80YR4-2 | Isoform 2 of Zinc finger protein 598 OS=Mus musculus GN=Znf598 - [ZN598_MOUSE] | 0,000E0 | 0,000E0 | 0,000E0 | 7,897E5 | 4,384E6 | 9,729E6 | Znf598 |
| Q03145 | Ephrin type-A receptor 2 OS=Mus musculus GN=Epha2 PE=1 SV=3 - [EPHA2_MOUSE] | 0,000E0 | 0,000E0 | 0,000E0 | 9,608E6 | 2,018E7 | 4,015E7 | Epha2 |
| E9Q3L2 | Phosphatidylinositol 4-kinase alpha OS=Mus musculus GN=Pi4ka PE=1 SV=2 - [PI4KA_MOUSE] | 0,000E0 | 0,000E0 | 0,000E0 | 3,502E6 | 1,026E7 | 2,568E7 | Pi4ka |
| Q80VI1 | E3 ubiquitin-protein ligase TRIM56 OS=Mus musculus GN=Trim56 PE=1 SV=1 - [TRI56_MOUSE] | 0,000E0 | 0,000E0 | 0,000E0 | 4,133E6 | 4,301E6 | 1,038E7 | Trim56 |
| Q5SUF2 | Luc7-like protein 3 OS=Mus musculus GN=Luc7l3 PE=1 SV=1 - [LC7L3_MOUSE] | 0,000E0 | 0,000E0 | 0,000E0 | 1,066E7 | 1,508E7 | 2,945E7 | Luc7l3 |
| Q8BGU5-2 | Isoform 2 of Cyclin-Y OS=Mus musculus GN=Ccny - [CCNY_MOUSE] | 0,000E0 | 0,000E0 | 0,000E0 | 1,658E6 | 8,696E6 | 1,143E7 | Ccny |
| Q6ZPF4 | Formin-like protein 3 OS=Mus musculus GN=Fmnl3 PE=1 SV=2 - [FMNL3_MOUSE] | 0,000E0 | 0,000E0 | 0,000E0 | 5,625E6 | 6,977E6 | 1,567E7 | Fmnl3 |
| Q8CH77 | Neuron navigator 1 OS=Mus musculus GN=Nav1 PE=1 SV=2 - [NAV1_MOUSE] | 0,000E0 | 0,000E0 | 0,000E0 | 3,656E6 | 6,640E6 | 1,178E7 | Nav1 |
| Q922R8 | Protein disulfide-isomerase A6 OS=Mus musculus GN=Pdia6 PE=1 SV=3 - [PDIA6_MOUSE] | 0,000E0 | 0,000E0 | 0,000E0 | 4,343E6 | 1,020E7 | 1,314E7 | Pdia6 |
| P28828 | Receptor-type tyrosine-protein phosphatase mu OS=Mus musculus GN=Ptprm PE=1 SV=2 - [PTPRM_MOUSE] | 0,000E0 | 0,000E0 | 0,000E0 | 1,893E7 | 2,155E7 | 5,201E7 | Ptprm |
| Q8BTW9 | Serine/threonine-protein kinase PAK 4 OS=Mus musculus GN=Pak4 PE=1 SV=1 - [PAK4_MOUSE] | 0,000E0 | 0,000E0 | 0,000E0 | 3,970E6 | 8,193E6 | 4,169E6 | Pak4 |
| Q8BWT5 | Disco-interacting protein 2 homolog A OS=Mus musculus GN=Dip2a PE=1 SV=3 - [DIP2A_MOUSE] | 0,000E0 | 0,000E0 | 0,000E0 | 6,147E6 | 1,679E7 | 2,390E7 | Dip2a |
| E9Q6P5 | Tetratricopeptide repeat protein 7B OS=Mus musculus GN=Ttc7b PE=1 SV=1 - [TTC7B_MOUSE] | 0,000E0 | 0,000E0 | 0,000E0 | 1,440E6 | 5,079E6 | 3,417E6 | Ttc7b |
| Q60973 | Histone-binding protein RBBP7 OS=Mus musculus GN=Rbbp7 PE=1 SV=1 - [RBBP7_MOUSE] | 0,000E0 | 0,000E0 | 0,000E0 | 5,163E6 | 8,607E6 | 8,565E6 | Rbbp7 |
| Q8BHL3 | TBC1 domain family member 10B OS=Mus musculus GN=Tbc1d10b PE=1 SV=2 - [TB10B_MOUSE] | 0,000E0 | 0,000E0 | 0,000E0 | 3,973E6 | 3,281E6 | 6,240E6 | Tbc1d10b |
| Q8VDM6-2 | Isoform 2 of Heterogeneous nuclear ribonucleoprotein U-like protein 1 OS=Mus musculus GN=Hnrnpul1 - [HNRL1_MOUSE] | 0,000E0 | 0,000E0 | 0,000E0 | 3,663E6 | 1,010E7 | 1,069E7 | Hnrnpul1 |
| Q8C854-2 | Isoform 4 of Myelin expression factor 2 OS=Mus musculus GN=Myef2 - [MYEF2_MOUSE] | 0,000E0 | 0,000E0 | 0,000E0 | 2,791E6 | 5,703E6 | 5,880E6 | Myef2 |
| Q3UFK8 | FERM domain-containing protein 8 OS=Mus musculus GN=Frmd8 PE=1 SV=2 - [FRMD8_MOUSE] | 0,000E0 | 0,000E0 | 0,000E0 | 1,696E6 | 6,759E6 | 8,161E6 | Frmd8 |
| Q5EG47 | 5'-AMP-activated protein kinase catalytic subunit alpha-1 OS=Mus musculus GN=Prkaa1 PE=1 SV=2 - [AAPK1_MOUSE] | 0,000E0 | 0,000E0 | 0,000E0 | 3,574E6 | 6,432E6 | 8,643E6 | Prkaa1 |
| Q78ZA7 | Nucleosome assembly protein 1-like 4 OS=Mus musculus GN=Nap1l4 PE=1 SV=1 - [NP1L4_MOUSE] | 0,000E0 | 0,000E0 | 0,000E0 | 2,814E6 | 7,931E6 | 9,935E6 | Nap1l4 |
| Q8CJ53-4 | Isoform 4 of Cdc42-interacting protein 4 OS=Mus musculus GN=Trip10 - [CIP4_MOUSE] | 0,000E0 | 0,000E0 | 0,000E0 | 2,371E6 | 5,979E6 | 8,934E6 | Trip10 |
| Q9D1E6 | Tubulin-folding cofactor B OS=Mus musculus GN=Tbcb PE=1 SV=2 - [TBCB_MOUSE] | 0,000E0 | 0,000E0 | 0,000E0 | 1,996E6 | 5,520E6 | 1,107E7 | Tbcb |
| Q91V64 | Isochorismatase domain-containing protein 1 OS=Mus musculus GN=Isoc1 PE=1 SV=1 - [ISOC1_MOUSE] | 0,000E0 | 0,000E0 | 0,000E0 | 6,309E5 | 9,475E5 | 1,217E7 | Isoc1 |
| Q9CZ30 | Obg-like ATPase 1 OS=Mus musculus GN=Ola1 PE=1 SV=1 - [OLA1_MOUSE] | 0,000E0 | 0,000E0 | 0,000E0 | 5,487E6 | 1,013E7 | 8,356E6 | Ola1 |
| O08810 | 116 kDa U5 small nuclear ribonucleoprotein component OS=Mus musculus GN=Eftud2 PE=1 SV=1 - [U5S1_MOUSE] | 0,000E0 | 0,000E0 | 0,000E0 | 1,263E6 | 4,777E6 | 1,038E7 | Eftud2 |
| P70261 | Paladin OS=Mus musculus GN=Pald1 PE=1 SV=1 - [PALD_MOUSE] | 0,000E0 | 0,000E0 | 0,000E0 | 3,772E6 | 1,473E7 | 2,316E7 | Pald1 |
| P61620 | Protein transport protein Sec61 subunit alpha isoform 1 OS=Mus musculus GN=Sec61a1 PE=1 SV=2 - [S61A1_MOUSE] | 0,000E0 | 0,000E0 | 0,000E0 | 8,238E6 | 1,329E7 | 1,101E7 | Sec61a1 |
| Q3UMU9-4 | Isoform 4 of Hepatoma-derived growth factor-related protein 2 OS=Mus musculus GN=Hdgfrp2 - [HDGR2_MOUSE] | 0,000E0 | 0,000E0 | 0,000E0 | 7,439E6 | 1,257E7 | 3,517E7 | Hdgfrp2 |
| Q8C166 | Copine-1 OS=Mus musculus GN=Cpne1 PE=1 SV=1 - [CPNE1_MOUSE] | 0,000E0 | 0,000E0 | 0,000E0 | 3,143E6 | 7,646E6 | 1,485E7 | Cpne1 |
| P05532-2 | Isoform 2 of Mast/stem cell growth factor receptor Kit OS=Mus musculus GN=Kit - [KIT_MOUSE] | 0,000E0 | 0,000E0 | 0,000E0 | 6,934E6 | 1,624E7 | 1,454E7 | Kit |
| P24788-2 | Isoform 2 of Cyclin-dependent kinase 11B OS=Mus musculus GN=Cdk11b - [CD11B_MOUSE] | 0,000E0 | 0,000E0 | 0,000E0 | 2,265E6 | 2,024E6 | 6,840E6 | Cdk11b |
| Q8BTH8 | Casein kinase I isoform gamma-1 OS=Mus musculus GN=Csnk1g1 PE=1 SV=2 - [KC1G1_MOUSE] | 0,000E0 | 0,000E0 | 0,000E0 | 1,557E6 | 6,473E6 | 6,647E6 | Csnk1g1 |
| Q8BSF4 | Phosphatidylserine decarboxylase proenzyme, mitochondrial OS=Mus musculus GN=Pisd PE=2 SV=1 - [PISD_MOUSE] | 0,000E0 | 0,000E0 | 0,000E0 | 1,467E6 | 9,084E6 | 9,000E5 | Pisd |
| Q80YW0 | Cytohesin-4 OS=Mus musculus GN=Cyth4 PE=1 SV=1 - [CYH4_MOUSE] | 0,000E0 | 0,000E0 | 0,000E0 | 6,843E6 | 2,783E6 | 1,430E6 | Cyth4 |
| Q8BGQ7 | Alanine--tRNA ligase, cytoplasmic OS=Mus musculus GN=Aars PE=1 SV=1 - [SYAC_MOUSE] | 0,000E0 | 0,000E0 | 2,770E6 | 1,262E6 | 2,349E6 | 4,970E6 | Aars |
| P54103 | DnaJ homolog subfamily C member 2 OS=Mus musculus GN=Dnajc2 PE=1 SV=2 - [DNJC2_MOUSE] | 0,000E0 | 0,000E0 | 0,000E0 | 2,778E6 | 8,690E6 | 9,383E6 | Dnajc2 |
| Q9Z130 | Heterogeneous nuclear ribonucleoprotein D-like OS=Mus musculus GN=Hnrnpdl PE=1 SV=1 - [HNRDL_MOUSE] | 0,000E0 | 0,000E0 | 0,000E0 | 2,962E6 | 3,767E7 | 8,374E5 | Hnrnpdl |
| O70591 | Prefoldin subunit 2 OS=Mus musculus GN=Pfdn2 PE=1 SV=2 - [PFD2_MOUSE] | 0,000E0 | 0,000E0 | 0,000E0 | 2,267E6 | 8,757E6 | 7,390E6 | Pfdn2 |
| Q99MU3-2 | Isoform 2 of Double-stranded RNA-specific adenosine deaminase OS=Mus musculus GN=Adar - [DSRAD_MOUSE] | 0,000E0 | 0,000E0 | 0,000E0 | 1,586E6 | 4,370E6 | 7,698E6 | Adar |
| Q922M3 | BTB/POZ domain-containing adapter for CUL3-mediated RhoA degradation protein 3 OS=Mus musculus GN=Kctd10 PE=1 SV=1 - [BACD3_MOUSE] | 0,000E0 | 0,000E0 | 0,000E0 | 1,638E6 | 1,853E6 | 2,809E6 | Kctd10 |
| Q9Z2N8 | Actin-like protein 6A OS=Mus musculus GN=Actl6a PE=1 SV=2 - [ACL6A_MOUSE] | 0,000E0 | 0,000E0 | 0,000E0 | 1,781E6 | 1,211E6 | 1,226E7 | Actl6a |
| Q8R0S2 | IQ motif and SEC7 domain-containing protein 1 OS=Mus musculus GN=Iqsec1 PE=1 SV=2 - [IQEC1_MOUSE] | 0,000E0 | 0,000E0 | 0,000E0 | 9,196E6 | 8,235E6 | 1,536E7 | Iqsec1 |
| P63158 | High mobility group protein B1 OS=Mus musculus GN=Hmgb1 PE=1 SV=2 - [HMGB1_MOUSE] | 0,000E0 | 0,000E0 | 1,350E6 | 5,490E6 | 1,478E7 | 1,527E7 | Hmgb1 |
| Q8C4X2 | Casein kinase I isoform gamma-3 OS=Mus musculus GN=Csnk1g3 PE=1 SV=2 - [KC1G3_MOUSE] | 0,000E0 | 0,000E0 | 0,000E0 | 8,887E5 | 7,405E6 | 1,273E7 | Csnk1g3 |
| Q8C0T5-2 | Isoform 2 of Signal-induced proliferation-associated 1-like protein 1 OS=Mus musculus GN=Sipa1l1 - [SI1L1_MOUSE] | 0,000E0 | 0,000E0 | 0,000E0 | 2,472E6 | 1,051E7 | 8,639E6 | Sipa1l1 |
| O54941 | SWI/SNF-related matrix-associated actin-dependent regulator of chromatin subfamily E member 1 OS=Mus musculus GN=Smarce1 PE=1 SV=1 - [SMCE1_MOUSE] | 0,000E0 | 0,000E0 | 0,000E0 | 1,832E6 | 6,045E6 | 5,037E6 | Smarce1 |
| Q569Z5 | Probable ATP-dependent RNA helicase DDX46 OS=Mus musculus GN=Ddx46 PE=1 SV=2 - [DDX46_MOUSE] | 0,000E0 | 0,000E0 | 0,000E0 | 1,133E6 | 4,288E6 | 1,951E7 | Ddx46 |
| Q99L47 | Hsc70-interacting protein OS=Mus musculus GN=St13 PE=1 SV=1 - [F10A1_MOUSE] | 0,000E0 | 0,000E0 | 0,000E0 | 3,360E6 | 1,326E7 | 1,360E7 | St13 |
| Q68FF6 | ARF GTPase-activating protein GIT1 OS=Mus musculus GN=Git1 PE=1 SV=1 - [GIT1_MOUSE] | 0,000E0 | 0,000E0 | 0,000E0 | 2,866E6 | 9,406E6 | 9,509E6 | Git1 |
| Q8CI08 | SLAIN motif-containing protein 2 OS=Mus musculus GN=Slain2 PE=1 SV=2 - [SLAI2_MOUSE] | 0,000E0 | 0,000E0 | 0,000E0 | 1,662E6 | 6,204E5 | 1,296E6 | Slain2 |
| Q9EQU5-2 | Isoform 2 of Protein SET OS=Mus musculus GN=Set - [SET_MOUSE] | 0,000E0 | 0,000E0 | 0,000E0 | 1,135E6 | 2,098E6 | 1,744E6 | Set |
| P40142 | Transketolase OS=Mus musculus GN=Tkt PE=1 SV=1 - [TKT_MOUSE] | 0,000E0 | 0,000E0 | 0,000E0 | 1,552E6 | 2,564E6 | 4,408E6 | Tkt |
| P28740 | Kinesin-like protein KIF2A OS=Mus musculus GN=Kif2a PE=1 SV=2 - [KIF2A_MOUSE] | 0,000E0 | 0,000E0 | 0,000E0 | 3,446E6 | 7,067E6 | 1,070E7 | Kif2a |
| P63073 | Eukaryotic translation initiation factor 4E OS=Mus musculus GN=Eif4e PE=1 SV=1 - [IF4E_MOUSE] | 0,000E0 | 0,000E0 | 0,000E0 | 1,825E6 | 1,121E7 | 1,342E7 | Eif4e |
| Q69ZR2 | E3 ubiquitin-protein ligase HECTD1 OS=Mus musculus GN=Hectd1 PE=1 SV=2 - [HECD1_MOUSE] | 0,000E0 | 0,000E0 | 0,000E0 | 4,761E6 | 8,592E6 | 4,090E7 | Hectd1 |
| Q62376 | U1 small nuclear ribonucleoprotein 70 kDa OS=Mus musculus GN=Snrnp70 PE=1 SV=2 - [RU17_MOUSE] | 0,000E0 | 0,000E0 | 0,000E0 | 3,976E6 | 8,504E6 | 1,748E7 | Snrnp70 |
| Q9DC70 | NADH dehydrogenase [ubiquinone] iron-sulfur protein 7, mitochondrial OS=Mus musculus GN=Ndufs7 PE=1 SV=1 - [NDUS7_MOUSE] | 0,000E0 | 0,000E0 | 0,000E0 | 1,121E6 | 2,305E6 | 4,994E6 | Ndufs7 |
| O09110 | Dual specificity mitogen-activated protein kinase kinase 3 OS=Mus musculus GN=Map2k3 PE=1 SV=2 - [MP2K3_MOUSE] | 0,000E0 | 0,000E0 | 0,000E0 | 3,822E6 | 2,368E6 | 2,177E6 | Map2k3 |
| Q3U9G9 | Lamin-B receptor OS=Mus musculus GN=Lbr PE=1 SV=2 - [LBR_MOUSE] | 0,000E0 | 0,000E0 | 0,000E0 | 9,738E6 | 2,503E7 | 3,335E7 | Lbr |
| Q8CH72 | E3 ubiquitin-protein ligase TRIM32 OS=Mus musculus GN=Trim32 PE=1 SV=2 - [TRI32_MOUSE] | 0,000E0 | 0,000E0 | 0,000E0 | 6,924E6 | 4,576E6 | 1,080E7 | Trim32 |
| Q8BNJ2 | A disintegrin and metalloproteinase with thrombospondin motifs 4 OS=Mus musculus GN=Adamts4 PE=2 SV=2 - [ATS4_MOUSE] | 0,000E0 | 0,000E0 | 0,000E0 | 1,303E7 | 2,145E7 | 1,972E7 | Adamts4 |
| E9Q555 | E3 ubiquitin-protein ligase RNF213 OS=Mus musculus GN=Rnf213 PE=1 SV=2 - [RN213_MOUSE] | 0,000E0 | 0,000E0 | 8,642E5 | 1,198E7 | 1,853E7 | 3,576E7 | Rnf213 |
| Q60790 | Ras GTPase-activating protein 3 OS=Mus musculus GN=Rasa3 PE=1 SV=2 - [RASA3_MOUSE] | 0,000E0 | 0,000E0 | 0,000E0 | 3,180E6 | 6,446E6 | 1,620E7 | Rasa3 |
| Q8BHD7-2 | Isoform 2 of Polypyrimidine tract-binding protein 3 OS=Mus musculus GN=Ptbp3 - [PTBP3_MOUSE] | 0,000E0 | 0,000E0 | 0,000E0 | 3,256E7 | 5,081E7 | 1,794E8 | Ptbp3 |
| P17095-1 | Isoform HMG-Y of High mobility group protein HMG-I/HMG-Y OS=Mus musculus GN=Hmga1 - [HMGA1_MOUSE] | 0,000E0 | 0,000E0 | 0,000E0 | 8,704E7 | 1,389E8 | 5,081E8 | Hmga1 |
| Q8BH88 | DEP domain-containing protein 1B OS=Mus musculus GN=Depdc1b PE=2 SV=1 - [DEP1B_MOUSE] | 0,000E0 | 0,000E0 | 0,000E0 | 3,815E5 | 4,607E6 | 1,198E6 | Depdc1b |
| Q3TIV5 | Zinc finger CCCH domain-containing protein 15 OS=Mus musculus GN=Zc3h15 PE=1 SV=2 - [ZC3HF_MOUSE] | 0,000E0 | 0,000E0 | 0,000E0 | 5,932E6 | 2,158E6 | 4,901E6 | Zc3h15 |
| Q8CGC7 | Bifunctional glutamate/proline--tRNA ligase OS=Mus musculus GN=Eprs PE=1 SV=4 - [SYEP_MOUSE] | 0,000E0 | 0,000E0 | 0,000E0 | 5,461E6 | 8,137E6 | 9,736E6 | Eprs |
| Q64737 | Trifunctional purine biosynthetic protein adenosine-3 OS=Mus musculus GN=Gart PE=1 SV=3 - [PUR2_MOUSE] | 0,000E0 | 0,000E0 | 0,000E0 | 5,563E6 | 1,099E7 | 1,584E7 | Gart |
| Q922D8 | C-1-tetrahydrofolate synthase, cytoplasmic OS=Mus musculus GN=Mthfd1 PE=1 SV=4 - [C1TC_MOUSE] | 0,000E0 | 0,000E0 | 0,000E0 | 2,757E6 | 1,379E7 | 1,235E7 | Mthfd1 |
| Q9ESL4-2 | Isoform 2 of Mitogen-activated protein kinase kinase kinase MLT OS=Mus musculus GN=Zak - [MLTK_MOUSE] | 0,000E0 | 0,000E0 | 0,000E0 | 2,924E6 | 6,027E6 | 8,104E6 | Zak |
| Q9R0E1 | Procollagen-lysine,2-oxoglutarate 5-dioxygenase 3 OS=Mus musculus GN=Plod3 PE=1 SV=1 - [PLOD3_MOUSE] | 0,000E0 | 0,000E0 | 0,000E0 | 1,599E6 | 5,597E6 | 7,792E6 | Plod3 |
| Q60953-2 | Isoform 2 of Protein PML OS=Mus musculus GN=Pml - [PML_MOUSE] | 0,000E0 | 0,000E0 | 0,000E0 | 5,147E6 | 1,029E7 | 1,552E7 | Pml |
| Q8BWY3 | Eukaryotic peptide chain release factor subunit 1 OS=Mus musculus GN=Etf1 PE=1 SV=4 - [ERF1_MOUSE] | 0,000E0 | 0,000E0 | 0,000E0 | 2,605E6 | 1,678E6 | 7,679E6 | Etf1 |
| Q9CZ52 | Anthrax toxin receptor 1 OS=Mus musculus GN=Antxr1 PE=1 SV=2 - [ANTR1_MOUSE] | 0,000E0 | 0,000E0 | 0,000E0 | 5,248E6 | 1,095E7 | 1,206E7 | Antxr1 |
| P62046 | Leucine-rich repeat and calponin homology domain-containing protein 1 OS=Mus musculus GN=Lrch1 PE=1 SV=2 - [LRCH1_MOUSE] | 0,000E0 | 0,000E0 | 0,000E0 | 4,710E7 | 8,244E6 | 3,222E6 | Lrch1 |
| Q8K363 | ATP-dependent RNA helicase DDX18 OS=Mus musculus GN=Ddx18 PE=1 SV=1 - [DDX18_MOUSE] | 0,000E0 | 0,000E0 | 0,000E0 | 2,499E6 | 5,402E6 | 2,078E7 | Ddx18 |
| Q9ERU9 | E3 SUMO-protein ligase RanBP2 OS=Mus musculus GN=Ranbp2 PE=1 SV=2 - [RBP2_MOUSE] | 0,000E0 | 0,000E0 | 0,000E0 | 3,740E6 | 6,148E6 | 2,168E7 | Ranbp2 |
| O35691 | Pinin OS=Mus musculus GN=Pnn PE=1 SV=4 - [PININ_MOUSE] | 0,000E0 | 0,000E0 | 0,000E0 | 3,341E6 | 7,965E6 | 9,962E6 | Pnn |
| P70255-5 | Isoform 5 of Nuclear factor 1 C-type OS=Mus musculus GN=Nfic - [NFIC_MOUSE] | 0,000E0 | 0,000E0 | 0,000E0 | 8,960E6 | 4,007E6 | 7,908E5 | Nfic |
| Q7TN60 | Transmembrane channel-like protein 6 OS=Mus musculus GN=Tmc6 PE=1 SV=2 - [TMC6_MOUSE] | 0,000E0 | 0,000E0 | 0,000E0 | 4,582E6 | 8,849E6 | 8,960E6 | Tmc6 |
| Q8CES0 | N-alpha-acetyltransferase 30 OS=Mus musculus GN=Naa30 PE=1 SV=2 - [NAA30_MOUSE] | 0,000E0 | 0,000E0 | 0,000E0 | 2,084E6 | 4,505E6 | 6,319E6 | Naa30 |
| P05132-2 | Isoform 2 of cAMP-dependent protein kinase catalytic subunit alpha OS=Mus musculus GN=Prkaca - [KAPCA_MOUSE] | 0,000E0 | 0,000E0 | 0,000E0 | 7,310E6 | 9,244E6 | 1,493E7 | Prkaca |
| P70429-2 | Isoform 1 of Ena/VASP-like protein OS=Mus musculus GN=Evl - [EVL_MOUSE] | 0,000E0 | 0,000E0 | 0,000E0 | 1,947E6 | 5,076E6 | 3,122E6 | Evl |
| Q3UMG5 | Leucine-rich repeat and calponin homology domain-containing protein 2 OS=Mus musculus GN=Lrch2 PE=2 SV=2 - [LRCH2_MOUSE] | 0,000E0 | 0,000E0 | 0,000E0 | 2,261E6 | 4,988E5 | 3,334E6 | Lrch2 |
| Q62432-2 | Isoform Short of Mothers against decapentaplegic homolog 2 OS=Mus musculus GN=Smad2 - [SMAD2_MOUSE] | 0,000E0 | 0,000E0 | 0,000E0 | 1,545E6 | 1,128E7 | 7,537E6 | Smad2 |

| | | | | | | | | |
|---|---|---|---|---|---|---|---|---|
| Q791V5 | Mitochondrial carrier homolog 2 OS=Mus musculus GN=Mtch2 PE=1 SV=1 - [MTCH2_MOUSE] | 0,000E0 | 0,000E0 | 0,000E0 | 2,528E6 | 6,857E6 | 1,589E7 | Mtch2 |
| P28740-2 | Isoform 2 of Kinesin-like protein KIF2A OS=Mus musculus GN=Kif2a - [KIF2A_MOUSE] | 0,000E0 | 0,000E0 | 0,000E0 | 3,446E6 | 7,067E6 | 1,070E7 | Kif2a |
| Q6PGH0 | Ubiquitin domain-containing protein 2 OS=Mus musculus GN=Ubtd2 PE=2 SV=1 - [UBTD2_MOUSE] | 0,000E0 | 0,000E0 | 0,000E0 | 1,088E6 | 9,254E5 | 2,288E6 | Ubtd2 |
| Q91VZ6 | Stromal membrane-associated protein 1 OS=Mus musculus GN=Smap1 PE=1 SV=1 - [SMAP1_MOUSE] | 0,000E0 | 0,000E0 | 0,000E0 | 7,338E6 | 5,100E6 | 1,010E7 | Smap1 |
| Q8K310 | Matrin-3 OS=Mus musculus GN=Matr3 PE=1 SV=1 - [MATR3_MOUSE] | 0,000E0 | 0,000E0 | 0,000E0 | 3,095E6 | 6,647E6 | 8,931E6 | Matr3 |
| Q9R1C7 | Pre-mRNA-processing factor 40 homolog A OS=Mus musculus GN=Prpf40a PE=1 SV=1 - [PR40A_MOUSE] | 0,000E0 | 0,000E0 | 0,000E0 | 3,490E6 | 1,286E6 | 6,385E6 | Prpf40a |
| Q8BP48 | Methionine aminopeptidase 1 OS=Mus musculus GN=Metap1 PE=1 SV=1 - [MAP11_MOUSE] | 0,000E0 | 0,000E0 | 0,000E0 | 5,609E6 | 8,226E6 | 1,541E7 | Metap1 |
| Q922H2 | [Pyruvate dehydrogenase (acetyl-transferring)] kinase isozyme 3, mitochondrial OS=Mus musculus GN=Pdk3 PE=1 SV=1 - [PDK3_MOUSE] | 0,000E0 | 0,000E0 | 0,000E0 | 1,334E6 | 2,837E6 | 4,537E6 | Pdk3 |
| O89032-3 | Isoform 3 of SH3 and PX domain-containing protein 2A OS=Mus musculus GN=Sh3pxd2a - [SPD2A_MOUSE] | 0,000E0 | 0,000E0 | 0,000E0 | 1,672E6 | 2,803E6 | 7,624E5 | Sh3pxd2a |
| Q7TNV0 | Protein DEK OS=Mus musculus GN=Dek PE=1 SV=1 - [DEK_MOUSE] | 0,000E0 | 0,000E0 | 0,000E0 | 9,174E6 | 1,565E7 | 4,193E7 | Dek |
| P58742 | Aladin OS=Mus musculus GN=Aaas PE=1 SV=1 - [AAAS_MOUSE] | 0,000E0 | 0,000E0 | 0,000E0 | 2,053E6 | 3,132E6 | 4,919E6 | Aaas |
| Q922B9 | Sperm-specific antigen 2 homolog OS=Mus musculus GN=Ssfa2 PE=1 SV=3 - [SSFA2_MOUSE] | 0,000E0 | 0,000E0 | 0,000E0 | 1,922E6 | 7,426E6 | 1,126E7 | Ssfa2 |
| Q9CX56 | 26S proteasome non-ATPase regulatory subunit 8 OS=Mus musculus GN=Psmd8 PE=1 SV=2 - [PSMD8_MOUSE] | 0,000E0 | 0,000E0 | 0,000E0 | 1,638E6 | 2,757E6 | 6,820E6 | Psmd8 |
| Q9D0R8 | Protein LSM12 homolog OS=Mus musculus GN=Lsm12 PE=1 SV=1 - [LSM12_MOUSE] | 0,000E0 | 0,000E0 | 0,000E0 | 6,599E6 | 1,335E7 | 4,919E7 | Lsm12 |
| P52479 | Ubiquitin carboxyl-terminal hydrolase 10 OS=Mus musculus GN=Usp10 PE=1 SV=3 - [UBP10_MOUSE] | 0,000E0 | 0,000E0 | 0,000E0 | 5,551E6 | 5,949E6 | 1,147E7 | Usp10 |
| E9QAM5 | Helicase with zinc finger domain 2 OS=Mus musculus GN=Helz2 PE=1 SV=1 - [HELZ2_MOUSE] | 0,000E0 | 0,000E0 | 0,000E0 | 4,552E6 | 7,940E6 | 1,770E7 | Helz2 |
| P97857 | A disintegrin and metalloproteinase with thrombospondin motifs 1 OS=Mus musculus GN=Adamts1 PE=1 SV=4 - [ATS1_MOUSE] | 0,000E0 | 0,000E0 | 0,000E0 | 1,311E6 | 1,522E7 | 2,013E7 | Adamts1 |
| Q91WF3 | Adenylate cyclase type 4 OS=Mus musculus GN=Adcy4 PE=1 SV=1 - [ADCY4_MOUSE] | 0,000E0 | 0,000E0 | 0,000E0 | 1,978E7 | 4,084E6 | 3,966E7 | Adcy4 |
| Q9QY06 | Unconventional myosin-IXb OS=Mus musculus GN=Myo9b PE=1 SV=2 - [MYO9B_MOUSE] | 0,000E0 | 0,000E0 | 0,000E0 | 1,760E7 | 5,584E6 | 1,397E7 | Myo9b |
| Q60930 | Voltage-dependent anion-selective channel protein 2 OS=Mus musculus GN=Vdac2 PE=1 SV=2 - [VDAC2_MOUSE] | 0,000E0 | 0,000E0 | 0,000E0 | 2,949E6 | 2,913E6 | 2,793E7 | Vdac2 |
| Q61211 | Eukaryotic translation initiation factor 2D OS=Mus musculus GN=Eif2d PE=1 SV=3 - [EIF2D_MOUSE] | 0,000E0 | 0,000E0 | 0,000E0 | 1,481E6 | 3,212E6 | 3,017E6 | Eif2d |
| P52624 | Uridine phosphorylase 1 OS=Mus musculus GN=Upp1 PE=1 SV=2 - [UPP1_MOUSE] | 0,000E0 | 0,000E0 | 0,000E0 | 1,649E7 | 1,939E7 | 3,489E7 | Upp1 |
| P31938 | Dual specificity mitogen-activated protein kinase kinase 1 OS=Mus musculus GN=Map2k1 PE=1 SV=2 - [MP2K1_MOUSE] | 0,000E0 | 0,000E0 | 0,000E0 | 2,900E6 | 5,955E6 | 5,631E6 | Map2k1 |
| Q9QUJ7-2 | Isoform Short of Long-chain-fatty-acid--CoA ligase 4 OS=Mus musculus GN=Acsl4 - [ACSL4_MOUSE] | 0,000E0 | 0,000E0 | 0,000E0 | 1,513E6 | 3,519E6 | 6,474E6 | Acsl4 |
| Q8K2Y3 | Protein eva-1 homolog B OS=Mus musculus GN=Eva1b PE=1 SV=1 - [EVA1B_MOUSE] | 0,000E0 | 0,000E0 | 0,000E0 | 7,679E6 | 1,947E7 | 2,912E7 | Eva1b |
| P61290 | Proteasome activator complex subunit 3 OS=Mus musculus GN=Psme3 PE=1 SV=1 - [PSME3_MOUSE] | 0,000E0 | 0,000E0 | 0,000E0 | 9,215E5 | 2,921E6 | 2,106E6 | Psme3 |
| Q64282 | Interferon-induced protein with tetratricopeptide repeats 1 OS=Mus musculus GN=Ifit1 PE=1 SV=2 - [IFIT1_MOUSE] | 0,000E0 | 0,000E0 | 0,000E0 | 2,261E6 | 2,178E6 | 3,218E6 | Ifit1 |
| Q6ZQ08-2 | Isoform 2 of CCR4-NOT transcription complex subunit 1 OS=Mus musculus GN=Cnot1 - [CNOT1_MOUSE] | 0,000E0 | 0,000E0 | 0,000E0 | 3,243E6 | 9,629E6 | 2,528E7 | Cnot1 |
| B2RXS4 | Plexin-B2 OS=Mus musculus GN=Plxnb2 PE=1 SV=1 - [PLXB2_MOUSE] | 0,000E0 | 0,000E0 | 0,000E0 | 8,971E6 | 2,376E7 | 2,615E7 | Plxnb2 |
| Q8BYZ1 | ABI gene family member 3 OS=Mus musculus GN=Abi3 PE=1 SV=3 - [ABI3_MOUSE] | 0,000E0 | 0,000E0 | 0,000E0 | 1,460E6 | 1,228E6 | 1,285E7 | Abi3 |
| Q9DBG6 | Dolichyl-diphosphooligosaccharide--protein glycosyltransferase subunit 2 OS=Mus musculus GN=Rpn2 PE=1 SV=1 - [RPN2_MOUSE] | 0,000E0 | 0,000E0 | 0,000E0 | 4,305E6 | 2,029E6 | 4,959E6 | Rpn2 |
| P70290 | 55 kDa erythrocyte membrane protein OS=Mus musculus GN=Mpp1 PE=1 SV=1 - [EM55_MOUSE] | 0,000E0 | 0,000E0 | 0,000E0 | 1,230E6 | 3,760E6 | 5,170E6 | Mpp1 |
| Q6A0A2-2 | Isoform 2 of La-related protein 4B OS=Mus musculus GN=Larp4b - [LAR4B_MOUSE] | 0,000E0 | 0,000E0 | 0,000E0 | 2,613E6 | 4,581E6 | 7,816E6 | Larp4b |
| Q8BGH2 | Sorting and assembly machinery component 50 homolog OS=Mus musculus GN=Samm50 PE=1 SV=1 - [SAM50_MOUSE] | 0,000E0 | 0,000E0 | 0,000E0 | 1,496E6 | 4,964E6 | 5,368E6 | Samm50 |
| P46662-2 | Isoform 2 of Merlin OS=Mus musculus GN=Nf2 - [MERL_MOUSE] | 0,000E0 | 0,000E0 | 0,000E0 | 3,500E6 | 7,885E6 | 5,335E6 | Nf2 |
| Q91XI1 | tRNA-dihydrouridine(47) synthase [NAD(P)(+)]-like OS=Mus musculus GN=Dus3l PE=1 SV=1 - [DUS3L_MOUSE] | 0,000E0 | 0,000E0 | 0,000E0 | 2,557E6 | 1,539E6 | 6,855E6 | Dus3l |
| Q99K41 | EMILIN-1 OS=Mus musculus GN=Emilin1 PE=1 SV=1 - [EMIL1_MOUSE] | 0,000E0 | 0,000E0 | 0,000E0 | 3,541E6 | 7,391E6 | 1,837E7 | Emilin1 |
| P23298 | Protein kinase C eta type OS=Mus musculus GN=Prkch PE=1 SV=2 - [KPCL_MOUSE] | 0,000E0 | 0,000E0 | 0,000E0 | 1,263E6 | 4,204E6 | 5,704E6 | Prkch |
| Q8K297 | Procollagen galactosyltransferase 1 OS=Mus musculus GN=Colgalt1 PE=1 SV=2 - [GT251_MOUSE] | 0,000E0 | 0,000E0 | 0,000E0 | 9,669E5 | 2,498E6 | 5,346E6 | Colgalt1 |
| P08113 | Endoplasmin OS=Mus musculus GN=Hsp90b1 PE=1 SV=2 - [ENPL_MOUSE] | 0,000E0 | 0,000E0 | 1,544E6 | 4,372E7 | 8,485E7 | 1,170E8 | Hsp90b1 |
| P62748 | Hippocalcin-like protein 1 OS=Mus musculus GN=Hpcal1 PE=1 SV=2 - [HPCL1_MOUSE] | 0,000E0 | 0,000E0 | 0,000E0 | 1,121E6 | 2,203E6 | 2,076E6 | Hpcal1 |
| P02088 | Hemoglobin subunit beta-1 OS=Mus musculus GN=Hbb-b1 PE=1 SV=2 - [HBB1_MOUSE] | 0,000E0 | 0,000E0 | 0,000E0 | 1,094E7 | 6,133E6 | 5,045E7 | Hbb-b1 |
| O54781 | SRSF protein kinase 2 OS=Mus musculus GN=Srpk2 PE=1 SV=2 - [SRPK2_MOUSE] | 0,000E0 | 0,000E0 | 0,000E0 | 3,952E6 | 8,114E6 | 2,294E7 | Srpk2 |
| P53811 | Phosphatidylinositol transfer protein beta isoform OS=Mus musculus GN=Pitpnb PE=1 SV=2 - [PIPNB_MOUSE] | 0,000E0 | 0,000E0 | 0,000E0 | 8,319E5 | 1,857E6 | 5,166E6 | Pitpnb |
| Q61286-2 | Isoform ALF1A of Transcription factor 12 OS=Mus musculus GN=Tcf12 - [HTF4_MOUSE] | 0,000E0 | 0,000E0 | 0,000E0 | 2,336E6 | 3,253E6 | 8,158E6 | Tcf12 |
| Q99P88 | Nuclear pore complex protein Nup155 OS=Mus musculus GN=Nup155 PE=1 SV=1 - [NU155_MOUSE] | 0,000E0 | 0,000E0 | 0,000E0 | 4,020E6 | 4,944E6 | 3,121E6 | Nup155 |
| Q9ESL4 | Mitogen-activated protein kinase kinase kinase MLT OS=Mus musculus GN=Zak PE=1 SV=1 - [MLTK_MOUSE] | 0,000E0 | 0,000E0 | 0,000E0 | 2,076E6 | 3,030E6 | 8,300E6 | Zak |
| P56399 | Ubiquitin carboxyl-terminal hydrolase 5 OS=Mus musculus GN=Usp5 PE=1 SV=1 - [UBP5_MOUSE] | 0,000E0 | 0,000E0 | 0,000E0 | 1,050E6 | 4,989E6 | 6,692E6 | Usp5 |
| P47856-2 | Isoform 2 of Glutamine--fructose-6-phosphate aminotransferase [isomerizing] 1 OS=Mus musculus GN=Gfpt1 - [GFPT1_MOUSE] | 0,000E0 | 0,000E0 | 0,000E0 | 6,250E5 | 1,727E6 | 4,308E6 | Gfpt1 |
| Q8BH43 | Wiskott-Aldrich syndrome protein family member 2 OS=Mus musculus GN=Wasf2 PE=1 SV=1 - [WASF2_MOUSE] | 0,000E0 | 0,000E0 | 0,000E0 | 6,262E6 | 4,434E7 | 7,108E7 | Wasf2 |
| Q99NH0 | Ankyrin repeat domain-containing protein 17 OS=Mus musculus GN=Ankrd17 PE=1 SV=2 - [ANR17_MOUSE] | 0,000E0 | 0,000E0 | 0,000E0 | 4,666E6 | 2,623E7 | 2,195E7 | Ankrd17 |
| Q9Z2F2 | 2'-5'-oligoadenylate synthase-like protein 2 OS=Mus musculus GN=Oasl2 PE=1 SV=2 - [OASL2_MOUSE] | 0,000E0 | 0,000E0 | 0,000E0 | 1,002E6 | 1,240E7 | 2,688E7 | Oasl2 |
| P18052-2 | Isoform 2 of Receptor-type tyrosine-protein phosphatase alpha OS=Mus musculus GN=Ptpra - [PTPRA_MOUSE] | 0,000E0 | 0,000E0 | 0,000E0 | 1,240E7 | 1,291E7 | 5,005E6 | Ptpra |
| Q9ESW4 | Acylglycerol kinase, mitochondrial OS=Mus musculus GN=Agk PE=1 SV=1 - [AGK_MOUSE] | 0,000E0 | 0,000E0 | 0,000E0 | 1,982E6 | 4,453E6 | 6,674E6 | Agk |
| Q6P8X1 | Sorting nexin-6 OS=Mus musculus GN=Snx6 PE=1 SV=2 - [SNX6_MOUSE] | 0,000E0 | 0,000E0 | 0,000E0 | 2,933E6 | 1,510E6 | 8,915E6 | Snx6 |
| Q8VCG3 | WD repeat-containing protein 74 OS=Mus musculus GN=Wdr74 PE=2 SV=1 - [WDR74_MOUSE] | 0,000E0 | 0,000E0 | 0,000E0 | 8,721E5 | 3,251E6 | 5,117E6 | Wdr74 |
| Q3TFK5-2 | Isoform 2 of G patch domain-containing protein 4 OS=Mus musculus GN=Gpatch4 - [GPTC4_MOUSE] | 0,000E0 | 0,000E0 | 0,000E0 | 1,561E6 | 4,151E6 | 7,693E6 | Gpatch4 |
| O35344 | Importin subunit alpha-4 OS=Mus musculus GN=Kpna3 PE=1 SV=1 - [IMA4_MOUSE] | 0,000E0 | 0,000E0 | 0,000E0 | 1,775E6 | 2,925E6 | 1,103E6 | Kpna3 |
| Q08093 | Calponin-2 OS=Mus musculus GN=Cnn2 PE=1 SV=1 - [CNN2_MOUSE] | 0,000E0 | 0,000E0 | 0,000E0 | 9,081E5 | 2,767E6 | 2,943E7 | Cnn2 |
| Q811D2 | Ankyrin repeat domain-containing protein 26 OS=Mus musculus GN=Ankrd26 PE=1 SV=2 - [ANR26_MOUSE] | 0,000E0 | 0,000E0 | 0,000E0 | 3,144E6 | 8,723E6 | 7,694E6 | Ankrd26 |
| Q8CG47 | Structural maintenance of chromosomes protein 4 OS=Mus musculus GN=Smc4 PE=1 SV=1 - [SMC4_MOUSE] | 0,000E0 | 0,000E0 | 0,000E0 | 1,233E6 | 5,219E6 | 1,148E7 | Smc4 |
| Q9R0M6 | Ras-related protein Rab-9A OS=Mus musculus GN=Rab9a PE=1 SV=1 - [RAB9A_MOUSE] | 0,000E0 | 0,000E0 | 0,000E0 | 9,937E5 | 4,799E6 | 5,981E6 | Rab9a |
| P01899 | H-2 class I histocompatibility antigen, D-B alpha chain OS=Mus musculus GN=H2-D1 PE=1 SV=2 - [HA11_MOUSE] | 0,000E0 | 0,000E0 | 0,000E0 | 3,338E6 | 3,956E6 | 5,133E6 | H2-D1 |
| Q60996-2 | Isoform 1 of Serine/threonine-protein phosphatase 2A 56 kDa regulatory subunit gamma isoform OS=Mus musculus GN=Ppp2r5c - [2A5G_MOUSE] | 0,000E0 | 0,000E0 | 0,000E0 | 1,138E6 | 5,271E6 | 8,010E6 | Ppp2r5c |
| P48725-3 | Isoform 3 of Pericentrin OS=Mus musculus GN=Pcnt - [PCNT_MOUSE] | 0,000E0 | 0,000E0 | 0,000E0 | 4,379E6 | 6,258E6 | 1,140E7 | Pcnt |
| Q8VEJ9 | Vacuolar protein sorting-associated protein 4A OS=Mus musculus GN=Vps4a PE=1 SV=1 - [VPS4A_MOUSE] | 0,000E0 | 0,000E0 | 0,000E0 | 6,175E6 | 1,626E7 | 6,757E6 | Vps4a |
| Q921F2 | TAR DNA-binding protein 43 OS=Mus musculus GN=Tardbp PE=1 SV=1 - [TADBP_MOUSE] | 0,000E0 | 0,000E0 | 0,000E0 | 1,031E7 | 7,503E6 | 1,321E7 | Tardbp |
| O88967 | ATP-dependent zinc metalloprotease YME1L1 OS=Mus musculus GN=Yme1l1 PE=1 SV=1 - [YMEL1_MOUSE] | 0,000E0 | 0,000E0 | 0,000E0 | 5,834E5 | 4,008E6 | 1,804E6 | Yme1l1 |
| Q7TPM1 | Protein PRRC2B OS=Mus musculus GN=Prrc2b PE=1 SV=1 - [PRC2B_MOUSE] | 0,000E0 | 0,000E0 | 0,000E0 | 3,412E6 | 4,394E6 | 1,053E7 | Prrc2b |
| Q3THK7 | GMP synthase [glutamine-hydrolyzing] OS=Mus musculus GN=Gmps PE=1 SV=2 - [GUAA_MOUSE] | 0,000E0 | 0,000E0 | 0,000E0 | 2,168E6 | 4,535E6 | 1,321E7 | Gmps |
| Q6ZPG2 | WD repeat-containing protein 90 OS=Mus musculus GN=Wdr90 PE=1 SV=2 - [WDR90_MOUSE] | 0,000E0 | 0,000E0 | 0,000E0 | 2,192E6 | 3,340E6 | 2,379E6 | Wdr90 |
| Q91ZR2 | Sorting nexin-18 OS=Mus musculus GN=Snx18 PE=1 SV=1 - [SNX18_MOUSE] | 0,000E0 | 0,000E0 | 0,000E0 | 5,857E6 | 1,448E7 | 2,441E7 | Snx18 |
| O70585-2 | Isoform 2 of Dystrobrevin beta OS=Mus musculus GN=Dtnb - [DTNB_MOUSE] | 0,000E0 | 0,000E0 | 0,000E0 | 1,979E6 | 1,019E7 | 1,152E7 | Dtnb |
| Q99JP0 | Mitogen-activated protein kinase kinase kinase kinase 3 OS=Mus musculus GN=Map4k3 PE=1 SV=4 - [M4K3_MOUSE] | 0,000E0 | 0,000E0 | 0,000E0 | 9,770E5 | 4,454E6 | 5,719E6 | Map4k3 |
| Q6A4J8-3 | Isoform 3 of Ubiquitin carboxyl-terminal hydrolase 7 OS=Mus musculus GN=Usp7 - [UBP7_MOUSE] | 0,000E0 | 0,000E0 | 0,000E0 | 5,889E6 | 2,984E6 | 4,744E6 | Usp7 |
| Q9CZJ2 | Heat shock 70 kDa protein 12B OS=Mus musculus GN=Hspa12b PE=1 SV=1 - [HS12B_MOUSE] | 0,000E0 | 0,000E0 | 0,000E0 | 3,971E6 | 5,995E6 | 1,628E7 | Hspa12b |
| Q8BZ03 | Serine/threonine-protein kinase D2 OS=Mus musculus GN=Prkd2 PE=1 SV=1 - [KPCD2_MOUSE] | 0,000E0 | 0,000E0 | 0,000E0 | 5,511E6 | 1,064E7 | 1,232E7 | Prkd2 |
| Q61210-2 | Isoform 2 of Rho guanine nucleotide exchange factor 1 OS=Mus musculus GN=Arhgef1 - [ARHG1_MOUSE] | 0,000E0 | 0,000E0 | 0,000E0 | 1,986E6 | 2,325E6 | 8,846E6 | Arhgef1 |
| Q8CG48 | Structural maintenance of chromosomes protein 2 OS=Mus musculus GN=Smc2 PE=1 SV=2 - [SMC2_MOUSE] | 0,000E0 | 0,000E0 | 0,000E0 | 3,193E6 | 3,833E6 | 5,829E6 | Smc2 |
| Q8CI51 | PDZ and LIM domain protein 5 OS=Mus musculus GN=Pdlim5 PE=1 SV=4 - [PDLI5_MOUSE] | 0,000E0 | 0,000E0 | 0,000E0 | 1,462E6 | 4,524E6 | 1,317E7 | Pdlim5 |
| P49615 | Cyclin-dependent-like kinase 5 OS=Mus musculus GN=Cdk5 PE=1 SV=1 - [CDK5_MOUSE] | 0,000E0 | 0,000E0 | 0,000E0 | 2,128E7 | 1,886E7 | 8,067E7 | Cdk5 |
| Q8K0V4 | CCR4-NOT transcription complex subunit 3 OS=Mus musculus GN=Cnot3 PE=1 SV=1 - [CNOT3_MOUSE] | 0,000E0 | 0,000E0 | 0,000E0 | 2,547E6 | 8,552E6 | 1,123E7 | Cnot3 |
| P43406 | Integrin alpha-V OS=Mus musculus GN=Itgav PE=1 SV=2 - [ITAV_MOUSE] | 0,000E0 | 0,000E0 | 0,000E0 | 4,718E6 | 5,063E6 | 7,399E6 | Itgav |
| E9PVX6 | Proliferation marker protein Ki-67 OS=Mus musculus GN=Mki67 PE=1 SV=1 - [KI67_MOUSE] | 0,000E0 | 0,000E0 | 0,000E0 | 1,850E6 | 3,812E6 | 1,244E7 | Mki67 |
| Q9DCN2-2 | Isoform 2 of NADH-cytochrome b5 reductase 3 OS=Mus musculus GN=Cyb5r3 - [NB5R3_MOUSE] | 0,000E0 | 0,000E0 | 0,000E0 | 4,026E6 | 4,921E6 | 7,402E6 | Cyb5r3 |
| P09103 | Protein disulfide-isomerase OS=Mus musculus GN=P4hb PE=1 SV=2 - [PDIA1_MOUSE] | 0,000E0 | 0,000E0 | 0,000E0 | 4,297E6 | 3,028E6 | 3,019E6 | P4hb |
| P30416 | Peptidyl-prolyl cis-trans isomerase FKBP4 OS=Mus musculus GN=Fkbp4 PE=1 SV=5 - [FKBP4_MOUSE] | 0,000E0 | 0,000E0 | 0,000E0 | 7,166E5 | 3,053E6 | 3,546E6 | Fkbp4 |
| Q91YT7 | YTH domain-containing family protein 2 OS=Mus musculus GN=Ythdf2 PE=1 SV=1 - [YTHD2_MOUSE] | 0,000E0 | 0,000E0 | 5,369E6 | 5,946E6 | 2,335E6 | 1,089E7 | Ythdf2 |
| Q8BVY0 | Ribosomal L1 domain-containing protein 1 OS=Mus musculus GN=Rsl1d1 PE=1 SV=1 - [RL1D1_MOUSE] | 0,000E0 | 0,000E0 | 0,000E0 | 1,323E6 | 2,043E6 | 1,577E7 | Rsl1d1 |
| Q9D883 | Splicing factor U2AF 35 kDa subunit OS=Mus musculus GN=U2af1 PE=1 SV=4 - [U2AF1_MOUSE] | 0,000E0 | 0,000E0 | 0,000E0 | 5,700E6 | 6,966E6 | 6,575E6 | U2af1 |
| A2RTL5 | Arginine/serine-rich coiled-coil protein 2 OS=Mus musculus GN=Rsrc2 PE=2 SV=1 - [RSRC2_MOUSE] | 0,000E0 | 0,000E0 | 0,000E0 | 2,971E6 | 7,963E6 | 1,383E7 | Rsrc2 |
| Q9CXW2 | 28S ribosomal protein S22, mitochondrial OS=Mus musculus GN=Mrps22 PE=1 SV=1 - [RT22_MOUSE] | 0,000E0 | 0,000E0 | 0,000E0 | 3,939E6 | 3,983E6 | 7,501E6 | Mrps22 |
| O70305-2 | Isoform 2 of Ataxin-2 OS=Mus musculus GN=Atxn2 - [ATX2_MOUSE] | 0,000E0 | 0,000E0 | 0,000E0 | 5,520E6 | 3,783E6 | 1,268E7 | Atxn2 |

| | | | | | | | | |
|---|---|---|---|---|---|---|---|---|
| Q9ERG2 | Striatin-3 OS=Mus musculus GN=Strn3 PE=1 SV=1 - [STRN3_MOUSE] | 0,000E0 | 0,000E0 | 0,000E0 | 1,089E6 | 3,985E6 | 9,441E5 | Strn3 |
| Q8BWB6 | Metalloreductase STEAP2 OS=Mus musculus GN=Steap2 PE=1 SV=1 - [STEA2_MOUSE] | 0,000E0 | 0,000E0 | 0,000E0 | 3,532E6 | 7,136E6 | 7,036E6 | Steap2 |
| P70248 | Unconventional myosin-If OS=Mus musculus GN=Myo1f PE=1 SV=1 - [MYO1F_MOUSE] | 0,000E0 | 0,000E0 | 2,601E6 | 1,353E8 | 1,824E8 | 1,442E8 | Myo1f |
| O54988-2 | Isoform 2 of STE20-like serine/threonine-protein kinase OS=Mus musculus GN=Slk - [SLK_MOUSE] | 0,000E0 | 0,000E0 | 0,000E0 | 2,328E6 | 1,076E7 | 1,029E7 | Slk |
| Q7TN58 | Transmembrane channel-like protein 8 OS=Mus musculus GN=Tmc8 PE=1 SV=2 - [TMC8_MOUSE] | 0,000E0 | 0,000E0 | 0,000E0 | 5,571E6 | 8,204E6 | 1,124E7 | Tmc8 |
| Q8JZX4 | Splicing factor 45 OS=Mus musculus GN=Rbm17 PE=1 SV=1 - [SPF45_MOUSE] | 0,000E0 | 0,000E0 | 0,000E0 | 1,023E6 | 1,279E6 | 3,236E6 | Rbm17 |
| Q9DCU6 | 39S ribosomal protein L4, mitochondrial OS=Mus musculus GN=Mrpl4 PE=1 SV=1 - [RM04_MOUSE] | 0,000E0 | 0,000E0 | 0,000E0 | 1,707E6 | 3,613E6 | 1,972E6 | Mrpl4 |
| P60766 | Cell division control protein 42 homolog OS=Mus musculus GN=Cdc42 PE=1 SV=2 - [CDC42_MOUSE] | 0,000E0 | 0,000E0 | 0,000E0 | 3,260E6 | 1,599E7 | 4,871E6 | Cdc42 |
| Q921M3 | Splicing factor 3B subunit 3 OS=Mus musculus GN=Sf3b3 PE=1 SV=1 - [SF3B3_MOUSE] | 0,000E0 | 0,000E0 | 0,000E0 | 2,382E6 | 5,391E6 | 1,208E7 | Sf3b3 |
| Q8C5Q4 | G-rich sequence factor 1 OS=Mus musculus GN=Grsf1 PE=1 SV=2 - [GRSF1_MOUSE] | 0,000E0 | 0,000E0 | 0,000E0 | 1,194E6 | 2,171E6 | 8,444E5 | Grsf1 |
| L0N7N1 | Kinesin-like protein KIF14 OS=Mus musculus GN=Kif14 PE=1 SV=1 - [KIF14_MOUSE] | 0,000E0 | 0,000E0 | 0,000E0 | 2,803E6 | 3,500E6 | 1,129E7 | Kif14 |
| Q8CG79 | Apoptosis-stimulating of p53 protein 2 OS=Mus musculus GN=Tp53bp2 PE=1 SV=3 - [ASPP2_MOUSE] | 0,000E0 | 0,000E0 | 0,000E0 | 4,976E6 | 6,521E6 | 1,030E7 | Tp53bp2 |
| Q9D0J4 | ADP-ribosylation factor-like protein 2 OS=Mus musculus GN=Arl2 PE=1 SV=1 - [ARL2_MOUSE] | 0,000E0 | 0,000E0 | 0,000E0 | 3,752E6 | 7,184E6 | 4,523E6 | Arl2 |
| Q8CB44 | GRAM domain-containing protein 4 OS=Mus musculus GN=Gramd4 PE=1 SV=1 - [GRAM4_MOUSE] | 0,000E0 | 0,000E0 | 0,000E0 | 1,775E6 | 2,846E6 | 3,503E6 | Gramd4 |
| Q9DBR4 | Amyloid beta A4 precursor protein-binding family B member 2 OS=Mus musculus GN=Apbb2 PE=1 SV=2 - [APBB2_MOUSE] | 0,000E0 | 0,000E0 | 0,000E0 | 1,942E6 | 2,801E6 | 6,187E6 | Apbb2 |
| P13864 | DNA (cytosine-5)-methyltransferase 1 OS=Mus musculus GN=Dnmt1 PE=1 SV=5 - [DNMT1_MOUSE] | 0,000E0 | 0,000E0 | 0,000E0 | 2,100E6 | 7,445E6 | 1,556E7 | Dnmt1 |
| P58660 | Caspase recruitment domain-containing protein 10 OS=Mus musculus GN=Card10 PE=1 SV=1 - [CAR10_MOUSE] | 0,000E0 | 0,000E0 | 0,000E0 | 1,067E6 | 2,822E6 | 6,974E7 | Card10 |
| Q9Z0H3-2 | Isoform B of SWI/SNF-related matrix-associated actin-dependent regulator of chromatin subfamily B member 1 OS=Mus musculus GN=Smarcb1 - [SNF5_MOUSE] | 0,000E0 | 0,000E0 | 0,000E0 | 1,930E6 | 5,985E6 | 8,512E6 | Smarcb1 |
| Q9CQ10 | Charged multivesicular body protein 3 OS=Mus musculus GN=Chmp3 PE=1 SV=3 - [CHMP3_MOUSE] | 0,000E0 | 0,000E0 | 0,000E0 | 9,984E6 | 1,585E7 | 3,655E7 | Chmp3 |
| Q9R0X4 | Acyl-coenzyme A thioesterase 9, mitochondrial OS=Mus musculus GN=Acot9 PE=1 SV=1 - [ACOT9_MOUSE] | 0,000E0 | 0,000E0 | 0,000E0 | 2,974E6 | 5,397E6 | 4,126E6 | Acot9 |
| Q7TNP2 | Serine/threonine-protein phosphatase 2A 65 kDa regulatory subunit A beta isoform OS=Mus musculus GN=Ppp2r1b PE=1 SV=2 - [2AAB_MOUSE] | 0,000E0 | 0,000E0 | 0,000E0 | 1,239E7 | 9,631E6 | 6,012E6 | Ppp2r1b |
| Q64322 | Neural proliferation differentiation and control protein 1 OS=Mus musculus GN=Npdc1 PE=2 SV=2 - [NPDC1_MOUSE] | 0,000E0 | 0,000E0 | 0,000E0 | 4,126E6 | 1,073E7 | 1,187E7 | Npdc1 |
| O35379 | Multidrug resistance-associated protein 1 OS=Mus musculus GN=Abcc1 PE=1 SV=1 - [MRP1_MOUSE] | 0,000E0 | 0,000E0 | 0,000E0 | 2,495E6 | 5,537E6 | 7,573E6 | Abcc1 |
| Q9CQC9 | GTP-binding protein SAR1b OS=Mus musculus GN=Sar1b PE=1 SV=1 - [SAR1B_MOUSE] | 0,000E0 | 0,000E0 | 0,000E0 | 5,084E6 | 4,001E6 | 7,373E6 | Sar1b |
| Q9WUN2 | Serine/threonine-protein kinase TBK1 OS=Mus musculus GN=Tbk1 PE=1 SV=1 - [TBK1_MOUSE] | 0,000E0 | 0,000E0 | 0,000E0 | 1,266E6 | 5,612E6 | 1,006E7 | Tbk1 |
| Q9QY76 | Vesicle-associated membrane protein-associated protein B OS=Mus musculus GN=Vapb PE=1 SV=3 - [VAPB_MOUSE] | 0,000E0 | 0,000E0 | 0,000E0 | 2,272E6 | 7,010E6 | 1,196E7 | Vapb |
| P70349 | Histidine triad nucleotide-binding protein 1 OS=Mus musculus GN=Hint1 PE=1 SV=3 - [HINT1_MOUSE] | 0,000E0 | 0,000E0 | 0,000E0 | 1,951E7 | 4,461E7 | 7,495E7 | Hint1 |
| Q61292 | Laminin subunit beta-2 OS=Mus musculus GN=Lamb2 PE=1 SV=2 - [LAMB2_MOUSE] | 0,000E0 | 0,000E0 | 0,000E0 | 1,502E6 | 4,312E6 | 1,824E7 | Lamb2 |
| Q8BMS1 | Trifunctional enzyme subunit alpha, mitochondrial OS=Mus musculus GN=Hadha PE=1 SV=1 - [ECHA_MOUSE] | 0,000E0 | 0,000E0 | 0,000E0 | 5,197E6 | 5,934E6 | 1,326E7 | Hadha |
| P10493 | Nidogen-1 OS=Mus musculus GN=Nid1 PE=1 SV=2 - [NID1_MOUSE] | 0,000E0 | 0,000E0 | 0,000E0 | 2,374E6 | 1,150E7 | 2,464E7 | Nid1 |
| Q9CZX0 | Elongator complex protein 3 OS=Mus musculus GN=Elp3 PE=1 SV=1 - [ELP3_MOUSE] | 0,000E0 | 0,000E0 | 0,000E0 | 9,901E5 | 2,217E6 | 2,652E6 | Elp3 |
| Q6PFQ7 | Ras GTPase-activating protein 4 OS=Mus musculus GN=Rasa4 PE=1 SV=1 - [RASL2_MOUSE] | 0,000E0 | 0,000E0 | 0,000E0 | 1,237E6 | 5,838E6 | 1,404E7 | Rasa4 |
| Q8BG02 | Serine/threonine-protein phosphatase 2A 55 kDa regulatory subunit B gamma isoform OS=Mus musculus GN=Ppp2r2c PE=1 SV=1 - [2ABG_MOUSE] | 0,000E0 | 0,000E0 | 0,000E0 | 2,757E6 | 1,944E7 | 1,721E7 | Ppp2r2c |
| Q5XJY5 | Coatomer subunit delta OS=Mus musculus GN=Arcn1 PE=1 SV=2 - [COPD_MOUSE] | 0,000E0 | 0,000E0 | 0,000E0 | 1,114E6 | 1,134E6 | 6,827E6 | Arcn1 |
| P54761 | Ephrin type-B receptor 4 OS=Mus musculus GN=Ephb4 PE=1 SV=2 - [EPHB4_MOUSE] | 0,000E0 | 0,000E0 | 0,000E0 | 4,637E6 | 1,105E7 | 2,906E7 | Ephb4 |
| Q9JIF7 | Coatomer subunit beta OS=Mus musculus GN=Copb1 PE=1 SV=1 - [COPB_MOUSE] | 0,000E0 | 0,000E0 | 0,000E0 | 4,996E5 | 7,296E5 | 5,339E6 | Copb1 |
| Q8BJU2 | Tetraspanin-9 OS=Mus musculus GN=Tspan9 PE=1 SV=1 - [TSN9_MOUSE] | 0,000E0 | 0,000E0 | 0,000E0 | 2,053E6 | 6,829E6 | 9,509E6 | Tspan9 |
| Q571I4 | Tyrosine-protein kinase SgK223 OS=Mus musculus GN=Sgk223 PE=1 SV=2 - [SG223_MOUSE] | 0,000E0 | 0,000E0 | 0,000E0 | 1,445E6 | 5,808E6 | 6,616E6 | Sgk223 |
| Q5SFM8 | RNA-binding protein 27 OS=Mus musculus GN=Rbm27 PE=1 SV=3 - [RBM27_MOUSE] | 0,000E0 | 0,000E0 | 0,000E0 | 4,092E6 | 6,751E6 | 1,214E7 | Rbm27 |
| Q8BRN9 | Coiled-coil and C2 domain-containing protein 1B OS=Mus musculus GN=Cc2d1b PE=1 SV=1 - [C2D1B_MOUSE] | 0,000E0 | 0,000E0 | 0,000E0 | 2,454E6 | 6,912E6 | 1,554E6 | Cc2d1b |
| Q8BX17 | Gem-associated protein 5 OS=Mus musculus GN=Gemin5 PE=1 SV=2 - [GEMI5_MOUSE] | 0,000E0 | 0,000E0 | 0,000E0 | 2,525E6 | 3,809E6 | 9,602E6 | Gemin5 |
| Q61545 | RNA-binding protein EWS OS=Mus musculus GN=Ewsr1 PE=1 SV=2 - [EWS_MOUSE] | 0,000E0 | 0,000E0 | 0,000E0 | 1,766E7 | 1,874E7 | 3,023E7 | Ewsr1 |
| Q9CZD3 | Glycine--tRNA ligase OS=Mus musculus GN=Gars PE=1 SV=1 - [SYG_MOUSE] | 0,000E0 | 0,000E0 | 0,000E0 | 2,348E6 | 3,909E6 | 9,834E6 | Gars |
| P10922 | Histone H1.0 OS=Mus musculus GN=H1f0 PE=2 SV=4 - [H10_MOUSE] | 0,000E0 | 0,000E0 | 0,000E0 | 6,867E6 | 9,401E6 | 9,565E6 | H1f0 |
| Q8R550-5 | Isoform 5 of SH3 domain-containing kinase-binding protein 1 OS=Mus musculus GN=Sh3kbp1 - [SH3K1_MOUSE] | 0,000E0 | 0,000E0 | 0,000E0 | 1,205E6 | 2,662E6 | 1,878E6 | Sh3kbp1 |
| Q91YE6 | Importin-9 OS=Mus musculus GN=Ipo9 PE=1 SV=3 - [IPO9_MOUSE] | 0,000E0 | 0,000E0 | 0,000E0 | 6,400E6 | 9,136E6 | 7,160E6 | Ipo9 |
| O35149 | Zinc transporter 4 OS=Mus musculus GN=Slc30a4 PE=1 SV=2 - [ZNT4_MOUSE] | 0,000E0 | 0,000E0 | 0,000E0 | 3,931E6 | 4,028E6 | 1,718E6 | Slc30a4 |
| Q3V1V3 | ESF1 homolog OS=Mus musculus GN=Esf1 PE=1 SV=1 - [ESF1_MOUSE] | 0,000E0 | 0,000E0 | 0,000E0 | 5,764E6 | 9,590E6 | 3,536E6 | Esf1 |
| Q6URW6-2 | Isoform 2 of Myosin-14 OS=Mus musculus GN=Myh14 - [MYH14_MOUSE] | 2,336E8 | 0,000E0 | 5,102E7 | 2,921E9 | 6,979E9 | 5,047E9 | Myh14 |
| Q9R1Z8 | Vinexin OS=Mus musculus GN=Sorbs3 PE=1 SV=1 - [VINEX_MOUSE] | 0,000E0 | 0,000E0 | 0,000E0 | 8,258E6 | 1,907E7 | 3,088E7 | Sorbs3 |
| P70207 | Plexin-A2 OS=Mus musculus GN=Plxna2 PE=1 SV=2 - [PLXA2_MOUSE] | 0,000E0 | 0,000E0 | 0,000E0 | 6,187E6 | 8,567E6 | 1,411E7 | Plxna2 |
| Q32P12-2 | Isoform 2 of Uncharacterized protein C17orf53 homolog OS=Mus musculus - [CQ053_MOUSE] | 0,000E0 | 0,000E0 | 0,000E0 | 2,390E6 | 9,890E5 | 7,516E6 | Sep2 |
| Q9CXF4 | TBC1 domain family member 15 OS=Mus musculus GN=Tbc1d15 PE=1 SV=1 - [TBC15_MOUSE] | 0,000E0 | 0,000E0 | 0,000E0 | 1,078E6 | 2,489E6 | 4,261E6 | Tbc1d15 |
| O88554 | Poly [ADP-ribose] polymerase 2 OS=Mus musculus GN=Parp2 PE=1 SV=3 - [PARP2_MOUSE] | 0,000E0 | 0,000E0 | 0,000E0 | 9,644E5 | 3,483E6 | 8,206E5 | Parp2 |
| Q99LU0 | Charged multivesicular body protein 1b-1 OS=Mus musculus GN=Chmp1b1 PE=1 SV=1 - [CH1B1_MOUSE] | 0,000E0 | 0,000E0 | 0,000E0 | 1,233E7 | 3,477E7 | 1,166E7 | Chmp1b1 |
| Q9CZX5 | PIN2/TERF1-interacting telomerase inhibitor 1 OS=Mus musculus GN=Pinx1 PE=1 SV=2 - [PINX1_MOUSE] | 0,000E0 | 0,000E0 | 0,000E0 | 1,479E6 | 7,902E6 | 2,114E7 | Pinx1 |
| Q8R326-2 | Isoform 2 of Paraspeckle component 1 OS=Mus musculus GN=Pspc1 - [PSPC1_MOUSE] | 0,000E0 | 0,000E0 | 0,000E0 | 7,780E6 | 1,248E7 | 3,724E7 | Pspc1 |
| Q9QZM4 | Tumor necrosis factor receptor superfamily member 10B OS=Mus musculus GN=Tnfrsf10b PE=2 SV=3 - [TR10B_MOUSE] | 0,000E0 | 0,000E0 | 0,000E0 | 2,374E6 | 4,057E6 | 4,490E6 | Tnfrsf10b |
| Q8BWG8-2 | Isoform 1B of Beta-arrestin-1 OS=Mus musculus GN=Arrb1 - [ARRB1_MOUSE] | 0,000E0 | 0,000E0 | 0,000E0 | 2,430E6 | 5,763E6 | 1,077E7 | Arrb1 |
| Q9JLB0-2 | Isoform Alpha of MAGUK p55 subfamily member 6 OS=Mus musculus GN=Mpp6 - [MPP6_MOUSE] | 0,000E0 | 0,000E0 | 0,000E0 | 1,225E6 | 5,467E6 | 7,884E6 | Mpp6 |
| Q80XI6 | Mitogen-activated protein kinase kinase kinase 11 OS=Mus musculus GN=Map3k11 PE=1 SV=1 - [M3K11_MOUSE] | 0,000E0 | 0,000E0 | 0,000E0 | 4,111E5 | 4,143E6 | 5,980E6 | Map3k11 |
| Q9D2Y4-2 | Isoform 2 of Mixed lineage kinase domain-like protein OS=Mus musculus GN=Mlkl - [MLKL_MOUSE] | 0,000E0 | 0,000E0 | 0,000E0 | 1,071E6 | 2,731E6 | 2,986E6 | Mlkl |
| P12894 | Probable Pol polyprotein OS=Mouse intracisternal a-particle IL3 PE=3 SV=1 - [POL_IPMAI] | 0,000E0 | 0,000E0 | 0,000E0 | 1,981E6 | 3,566E6 | 4,052E6 | [POL |
| Q8BHF7 | CDP-diacylglycerol--glycerol-3-phosphate 3-phosphatidyltransferase, mitochondrial OS=Mus musculus GN=Pgs1 PE=1 SV=1 - [PGPS1_MOUSE] | 0,000E0 | 0,000E0 | 0,000E0 | 3,374E6 | 1,843E6 | 1,678E6 | Pgs1 |
| Q6PB44-2 | Isoform 2 of Tyrosine-protein phosphatase non-receptor type 23 OS=Mus musculus GN=Ptpn23 - [PTN23_MOUSE] | 0,000E0 | 0,000E0 | 0,000E0 | 2,543E6 | 5,850E6 | 9,455E6 | Ptpn23 |
| O35368 | Interferon-activable protein 203 OS=Mus musculus GN=Ifi203 PE=1 SV=1 - [IFI3_MOUSE] | 0,000E0 | 0,000E0 | 0,000E0 | 3,036E6 | 5,717E6 | 3,630E6 | Ifi203 |
| Q8K019-3 | Isoform 3 of Bcl-2-associated transcription factor 1 OS=Mus musculus GN=Bclaf1 - [BCLF1_MOUSE] | 0,000E0 | 0,000E0 | 0,000E0 | 6,968E5 | 1,412E6 | 6,045E6 | Bclaf1 |
| Q8BG95 | Protein phosphatase 1 regulatory subunit 12B OS=Mus musculus GN=Ppp1r12b PE=1 SV=2 - [MYPT2_MOUSE] | 0,000E0 | 0,000E0 | 0,000E0 | 6,992E7 | 9,881E7 | 1,253E8 | Ppp1r12b |
| O09172 | Glutamate--cysteine ligase regulatory subunit OS=Mus musculus GN=Gclm PE=1 SV=1 - [GSH0_MOUSE] | 0,000E0 | 0,000E0 | 0,000E0 | 2,737E6 | 6,511E6 | 6,788E6 | Gclm |
| Q9CX00 | IST1 homolog OS=Mus musculus GN=Ist1 PE=1 SV=1 - [IST1_MOUSE] | 0,000E0 | 0,000E0 | 0,000E0 | 4,093E6 | 7,738E6 | 1,106E7 | Ist1 |
| O09106 | Histone deacetylase 1 OS=Mus musculus GN=Hdac1 PE=1 SV=1 - [HDAC1_MOUSE] | 0,000E0 | 0,000E0 | 0,000E0 | 4,690E6 | 8,142E6 | 1,634E7 | Hdac1 |
| Q8BRT1-5 | Isoform 2 of CLIP-associating protein 2 OS=Mus musculus GN=Clasp2 - [CLAP2_MOUSE] | 0,000E0 | 0,000E0 | 0,000E0 | 4,562E5 | 3,844E6 | 1,018E7 | Clasp2 |
| Q80UY2-2 | Isoform 2 of E3 ubiquitin-protein ligase KCMF1 OS=Mus musculus GN=Kcmf1 - [KCMF1_MOUSE] | 0,000E0 | 0,000E0 | 0,000E0 | 5,965E6 | 9,529E6 | 2,306E7 | Kcmf1 |
| P51912 | Neutral amino acid transporter B(0) OS=Mus musculus GN=Slc1a5 PE=1 SV=2 - [AAAT_MOUSE] | 0,000E0 | 0,000E0 | 0,000E0 | 2,869E6 | 8,670E6 | 1,370E7 | Slc1a5 |
| P46460 | Vesicle-fusing ATPase OS=Mus musculus GN=Nsf PE=1 SV=2 - [NSF_MOUSE] | 0,000E0 | 0,000E0 | 0,000E0 | 1,126E6 | 4,179E6 | 2,901E6 | Nsf |
| Q9D0R4 | Probable ATP-dependent RNA helicase DDX56 OS=Mus musculus GN=Ddx56 PE=2 SV=1 - [DDX56_MOUSE] | 0,000E0 | 0,000E0 | 0,000E0 | 2,047E6 | 2,882E6 | 3,378E6 | Ddx56 |
| A2AGT5-2 | Isoform 2 of Cytoskeleton-associated protein 5 OS=Mus musculus GN=Ckap5 - [CKAP5_MOUSE] | 0,000E0 | 0,000E0 | 0,000E0 | 3,313E6 | 6,127E6 | 9,096E6 | Ckap5 |
| P70288 | Histone deacetylase 2 OS=Mus musculus GN=Hdac2 PE=1 SV=1 - [HDAC2_MOUSE] | 0,000E0 | 0,000E0 | 0,000E0 | 2,860E6 | 8,451E6 | 1,402E7 | Hdac2 |
| O35963 | Ras-related protein Rab-33B OS=Mus musculus GN=Rab33b PE=1 SV=1 - [RB33B_MOUSE] | 0,000E0 | 0,000E0 | 3,456E5 | 1,027E8 | 2,549E8 | 1,841E8 | Rab33b |
| P23475 | X-ray repair cross-complementing protein 6 OS=Mus musculus GN=Xrcc6 PE=1 SV=5 - [XRCC6_MOUSE] | 0,000E0 | 0,000E0 | 0,000E0 | 1,892E6 | 5,125E6 | 8,931E6 | Xrcc6 |
| Q8VC03 | Echinoderm microtubule-associated protein-like 3 OS=Mus musculus GN=Eml3 PE=1 SV=1 - [EMAL3_MOUSE] | 0,000E0 | 0,000E0 | 0,000E0 | 2,787E6 | 3,125E6 | 8,089E6 | Eml3 |
| Q3UDE2 | Tubulin--tyrosine ligase-like protein 12 OS=Mus musculus GN=Ttll12 PE=1 SV=1 - [TTL12_MOUSE] | 0,000E0 | 0,000E0 | 0,000E0 | 7,815E5 | 2,796E6 | 1,890E6 | Ttll12 |
| Q6ZPR6 | Inhibitor of Bruton tyrosine kinase OS=Mus musculus GN=Ibtk PE=1 SV=3 - [IBTK_MOUSE] | 0,000E0 | 0,000E0 | 0,000E0 | 1,539E6 | 3,276E6 | 6,271E6 | Ibtk |
| Q9CUN6 | E3 ubiquitin-protein ligase SMURF1 OS=Mus musculus GN=Smurf1 PE=1 SV=2 - [SMUF1_MOUSE] | 0,000E0 | 0,000E0 | 0,000E0 | 9,328E6 | 1,463E7 | 1,269E7 | Smurf1 |
| P39876 | Metalloproteinase inhibitor 3 OS=Mus musculus GN=Timp3 PE=1 SV=1 - [TIMP3_MOUSE] | 0,000E0 | 0,000E0 | 0,000E0 | 4,363E6 | 7,335E6 | 1,727E7 | Timp3 |
| P34152-3 | Isoform 3 of Focal adhesion kinase 1 OS=Mus musculus GN=Ptk2 - [FAK1_MOUSE] | 0,000E0 | 0,000E0 | 0,000E0 | 1,661E6 | 6,878E6 | 1,008E7 | Ptk2 |
| P40201 | Chromodomain-helicase-DNA-binding protein 1 OS=Mus musculus GN=Chd1 PE=1 SV=3 - [CHD1_MOUSE] | 0,000E0 | 0,000E0 | 0,000E0 | 3,349E6 | 1,313E6 | 2,135E6 | Chd1 |
| P97440 | Histone RNA hairpin-binding protein OS=Mus musculus GN=Slbp PE=1 SV=1 - [SLBP_MOUSE] | 0,000E0 | 0,000E0 | 0,000E0 | 4,769E6 | 1,498E6 | 6,832E5 | Slbp |
| Q8VCL2 | Protein SCO2 homolog, mitochondrial OS=Mus musculus GN=Sco2 PE=1 SV=1 - [SCO2_MOUSE] | 0,000E0 | 0,000E0 | 0,000E0 | 7,012E5 | 5,956E6 | 3,012E6 | Sco2 |
| Q6Q477 | Plasma membrane calcium-transporting ATPase 4 OS=Mus musculus GN=Atp2b4 PE=1 SV=1 - [AT2B4_MOUSE] | 0,000E0 | 0,000E0 | 0,000E0 | 8,105E6 | 9,547E6 | 1,788E7 | Atp2b4 |
| Q9ER72 | Cysteine--tRNA ligase, cytoplasmic OS=Mus musculus GN=Cars PE=1 SV=2 - [SYCC_MOUSE] | 0,000E0 | 0,000E0 | 0,000E0 | 1,909E6 | 6,499E6 | 1,030E7 | Cars |
| Q5SWD9 | Pre-rRNA-processing protein TSR1 homolog OS=Mus musculus GN=Tsr1 PE=1 SV=1 - [TSR1_MOUSE] | 0,000E0 | 0,000E0 | 0,000E0 | 3,056E6 | 1,380E6 | 1,192E7 | Tsr1 |
| P0DOV2 | Interferon-activable protein 204 OS=Mus musculus GN=Ifi204 PE=1 SV=1 - [IFI4_MOUSE] | 0,000E0 | 0,000E0 | 0,000E0 | 2,361E6 | 2,647E6 | 3,973E5 | Ifi204 |

| | | | | | | | | |
|---|---|---|---|---|---|---|---|---|
| Q9ERK4 | Exportin-2 OS=Mus musculus GN=Cse1l PE=1 SV=1 - [XPO2_MOUSE] | 0,000E0 | 0,000E0 | 0,000E0 | 2,980E6 | 7,246E6 | 9,950E6 | Cse1l |
| P11365 | Retrovirus-related Gag polyprotein OS=Mouse intracisternal a-particle MIA14 GN=gag PE=4 SV=2 - [GAG_IPMA] | 0,000E0 | 0,000E0 | 0,000E0 | 1,226E7 | 1,761E7 | 2,545E7 | gag |
| Q80X73 | Protein pelota homolog OS=Mus musculus GN=Pelo PE=1 SV=3 - [PELO_MOUSE] | 0,000E0 | 0,000E0 | 0,000E0 | 4,291E6 | 6,583E6 | 7,123E6 | Pelo |
| Q7TT37 | Elongator complex protein 1 OS=Mus musculus GN=Ikbkap PE=1 SV=2 - [ELP1_MOUSE] | 0,000E0 | 0,000E0 | 0,000E0 | 2,022E6 | 3,183E6 | 3,921E6 | Ikbkap |
| Q6PIX5 | Inactive rhomboid protein 1 OS=Mus musculus GN=Rhbdf1 PE=1 SV=2 - [RHDF1_MOUSE] | 0,000E0 | 0,000E0 | 0,000E0 | 3,128E6 | 6,078E6 | 6,945E6 | Rhbdf1 |
| Q62191 | E3 ubiquitin-protein ligase TRIM21 OS=Mus musculus GN=Trim21 PE=1 SV=1 - [RO52_MOUSE] | 0,000E0 | 0,000E0 | 1,073E6 | 1,027E7 | 4,860E6 | 5,318E6 | Trim21 |
| P35123 | Ubiquitin carboxyl-terminal hydrolase 4 OS=Mus musculus GN=Usp4 PE=1 SV=3 - [UBP4_MOUSE] | 0,000E0 | 0,000E0 | 0,000E0 | 3,696E5 | 1,566E6 | 3,832E6 | Usp4 |
| Q6PGL7-2 | Isoform 2 of WASH complex subunit 2 OS=Mus musculus GN=Washc2 - [WASC2_MOUSE] | 0,000E0 | 0,000E0 | 0,000E0 | 1,493E6 | 1,827E6 | 2,271E6 | Washc2 |
| Q99P72 | Reticulon-4 OS=Mus musculus GN=Rtn4 PE=1 SV=2 - [RTN4_MOUSE] | 0,000E0 | 0,000E0 | 0,000E0 | 6,569E6 | 6,637E6 | 2,184E7 | Rtn4 |
| O08919 | Numb-like protein OS=Mus musculus GN=Numbl PE=1 SV=3 - [NUMBL_MOUSE] | 0,000E0 | 0,000E0 | 0,000E0 | 7,608E6 | 9,439E6 | 1,152E7 | Numbl |
| O08582 | GTP-binding protein 1 OS=Mus musculus GN=Gtpbp1 PE=1 SV=2 - [GTPB1_MOUSE] | 0,000E0 | 0,000E0 | 0,000E0 | 1,521E6 | 6,914E6 | 2,708E6 | Gtpbp1 |
| Q8R3B1 | 1-phosphatidylinositol 4,5-bisphosphate phosphodiesterase delta-1 OS=Mus musculus GN=Plcd1 PE=1 SV=2 - [PLCD1_MOUSE] | 0,000E0 | 0,000E0 | 0,000E0 | 1,428E6 | 4,103E6 | 5,665E6 | Plcd1 |
| Q02858 | Angiopoietin-1 receptor OS=Mus musculus GN=Tek PE=1 SV=2 - [TIE2_MOUSE] | 0,000E0 | 0,000E0 | 0,000E0 | 7,184E6 | 8,964E6 | 1,756E7 | Tek |
| Q3TXS7 | 26S proteasome non-ATPase regulatory subunit 1 OS=Mus musculus GN=Psmd1 PE=1 SV=1 - [PSMD1_MOUSE] | 0,000E0 | 0,000E0 | 0,000E0 | 8,673E5 | 5,261E6 | 5,129E6 | Psmd1 |
| P07742 | Ribonucleoside-diphosphate reductase large subunit OS=Mus musculus GN=Rrm1 PE=1 SV=2 - [RIR1_MOUSE] | 0,000E0 | 0,000E0 | 0,000E0 | 3,894E6 | 1,027E7 | 1,104E7 | Rrm1 |
| P49962 | Signal recognition particle 9 kDa protein OS=Mus musculus GN=Srp9 PE=1 SV=2 - [SRP09_MOUSE] | 0,000E0 | 0,000E0 | 0,000E0 | 5,072E6 | 5,480E6 | 8,531E6 | Srp9 |
| Q9QWY8-2 | Isoform 2 of Arf-GAP with SH3 domain, ANK repeat and PH domain-containing protein 1 OS=Mus musculus GN=Asap1 - [ASAP1_MOUSE] | 0,000E0 | 0,000E0 | 0,000E0 | 8,234E5 | 4,151E6 | 4,461E6 | Asap1 |
| Q3UH93 | Plexin-D1 OS=Mus musculus GN=Plxnd1 PE=1 SV=1 - [PLXD1_MOUSE] | 0,000E0 | 0,000E0 | 0,000E0 | 6,711E6 | 1,528E7 | 2,068E7 | Plxnd1 |
| P32507 | Nectin-2 OS=Mus musculus GN=Nectin2 PE=1 SV=2 - [NECT2_MOUSE] | 0,000E0 | 0,000E0 | 0,000E0 | 1,861E6 | 5,041E5 | 9,776E5 | Nectin2 |
| Q8BKG3 | Inactive tyrosine-protein kinase 7 OS=Mus musculus GN=Ptk7 PE=1 SV=1 - [PTK7_MOUSE] | 0,000E0 | 0,000E0 | 0,000E0 | 7,024E6 | 1,317E7 | 4,140E7 | Ptk7 |
| Q8C7Q4 | RNA-binding protein 4 OS=Mus musculus GN=Rbm4 PE=1 SV=1 - [RBM4_MOUSE] | 0,000E0 | 0,000E0 | 0,000E0 | 7,181E6 | 7,197E6 | 6,225E6 | Rbm4 |
| P35283 | Ras-related protein Rab-12 OS=Mus musculus GN=Rab12 PE=1 SV=3 - [RAB12_MOUSE] | 0,000E0 | 0,000E0 | 3,456E5 | 1,027E8 | 2,549E8 | 3,658E8 | Rab12 |
| Q6PB66 | Leucine-rich PPR motif-containing protein, mitochondrial OS=Mus musculus GN=Lrpprc PE=1 SV=2 - [LPPRC_MOUSE] | 0,000E0 | 0,000E0 | 0,000E0 | 1,442E6 | 1,610E6 | 3,379E6 | Lrpprc |
| Q8R3P2 | Probable E3 ubiquitin-protein ligase DTX2 OS=Mus musculus GN=Dtx2 PE=1 SV=2 - [DTX2_MOUSE] | 0,000E0 | 0,000E0 | 0,000E0 | 2,626E6 | 3,255E6 | 8,966E5 | Dtx2 |
| A2APV2-2 | Isoform 2 of Formin-like protein 2 OS=Mus musculus GN=Fmnl2 - [FMNL2_MOUSE] | 0,000E0 | 0,000E0 | 0,000E0 | 5,462E6 | 4,438E6 | 8,570E6 | Fmnl2 |
| Q9JJA4 | Ribosome biogenesis protein WDR12 OS=Mus musculus GN=Wdr12 PE=1 SV=1 - [WDR12_MOUSE] | 0,000E0 | 0,000E0 | 0,000E0 | 6,465E5 | 8,510E5 | 2,441E6 | Wdr12 |
| Q6NSQ7 | Protein LTV1 homolog OS=Mus musculus GN=Ltv1 PE=1 SV=2 - [LTV1_MOUSE] | 0,000E0 | 0,000E0 | 0,000E0 | 2,306E6 | 1,807E6 | 2,596E6 | Ltv1 |
| Q8CB77 | Elongin-A OS=Mus musculus GN=Eloa PE=1 SV=3 - [ELOA1_MOUSE] | 0,000E0 | 0,000E0 | 0,000E0 | 2,946E6 | 4,660E6 | 1,078E7 | Eloa |
| Q99LI8 | Hepatocyte growth factor-regulated tyrosine kinase substrate OS=Mus musculus GN=Hgs PE=1 SV=2 - [HGS_MOUSE] | 0,000E0 | 0,000E0 | 0,000E0 | 9,902E6 | 1,667E7 | 2,330E7 | Hgs |
| Q9D832 | DnaJ homolog subfamily B member 4 OS=Mus musculus GN=Dnajb4 PE=1 SV=1 - [DNJB4_MOUSE] | 0,000E0 | 0,000E0 | 0,000E0 | 2,647E6 | 3,081E6 | 7,481E6 | Dnajb4 |
| Q8BRT1 | CLIP-associating protein 2 OS=Mus musculus GN=Clasp2 PE=1 SV=1 - [CLAP2_MOUSE] | 0,000E0 | 0,000E0 | 0,000E0 | 1,931E6 | 5,136E6 | 6,954E6 | Clasp2 |
| Q8R0A0 | General transcription factor IIF subunit 2 OS=Mus musculus GN=Gtf2f2 PE=1 SV=1 - [T2FB_MOUSE] | 0,000E0 | 0,000E0 | 0,000E0 | 1,547E6 | 2,185E6 | 6,953E6 | Gtf2f2 |
| Q9QZE5 | Coatomer subunit gamma-1 OS=Mus musculus GN=Copg1 PE=1 SV=1 - [COPG1_MOUSE] | 0,000E0 | 0,000E0 | 0,000E0 | 3,576E6 | 1,021E7 | 1,051E7 | Copg1 |
| O35593 | 26S proteasome non-ATPase regulatory subunit 14 OS=Mus musculus GN=Psmd14 PE=1 SV=2 - [PSDE_MOUSE] | 0,000E0 | 0,000E0 | 0,000E0 | 3,273E6 | 4,028E6 | 1,070E7 | Psmd14 |
| Q52KI8-2 | Isoform 2 of Serine/arginine repetitive matrix protein 1 OS=Mus musculus GN=Srrm1 - [SRRM1_MOUSE] | 0,000E0 | 0,000E0 | 0,000E0 | 3,972E6 | 5,575E5 | 2,050E7 | Srrm1 |
| Q6PAJ1 | Breakpoint cluster region protein OS=Mus musculus GN=Bcr PE=1 SV=3 - [BCR_MOUSE] | 0,000E0 | 0,000E0 | 0,000E0 | 1,494E6 | 1,814E6 | 3,458E6 | Bcr |
| P14211 | Calreticulin OS=Mus musculus GN=Calr PE=1 SV=1 - [CALR_MOUSE] | 0,000E0 | 0,000E0 | 0,000E0 | 2,109E6 | 9,833E6 | 1,378E7 | Calr |
| Q9CPP0 | Nucleoplasmin-3 OS=Mus musculus GN=Npm3 PE=1 SV=3 - [NPM3_MOUSE] | 0,000E0 | 0,000E0 | 0,000E0 | 1,052E7 | 3,717E7 | 8,792E7 | Npm3 |
| Q8BHJ9 | Pre-mRNA-splicing factor SLU7 OS=Mus musculus GN=Slu7 PE=1 SV=1 - [SLU7_MOUSE] | 0,000E0 | 0,000E0 | 0,000E0 | 3,418E6 | 5,502E6 | 1,383E6 | Slu7 |
| Q5ND28 | Scavenger receptor class F member 1 OS=Mus musculus GN=Scarf1 PE=1 SV=1 - [SREC_MOUSE] | 0,000E0 | 0,000E0 | 0,000E0 | 1,314E6 | 9,480E6 | 1,543E6 | Scarf1 |
| Q6NVF0-2 | Isoform 2 of Inositol polyphosphate 5-phosphatase OCRL-1 OS=Mus musculus GN=Ocrl - [OCRL_MOUSE] | 0,000E0 | 0,000E0 | 0,000E0 | 1,591E6 | 5,005E6 | 3,078E6 | Ocrl |
| Q8CAF4-3 | Isoform 3 of NHS-like protein 1 OS=Mus musculus GN=Nhsl1 - [NHSL1_MOUSE] | 0,000E0 | 0,000E0 | 0,000E0 | 2,637E6 | 5,915E6 | 2,842E7 | Nhsl1 |
| O54890 | Integrin beta-3 OS=Mus musculus GN=Itgb3 PE=1 SV=2 - [ITB3_MOUSE] | 0,000E0 | 0,000E0 | 0,000E0 | 7,201E6 | 7,933E6 | 2,257E7 | Itgb3 |
| Q640N1 | Adipocyte enhancer-binding protein 1 OS=Mus musculus GN=Aebp1 PE=1 SV=1 - [AEBP1_MOUSE] | 0,000E0 | 0,000E0 | 0,000E0 | 1,332E6 | 3,504E6 | 7,733E6 | Aebp1 |
| Q9ES52-3 | Isoform 3 of Phosphatidylinositol 3,4,5-trisphosphate 5-phosphatase 1 OS=Mus musculus GN=Inpp5d - [SHIP1_MOUSE] | 0,000E0 | 0,000E0 | 0,000E0 | 2,468E6 | 2,990E6 | 3,909E6 | Inpp5d |
| Q9JKP5 | Muscleblind-like protein 1 OS=Mus musculus GN=Mbnl1 PE=1 SV=1 - [MBNL1_MOUSE] | 0,000E0 | 0,000E0 | 0,000E0 | 9,110E5 | 9,671E6 | 7,498E6 | Mbnl1 |
| Q80W68 | Kin of IRRE-like protein 1 OS=Mus musculus GN=Kirrel PE=1 SV=1 - [KIRR1_MOUSE] | 0,000E0 | 0,000E0 | 0,000E0 | 3,203E6 | 1,306E7 | 2,271E7 | Kirrel |
| Q9D662 | Protein transport protein Sec23B OS=Mus musculus GN=Sec23b PE=1 SV=1 - [SC23B_MOUSE] | 0,000E0 | 0,000E0 | 0,000E0 | 4,185E6 | 5,073E6 | 6,851E6 | Sec23b |
| P23949 | mRNA decay activator protein ZFP36L2 OS=Mus musculus GN=Zfp36l2 PE=1 SV=2 - [TISD_MOUSE] | 0,000E0 | 0,000E0 | 0,000E0 | 2,162E6 | 1,397E7 | 5,862E6 | Zfp36l2 |
| Q9EPJ9 | ADP-ribosylation factor GTPase-activating protein 1 OS=Mus musculus GN=Arfgap1 PE=1 SV=2 - [ARFG1_MOUSE] | 0,000E0 | 0,000E0 | 0,000E0 | 1,061E6 | 2,219E6 | 4,447E6 | Arfgap1 |
| Q9CY27 | Very-long-chain enoyl-CoA reductase OS=Mus musculus GN=Tecr PE=1 SV=1 - [TECR_MOUSE] | 0,000E0 | 0,000E0 | 0,000E0 | 1,247E7 | 2,198E7 | 1,537E7 | Tecr |
| Q8BMJ2 | Leucine--tRNA ligase, cytoplasmic OS=Mus musculus GN=Lars PE=1 SV=2 - [SYLC_MOUSE] | 0,000E0 | 0,000E0 | 0,000E0 | 7,083E5 | 3,374E6 | 6,364E6 | Lars |
| Q9R1Q6 | Transmembrane protein 176B OS=Mus musculus GN=Tmem176b PE=1 SV=1 - [T176B_MOUSE] | 0,000E0 | 0,000E0 | 0,000E0 | 3,373E6 | 4,638E6 | 7,472E6 | Tmem176b |
| E9Q557 | Desmoplakin OS=Mus musculus GN=Dsp PE=1 SV=1 - [DESP_MOUSE] | 0,000E0 | 0,000E0 | 8,274E6 | 5,286E6 | 1,965E6 | 2,370E6 | Dsp |
| Q6ZPZ3-2 | Isoform 2 of Zinc finger CCCH domain-containing protein 4 OS=Mus musculus GN=Zc3h4 - [ZC3H4_MOUSE] | 0,000E0 | 0,000E0 | 0,000E0 | 2,382E6 | 3,241E6 | 1,269E7 | Zc3h4 |
| Q76LS9-2 | Isoform 2 of Ubiquitin carboxyl-terminal hydrolase MINDY-1 OS=Mus musculus GN=Fam63a - [FA63A_MOUSE] | 0,000E0 | 0,000E0 | 0,000E0 | 4,213E6 | 1,147E6 | 6,166E6 | Fam63a |
| Q61214 | Dual specificity tyrosine-phosphorylation-regulated kinase 1A OS=Mus musculus GN=Dyrk1a PE=1 SV=1 - [DYR1A_MOUSE] | 0,000E0 | 0,000E0 | 0,000E0 | 1,151E6 | 4,814E6 | 9,037E6 | Dyrk1a |
| P35822 | Receptor-type tyrosine-protein phosphatase kappa OS=Mus musculus GN=Ptprk PE=1 SV=1 - [PTPRK_MOUSE] | 0,000E0 | 0,000E0 | 0,000E0 | 3,513E6 | 4,003E6 | 1,329E7 | Ptprk |
| Q9JKY5 | Huntingtin-interacting protein 1-related protein OS=Mus musculus GN=Hip1r PE=1 SV=2 - [HIP1R_MOUSE] | 0,000E0 | 0,000E0 | 0,000E0 | 1,761E7 | 4,116E7 | 2,839E6 | Hip1r |
| Q6PFD9 | Nuclear pore complex protein Nup98-Nup96 OS=Mus musculus GN=Nup98 PE=1 SV=2 - [NUP98_MOUSE] | 0,000E0 | 0,000E0 | 0,000E0 | 1,011E6 | 3,277E6 | 6,815E6 | Nup98 |
| Q5FWK3 | Rho GTPase-activating protein 1 OS=Mus musculus GN=Arhgap1 PE=1 SV=1 - [RHG01_MOUSE] | 0,000E0 | 0,000E0 | 0,000E0 | 1,877E6 | 8,360E5 | 7,619E6 | Arhgap1 |
| Q8K3W3 | Protein CASC3 OS=Mus musculus GN=Casc3 PE=1 SV=3 - [CASC3_MOUSE] | 0,000E0 | 0,000E0 | 0,000E0 | 6,702E6 | 1,400E7 | 1,137E7 | Casc3 |
| Q60805 | Tyrosine-protein kinase Mer OS=Mus musculus GN=Mertk PE=1 SV=1 - [MERTK_MOUSE] | 0,000E0 | 0,000E0 | 0,000E0 | 9,170E6 | 4,119E6 | 1,162E7 | Mertk |
| Q921Y2 | U3 small nucleolar ribonucleoprotein protein IMP3 OS=Mus musculus GN=Imp3 PE=2 SV=1 - [IMP3_MOUSE] | 0,000E0 | 0,000E0 | 0,000E0 | 4,491E6 | 1,072E7 | 1,033E7 | Imp3 |
| P02468 | Laminin subunit gamma-1 OS=Mus musculus GN=Lamc1 PE=1 SV=2 - [LAMC1_MOUSE] | 0,000E0 | 0,000E0 | 0,000E0 | 1,063E6 | 3,982E6 | 1,833E7 | Lamc1 |
| Q8CJ19 | [F-actin]-methionine sulfoxide oxidase MICAL3 OS=Mus musculus GN=Mical3 PE=1 SV=2 - [MICA3_MOUSE] | 0,000E0 | 0,000E0 | 0,000E0 | 2,930E6 | 4,563E6 | 6,390E6 | Mical3 |
| Q8BHD8 | Protein-L-isoaspartate O-methyltransferase domain-containing protein 2 OS=Mus musculus GN=Pcmtd2 PE=2 SV=1 - [PCMD2_MOUSE] | 0,000E0 | 0,000E0 | 0,000E0 | 7,963E5 | 2,480E6 | 1,124E6 | Pcmtd2 |
| Q9CQN1 | Heat shock protein 75 kDa, mitochondrial OS=Mus musculus GN=Trap1 PE=1 SV=1 - [TRAP1_MOUSE] | 0,000E0 | 0,000E0 | 0,000E0 | 9,790E7 | 1,385E8 | 2,929E8 | Trap1 |
| Q80UZ0 | FYVE, RhoGEF and PH domain-containing protein 5 OS=Mus musculus GN=Fgd5 PE=1 SV=2 - [FGD5_MOUSE] | 0,000E0 | 0,000E0 | 0,000E0 | 4,539E6 | 7,869E6 | 4,507E6 | Fgd5 |
| Q920B9 | FACT complex subunit SPT16 OS=Mus musculus GN=Supt16h PE=1 SV=2 - [SP16H_MOUSE] | 0,000E0 | 0,000E0 | 0,000E0 | 2,583E6 | 2,788E6 | 2,679E7 | Supt16h |
| Q91V81 | RNA-binding protein 42 OS=Mus musculus GN=Rbm42 PE=1 SV=1 - [RBM42_MOUSE] | 0,000E0 | 0,000E0 | 0,000E0 | 1,869E6 | 1,776E6 | 2,204E6 | Rbm42 |
| Q68FE6 | Protein FAM65A OS=Mus musculus GN=Fam65a PE=1 SV=2 - [FA65A_MOUSE] | 0,000E0 | 0,000E0 | 0,000E0 | 2,239E6 | 5,659E6 | 6,782E6 | Fam65a |
| Q5SSZ5-2 | Isoform 2 of Tensin-3 OS=Mus musculus GN=Tns3 - [TENS3_MOUSE] | 0,000E0 | 0,000E0 | 0,000E0 | 3,158E7 | 1,846E6 | 4,529E7 | Tns3 |
| P70227 | Inositol 1,4,5-trisphosphate receptor type 3 OS=Mus musculus GN=Itpr3 PE=1 SV=3 - [ITPR3_MOUSE] | 0,000E0 | 0,000E0 | 0,000E0 | 4,207E6 | 7,277E6 | 1,349E7 | Itpr3 |
| Q9DAX2 | Phospholipid phosphatase 2 OS=Mus musculus GN=Plpp2 PE=1 SV=1 - [PLPP2_MOUSE] | 0,000E0 | 0,000E0 | 0,000E0 | 6,375E5 | 3,376E6 | 6,159E6 | Plpp2 |
| Q00560 | Interleukin-6 receptor subunit beta OS=Mus musculus GN=Il6st PE=1 SV=2 - [IL6RB_MOUSE] | 0,000E0 | 0,000E0 | 0,000E0 | 1,181E6 | 3,818E6 | 5,858E6 | Il6st |
| Q00612 | Glucose-6-phosphate 1-dehydrogenase X OS=Mus musculus GN=G6pdx PE=1 SV=3 - [G6PD1_MOUSE] | 0,000E0 | 0,000E0 | 0,000E0 | 1,193E6 | 4,309E6 | 2,511E6 | G6pdx |
| P51432 | 1-phosphatidylinositol 4,5-bisphosphate phosphodiesterase beta-3 OS=Mus musculus GN=Plcb3 PE=1 SV=2 - [PLCB3_MOUSE] | 0,000E0 | 0,000E0 | 0,000E0 | 2,362E6 | 4,868E6 | 7,343E6 | Plcb3 |
| O35680 | 28S ribosomal protein S12, mitochondrial OS=Mus musculus GN=Mrps12 PE=2 SV=1 - [RT12_MOUSE] | 0,000E0 | 0,000E0 | 0,000E0 | 5,814E6 | 9,954E6 | 7,431E6 | Mrps12 |
| Q8R5G7 | Arf-GAP with Rho-GAP domain, ANK repeat and PH domain-containing protein 3 OS=Mus musculus GN=Arap3 PE=1 SV=3 - [ARAP3_MOUSE] | 0,000E0 | 0,000E0 | 0,000E0 | 1,388E6 | 3,386E6 | 4,623E6 | Arap3 |
| A2RT67 | DENN domain-containing protein 3 OS=Mus musculus GN=Dennd3 PE=1 SV=2 - [DEND3_MOUSE] | 0,000E0 | 0,000E0 | 0,000E0 | 1,304E6 | 2,968E6 | 2,659E6 | Dennd3 |
| Q8R010 | Aminoacyl tRNA synthase complex-interacting multifunctional protein 2 OS=Mus musculus GN=Aimp2 PE=1 SV=2 - [AIMP2_MOUSE] | 0,000E0 | 0,000E0 | 0,000E0 | 4,011E6 | 2,240E6 | 4,491E6 | Aimp2 |
| Q8R4Y4-2 | Isoform 2 of Stabilin-1 OS=Mus musculus GN=Stab1 - [STAB1_MOUSE] | 0,000E0 | 0,000E0 | 0,000E0 | 2,266E7 | 2,584E7 | 4,512E7 | Stab1 |
| B1AXH1 | NHS-like protein 2 OS=Mus musculus GN=Nhsl2 PE=1 SV=1 - [NHSL2_MOUSE] | 0,000E0 | 0,000E0 | 0,000E0 | 5,183E6 | 9,820E6 | 3,006E6 | Nhsl2 |
| A2A4P0 | ATP-dependent RNA helicase DHX8 OS=Mus musculus GN=Dhx8 PE=2 SV=1 - [DHX8_MOUSE] | 0,000E0 | 0,000E0 | 0,000E0 | 4,776E6 | 7,091E6 | 1,020E7 | Dhx8 |
| Q02105 | Complement C1q subcomponent subunit C OS=Mus musculus GN=C1qc PE=1 SV=2 - [C1QC_MOUSE] | 0,000E0 | 0,000E0 | 0,000E0 | 1,832E7 | 3,056E8 | 1,616E8 | C1qc |
| O54734 | Dolichyl-diphosphooligosaccharide--protein glycosyltransferase 48 kDa subunit OS=Mus musculus GN=Ddost PE=1 SV=2 - [OST48_MOUSE] | 0,000E0 | 0,000E0 | 0,000E0 | 1,655E6 | 2,711E6 | 3,503E6 | Ddost |
| Q3TTA7-2 | Isoform 2 of E3 ubiquitin-protein ligase CBL-B OS=Mus musculus GN=Cblb - [CBLB_MOUSE] | 0,000E0 | 0,000E0 | 0,000E0 | 1,702E6 | 2,150E6 | 2,271E6 | Cblb |
| Q80Z25 | Oral-facial-digital syndrome 1 protein homolog OS=Mus musculus GN=Ofd1 PE=1 SV=1 - [OFD1_MOUSE] | 0,000E0 | 0,000E0 | 0,000E0 | 6,329E5 | 1,646E6 | 4,614E7 | Ofd1 |
| Q9JII5-2 | Isoform 2 of DAZ-associated protein 1 OS=Mus musculus GN=Dazap1 - [DAZP1_MOUSE] | 0,000E0 | 0,000E0 | 0,000E0 | 5,965E6 | 7,044E6 | 1,613E7 | Dazap1 |
| Q6DVA0 | LEM domain-containing protein 2 OS=Mus musculus GN=Lemd2 PE=1 SV=1 - [LEMD2_MOUSE] | 0,000E0 | 0,000E0 | 0,000E0 | 8,142E5 | 2,401E6 | 9,327E5 | Lemd2 |
| B2RU80 | Receptor-type tyrosine-protein phosphatase beta OS=Mus musculus GN=Ptprb PE=1 SV=1 - [PTPRB_MOUSE] | 0,000E0 | 0,000E0 | 0,000E0 | 2,030E6 | 1,113E7 | 1,904E7 | Ptprb |
| Q9CZD5 | Translation initiation factor IF-3, mitochondrial OS=Mus musculus GN=Mtif3 PE=1 SV=1 - [IF3M_MOUSE] | 0,000E0 | 0,000E0 | 0,000E0 | 2,908E6 | 7,095E6 | 1,633E7 | Mtif3 |

| | | | | | | | | |
|---|---|---|---|---|---|---|---|---|
| Q01705-4 | Isoform 4 of Neurogenic locus notch homolog protein 1 OS=Mus musculus GN=Notch1 - [NOTC1_MOUSE] | 0,000E0 | 0,000E0 | 0,000E0 | 1,475E7 | 2,933E7 | 1,564E7 | Notch1 |
| Q8JZW4 | Copine-5 OS=Mus musculus GN=Cpne5 PE=1 SV=1 - [CPNE5_MOUSE] | 0,000E0 | 0,000E0 | 0,000E0 | 2,218E6 | 3,588E6 | 7,389E6 | Cpne5 |
| Q9QXZ0 | Microtubule-actin cross-linking factor 1 OS=Mus musculus GN=Macf1 PE=1 SV=2 - [MACF1_MOUSE] | 0,000E0 | 0,000E0 | 0,000E0 | 4,223E6 | 1,297E8 | 1,338E8 | Macf1 |
| Q80U78-2 | Isoform 2 of Pumilio homolog 1 OS=Mus musculus GN=Pum1 - [PUM1_MOUSE] | 0,000E0 | 0,000E0 | 0,000E0 | 1,253E6 | 6,252E5 | 8,476E6 | Pum1 |
| Q07139-2 | Isoform 2 of Protein ECT2 OS=Mus musculus GN=Ect2 - [ECT2_MOUSE] | 0,000E0 | 0,000E0 | 0,000E0 | 5,244E5 | 1,416E6 | 2,241E6 | Ect2 |
| P61079 | Ubiquitin-conjugating enzyme E2 D3 OS=Mus musculus GN=Ube2d3 PE=1 SV=1 - [UB2D3_MOUSE] | 0,000E0 | 0,000E0 | 0,000E0 | 1,082E7 | 3,589E7 | 5,207E7 | Ube2d3 |
| Q8C5L3-3 | Isoform 3 of CCR4-NOT transcription complex subunit 2 OS=Mus musculus GN=Cnot2 - [CNOT2_MOUSE] | 0,000E0 | 0,000E0 | 0,000E0 | 1,973E6 | 8,405E6 | 1,798E7 | Cnot2 |
| Q91WM1-2 | Isoform 2 of Spermatid perinuclear RNA-binding protein OS=Mus musculus GN=Strbp - [STRBP_MOUSE] | 0,000E0 | 0,000E0 | 0,000E0 | 1,346E7 | 9,306E6 | 2,199E7 | Strbp |
| O35660 | Glutathione S-transferase Mu 6 OS=Mus musculus GN=Gstm6 PE=1 SV=4 - [GSTM6_MOUSE] | 0,000E0 | 0,000E0 | 0,000E0 | 5,973E5 | 4,301E5 | 5,492E5 | Gstm6 |
| Q99MS7 | EH domain-binding protein 1-like protein 1 OS=Mus musculus GN=Ehbp1l1 PE=1 SV=1 - [EH1L1_MOUSE] | 0,000E0 | 0,000E0 | 0,000E0 | 3,964E6 | 9,571E6 | 1,144E7 | Ehbp1l1 |
| P23463 | Homeobox protein Hox-D8 OS=Mus musculus GN=Hoxd8 PE=3 SV=4 - [HXD8_MOUSE] | 0,000E0 | 0,000E0 | 0,000E0 | 2,880E6 | 7,146E6 | 4,389E6 | Hoxd8 |
| P83093 | Stromal interaction molecule 2 OS=Mus musculus GN=Stim2 PE=1 SV=2 - [STIM2_MOUSE] | 0,000E0 | 0,000E0 | 0,000E0 | 1,603E6 | 2,805E6 | 2,647E6 | Stim2 |
| Q61001 | Laminin subunit alpha-5 OS=Mus musculus GN=Lama5 PE=1 SV=4 - [LAMA5_MOUSE] | 0,000E0 | 0,000E0 | 0,000E0 | 1,300E6 | 4,150E6 | 1,434E7 | Lama5 |
| P35821 | Tyrosine-protein phosphatase non-receptor type 1 OS=Mus musculus GN=Ptpn1 PE=1 SV=2 - [PTN1_MOUSE] | 0,000E0 | 0,000E0 | 0,000E0 | 8,743E5 | 1,694E6 | 3,833E6 | Ptpn1 |
| Q80SW1 | Putative adenosylhomocysteinase 2 OS=Mus musculus GN=Ahcyl1 PE=1 SV=1 - [SAHH2_MOUSE] | 0,000E0 | 0,000E0 | 0,000E0 | 2,795E6 | 2,412E6 | 2,775E6 | Ahcyl1 |
| Q9DC69 | NADH dehydrogenase [ubiquinone] 1 alpha subcomplex subunit 9, mitochondrial OS=Mus musculus GN=Ndufa9 PE=1 SV=2 - [NDUA9_MOUSE] | 0,000E0 | 0,000E0 | 0,000E0 | 1,643E6 | 2,321E6 | 3,761E6 | Ndufa9 |
| Q99KN1-2 | Isoform 2 of Arrestin domain-containing protein 1 OS=Mus musculus GN=Arrdc1 - [ARRD1_MOUSE] | 0,000E0 | 0,000E0 | 0,000E0 | 9,543E5 | 4,932E6 | 6,451E6 | Arrdc1 |
| Q80Y56 | Rabenosyn-5 OS=Mus musculus GN=Rbsn PE=1 SV=1 - [RBNS5_MOUSE] | 0,000E0 | 0,000E0 | 0,000E0 | 9,590E5 | 1,130E6 | 1,492E6 | Rbsn |
| Q64430 | Copper-transporting ATPase 1 OS=Mus musculus GN=Atp7a PE=1 SV=3 - [ATP7A_MOUSE] | 0,000E0 | 0,000E0 | 0,000E0 | 1,294E6 | 2,214E6 | 3,588E6 | Atp7a |
| Q8BG79 | CWF19-like protein 2 OS=Mus musculus GN=Cwf19l2 PE=1 SV=1 - [C19L2_MOUSE] | 0,000E0 | 0,000E0 | 0,000E0 | 7,675E5 | 1,405E6 | 5,503E6 | Cwf19l2 |
| P35918 | Vascular endothelial growth factor receptor 2 OS=Mus musculus GN=Kdr PE=1 SV=1 - [VGFR2_MOUSE] | 0,000E0 | 0,000E0 | 0,000E0 | 1,139E7 | 9,802E6 | 2,165E7 | Kdr |
| Q8BZX4 | Splicing regulatory glutamine/lysine-rich protein 1 OS=Mus musculus GN=Srek1 PE=2 SV=1 - [SREK1_MOUSE] | 0,000E0 | 0,000E0 | 0,000E0 | 1,378E6 | 5,429E5 | 6,565E6 | Srek1 |
| Q63850 | Nuclear pore glycoprotein p62 OS=Mus musculus GN=Nup62 PE=1 SV=2 - [NUP62_MOUSE] | 0,000E0 | 0,000E0 | 0,000E0 | 1,735E6 | 2,664E6 | 3,891E6 | Nup62 |
| Q8BKI2 | Trinucleotide repeat-containing gene 6B protein OS=Mus musculus GN=Tnrc6b PE=1 SV=2 - [TNR6B_MOUSE] | 0,000E0 | 0,000E0 | 0,000E0 | 2,376E6 | 3,240E6 | 2,598E6 | Tnrc6b |
| P46664 | Adenylosuccinate synthetase isozyme 2 OS=Mus musculus GN=Adss PE=1 SV=2 - [PURA2_MOUSE] | 0,000E0 | 0,000E0 | 0,000E0 | 2,239E6 | 3,514E6 | 7,118E6 | Adss |
| Q8BM55 | Transmembrane protein 214 OS=Mus musculus GN=Tmem214 PE=1 SV=1 - [TM214_MOUSE] | 0,000E0 | 0,000E0 | 0,000E0 | 2,949E6 | 3,795E6 | 5,973E6 | Tmem214 |
| Q9D2R0 | Acetoacetyl-CoA synthetase OS=Mus musculus GN=Aacs PE=1 SV=1 - [AACS_MOUSE] | 0,000E0 | 0,000E0 | 0,000E0 | 1,939E6 | 1,747E6 | 4,128E6 | Aacs |
| Q9Z1T1 | AP-3 complex subunit beta-1 OS=Mus musculus GN=Ap3b1 PE=1 SV=2 - [AP3B1_MOUSE] | 0,000E0 | 0,000E0 | 0,000E0 | 3,649E6 | 6,529E6 | 5,274E6 | Ap3b1 |
| A2AAY5 | SH3 and PX domain-containing protein 2B OS=Mus musculus GN=Sh3pxd2b PE=1 SV=1 - [SPD2B_MOUSE] | 0,000E0 | 0,000E0 | 0,000E0 | 1,672E6 | 3,423E6 | 4,022E6 | Sh3pxd2b |
| Q8K2B3 | Succinate dehydrogenase [ubiquinone] flavoprotein subunit, mitochondrial OS=Mus musculus GN=Sdha PE=1 SV=1 - [SDHA_MOUSE] | 0,000E0 | 0,000E0 | 0,000E0 | 3,018E6 | 4,687E6 | 2,565E6 | Sdha |
| Q3UFM5 | Nucleolar MIF4G domain-containing protein 1 OS=Mus musculus GN=Nom1 PE=1 SV=2 - [NOM1_MOUSE] | 0,000E0 | 0,000E0 | 0,000E0 | 4,975E5 | 5,572E6 | 6,752E6 | Nom1 |
| Q9D706 | RNA polymerase II-associated protein 3 OS=Mus musculus GN=Rpap3 PE=1 SV=1 - [RPAP3_MOUSE] | 0,000E0 | 0,000E0 | 0,000E0 | 3,438E6 | 5,609E6 | 1,368E7 | Rpap3 |
| Q5I1X5 | RelA-associated inhibitor OS=Mus musculus GN=Ppp1r13l PE=1 SV=1 - [IASPP_MOUSE] | 0,000E0 | 0,000E0 | 0,000E0 | 8,316E5 | 5,868E5 | 3,700E6 | Ppp1r13l |
| Q6ZPE2-2 | Isoform 2 of Myotubularin-related protein 5 OS=Mus musculus GN=Sbf1 - [MTMR5_MOUSE] | 0,000E0 | 0,000E0 | 0,000E0 | 1,716E6 | 1,606E6 | 3,705E6 | Sbf1 |
| Q3UPH7-3 | Isoform 3 of Rho guanine nucleotide exchange factor 40 OS=Mus musculus GN=Arhgef40 - [ARH40_MOUSE] | 0,000E0 | 0,000E0 | 0,000E0 | 2,345E6 | 6,154E6 | 8,629E6 | Arhgef40 |
| Q8BX57-3 | Isoform 3 of PX domain-containing protein kinase-like protein OS=Mus musculus GN=Pxk - [PXK_MOUSE] | 0,000E0 | 0,000E0 | 0,000E0 | 1,065E6 | 2,147E6 | 2,671E6 | Pxk |
| Q8CIP4-2 | Isoform 2 of MAP/microtubule affinity-regulating kinase 4 OS=Mus musculus GN=Mark4 - [MARK4_MOUSE] | 0,000E0 | 0,000E0 | 0,000E0 | 9,286E6 | 1,155E7 | 1,407E7 | Mark4 |
| Q80U93 | Nuclear pore complex protein Nup214 OS=Mus musculus GN=Nup214 PE=1 SV=2 - [NU214_MOUSE] | 0,000E0 | 0,000E0 | 0,000E0 | 5,193E5 | 2,763E6 | 5,939E6 | Nup214 |
| Q6ZQ38 | Cullin-associated NEDD8-dissociated protein 1 OS=Mus musculus GN=Cand1 PE=1 SV=2 - [CAND1_MOUSE] | 0,000E0 | 0,000E0 | 0,000E0 | 1,241E6 | 3,688E6 | 3,377E6 | Cand1 |
| Q80SU7 | Interferon-induced very large GTPase 1 OS=Mus musculus GN=Gvin1 PE=1 SV=1 - [GVIN1_MOUSE] | 0,000E0 | 0,000E0 | 0,000E0 | 1,487E6 | 3,487E6 | 8,673E6 | Gvin1 |
| Q99LI5 | Zinc finger protein 281 OS=Mus musculus GN=Znf281 PE=1 SV=1 - [ZN281_MOUSE] | 0,000E0 | 0,000E0 | 0,000E0 | 2,058E6 | 4,160E6 | 5,764E6 | Znf281 |
| Q9R0U0-3 | Isoform 3 of Serine/arginine-rich splicing factor 10 OS=Mus musculus GN=Srsf10 - [SRS10_MOUSE] | 0,000E0 | 0,000E0 | 0,000E0 | 1,277E6 | 5,243E6 | 1,221E7 | Srsf10 |
| P47226-2 | Isoform TES1 of Testin OS=Mus musculus GN=Tes - [TES_MOUSE] | 0,000E0 | 0,000E0 | 0,000E0 | 6,851E5 | 2,694E6 | 3,421E6 | Tes |
| Q6ZQ73 | Cullin-associated NEDD8-dissociated protein 2 OS=Mus musculus GN=Cand2 PE=1 SV=2 - [CAND2_MOUSE] | 0,000E0 | 0,000E0 | 0,000E0 | 2,120E6 | 6,679E6 | 8,745E6 | Cand2 |
| Q8BP22 | Protein FAM92A1 OS=Mus musculus GN=Fam92a1 PE=1 SV=1 - [F92A1_MOUSE] | 0,000E0 | 0,000E0 | 0,000E0 | 1,141E6 | 1,944E6 | 6,828E6 | Fam92a1 |
| P14873 | Microtubule-associated protein 1B OS=Mus musculus GN=Map1b PE=1 SV=2 - [MAP1B_MOUSE] | 0,000E0 | 0,000E0 | 2,559E6 | 2,404E6 | 4,708E6 | 5,312E6 | Map1b |
| Q8C0D4 | Rho GTPase-activating protein 12 OS=Mus musculus GN=Arhgap12 PE=1 SV=2 - [RHG12_MOUSE] | 0,000E0 | 0,000E0 | 0,000E0 | 1,616E6 | 4,145E6 | 1,531E7 | Arhgap12 |
| Q8BU30 | Isoleucine--tRNA ligase, cytoplasmic OS=Mus musculus GN=Iars PE=1 SV=2 - [SYIC_MOUSE] | 0,000E0 | 0,000E0 | 0,000E0 | 5,789E5 | 3,740E6 | 9,802E6 | Iars |
| P70398 | Probable ubiquitin carboxyl-terminal hydrolase FAF-X OS=Mus musculus GN=Usp9x PE=1 SV=2 - [USP9X_MOUSE] | 0,000E0 | 0,000E0 | 8,255E5 | 2,010E6 | 1,172E6 | 3,421E6 | Usp9x |
| Q8K389 | CDK5 regulatory subunit-associated protein 2 OS=Mus musculus GN=Cdk5rap2 PE=1 SV=3 - [CK5P2_MOUSE] | 0,000E0 | 0,000E0 | 0,000E0 | 8,224E5 | 3,340E6 | 2,110E6 | Cdk5rap2 |
| Q8K2D3 | Enhancer of mRNA-decapping protein 3 OS=Mus musculus GN=Edc3 PE=1 SV=1 - [EDC3_MOUSE] | 0,000E0 | 0,000E0 | 0,000E0 | 1,428E6 | 2,141E7 | 4,625E6 | Edc3 |
| P22777 | Plasminogen activator inhibitor 1 OS=Mus musculus GN=Serpine1 PE=1 SV=1 - [PAI1_MOUSE] | 0,000E0 | 0,000E0 | 0,000E0 | 4,451E6 | 4,869E6 | 1,024E7 | Serpine1 |
| F6ZDS4 | Nucleoprotein TPR OS=Mus musculus GN=Tpr PE=1 SV=1 - [TPR_MOUSE] | 0,000E0 | 0,000E0 | 0,000E0 | 7,245E5 | 3,571E6 | 5,196E6 | Tpr |
| Q8C5N3-2 | Isoform 2 of Pre-mRNA-splicing factor CWC22 homolog OS=Mus musculus GN=Cwc22 - [CWC22_MOUSE] | 0,000E0 | 0,000E0 | 0,000E0 | 6,254E6 | 7,502E6 | 1,230E7 | Cwc22 |
| Q9DBH0 | NEDD4-like E3 ubiquitin-protein ligase WWP2 OS=Mus musculus GN=Wwp2 PE=1 SV=1 - [WWP2_MOUSE] | 0,000E0 | 0,000E0 | 0,000E0 | 2,994E6 | 4,391E6 | 1,077E7 | Wwp2 |
| P10404 | MLV-related proviral Env polyprotein OS=Mus musculus PE=1 SV=3 - [ENV1_MOUSE] | 0,000E0 | 0,000E0 | 0,000E0 | 4,281E6 | 6,622E6 | 4,626E6 | ENV1 |
| P24270 | Catalase OS=Mus musculus GN=Cat PE=1 SV=4 - [CATA_MOUSE] | 0,000E0 | 0,000E0 | 0,000E0 | 2,870E6 | 1,661E6 | 6,442E5 | Cat |
| Q9R1Q9 | V-type proton ATPase subunit S1 OS=Mus musculus GN=Atp6ap1 PE=1 SV=1 - [VAS1_MOUSE] | 0,000E0 | 0,000E0 | 0,000E0 | 1,220E6 | 2,027E6 | 5,582E6 | Atp6ap1 |
| Q64343 | ATP-binding cassette sub-family G member 1 OS=Mus musculus GN=Abcg1 PE=1 SV=1 - [ABCG1_MOUSE] | 0,000E0 | 0,000E0 | 0,000E0 | 5,462E6 | 1,148E7 | 6,352E6 | Abcg1 |
| Q06806 | Tyrosine-protein kinase receptor Tie-1 OS=Mus musculus GN=Tie1 PE=1 SV=3 - [TIE1_MOUSE] | 0,000E0 | 0,000E0 | 0,000E0 | 8,554E6 | 1,148E7 | 1,662E7 | Tie1 |
| Q8BKF1 | DNA-directed RNA polymerase, mitochondrial OS=Mus musculus GN=Polrmt PE=1 SV=1 - [RPOM_MOUSE] | 0,000E0 | 0,000E0 | 0,000E0 | 2,080E6 | 5,330E6 | 1,317E8 | Polrmt |
| O54825 | Bystin OS=Mus musculus GN=Bysl PE=1 SV=3 - [BYST_MOUSE] | 0,000E0 | 0,000E0 | 0,000E0 | 7,798E5 | 1,247E6 | 2,392E6 | Bysl |
| Q3UU96 | Serine/threonine-protein kinase MRCK alpha OS=Mus musculus GN=Cdc42bpa PE=1 SV=2 - [MRCKA_MOUSE] | 0,000E0 | 0,000E0 | 0,000E0 | 2,825E6 | 1,123E7 | 2,096E7 | Cdc42bpa |
| Q6DFV3 | Rho GTPase-activating protein 21 OS=Mus musculus GN=Arhgap21 PE=1 SV=1 - [RHG21_MOUSE] | 0,000E0 | 0,000E0 | 0,000E0 | 2,212E6 | 2,860E6 | 3,103E6 | Arhgap21 |
| Q8C156 | Condensin complex subunit 2 OS=Mus musculus GN=Ncaph PE=1 SV=1 - [CND2_MOUSE] | 0,000E0 | 0,000E0 | 0,000E0 | 2,078E6 | 8,302E5 | 3,655E6 | Ncaph |
| B2RX88-2 | Isoform 2 of Centrosome and spindle pole associated protein 1 OS=Mus musculus GN=Cspp1 - [CSPP1_MOUSE] | 0,000E0 | 0,000E0 | 0,000E0 | 1,909E6 | 2,755E6 | 1,054E6 | Cspp1 |
| Q8BHY3-2 | Isoform 2 of Anoctamin-1 OS=Mus musculus GN=Ano1 - [ANO1_MOUSE] | 0,000E0 | 0,000E0 | 0,000E0 | 1,282E6 | 3,161E6 | 1,033E7 | Ano1 |
| Q62077 | 1-phosphatidylinositol 4,5-bisphosphate phosphodiesterase gamma-1 OS=Mus musculus GN=Plcg1 PE=1 SV=2 - [PLCG1_MOUSE] | 0,000E0 | 0,000E0 | 0,000E0 | 1,321E6 | 9,439E5 | 2,369E6 | Plcg1 |
| Q8BPM0-2 | Isoform 2 of Disheveled-associated activator of morphogenesis 1 OS=Mus musculus GN=Daam1 - [DAAM1_MOUSE] | 0,000E0 | 0,000E0 | 0,000E0 | 1,516E6 | 2,309E6 | 5,149E6 | Daam1 |
| Q3U0V2 | Tumor necrosis factor receptor type 1-associated DEATH domain protein OS=Mus musculus GN=Tradd PE=1 SV=1 - [TRADD_MOUSE] | 0,000E0 | 0,000E0 | 0,000E0 | 2,135E5 | 1,301E6 | 3,664E6 | Tradd |
| Q80TP3 | E3 ubiquitin-protein ligase UBR5 OS=Mus musculus GN=Ubr5 PE=1 SV=2 - [UBR5_MOUSE] | 0,000E0 | 0,000E0 | 0,000E0 | 1,013E6 | 7,372E6 | 4,983E6 | Ubr5 |
| Q61233 | Plastin-2 OS=Mus musculus GN=Lcp1 PE=1 SV=4 - [PLSL_MOUSE] | 0,000E0 | 0,000E0 | 0,000E0 | 6,535E6 | 3,327E7 | 1,948E7 | Lcp1 |
| Q8VE97 | Serine/arginine-rich splicing factor 4 OS=Mus musculus GN=Srsf4 PE=2 SV=1 - [SRSF4_MOUSE] | 0,000E0 | 0,000E0 | 0,000E0 | 1,028E8 | 7,600E7 | 3,874E6 | Srsf4 |
| O08911 | Mitogen-activated protein kinase 12 OS=Mus musculus GN=Mapk12 PE=1 SV=1 - [MK12_MOUSE] | 0,000E0 | 0,000E0 | 0,000E0 | 4,305E6 | 1,014E7 | 8,472E6 | Mapk12 |
| Q4U2R1 | E3 ubiquitin-protein ligase HERC2 OS=Mus musculus GN=Herc2 PE=1 SV=3 - [HERC2_MOUSE] | 0,000E0 | 0,000E0 | 0,000E0 | 1,073E6 | 3,650E6 | 5,635E6 | Herc2 |
| P59764 | Dedicator of cytokinesis protein 4 OS=Mus musculus GN=Dock4 PE=1 SV=1 - [DOCK4_MOUSE] | 0,000E0 | 0,000E0 | 0,000E0 | 1,700E6 | 6,895E5 | 2,725E6 | Dock4 |
| P47811-4 | Isoform 4 of Mitogen-activated protein kinase 14 OS=Mus musculus GN=Mapk14 - [MK14_MOUSE] | 0,000E0 | 0,000E0 | 0,000E0 | 4,305E6 | 1,014E7 | 8,472E6 | Mapk14 |
| Q9JMD0-4 | Isoform 4 of BUB3-interacting and GLEBS motif-containing protein ZNF207 OS=Mus musculus GN=Znf207 - [ZN207_MOUSE] | 0,000E0 | 0,000E0 | 0,000E0 | 9,455E6 | 1,236E7 | 2,380E7 | Znf207 |
| Q8CCP0 | Nuclear export mediator factor Nemf OS=Mus musculus GN=Nemf PE=1 SV=2 - [NEMF_MOUSE] | 0,000E0 | 0,000E0 | 0,000E0 | 3,048E6 | 1,315E6 | 6,649E6 | Nemf |
| Q8BT60 | Copine-3 OS=Mus musculus GN=Cpne3 PE=1 SV=2 - [CPNE3_MOUSE] | 0,000E0 | 0,000E0 | 0,000E0 | 2,218E6 | 4,070E6 | 2,504E6 | Cpne3 |
| Q6P5F9 | Exportin-1 OS=Mus musculus GN=Xpo1 PE=1 SV=1 - [XPO1_MOUSE] | 0,000E0 | 0,000E0 | 0,000E0 | 2,028E6 | 4,358E6 | 3,288E6 | Xpo1 |
| P27460 | Gag polyprotein OS=Cas-Br-E murine leukemia virus GN=gag PE=3 SV=3 - [GAG_MLVCB] | 0,000E0 | 0,000E0 | 0,000E0 | 3,382E6 | 5,988E6 | 9,946E6 | gag |
| E9Q5K9-3 | Isoform 3 of YTH domain-containing protein 1 OS=Mus musculus GN=Ythdc1 - [YTDC1_MOUSE] | 0,000E0 | 0,000E0 | 0,000E0 | 2,534E6 | 6,894E6 | 1,071E7 | Ythdc1 |
| Q8K2J0 | 1-phosphatidylinositol 4,5-bisphosphate phosphodiesterase delta-3 OS=Mus musculus GN=Plcd3 PE=1 SV=2 - [PLCD3_MOUSE] | 0,000E0 | 0,000E0 | 0,000E0 | 9,401E5 | 2,877E6 | 4,154E6 | Plcd3 |
| Q5SUA5 | Unconventional myosin-Ig OS=Mus musculus GN=Myo1g PE=1 SV=1 - [MYO1G_MOUSE] | 0,000E0 | 0,000E0 | 0,000E0 | 1,010E7 | 1,075E8 | 1,460E8 | Myo1g |
| O55103 | Periaxin OS=Mus musculus GN=Prx PE=1 SV=1 - [PRAX_MOUSE] | 0,000E0 | 0,000E0 | 0,000E0 | 3,868E6 | 7,077E6 | 7,313E6 | Prx |
| O88532 | Zinc finger RNA-binding protein OS=Mus musculus GN=Zfr PE=1 SV=2 - [ZFR_MOUSE] | 0,000E0 | 0,000E0 | 0,000E0 | 1,465E6 | 3,810E6 | 4,531E6 | Zfr |
| Q9CZU3 | Superkiller viralicidic activity 2-like 2 OS=Mus musculus GN=Skiv2l2 PE=1 SV=1 - [SK2L2_MOUSE] | 0,000E0 | 0,000E0 | 0,000E0 | 4,226E5 | 6,021E5 | 1,874E6 | Skiv2l2 |
| Q9ESV0 | ATP-dependent RNA helicase DDX24 OS=Mus musculus GN=Ddx24 PE=1 SV=2 - [DDX24_MOUSE] | 0,000E0 | 0,000E0 | 0,000E0 | 1,063E6 | 1,765E6 | 4,905E6 | Ddx24 |
| Q9D8S3 | ADP-ribosylation factor GTPase-activating protein 3 OS=Mus musculus GN=Arfgap3 PE=1 SV=2 - [ARFG3_MOUSE] | 0,000E0 | 0,000E0 | 0,000E0 | 1,234E6 | 2,397E6 | 5,351E6 | Arfgap3 |
| Q70FJ1-3 | Isoform 3 of A-kinase anchor protein 9 OS=Mus musculus GN=Akap9 - [AKAP9_MOUSE] | 0,000E0 | 0,000E0 | 0,000E0 | 1,105E6 | 2,762E6 | 3,939E6 | Akap9 |
| Q8BFY9-2 | Isoform 2 of Transportin-1 OS=Mus musculus GN=Tnpo1 - [TNPO1_MOUSE] | 0,000E0 | 0,000E0 | 0,000E0 | 2,208E6 | 3,223E6 | 3,676E6 | Tnpo1 |
| Q8CJ96 | Ras association domain-containing protein 8 OS=Mus musculus GN=Rassf8 PE=2 SV=1 - [RASF8_MOUSE] | 0,000E0 | 0,000E0 | 0,000E0 | 3,567E6 | 5,238E6 | 9,753E6 | Rassf8 |

| | | | | | | | | |
|---|---|---|---|---|---|---|---|---|
| Q920Q4-2 | Isoform 2 of Vacuolar protein sorting-associated protein 16 homolog OS=Mus musculus GN=Vps16 - [VPS16_MOUSE] | 0,000E0 | 0,000E0 | 0,000E0 | 1,449E6 | 2,497E5 | 4,894E5 | Vps16 |
| P97789-3 | Isoform 3 of 5'-3' exoribonuclease 1 OS=Mus musculus GN=Xrn1 - [XRN1_MOUSE] | 0,000E0 | 0,000E0 | 0,000E0 | 4,721E7 | 5,406E7 | 8,493E7 | Xrn1 |
| P97798-5 | Isoform 5 of Neogenin OS=Mus musculus GN=Neo1 - [NEO1_MOUSE] | 0,000E0 | 0,000E0 | 0,000E0 | 1,446E6 | 4,804E6 | 2,924E6 | Neo1 |
| Q8JZS9 | 39S ribosomal protein L48, mitochondrial OS=Mus musculus GN=Mrpl48 PE=1 SV=1 - [RM48_MOUSE] | 0,000E0 | 0,000E0 | 0,000E0 | 1,058E6 | 2,448E6 | 1,067E7 | Mrpl48 |
| Q6PDH0 | Pleckstrin homology-like domain family B member 1 OS=Mus musculus GN=Phldb1 PE=1 SV=1 - [PHLB1_MOUSE] | 0,000E0 | 0,000E0 | 0,000E0 | 6,779E5 | 2,258E6 | 3,730E6 | Phldb1 |
| Q8R418-2 | Isoform 2 of Endoribonuclease Dicer OS=Mus musculus GN=Dicer1 - [DICER_MOUSE] | 0,000E0 | 0,000E0 | 0,000E0 | 1,087E6 | 2,393E6 | 2,349E6 | Dicer1 |
| P58404-2 | Isoform 2 of Striatin-4 OS=Mus musculus GN=Strn4 - [STRN4_MOUSE] | 0,000E0 | 0,000E0 | 0,000E0 | 5,413E5 | 2,563E6 | 4,056E6 | Strn4 |
| O54774 | AP-3 complex subunit delta-1 OS=Mus musculus GN=Ap3d1 PE=1 SV=1 - [AP3D1_MOUSE] | 0,000E0 | 0,000E0 | 0,000E0 | 3,698E6 | 7,873E5 | 3,980E6 | Ap3d1 |
| Q9WVL3 | Solute carrier family 12 member 7 OS=Mus musculus GN=Slc12a7 PE=1 SV=1 - [S12A7_MOUSE] | 0,000E0 | 0,000E0 | 0,000E0 | 3,468E6 | 4,885E6 | 3,674E6 | Slc12a7 |
| Q6PAR5-5 | Isoform 5 of GTPase-activating protein and VPS9 domain-containing protein 1 OS=Mus musculus GN=Gapvd1 - [GAPD1_MOUSE] | 0,000E0 | 0,000E0 | 0,000E0 | 1,895E6 | 3,290E6 | 4,345E6 | Gapvd1 |
| Q8VE73-3 | Isoform 2 of Cullin-7 OS=Mus musculus GN=Cul7 - [CUL7_MOUSE] | 0,000E0 | 0,000E0 | 0,000E0 | 1,794E6 | 2,435E6 | 5,081E6 | Cul7 |
| Q6DFX2 | Anthrax toxin receptor 2 OS=Mus musculus GN=Antxr2 PE=1 SV=1 - [ANTR2_MOUSE] | 0,000E0 | 0,000E0 | 0,000E0 | 5,963E6 | 7,414E6 | 1,849E7 | Antxr2 |
| Q6P5E4 | UDP-glucose:glycoprotein glucosyltransferase 1 OS=Mus musculus GN=Uggt1 PE=1 SV=4 - [UGGG1_MOUSE] | 0,000E0 | 0,000E0 | 0,000E0 | 2,985E6 | 4,484E6 | 3,749E6 | Uggt1 |
| Q9DBB4 | N-alpha-acetyltransferase 16, NatA auxiliary subunit OS=Mus musculus GN=Naa16 PE=2 SV=1 - [NAA16_MOUSE] | 0,000E0 | 0,000E0 | 0,000E0 | 4,245E6 | 1,204E7 | 1,929E7 | Naa16 |
| Q6PDQ2 | Chromodomain-helicase-DNA-binding protein 4 OS=Mus musculus GN=Chd4 PE=1 SV=1 - [CHD4_MOUSE] | 0,000E0 | 0,000E0 | 0,000E0 | 5,787E6 | 4,604E6 | 7,235E6 | Chd4 |
| Q8C0E3 | Tripartite motif-containing protein 47 OS=Mus musculus GN=Trim47 PE=1 SV=2 - [TRI47_MOUSE] | 0,000E0 | 0,000E0 | 0,000E0 | 2,201E6 | 1,115E6 | 1,048E6 | Trim47 |
| Q99MD9-2 | Isoform 2 of Nuclear autoantigenic sperm protein OS=Mus musculus GN=Nasp - [NASP_MOUSE] | 0,000E0 | 0,000E0 | 0,000E0 | 1,443E6 | 2,879E6 | 3,129E6 | Nasp |
| P06795 | Multidrug resistance protein 1B OS=Mus musculus GN=Abcb1b PE=1 SV=1 - [MDR1B_MOUSE] | 0,000E0 | 0,000E0 | 0,000E0 | 8,525E5 | 2,665E6 | 4,124E6 | Abcb1b |
| Q3UYV9 | Nuclear cap-binding protein subunit 1 OS=Mus musculus GN=Ncbp1 PE=1 SV=2 - [NCBP1_MOUSE] | 0,000E0 | 0,000E0 | 0,000E0 | 3,681E6 | 5,270E6 | 1,785E6 | Ncbp1 |
| A2AR02 | Peptidyl-prolyl cis-trans isomerase G OS=Mus musculus GN=Ppig PE=1 SV=1 - [PPIG_MOUSE] | 0,000E0 | 0,000E0 | 0,000E0 | 3,317E6 | 5,165E6 | 8,024E6 | Ppig |
| P17809 | Solute carrier family 2, facilitated glucose transporter member 1 OS=Mus musculus GN=Slc2a1 PE=1 SV=4 - [GTR1_MOUSE] | 0,000E0 | 0,000E0 | 0,000E0 | 1,538E6 | 4,671E6 | 7,468E6 | Slc2a1 |
| Q7SIG6 | Arf-GAP with SH3 domain, ANK repeat and PH domain-containing protein 2 OS=Mus musculus GN=Asap2 PE=1 SV=3 - [ASAP2_MOUSE] | 0,000E0 | 0,000E0 | 0,000E0 | 7,767E5 | 3,134E6 | 1,793E6 | Asap2 |
| Q91Z69 | SLIT-ROBO Rho GTPase-activating protein 1 OS=Mus musculus GN=Srgap1 PE=1 SV=2 - [SRGP1_MOUSE] | 0,000E0 | 0,000E0 | 0,000E0 | 1,599E7 | 2,463E7 | 5,061E7 | Srgap1 |
| Q640Q5 | PAB-dependent poly(A)-specific ribonuclease subunit PAN3 OS=Mus musculus GN=Pan3 PE=1 SV=3 - [PAN3_MOUSE] | 0,000E0 | 0,000E0 | 0,000E0 | 1,110E6 | 2,124E6 | 1,323E6 | Pan3 |
| Q8BUR4 | Dedicator of cytokinesis protein 1 OS=Mus musculus GN=Dock1 PE=1 SV=3 - [DOCK1_MOUSE] | 0,000E0 | 0,000E0 | 0,000E0 | 1,558E6 | 4,658E6 | 4,354E6 | Dock1 |
| P18406 | Protein CYR61 OS=Mus musculus GN=Cyr61 PE=1 SV=1 - [CYR61_MOUSE] | 0,000E0 | 0,000E0 | 0,000E0 | 8,365E5 | 3,064E6 | 8,443E6 | Cyr61 |
| Q9D5V5 | Cullin-5 OS=Mus musculus GN=Cul5 PE=1 SV=3 - [CUL5_MOUSE] | 0,000E0 | 0,000E0 | 0,000E0 | 6,775E6 | 2,137E6 | 5,404E6 | Cul5 |
| Q08297 | DNA repair protein RAD51 homolog 1 OS=Mus musculus GN=Rad51 PE=1 SV=1 - [RAD51_MOUSE] | 0,000E0 | 0,000E0 | 0,000E0 | 5,634E5 | 3,666E6 | 2,870E6 | Rad51 |
| Q9WTU6-2 | Isoform Alpha-1 of Mitogen-activated protein kinase 9 OS=Mus musculus GN=Mapk9 - [MK09_MOUSE] | 0,000E0 | 0,000E0 | 0,000E0 | 4,305E6 | 6,578E6 | 8,472E6 | Mapk9 |
| Q9D742 | AP-5 complex subunit sigma-1 OS=Mus musculus GN=Ap5s1 PE=1 SV=1 - [AP5S1_MOUSE] | 0,000E0 | 0,000E0 | 0,000E0 | 1,281E7 | 4,401E7 | 9,475E7 | Ap5s1 |
| P08122 | Collagen alpha-2(IV) chain OS=Mus musculus GN=Col4a2 PE=1 SV=4 - [CO4A2_MOUSE] | 0,000E0 | 0,000E0 | 0,000E0 | 8,132E5 | 8,923E6 | 2,293E7 | Col4a2 |
| Q14AX6-3 | Isoform 3 of Cyclin-dependent kinase 12 OS=Mus musculus GN=Cdk12 - [CDK12_MOUSE] | 0,000E0 | 0,000E0 | 0,000E0 | 7,773E6 | 1,847E7 | 3,097E7 | Cdk12 |
| P48722-2 | Isoform 2 of Heat shock 70 kDa protein 4L OS=Mus musculus GN=Hspa4l - [HS74L_MOUSE] | 0,000E0 | 0,000E0 | 0,000E0 | 3,849E6 | 1,344E7 | 2,462E7 | Hspa4l |
| Q8K2Z4 | Condensin complex subunit 1 OS=Mus musculus GN=Ncapd2 PE=1 SV=2 - [CND1_MOUSE] | 0,000E0 | 0,000E0 | 0,000E0 | 3,405E6 | 4,844E6 | 3,110E6 | Ncapd2 |
| Q9D0G0 | 28S ribosomal protein S30, mitochondrial OS=Mus musculus GN=Mrps30 PE=1 SV=1 - [RT30_MOUSE] | 0,000E0 | 0,000E0 | 0,000E0 | 1,736E6 | 6,170E6 | 5,311E6 | Mrps30 |
| Q9DBR0 | A-kinase anchor protein 8 OS=Mus musculus GN=Akap8 PE=1 SV=1 - [AKAP8_MOUSE] | 0,000E0 | 0,000E0 | 0,000E0 | 4,310E6 | 2,684E6 | 2,331E6 | Akap8 |
| Q8K4R9-3 | Isoform 3 of Disks large-associated protein 5 OS=Mus musculus GN=Dlgap5 - [DLGP5_MOUSE] | 0,000E0 | 0,000E0 | 0,000E0 | 7,256E5 | 1,115E6 | 6,524E5 | Dlgap5 |
| P11881-8 | Isoform 8 of Inositol 1,4,5-trisphosphate receptor type 1 OS=Mus musculus GN=Itpr1 - [ITPR1_MOUSE] | 0,000E0 | 0,000E0 | 0,000E0 | 3,231E6 | 6,095E6 | 9,739E6 | Itpr1 |
| Q9DC23 | DnaJ homolog subfamily C member 10 OS=Mus musculus GN=Dnajc10 PE=1 SV=2 - [DJC10_MOUSE] | 0,000E0 | 0,000E0 | 0,000E0 | 8,297E5 | 6,589E6 | 1,239E7 | Dnajc10 |
| E9PZJ8 | Activating signal cointegrator 1 complex subunit 3 OS=Mus musculus GN=Ascc3 PE=1 SV=1 - [ASCC3_MOUSE] | 0,000E0 | 0,000E0 | 0,000E0 | 1,845E6 | 3,609E6 | 8,034E6 | Ascc3 |
| Q99KQ4 | Nicotinamide phosphoribosyltransferase OS=Mus musculus GN=Nampt PE=1 SV=1 - [NAMPT_MOUSE] | 0,000E0 | 0,000E0 | 0,000E0 | 1,363E6 | 7,218E5 | 3,470E6 | Nampt |
| P33174 | Chromosome-associated kinesin KIF4 OS=Mus musculus GN=Kif4 PE=1 SV=3 - [KIF4_MOUSE] | 0,000E0 | 0,000E0 | 0,000E0 | 1,563E6 | 2,604E6 | 4,897E6 | Kif4 |
| Q8BQN6 | Zinc finger protein OZF OS=Mus musculus GN=Znf146 PE=2 SV=2 - [OZF_MOUSE] | 0,000E0 | 0,000E0 | 0,000E0 | 1,770E6 | 5,388E6 | 7,491E6 | Znf146 |
| P52431 | DNA polymerase delta catalytic subunit OS=Mus musculus GN=Pold1 PE=1 SV=2 - [DPOD1_MOUSE] | 0,000E0 | 0,000E0 | 0,000E0 | 5,264E5 | 2,011E6 | 3,008E6 | Pold1 |
| Q9QXD8 | LIM domain-containing protein 1 OS=Mus musculus GN=Limd1 PE=1 SV=2 - [LIMD1_MOUSE] | 0,000E0 | 0,000E0 | 0,000E0 | 1,322E6 | 5,519E6 | 4,380E6 | Limd1 |
| P15306 | Thrombomodulin OS=Mus musculus GN=Thbd PE=1 SV=1 - [TRBM_MOUSE] | 0,000E0 | 0,000E0 | 0,000E0 | 3,583E7 | 1,059E7 | 7,472E6 | Thbd |
| Q8CDJ8 | Stonin-1 OS=Mus musculus GN=Ston1 PE=1 SV=2 - [STON1_MOUSE] | 0,000E0 | 0,000E0 | 0,000E0 | 1,355E6 | 3,264E6 | 8,801E6 | Ston1 |
| P42225 | Signal transducer and activator of transcription 1 OS=Mus musculus GN=Stat1 PE=1 SV=1 - [STAT1_MOUSE] | 0,000E0 | 0,000E0 | 0,000E0 | 1,240E6 | 8,767E5 | 1,237E8 | Stat1 |
| Q9Z329-3 | Isoform 3 of Inositol 1,4,5-trisphosphate receptor type 2 OS=Mus musculus GN=Itpr2 - [ITPR2_MOUSE] | 0,000E0 | 0,000E0 | 0,000E0 | 3,667E6 | 4,245E6 | 6,724E6 | Itpr2 |
| Q99PV0 | Pre-mRNA-processing-splicing factor 8 OS=Mus musculus GN=Prpf8 PE=1 SV=2 - [PRP8_MOUSE] | 0,000E0 | 0,000E0 | 0,000E0 | 8,824E5 | 9,027E5 | 2,375E6 | Prpf8 |
| Q8BFZ2-2 | Isoform 2 of Phospholipid phosphatase-related protein type 1 OS=Mus musculus GN=Plppr1 - [PLPR1_MOUSE] | 0,000E0 | 0,000E0 | 0,000E0 | 5,935E6 | 9,569E6 | 2,766E6 | Plppr1 |
| Q99JW4 | LIM and senescent cell antigen-like-containing domain protein 1 OS=Mus musculus GN=Lims1 PE=1 SV=3 - [LIMS1_MOUSE] | 0,000E0 | 0,000E0 | 0,000E0 | 5,696E6 | 9,240E6 | 1,772E7 | Lims1 |
| Q7TPD1-3 | Isoform 3 of F-box only protein 11 OS=Mus musculus GN=Fbxo11 - [FBX11_MOUSE] | 0,000E0 | 0,000E0 | 0,000E0 | 6,609E5 | 2,284E6 | 5,642E6 | Fbxo11 |
| Q9DCC4 | Pyrroline-5-carboxylate reductase 3 OS=Mus musculus GN=Pycrl PE=1 SV=2 - [P5CR3_MOUSE] | 0,000E0 | 0,000E0 | 0,000E0 | 1,854E6 | 1,452E6 | 4,273E6 | Pycrl |
| Q9R1S8 | Calpain-7 OS=Mus musculus GN=Capn7 PE=2 SV=1 - [CAN7_MOUSE] | 0,000E0 | 0,000E0 | 0,000E0 | 1,762E6 | 2,080E6 | 1,511E6 | Capn7 |
| Q9WTR2 | Mitogen-activated protein kinase kinase kinase 6 OS=Mus musculus GN=Map3k6 PE=1 SV=4 - [M3K6_MOUSE] | 0,000E0 | 0,000E0 | 0,000E0 | 7,774E8 | 1,237E9 | 1,635E8 | Map3k6 |
| Q9QYJ3 | DnaJ homolog subfamily B member 1 OS=Mus musculus GN=Dnajb1 PE=1 SV=3 - [DNJB1_MOUSE] | 0,000E0 | 0,000E0 | 0,000E0 | 2,528E6 | 1,835E6 | 4,638E6 | Dnajb1 |
| Q71LX4 | Talin-2 OS=Mus musculus GN=Tln2 PE=1 SV=3 - [TLN2_MOUSE] | 0,000E0 | 0,000E0 | 0,000E0 | 3,980E6 | 1,115E7 | 5,270E6 | Tln2 |
| F8VPU2 | FERM, RhoGEF and pleckstrin domain-containing protein 1 OS=Mus musculus GN=Farp1 PE=1 SV=1 - [FARP1_MOUSE] | 0,000E0 | 0,000E0 | 0,000E0 | 6,273E5 | 1,829E6 | 3,045E6 | Farp1 |
| P83741-4 | Isoform 4 of Serine/threonine-protein kinase WNK1 OS=Mus musculus GN=Wnk1 - [WNK1_MOUSE] | 0,000E0 | 0,000E0 | 0,000E0 | 1,033E6 | 1,671E6 | 2,138E6 | Wnk1 |
| P70206 | Plexin-A1 OS=Mus musculus GN=Plxna1 PE=1 SV=1 - [PLXA1_MOUSE] | 0,000E0 | 0,000E0 | 0,000E0 | 2,637E6 | 3,080E6 | 8,465E6 | Plxna1 |
| P53995 | Anaphase-promoting complex subunit 1 OS=Mus musculus GN=Anapc1 PE=1 SV=2 - [APC1_MOUSE] | 0,000E0 | 0,000E0 | 0,000E0 | 2,948E6 | 2,756E6 | 3,839E6 | Anapc1 |
| Q9QY73 | Transmembrane protein 59 OS=Mus musculus GN=Tmem59 PE=1 SV=2 - [TMM59_MOUSE] | 0,000E0 | 0,000E0 | 0,000E0 | 8,690E6 | 5,998E6 | 1,482E6 | Tmem59 |
| P00532 | Serine/threonine-protein kinase-transforming protein raf OS=Murine sarcoma virus 3611 GN=V-RAF PE=3 SV=1 - [RAF_MSV36] | 0,000E0 | 0,000E0 | 0,000E0 | 3,362E6 | 6,219E6 | 1,139E7 | V-RAF |
| Q9JIS8 | Solute carrier family 12 member 4 OS=Mus musculus GN=Slc12a4 PE=1 SV=2 - [S12A4_MOUSE] | 0,000E0 | 0,000E0 | 0,000E0 | 2,203E6 | 7,992E6 | 1,280E7 | Slc12a4 |
| Q6P549 | Phosphatidylinositol 3,4,5-trisphosphate 5-phosphatase 2 OS=Mus musculus GN=Inppl1 PE=1 SV=1 - [SHIP2_MOUSE] | 0,000E0 | 0,000E0 | 0,000E0 | 4,295E6 | 6,776E6 | 1,432E7 | Inppl1 |
| Q5SSK3 | Transcription elongation factor, mitochondrial OS=Mus musculus GN=Tefm PE=1 SV=1 - [TEFM_MOUSE] | 0,000E0 | 0,000E0 | 0,000E0 | 5,918E5 | 1,960E6 | 3,581E6 | Tefm |
| Q9Z0R4 | Intersectin-1 OS=Mus musculus GN=Itsn1 PE=1 SV=2 - [ITSN1_MOUSE] | 0,000E0 | 0,000E0 | 0,000E0 | 4,169E6 | 2,358E6 | 6,731E6 | Itsn1 |
| D3YXK2 | Scaffold attachment factor B1 OS=Mus musculus GN=Safb PE=1 SV=2 - [SAFB1_MOUSE] | 0,000E0 | 0,000E0 | 0,000E0 | 2,573E6 | 5,010E6 | 1,005E7 | Safb |
| P55937-1 | Isoform 1 of Golgin subfamily A member 3 OS=Mus musculus GN=Golga3 - [GOGA3_MOUSE] | 0,000E0 | 0,000E0 | 0,000E0 | 1,080E6 | 1,569E6 | 2,899E6 | Golga3 |
| Q9Z280-2 | Isoform PLD1B of Phospholipase D1 OS=Mus musculus GN=Pld1 - [PLD1_MOUSE] | 0,000E0 | 0,000E0 | 0,000E0 | 2,111E6 | 2,099E6 | 3,634E6 | Pld1 |
| B1AVY7 | Kinesin-like protein KIF16B OS=Mus musculus GN=Kif16b PE=1 SV=1 - [KI16B_MOUSE] | 0,000E0 | 0,000E0 | 0,000E0 | 1,563E6 | 2,138E6 | 4,571E6 | Kif16b |
| Q4JIM5 | Abelson tyrosine-protein kinase 2 OS=Mus musculus GN=Abl2 PE=1 SV=1 - [ABL2_MOUSE] | 0,000E0 | 0,000E0 | 0,000E0 | 1,355E6 | 1,597E6 | 3,912E6 | Abl2 |
| Q9R117 | Non-receptor tyrosine-protein kinase TYK2 OS=Mus musculus GN=Tyk2 PE=1 SV=2 - [TYK2_MOUSE] | 0,000E0 | 0,000E0 | 0,000E0 | 2,786E6 | 6,157E6 | 8,241E6 | Tyk2 |
| Q8BWY9 | Protein CIP2A OS=Mus musculus GN=Kiaa1524 PE=1 SV=3 - [CIP2A_MOUSE] | 0,000E0 | 0,000E0 | 0,000E0 | 1,554E6 | 2,989E6 | 4,467E6 | Kiaa1524 |
| O88738-3 | Isoform 3 of Baculoviral IAP repeat-containing protein 6 OS=Mus musculus GN=Birc6 - [BIRC6_MOUSE] | 0,000E0 | 0,000E0 | 0,000E0 | 5,657E6 | 4,645E6 | 5,420E6 | Birc6 |
| Q80TV8-2 | Isoform 2 of CLIP-associating protein 1 OS=Mus musculus GN=Clasp1 - [CLAP1_MOUSE] | 0,000E0 | 0,000E0 | 0,000E0 | 1,014E6 | 1,426E6 | 5,115E5 | Clasp1 |
| P59913 | Protein-L-isoaspartate O-methyltransferase domain-containing protein 1 OS=Mus musculus GN=Pcmtd1 PE=1 SV=1 - [PCMD1_MOUSE] | 0,000E0 | 0,000E0 | 0,000E0 | 3,097E6 | 2,367E6 | 1,530E6 | Pcmtd1 |
| P01872 | Ig mu chain C region OS=Mus musculus GN=Ighm PE=1 SV=2 - [IGHM_MOUSE] | 0,000E0 | 0,000E0 | 0,000E0 | 4,839E7 | 4,957E8 | 4,273E8 | Ighm |
| Q99M51 | Cytoplasmic protein NCK1 OS=Mus musculus GN=Nck1 PE=1 SV=1 - [NCK1_MOUSE] | 0,000E0 | 0,000E0 | 0,000E0 | 1,839E6 | 1,213E6 | 3,944E6 | Nck1 |
| P35917 | Vascular endothelial growth factor receptor 3 OS=Mus musculus GN=Flt4 PE=1 SV=1 - [VGFR3_MOUSE] | 0,000E0 | 0,000E0 | 0,000E0 | 3,523E6 | 1,319E7 | 1,225E7 | Flt4 |
| Q8CGY8-2 | Isoform 2 of UDP-N-acetylglucosamine--peptide N-acetylglucosaminyltransferase 110 kDa subunit OS=Mus musculus GN=Ogt - [OGT1_MOUSE] | 0,000E0 | 0,000E0 | 0,000E0 | 1,460E6 | 3,095E6 | 5,330E6 | Ogt |
| Q8VDG3 | Poly(A)-specific ribonuclease PARN OS=Mus musculus GN=Parn PE=1 SV=1 - [PARN_MOUSE] | 0,000E0 | 0,000E0 | 0,000E0 | 1,275E6 | 3,004E6 | 3,737E6 | Parn |
| Q7TMY8-4 | Isoform 4 of E3 ubiquitin-protein ligase HUWE1 OS=Mus musculus GN=Huwe1 - [HUWE1_MOUSE] | 0,000E0 | 0,000E0 | 0,000E0 | 2,401E6 | 1,430E6 | 3,032E6 | Huwe1 |
| Q3U214 | Microtubule-associated serine/threonine-protein kinase 3 OS=Mus musculus GN=Mast3 PE=1 SV=3 - [MAST3_MOUSE] | 0,000E0 | 0,000E0 | 0,000E0 | 3,096E6 | 2,839E6 | 3,860E6 | Mast3 |
| Q640M1 | U3 small nucleolar RNA-associated protein 14 homolog A OS=Mus musculus GN=Utp14a PE=1 SV=1 - [UT14A_MOUSE] | 0,000E0 | 0,000E0 | 0,000E0 | 1,773E6 | 8,582E5 | 1,322E6 | Utp14a |
| Q8BZQ7 | Anaphase-promoting complex subunit 2 OS=Mus musculus GN=Anapc2 PE=1 SV=2 - [ANC2_MOUSE] | 0,000E0 | 0,000E0 | 0,000E0 | 6,631E6 | 8,064E6 | 1,735E7 | Anapc2 |
| A2AN08-3 | Isoform 3 of E3 ubiquitin-protein ligase UBR4 OS=Mus musculus GN=Ubr4 - [UBR4_MOUSE] | 0,000E0 | 0,000E0 | 0,000E0 | 2,192E6 | 3,358E6 | 6,423E6 | Ubr4 |
| Q99MR1 | PERQ amino acid-rich with GYF domain-containing protein 1 OS=Mus musculus GN=Gigyf1 PE=1 SV=2 - [PERQ1_MOUSE] | 0,000E0 | 0,000E0 | 0,000E0 | 2,615E6 | 5,911E6 | 9,832E6 | Gigyf1 |
| Q91ZX7 | Prolow-density lipoprotein receptor-related protein 1 OS=Mus musculus GN=Lrp1 PE=1 SV=1 - [LRP1_MOUSE] | 0,000E0 | 0,000E0 | 0,000E0 | 2,824E6 | 7,341E6 | 6,997E6 | Lrp1 |
| Q80WQ6 | Inactive rhomboid protein 2 OS=Mus musculus GN=Rhbdf2 PE=1 SV=1 - [RHDF2_MOUSE] | 0,000E0 | 0,000E0 | 0,000E0 | 2,015E6 | 5,360E6 | 8,553E6 | Rhbdf2 |
| A2ASS6 | Titin OS=Mus musculus GN=Ttn PE=1 SV=1 - [TITIN_MOUSE] | 0,000E0 | 0,000E0 | 0,000E0 | 5,187E6 | 8,800E6 | 1,166E7 | Ttn |

**Supplementary Table 2. Mass-spec analysis of AmotL2 immunoprecipitation in BAE cells**

| Accession | Description | Area_Ctrl_1 | Area_Ctrl_2 | Area_Ctrl_3 | Area_AmotL2_1 | Area_AmotL2_2 | Area_AmotL2_3 | Gene name |
|---|---|---|---|---|---|---|---|---|
| F1MQ37 | Myosin heavy chain 9 OS=Bos taurus OX=9913 GN=MYH9 PE=1 SV=2 - [F1MQ37_BOVIN] | 6,020E7 | 4,102E6 | 6,816E6 | 5,610E9 | 1,448E10 | 3,618E10 | MYH9 |
| E1BF59 | Plectin OS=Bos taurus OX=9913 GN=PLEC PE=1 SV=2 - [E1BF59_BOVIN] | 0,000E0 | 0,000E0 | 0,000E0 | 2,215E8 | 1,471E9 | 1,320E9 | PLEC |
| Q27991 | Myosin-10 OS=Bos taurus OX=9913 GN=MYH10 PE=2 SV=2 - [MYH10_BOVIN] | 5,663E7 | 1,987E6 | 0,000E0 | 3,047E9 | 8,568E9 | 9,278E9 | MYH10 |
| B8Y9S9 | Embryo-specific fibronectin 1 transcript variant OS=Bos taurus OX=9913 GN=FN1 PE=2 SV=1 - [B8Y9S9_BOVIN] | 0,000E0 | 0,000E0 | 0,000E0 | 3,601E8 | 2,960E9 | 2,496E9 | FN1 |
| O02717 | Non-muscle myosin heavy chain (Fragment) OS=Bos taurus OX=9913 PE=2 SV=2 - [O02717_BOVIN] | 5,457E7 | 0,000E0 | 7,722E6 | 4,326E9 | 1,312E10 | 2,392E10 | O02717 |
| E1BFB0 | Spectrin alpha, non-erythrocytic 1 OS=Bos taurus OX=9913 GN=SPTAN1 PE=1 SV=2 - [E1BFB0_BOVIN] | 0,000E0 | 0,000E0 | 0,000E0 | 2,591E7 | 3,456E8 | 3,012E8 | SPTAN1 |
| E1BKX7 | Filamin B OS=Bos taurus OX=9913 GN=FLNB PE=1 SV=2 - [E1BKX7_BOVIN] | 0,000E0 | 0,000E0 | 0,000E0 | 1,752E7 | 3,028E8 | 3,291E8 | FLNB |
| F1MYC9 | Spectrin beta chain OS=Bos taurus OX=9913 GN=SPTBN1 PE=1 SV=1 - [F1MYC9_BOVIN] | 0,000E0 | 0,000E0 | 0,000E0 | 2,334E7 | 2,665E8 | 2,825E8 | SPTBN1 |
| F1MER7 | Heparan sulfate proteoglycan 2 OS=Bos taurus OX=9913 GN=HSPG2 PE=1 SV=1 - [F1MER7_BOVIN] | 0,000E0 | 0,000E0 | 0,000E0 | 3,977E7 | 4,752E8 | 4,630E8 | HSPG2 |
| P48616 | Vimentin OS=Bos taurus OX=9913 GN=VIM PE=1 SV=3 - [VIME_BOVIN] | 1,773E7 | 3,057E7 | 0,000E0 | 4,604E9 | 1,087E10 | 1,784E10 | VIM |
| Q27966-3 | Isoform 3 of Unconventional myosin-Ic OS=Bos taurus OX=9913 GN=MYO1C - [MYO1C_BOVIN] | 1,185E6 | 0,000E0 | 0,000E0 | 5,697E8 | 4,504E9 | 4,765E9 | MYO1C |
| F1MHT3 | Myosin VA OS=Bos taurus OX=9913 GN=MYO5A PE=3 SV=2 - [F1MHT3_BOVIN] | 0,000E0 | 0,000E0 | 0,000E0 | 7,743E7 | 5,367E8 | 3,435E8 | MYO5A |
| A5D7D1 | Alpha-actinin-4 OS=Bos taurus OX=9913 GN=ACTN4 PE=2 SV=1 - [ACTN4_BOVIN] | 0,000E0 | 0,000E0 | 0,000E0 | 1,224E8 | 1,008E9 | 9,257E8 | ACTN4 |
| F1MQ21 | Nestin OS=Bos taurus OX=9913 GN=NES PE=3 SV=2 - [F1MQ21_BOVIN] | 0,000E0 | 0,000E0 | 0,000E0 | 2,002E8 | 8,770E8 | 9,379E8 | NES |
| P60712 | Actin, cytoplasmic 1 OS=Bos taurus OX=9913 GN=ACTB PE=1 SV=1 - [ACTB_BOVIN] | 2,164E8 | 2,069E7 | 0,000E0 | 2,369E10 | 1,389E10 | 9,397E9 | ACTB |
| F1MJ56 | Unconventional myosin-Id OS=Bos taurus OX=9913 GN=MYO1D PE=3 SV=1 - [F1MJ56_BOVIN] | 0,000E0 | 0,000E0 | 0,000E0 | 2,315E8 | 8,567E8 | 5,625E8 | MYO1D |
| F1MQ65 | Myosin IE OS=Bos taurus OX=9913 GN=MYO1E PE=3 SV=2 - [F1MQ65_BOVIN] | 0,000E0 | 0,000E0 | 0,000E0 | 8,994E7 | 7,000E8 | 6,884E8 | MYO1E |
| G5E631 | Uncharacterized protein OS=Bos taurus OX=9913 GN=DDX3X PE=1 SV=1 - [G5E631_BOVIN] | 1,741E8 | 5,158E6 | 0,000E0 | 8,700E8 | 3,134E8 | 4,333E8 | DDX3X |
| F1MRD0 | Actin, cytoplasmic 1 OS=Bos taurus OX=9913 GN=ACTB PE=3 SV=2 - [F1MRD0_BOVIN] | 2,164E8 | 2,069E7 | 0,000E0 | 1,916E10 | 9,283E9 | 9,397E9 | ACTB |
| Q3B7N2 | Alpha-actinin-1 OS=Bos taurus OX=9913 GN=ACTN1 PE=2 SV=1 - [ACTN1_BOVIN] | 0,000E0 | 0,000E0 | 0,000E0 | 8,213E7 | 5,927E8 | 6,215E8 | ACTN1 |
| E1BDF0 | Nexilin F-actin binding protein OS=Bos taurus OX=9913 GN=NEXN PE=4 SV=2 - [E1BDF0_BOVIN] | 0,000E0 | 0,000E0 | 0,000E0 | 1,259E8 | 7,540E8 | 1,343E9 | NEXN |
| Q1RMT6 | Drebrin 1 OS=Bos taurus OX=9913 GN=DBN1 PE=1 SV=1 - [Q1RMT6_BOVIN] | 0,000E0 | 0,000E0 | 0,000E0 | 8,127E8 | 1,487E9 | 1,750E9 | DBN1 |
| E1BPK6 | Myosin VI OS=Bos taurus OX=9913 GN=MYO6 PE=3 SV=2 - [E1BPK6_BOVIN] | 0,000E0 | 0,000E0 | 0,000E0 | 5,921E7 | 3,677E8 | 5,488E8 | MYO6 |
| F1N6K7 | Protein transport protein sec16 OS=Bos taurus OX=9913 GN=SEC16A PE=3 SV=2 - [F1N6K7_BOVIN] | 1,024E7 | 0,000E0 | 8,998E6 | 5,670E7 | 4,123E7 | 4,923E7 | SEC16A |
| E1BDU8 | LIM domain and actin binding 1 OS=Bos taurus OX=9913 GN=LIMA1 PE=1 SV=2 - [E1BDU8_BOVIN] | 0,000E0 | 0,000E0 | 0,000E0 | 8,139E7 | 1,247E9 | 1,051E9 | LIMA1 |
| F1MY06 | Trinucleotide repeat containing 6B OS=Bos taurus OX=9913 GN=TNRC6B PE=4 SV=2 - [F1MY06_BOVIN] | 9,599E6 | 0,000E0 | 1,379E7 | 4,931E7 | 3,929E7 | 3,542E7 | TNRC6B |
| P19120 | Heat shock cognate 71 kDa protein OS=Bos taurus OX=9913 GN=HSPA8 PE=1 SV=2 - [HSP7C_BOVIN] | 2,063E7 | 8,924E6 | 1,636E7 | 4,185E8 | 1,116E9 | 1,126E9 | HSPA8 |
| E1BC59 | Protein phosphatase 1 regulatory subunit 12A OS=Bos taurus OX=9913 GN=PPP1R12A PE=4 SV=2 - [E1BC59_BOVIN] | 0,000E0 | 0,000E0 | 0,000E0 | 8,072E7 | 7,197E8 | 6,971E8 | PPP1R12A |
| A0A140T897 | Serum albumin OS=Bos taurus OX=9913 GN=ALB PE=4 SV=1 - [A0A140T897_BOVIN] | 1,543E8 | 0,000E0 | 3,453E7 | 1,193E8 | 5,507E7 | 2,688E7 | ALB |
| G3MWI5 | Coronin OS=Bos taurus OX=9913 GN=CORO1C PE=3 SV=1 - [G3MWI5_BOVIN] | 0,000E0 | 0,000E0 | 0,000E0 | 1,239E8 | 9,689E8 | 5,237E8 | CORO1C |
| E1BKT0 | Leucine zipper protein 1 OS=Bos taurus OX=9913 GN=LUZP1 PE=4 SV=2 - [E1BKT0_BOVIN] | 0,000E0 | 0,000E0 | 0,000E0 | 1,877E7 | 9,483E7 | 9,462E7 | LUZP1 |
| Q3ZCH0 | Stress-70 protein, mitochondrial OS=Bos taurus OX=9913 GN=HSPA9 PE=2 SV=1 - [GRP75_BOVIN] | 2,855E7 | 1,100E7 | 2,041E7 | 3,600E8 | 3,932E8 | 3,923E8 | HSPA9 |
| F1N2D3 | Tight junction protein 1 OS=Bos taurus OX=9913 GN=TJP1 PE=3 SV=2 - [F1N2D3_BOVIN] | 0,000E0 | 0,000E0 | 0,000E0 | 1,086E7 | 7,229E7 | 1,165E8 | TJP1 |
| P49951 | Clathrin heavy chain 1 OS=Bos taurus OX=9913 GN=CLTC PE=1 SV=1 - [CLH1_BOVIN] | 0,000E0 | 0,000E0 | 0,000E0 | 1,570E7 | 1,786E8 | 2,593E8 | CLTC |
| A6QLD6 | MYO1B protein OS=Bos taurus OX=9913 GN=MYO1B PE=2 SV=1 - [A6QLD6_BOVIN] | 0,000E0 | 0,000E0 | 0,000E0 | 8,800E6 | 1,984E8 | 1,939E8 | MYO1B |
| Q5KR49 | Tropomyosin alpha-1 chain OS=Bos taurus OX=9913 GN=TPM1 PE=2 SV=1 - [TPM1_BOVIN] | 0,000E0 | 0,000E0 | 0,000E0 | 1,441E9 | 8,777E9 | 7,273E9 | TPM1 |
| A4IFM8 | Actin, alpha 1, skeletal muscle OS=Bos taurus OX=9913 GN=ACTA1 PE=2 SV=1 - [A4IFM8_BOVIN] | 2,426E8 | 2,069E7 | 0,000E0 | 1,945E10 | 1,203E10 | 7,861E9 | ACTA1 |
| F1N365 | FLII, actin remodeling protein OS=Bos taurus OX=9913 GN=FLII PE=4 SV=1 - [F1N365_BOVIN] | 0,000E0 | 0,000E0 | 0,000E0 | 2,696E7 | 2,339E8 | 1,845E8 | FLII |
| Q5KR47-2 | Isoform 2 of Tropomyosin alpha-3 chain OS=Bos taurus OX=9913 GN=TPM3 - [TPM3_BOVIN] | 0,000E0 | 0,000E0 | 0,000E0 | 1,349E9 | 9,210E9 | 8,943E9 | TPM3 |
| P61286 | Polyadenylate-binding protein 1 OS=Bos taurus OX=9913 GN=PABPC1 PE=2 SV=1 - [PABP1_BOVIN] | 1,166E7 | 0,000E0 | 0,000E0 | 2,575E8 | 1,549E8 | 1,684E8 | PABPC1 |
| Q0V8H6 | Putative helicase MOV-10 OS=Bos taurus OX=9913 GN=MOV10 PE=2 SV=1 - [MOV10_BOVIN] | 0,000E0 | 0,000E0 | 0,000E0 | 9,763E7 | 3,227E7 | 6,965E7 | MOV10 |
| F1MDC1 | Uncharacterized protein OS=Bos taurus OX=9913 PE=1 SV=2 - [F1MDC1_BOVIN] | 0,000E0 | 0,000E0 | 0,000E0 | 1,287E7 | 4,050E8 | 4,074E8 | F1mdc1 |
| E1BEK9 | UPF1, RNA helicase and ATPase OS=Bos taurus OX=9913 GN=UPF1 PE=4 SV=2 - [E1BEK9_BOVIN] | 0,000E0 | 0,000E0 | 0,000E0 | 3,604E7 | 5,350E7 | 1,193E7 | UPF1 |
| F1MBA3 | Integrin alpha-2 OS=Bos taurus OX=9913 GN=ITGA2 PE=3 SV=2 - [F1MBA3_BOVIN] | 1,531E7 | 0,000E0 | 2,417E7 | 9,388E7 | 3,219E7 | 7,963E7 | ITGA2 |
| G3X757 | Transitional endoplasmic reticulum ATPase OS=Bos taurus OX=9913 GN=VCP PE=3 SV=1 - [G3X757_BOVIN] | 0,000E0 | 0,000E0 | 0,000E0 | 3,692E7 | 9,752E7 | 1,599E8 | VCP |
| Q0VCX2 | Endoplasmic reticulum chaperone BiP OS=Bos taurus OX=9913 GN=HSPA5 PE=2 SV=1 - [BIP_BOVIN] | 2,051E7 | 0,000E0 | 2,976E7 | 1,622E8 | 1,787E8 | 1,935E8 | HSPA5 |
| A4FUC2 | HNRNPUL1 protein OS=Bos taurus OX=9913 GN=HNRNPUL1 PE=1 SV=1 - [A4FUC2_BOVIN] | 6,245E7 | 6,810E7 | 0,000E0 | 4,349E8 | 2,130E8 | 6,945E7 | HNRNPUL1 |
| P53712 | Integrin beta-1 OS=Bos taurus OX=9913 GN=ITGB1 PE=1 SV=3 - [ITB1_BOVIN] | 8,016E7 | 4,170E7 | 7,723E7 | 5,275E8 | 3,133E8 | 3,005E8 | ITGB1 |
| **F1MRK3** | **Angiomotin-like protein 2 OS=Bos taurus OX=9913 GN=AMOTL2 PE=2 SV=3 - [AMOL2_BOVIN]** | **0,000E0** | **0,000E0** | **0,000E0** | **1,919E8** | **8,611E7** | **1,409E8** | AMOTL2 |
| Q2HJ60 | Heterogeneous nuclear ribonucleoproteins A2/B1 OS=Bos taurus OX=9913 GN=HNRNPA2B1 PE=2 SV=1 - [ROA2_BOVIN] | 3,004E8 | 5,764E6 | 1,665E8 | 5,180E8 | 4,291E7 | 1,664E7 | HNRNPA2B1 |
| F1MC11 | Keratin, type I cytoskeletal 14 OS=Bos taurus OX=9913 GN=KRT14 PE=1 SV=2 - [F1MC11_BOVIN] | 4,111E8 | 6,366E7 | 2,939E8 | 6,043E8 | 3,424E7 | 1,067E7 | KRT14 |
| F1MM57 | Myosin heavy chain 14 OS=Bos taurus OX=9913 GN=MYH14 PE=1 SV=2 - [F1MM57_BOVIN] | 3,181E7 | 0,000E0 | 0,000E0 | 2,032E9 | 9,526E9 | 1,439E10 | MYH14 |
| E1BET5 | Sperm specific antigen 2 OS=Bos taurus OX=9913 GN=ITPRID2 PE=4 SV=2 - [E1BET5_BOVIN] | 0,000E0 | 0,000E0 | 0,000E0 | 1,495E6 | 1,186E8 | 7,795E7 | ITPRID2 |
| E1BNE7 | Caveolae associated protein 1 OS=Bos taurus OX=9913 GN=CAVIN1 PE=1 SV=1 - [E1BNE7_BOVIN] | 5,895E6 | 0,000E0 | 0,000E0 | 4,678E8 | 4,985E9 | 4,407E9 | CAVIN1 |
| Q1RMR3 | Cortactin OS=Bos taurus OX=9913 GN=CTTN PE=1 SV=1 - [Q1RMR3_BOVIN] | 0,000E0 | 0,000E0 | 0,000E0 | 1,909E8 | 1,216E9 | 1,528E9 | CTTN |
| A7MBI5 | DPYSL3 protein OS=Bos taurus OX=9913 GN=DPYSL3 PE=1 SV=1 - [A7MBI5_BOVIN] | 0,000E0 | 1,643E6 | 7,066E6 | 6,500E7 | 4,821E7 | 4,016E7 | DPYSL3 |
| E1BF20 | Heterogeneous nuclear ribonucleoprotein H1 OS=Bos taurus OX=9913 GN=HNRNPH1 PE=1 SV=2 - [E1BF20_BOVIN] | 3,774E8 | 3,634E7 | 1,022E8 | 1,917E9 | 7,770E8 | 9,580E8 | HNRNPH1 |
| M0QVZ6 | Keratin, type II cytoskeletal 5 OS=Bos taurus OX=9913 GN=KRT5 PE=3 SV=1 - [M0QVZ6_BOVIN] | 1,295E9 | 2,610E7 | 2,830E7 | 4,192E8 | 1,092E7 | 1,227E7 | KRT5 |
| Q5KR47 | Tropomyosin alpha-3 chain OS=Bos taurus OX=9913 GN=TPM3 PE=2 SV=1 - [TPM3_BOVIN] | 0,000E0 | 0,000E0 | 0,000E0 | 1,349E9 | 8,165E9 | 6,151E9 | TPM3 |
| A7E307 | DDX17 protein OS=Bos taurus OX=9913 GN=DDX17 PE=1 SV=1 - [A7E307_BOVIN] | 0,000E0 | 0,000E0 | 0,000E0 | 7,125E7 | 1,596E8 | 1,779E8 | DDX17 |
| F1MBQ8 | DEAD-box helicase 5 OS=Bos taurus OX=9913 GN=DDX5 PE=3 SV=2 - [F1MBQ8_BOVIN] | 0,000E0 | 0,000E0 | 0,000E0 | 1,226E8 | 2,143E8 | 2,056E8 | DDX5 |
| F1N3L7 | Dedicator of cytokinesis 7 OS=Bos taurus OX=9913 GN=DOCK7 PE=3 SV=2 - [F1N3L7_BOVIN] | 0,000E0 | 0,000E0 | 0,000E0 | 6,523E6 | 6,343E7 | 3,671E7 | DOCK7 |
| K9ZRL4 | Heat shock 70kDa protein 1A OS=Bos indicus x Bos taurus OX=30522 GN=HSPA1A PE=2 SV=1 - [K9ZRL4_BOBOX] | 3,446E7 | 0,000E0 | 2,976E7 | 2,943E8 | 4,032E8 | 4,354E8 | HSPA1A |
| P79136-2 | Isoform Beta-2 of F-actin-capping protein subunit beta OS=Bos taurus OX=9913 GN=CAPZB - [CAPZB_BOVIN] | 0,000E0 | 0,000E0 | 0,000E0 | 2,853E8 | 1,487E9 | 2,174E9 | CAPZB |
| A8E4L8 | LIM domain 7 OS=Bos taurus OX=9913 GN=LMO7 PE=1 SV=1 - [A8E4L8_BOVIN] | 0,000E0 | 0,000E0 | 0,000E0 | 5,743E6 | 7,540E7 | 1,125E8 | LMO7 |
| A6QR15 | LOC535277 protein OS=Bos taurus OX=9913 GN=TPM4 PE=1 SV=1 - [A6QR15_BOVIN] | 0,000E0 | 0,000E0 | 0,000E0 | 1,049E9 | 5,247E9 | 4,464E9 | TPM4 |
| E1BKQ6 | Myosin XVIIIA OS=Bos taurus OX=9913 GN=MYO18A PE=3 SV=2 - [E1BKQ6_BOVIN] | 0,000E0 | 0,000E0 | 0,000E0 | 4,855E6 | 1,231E8 | 1,014E8 | MYO18A |
| Q3SX36 | TFG protein OS=Bos taurus OX=9913 GN=TFG PE=2 SV=1 - [Q3SX36_BOVIN] | 7,425E8 | 1,859E8 | 8,084E7 | 3,202E9 | 5,176E8 | 9,231E8 | TFG |
| F1N3A1 | Thrombospondin-1 OS=Bos taurus OX=9913 GN=THBS1 PE=4 SV=1 - [F1N3A1_BOVIN] | 9,682E5 | 0,000E0 | 0,000E0 | 1,277E7 | 1,156E8 | 1,095E8 | THBS1 |
| F1MTP5 | WD repeat-containing protein 1 OS=Bos taurus OX=9913 GN=WDR1 PE=4 SV=2 - [F1MTP5_BOVIN] | 0,000E0 | 0,000E0 | 0,000E0 | 2,044E7 | 3,156E8 | 4,507E8 | WDR1 |
| Q3MHR7 | Actin-related protein 2/3 complex subunit 2 OS=Bos taurus OX=9913 GN=ARPC2 PE=1 SV=1 - [ARPC2_BOVIN] | 0,000E0 | 0,000E0 | 0,000E0 | 8,576E7 | 5,033E8 | 9,465E8 | ARPC2 |
| P61157 | Actin-related protein 3 OS=Bos taurus OX=9913 GN=ACTR3 PE=1 SV=3 - [ARP3_BOVIN] | 0,000E0 | 0,000E0 | 0,000E0 | 1,341E8 | 2,483E8 | 1,186E8 | ACTR3 |
| A8E654 | COL18A1 protein OS=Bos taurus OX=9913 GN=COL18A1 PE=2 SV=1 - [A8E654_BOVIN] | 0,000E0 | 0,000E0 | 0,000E0 | 5,072E7 | 3,067E8 | 3,942E8 | COL18A1 |
| F1ME65 | Cytoskeleton associated protein 4 OS=Bos taurus OX=9913 GN=CKAP4 PE=1 SV=1 - [F1ME65_BOVIN] | 0,000E0 | 0,000E0 | 0,000E0 | 1,215E7 | 9,264E7 | 1,071E8 | CKAP4 |
| Q7YRV4 | E3 ubiquitin-protein ligase TRIM21 OS=Bos taurus OX=9913 GN=TRIM21 PE=2 SV=1 - [RO52_BOVIN] | 0,000E0 | 0,000E0 | 0,000E0 | 8,054E7 | 1,599E8 | 5,598E7 | TRIM21 |
| E1BPW2 | ELKS/RAB6-interacting/CAST family member 1 OS=Bos taurus OX=9913 GN=ERC1 PE=4 SV=2 - [E1BPW2_BOVIN] | 0,000E0 | 0,000E0 | 0,000E0 | 1,419E7 | 9,191E6 | 5,451E6 | ERC1 |
| A7MB80 | General transcription factor II-I OS=Bos taurus OX=9913 GN=GTF2I PE=2 SV=1 - [GTF2I_BOVIN] | 0,000E0 | 0,000E0 | 0,000E0 | 1,851E7 | 2,968E7 | 2,869E7 | GTF2I |
| F1MD77 | Laminin subunit gamma 1 OS=Bos taurus OX=9913 GN=LAMC1 PE=1 SV=2 - [F1MD77_BOVIN] | 0,000E0 | 0,000E0 | 0,000E0 | 1,297E7 | 1,487E8 | 1,619E8 | LAMC1 |
| F1MP70 | Leucine-rich repeat flightless-interacting protein 2 OS=Bos taurus OX=9913 GN=LRRFIP2 PE=4 SV=1 - [F1MP70_BOVIN] | 0,000E0 | 0,000E0 | 0,000E0 | 7,899E7 | 1,612E8 | 1,175E8 | LRRFIP2 |
| E1BDB1 | Ras interacting protein 1 OS=Bos taurus OX=9913 GN=RASIP1 PE=4 SV=1 - [E1BDB1_BOVIN] | 0,000E0 | 0,000E0 | 0,000E0 | 3,113E7 | 1,703E8 | 1,564E8 | RASIP1 |
| F1MI47 | RNA-binding protein 14 OS=Bos taurus OX=9913 GN=RBM14 PE=4 SV=1 - [F1MI47_BOVIN] | 0,000E0 | 1,814E7 | 0,000E0 | 6,851E7 | 7,143E7 | 5,077E7 | RBM14 |
| F1MUZ9 | 60 kDa heat shock protein, mitochondrial OS=Bos taurus OX=9913 GN=HSPD1 PE=1 SV=1 - [F1MUZ9_BOVIN] | 0,000E0 | 0,000E0 | 0,000E0 | 2,846E7 | 5,329E7 | 5,990E7 | HSPD1 |
| A7YWC4 | ATPase family AAA domain-containing protein 3 OS=Bos taurus OX=9913 GN=ATAD3 PE=2 SV=1 - [ATAD3_BOVIN] | 0,000E0 | 0,000E0 | 0,000E0 | 2,815E7 | 4,185E7 | 5,686E7 | ATAD3 |
| A5D9B4 | Heterogeneous nuclear ribonucleoprotein H2 OS=Bos taurus OX=9913 GN=HNRNPH2 PE=2 SV=1 - [A5D9B4_BOVIN] | 3,774E8 | 3,634E7 | 3,481E7 | 1,917E9 | 7,770E8 | 7,314E8 | HNRNPH2 |
| E1BIB4 | Heterogeneous nuclear ribonucleoprotein L OS=Bos taurus OX=9913 GN=HNRNPL PE=1 SV=2 - [E1BIB4_BOVIN] | 0,000E0 | 0,000E0 | 0,000E0 | 9,256E7 | 2,010E7 | 3,332E7 | HNRNPL |
| A4IFC3 | Polyadenylate-binding protein OS=Bos taurus OX=9913 GN=PABPC4 PE=2 SV=1 - [A4IFC3_BOVIN] | 0,000E0 | 0,000E0 | 0,000E0 | 1,343E8 | 1,461E8 | 6,130E7 | PABPC4 |
| P63243 | Receptor of activated protein C kinase 1 OS=Bos taurus OX=9913 GN=RACK1 PE=2 SV=3 - [RACK1_BOVIN] | 7,575E6 | 0,000E0 | 0,000E0 | 2,907E8 | 4,730E8 | 2,448E8 | RACK1 |
| E1BA93 | Synaptopodin OS=Bos taurus OX=9913 GN=SYNPO PE=4 SV=2 - [E1BA93_BOVIN] | 0,000E0 | 0,000E0 | 0,000E0 | 2,388E7 | 2,559E8 | 1,774E8 | SYNPO |
| A7E355 | SYNCRIP protein (Fragment) OS=Bos taurus OX=9913 GN=SYNCRIP PE=2 SV=1 - [A7E355_BOVIN] | 0,000E0 | 0,000E0 | 1,715E7 | 2,762E7 | 7,609E6 | 1,022E7 | SYNCRIP |
| F1MBN2 | Tight junction protein 2 OS=Bos taurus OX=9913 GN=TJP2 PE=1 SV=2 - [F1MBN2_BOVIN] | 0,000E0 | 0,000E0 | 0,000E0 | 1,036E7 | 3,376E7 | 3,495E7 | TJP2 |
| E1BGI2 | Centrosomal protein 170 OS=Bos taurus OX=9913 GN=CEP170 PE=4 SV=2 - [E1BGI2_BOVIN] | 0,000E0 | 0,000E0 | 0,000E0 | 2,188E7 | 4,878E7 | 5,161E7 | CEP170 |
| E1BK77 | Dedicator of cytokinesis 6 OS=Bos taurus OX=9913 GN=DOCK6 PE=3 SV=2 - [E1BK77_BOVIN] | 0,000E0 | 0,000E0 | 0,000E0 | 1,966E6 | 5,692E7 | 2,415E7 | DOCK6 |
| Q5E997 | F-actin-capping protein subunit alpha-2 OS=Bos taurus OX=9913 GN=CAPZA2 PE=1 SV=3 - [CAZA2_BOVIN] | 0,000E0 | 0,000E0 | 0,000E0 | 8,090E7 | 1,144E8 | 1,183E8 | CAPZA2 |
| F1MF97 | Nidogen 2 OS=Bos taurus OX=9913 GN=NID2 PE=4 SV=2 - [F1MF97_BOVIN] | 0,000E0 | 0,000E0 | 0,000E0 | 8,613E5 | 1,788E8 | 1,692E8 | NID2 |
| Q3SZN2 | Protein transport protein Sec23B OS=Bos taurus OX=9913 GN=SEC23B PE=2 SV=1 - [SC23B_BOVIN] | 2,526E7 | 0,000E0 | 0,000E0 | 4,841E7 | 4,278E6 | 5,889E6 | SEC23B |
| A6QQF3 | TJP2 protein OS=Bos taurus OX=9913 GN=TJP2 PE=2 SV=1 - [A6QQF3_BOVIN] | 0,000E0 | 0,000E0 | 0,000E0 | 1,036E7 | 1,655E7 | 2,481E7 | TJP2 |
| Q2KJH0 | Tropomodulin 3 (Ubiquitous) OS=Bos taurus OX=9913 GN=TMOD3 PE=2 SV=1 - [Q2KJH0_BOVIN] | 0,000E0 | 0,000E0 | 0,000E0 | 8,792E7 | 1,651E9 | 9,341E8 | TMOD3 |
| F1MY44 | Uncharacterized protein OS=Bos taurus OX=9913 GN=HNRNPM PE=1 SV=2 - [F1MY44_BOVIN] | 0,000E0 | 0,000E0 | 0,000E0 | 2,418E7 | 5,831E7 | 6,064E7 | HNRNPM |
| E1BCF8 | Zinc finger CCCH-type containing, antiviral 1 OS=Bos taurus OX=9913 GN=ZC3HAV1 PE=4 SV=1 - [E1BCF8_BOVIN] | 0,000E0 | 0,000E0 | 0,000E0 | 1,352E7 | 2,264E7 | 3,873E7 | ZC3HAV1 |

| | | | | | | | | |
|---|---|---|---|---|---|---|---|---|
| Q3T169 | 40S ribosomal protein S3 OS=Bos taurus OX=9913 GN=RPS3 PE=2 SV=1 - [RS3_BOVIN] | 8,535E6 | 8,993E6 | 5,061E6 | 1,842E8 | 5,501E8 | 5,680E8 | RPS3 |
| Q56JV9 | 40S ribosomal protein S3a OS=Bos taurus OX=9913 GN=RPS3A PE=2 SV=3 - [RS3A_BOVIN] | 0,000E0 | 0,000E0 | 0,000E0 | 8,427E7 | 4,585E8 | 2,354E8 | RPS3A |
| A7MB62 | Actin-related protein 2 OS=Bos taurus OX=9913 GN=ACTR2 PE=1 SV=1 - [ARP2_BOVIN] | 0,000E0 | 0,000E0 | 0,000E0 | 2,174E8 | 1,042E9 | 8,179E8 | ACTR2 |
| Q0IIK5 | ATP-dependent RNA helicase DDX1 OS=Bos taurus OX=9913 GN=DDX1 PE=2 SV=1 - [DDX1_BOVIN] | 0,000E0 | 0,000E0 | 0,000E0 | 1,787E7 | 1,344E8 | 1,009E8 | DDX1 |
| A6H7H1 | Constitutive coactivator of PPAR-gamma-like protein 1 OS=Bos taurus OX=9913 GN=FAM120A PE=2 SV=1 - [F120A_BOVIN] | 0,000E0 | 0,000E0 | 0,000E0 | 7,518E6 | 2,471E6 | 4,423E7 | FAM120A |
| Q76LV1 | Heat shock protein HSP 90-beta OS=Bos taurus OX=9913 GN=HSP90AB1 PE=2 SV=3 - [HS90B_BOVIN] | 1,281E7 | 0,000E0 | 0,000E0 | 3,924E7 | 2,724E7 | 1,981E7 | HSP90AB1 |
| P09867 | Heterogeneous nuclear ribonucleoprotein A1 OS=Bos taurus OX=9913 GN=HNRNPA1 PE=1 SV=2 - [ROA1_BOVIN] | 8,950E6 | 0,000E0 | 0,000E0 | 7,875E7 | 1,325E8 | 1,223E8 | HNRNPA1 |
| E1BEG2 | Heterogeneous nuclear ribonucleoprotein A3 OS=Bos taurus OX=9913 GN=HNRNPA3 PE=1 SV=2 - [E1BEG2_BOVIN] | 3,383E7 | 0,000E0 | 0,000E0 | 3,610E8 | 1,092E7 | 3,142E8 | HNRNPA3 |
| E1BB17 | Heterogeneous nuclear ribonucleoprotein H3 OS=Bos taurus OX=9913 GN=HNRNPH3 PE=1 SV=1 - [E1BB17_BOVIN] | 8,945E7 | 0,000E0 | 1,443E7 | 3,082E8 | 5,168E6 | 3,529E6 | HNRNPH3 |
| Q3T0D0 | Heterogeneous nuclear ribonucleoprotein K OS=Bos taurus OX=9913 GN=HNRNPK PE=2 SV=1 - [HNRPK_BOVIN] | 1,081E7 | 0,000E0 | 0,000E0 | 1,077E8 | 4,562E7 | 6,817E7 | HNRNPK |
| F1MMS9 | Integrin alpha-3 OS=Bos taurus OX=9913 GN=ITGA3 PE=3 SV=1 - [ITA3_BOVIN] | 2,909E7 | 3,732E7 | 3,998E7 | 1,858E8 | 2,556E7 | 4,198E7 | ITGA3 |
| B0JYP5 | Protein argonaute-2 OS=Bos taurus OX=9913 GN=AGO2 PE=2 SV=1 - [B0JYP5_BOVIN] | 0,000E0 | 0,000E0 | 0,000E0 | 4,797E7 | 7,436E7 | 4,258E7 | AGO2 |
| F1MVW5 | Protein transport protein SEC23 OS=Bos taurus OX=9913 GN=SEC23A PE=3 SV=2 - [F1MVW5_BOVIN] | 3,039E7 | 0,000E0 | 0,000E0 | 3,944E7 | 8,283E7 | 3,504E7 | SEC23A |
| Q5E9T9 | tRNA-splicing ligase RtcB homolog OS=Bos taurus OX=9913 GN=RTCB PE=2 SV=1 - [RTCB_BOVIN] | 0,000E0 | 0,000E0 | 0,000E0 | 4,540E7 | 1,468E8 | 6,082E7 | RTCB |
| F1N6C0 | Uncharacterized protein OS=Bos taurus OX=9913 GN=CALM2 PE=4 SV=2 - [F1N6C0_BOVIN] | 0,000E0 | 0,000E0 | 0,000E0 | 3,826E8 | 1,006E9 | 1,923E9 | CALM2 |
| Q32PD5 | 40S ribosomal protein S19 OS=Bos taurus OX=9913 GN=RPS19 PE=2 SV=3 - [RS19_BOVIN] | 0,000E0 | 0,000E0 | 0,000E0 | 1,928E8 | 5,144E8 | 1,205E8 | RPS19 |
| Q58DW0 | 60S ribosomal protein L4 OS=Bos taurus OX=9913 GN=RPL4 PE=2 SV=3 - [RL4_BOVIN] | 0,000E0 | 0,000E0 | 0,000E0 | 3,119E7 | 2,577E8 | 1,279E8 | RPL4 |
| Q1JPK9 | Actin related protein 2/3 complex subunit 1B (Fragment) OS=Bos taurus OX=9913 GN=ARPC1B PE=2 SV=1 - [Q1JPK9_BOVIN] | 0,000E0 | 0,000E0 | 0,000E0 | 8,267E7 | 3,873E8 | 1,994E8 | ARPC1B |
| G5E5N3 | ATP-dependent RNA helicase A OS=Bos taurus OX=9913 GN=DHX9 PE=4 SV=1 - [G5E5N3_BOVIN] | 0,000E0 | 0,000E0 | 0,000E0 | 1,991E7 | 5,330E7 | 6,292E7 | DHX9 |
| A4FUA8 | F-actin-capping protein subunit alpha-1 OS=Bos taurus OX=9913 GN=CAPZA1 PE=2 SV=1 - [CAZA1_BOVIN] | 0,000E0 | 0,000E0 | 0,000E0 | 3,232E8 | 1,746E9 | 1,373E9 | CAPZA1 |
| Q8WMQ6 | Filamin A (Fragment) OS=Bos taurus OX=9913 PE=2 SV=1 - [Q8WMQ6_BOVIN] | 0,000E0 | 0,000E0 | 0,000E0 | 1,438E7 | 1,104E8 | 3,854E7 | FLNA |
| A2VDN7 | Heterogeneous nuclear ribonucleoprotein U (Scaffold attachment factor A) OS=Bos taurus OX=9913 GN=HNRNPU PE=2 SV=1 - [A2VDN7_BOVIN] | 0,000E0 | 0,000E0 | 0,000E0 | 9,752E7 | 2,376E8 | 1,282E8 | HNRNPU |
| A3KMV6 | HNRNPR protein OS=Bos taurus OX=9913 GN=HNRNPR PE=1 SV=1 - [A3KMV6_BOVIN] | 0,000E0 | 0,000E0 | 1,715E7 | 4,367E7 | 3,361E7 | 1,057E7 | HNRNPR |
| P60661-2 | Isoform Smooth muscle of Myosin light polypeptide 6 OS=Bos taurus OX=9913 GN=MYL6 - [MYL6_BOVIN] | 7,341E6 | 1,354E7 | 0,000E0 | 3,112E9 | 1,484E10 | 1,581E10 | MYL6 |
| F1MXX0 | MICOS complex subunit MIC60 OS=Bos taurus OX=9913 GN=IMMT PE=1 SV=1 - [F1MXX0_BOVIN] | 0,000E0 | 0,000E0 | 0,000E0 | 6,081E6 | 1,407E8 | 1,487E8 | IMMT |
| P60661 | Myosin light polypeptide 6 OS=Bos taurus OX=9913 GN=MYL6 PE=2 SV=2 - [MYL6_BOVIN] | 7,341E6 | 1,354E7 | 0,000E0 | 3,112E9 | 1,484E10 | 1,581E10 | MYL6 |
| F1MZ92 | Nuclease-sensitive element-binding protein 1 OS=Bos taurus OX=9913 GN=YBX1 PE=4 SV=2 - [F1MZ92_BOVIN] | 2,620E7 | 0,000E0 | 5,410E7 | 1,238E9 | 7,883E8 | 1,185E9 | YBX1 |
| A0A140T861 | RNA-binding protein FUS OS=Bos taurus OX=9913 GN=FUS PE=4 SV=1 - [A0A140T861_BOVIN] | 9,088E8 | 0,000E0 | 5,559E8 | 1,321E10 | 1,780E9 | 2,558E9 | FUS |
| E1BIU0 | SEC24 homolog C, COPII coat complex component OS=Bos taurus OX=9913 GN=SEC24C PE=1 SV=1 - [E1BIU0_BOVIN] | 1,279E7 | 1,198E7 | 1,106E7 | 8,100E7 | 6,415E7 | 5,408E7 | SEC24C |
| E1BGX1 | TATA-box binding protein associated factor 15 OS=Bos taurus OX=9913 GN=TAF15 PE=4 SV=2 - [E1BGX1_BOVIN] | 2,706E8 | 5,179E6 | 5,559E8 | 1,737E8 | 1,158E9 | 9,225E8 | TAF15 |
| F2Z4H8 | TIA1 cytotoxic granule associated RNA binding protein like 1 OS=Bos taurus OX=9913 GN=TIAL1 PE=4 SV=2 - [F2Z4H8_BOVIN] | 3,259E7 | 0,000E0 | 6,197E7 | 1,303E8 | 5,732E6 | 1,610E8 | TIAL1 |
| E1BJB1 | Tubulin beta chain OS=Bos taurus OX=9913 GN=TUBB2A PE=1 SV=1 - [E1BJB1_BOVIN] | 2,869E7 | 0,000E0 | 0,000E0 | 6,851E7 | 1,284E8 | 1,337E8 | TUBB2A |
| Q3MHM5 | Tubulin beta-4B chain OS=Bos taurus OX=9913 GN=TUBB4B PE=2 SV=1 - [TBB4B_BOVIN] | 2,869E7 | 0,000E0 | 0,000E0 | 8,312E7 | 1,406E8 | 1,337E8 | TUBB4B |
| Q2KJD0 | Tubulin beta-5 chain OS=Bos taurus OX=9913 GN=TUBB5 PE=2 SV=1 - [TBB5_BOVIN] | 2,220E7 | 0,000E0 | 0,000E0 | 9,628E7 | 1,479E8 | 1,067E8 | TUBB5 |
| F1MYI2 | Uncharacterized protein OS=Bos taurus OX=9913 GN=HNRNPM PE=1 SV=2 - [F1MYI2_BOVIN] | 0,000E0 | 0,000E0 | 0,000E0 | 2,372E7 | 4,213E7 | 4,233E7 | HNRNPM |
| A5D7Q4 | CSDA protein OS=Bos taurus OX=9913 GN=YBX3 PE=1 SV=1 - [A5D7Q4_BOVIN] | 3,000E7 | 0,000E0 | 0,000E0 | 8,555E8 | 2,257E8 | 7,208E8 | YBX3 |
| F1MJH1 | Gelsolin OS=Bos taurus OX=9913 GN=GSN PE=4 SV=1 - [F1MJH1_BOVIN] | 1,209E7 | 0,000E0 | 0,000E0 | 1,101E8 | 2,819E8 | 8,336E7 | GSN |
| A0A088Q0F1 | Heat shock protein 90kDa alpha OS=Bos taurus x Bos grunniens OX=1167295 PE=2 SV=1 - [A0A088Q0F1_9CETA] | 0,000E0 | 0,000E0 | 0,000E0 | 4,405E7 | 1,283E7 | 1,617E7 | HSP90AB1 |
| Q5E9J1 | Heterogeneous nuclear ribonucleoprotein F OS=Bos taurus OX=9913 GN=HNRNPF PE=2 SV=3 - [HNRPF_BOVIN] | 2,343E8 | 0,000E0 | 0,000E0 | 7,057E8 | 2,517E7 | 2,077E7 | HNRNPF |
| F1MTN1 | Integrin beta OS=Bos taurus OX=9913 GN=ITGB3 PE=3 SV=1 - [F1MTN1_BOVIN] | 0,000E0 | 0,000E0 | 0,000E0 | 3,463E7 | 1,371E7 | 1,751E7 | ITGB3 |
| F1MNT4 | Laminin subunit beta 1 OS=Bos taurus OX=9913 GN=LAMB1 PE=4 SV=2 - [F1MNT4_BOVIN] | 0,000E0 | 0,000E0 | 0,000E0 | 6,481E6 | 3,346E7 | 1,099E7 | LAMB1 |
| A4IF97 | Myosin regulatory light chain 12B OS=Bos taurus OX=9913 GN=MYL12B PE=2 SV=1 - [ML12B_BOVIN] | 1,003E7 | 0,000E0 | 0,000E0 | 4,282E9 | 1,172E10 | 8,380E9 | MYL12B |
| F1MU08 | Poly [ADP-ribose] polymerase OS=Bos taurus OX=9913 GN=PARP1 PE=4 SV=2 - [F1MU08_BOVIN] | 0,000E0 | 0,000E0 | 0,000E0 | 1,386E7 | 9,516E7 | 1,115E8 | PARP1 |
| F1N6W6 | Sorting nexin OS=Bos taurus OX=9913 GN=SNX9 PE=3 SV=2 - [F1N6W6_BOVIN] | 4,182E7 | 6,389E7 | 5,097E6 | 2,603E8 | 2,083E8 | 2,120E8 | SNX9 |
| F2Z4C1 | Tubulin alpha chain OS=Bos taurus OX=9913 GN=TUBA1A PE=3 SV=1 - [F2Z4C1_BOVIN] | 2,350E7 | 0,000E0 | 0,000E0 | 1,890E8 | 1,757E8 | 9,255E7 | TUBA1A |
| P81947 | Tubulin alpha-1B chain OS=Bos taurus OX=9913 PE=1 SV=2 - [TBA1B_BOVIN] | 2,350E7 | 0,000E0 | 0,000E0 | 1,890E8 | 1,757E8 | 9,255E7 | TUBA1B |
| Q3T0X6 | 40S ribosomal protein S16 OS=Bos taurus OX=9913 GN=RPS16 PE=2 SV=3 - [RS16_BOVIN] | 6,254E6 | 0,000E0 | 0,000E0 | 2,518E7 | 3,647E7 | 4,029E7 | RPS16 |
| Q58DT1 | 60S ribosomal protein L7 OS=Bos taurus OX=9913 GN=RPL7 PE=2 SV=1 - [RL7_BOVIN] | 0,000E0 | 0,000E0 | 0,000E0 | 6,401E7 | 7,190E8 | 8,174E8 | RPL7 |
| A4FV37 | Caveolae-associated protein 3 OS=Bos taurus OX=9913 GN=CAVIN3 PE=2 SV=1 - [CAVN3_BOVIN] | 0,000E0 | 0,000E0 | 0,000E0 | 1,970E6 | 2,871E7 | 2,432E7 | CAVIN3 |
| E1BDM8 | DEAD-box helicase 6 OS=Bos taurus OX=9913 GN=DDX6 PE=1 SV=1 - [E1BDM8_BOVIN] | 0,000E0 | 0,000E0 | 0,000E0 | 3,633E7 | 2,172E7 | 6,322E7 | DDX6 |
| A6QNZ7 | Keratin 10 (Epidermolytic hyperkeratosis; keratosis palmaris et plantaris) OS=Bos taurus OX=9913 GN=KRT10 PE=2 SV=1 - [A6QNZ7_BOVIN] | 5,671E8 | 5,145E8 | 2,270E7 | 3,371E9 | 9,223E7 | 5,738E8 | KRT10 |
| G3X6B3 | Proline rich coiled-coil 2C OS=Bos taurus OX=9913 GN=PRRC2C PE=4 SV=1 - [G3X6B3_BOVIN] | 0,000E0 | 0,000E0 | 0,000E0 | 2,014E6 | 1,997E7 | 1,534E7 | PRRC2C |
| A1L5B7 | SERPINE1 mRNA binding protein 1 OS=Bos taurus OX=9913 GN=SERBP1 PE=1 SV=1 - [A1L5B7_BOVIN] | 0,000E0 | 0,000E0 | 0,000E0 | 2,193E7 | 1,771E7 | 3,206E7 | SERBP1 |
| Q862S8 | Similar to ribosomal protein S18 (Fragment) OS=Bos taurus OX=9913 PE=2 SV=1 - [Q862S8_BOVIN] | 0,000E0 | 0,000E0 | 8,945E6 | 5,566E7 | 7,652E6 | 8,803E7 | RPS18 |
| Q0VBZ6 | TIA1 cytotoxic granule-associated RNA binding protein OS=Bos taurus OX=9913 GN=TIA1 PE=2 SV=1 - [Q0VBZ6_BOVIN] | 3,259E7 | 0,000E0 | 6,825E7 | 1,655E8 | 6,116E7 | 2,315E8 | TIA1 |
| A1L505 | Ubiquitin associated protein 2-like OS=Bos taurus OX=9913 GN=UBAP2L PE=2 SV=1 - [A1L505_BOVIN] | 8,334E6 | 0,000E0 | 0,000E0 | 6,526E7 | 3,231E7 | 3,661E7 | UBAP2L |
| E1B8K6 | Uncharacterized protein OS=Bos taurus OX=9913 GN=NCL PE=1 SV=1 - [E1B8K6_BOVIN] | 0,000E0 | 0,000E0 | 0,000E0 | 3,799E7 | 2,301E8 | 1,156E8 | NCL |
| E1BP85 | Uridine-cytidine kinase OS=Bos taurus OX=9913 GN=UCKL1 PE=3 SV=1 - [E1BP85_BOVIN] | 0,000E0 | 0,000E0 | 0,000E0 | 2,367E7 | 9,340E6 | 9,001E6 | UCKL1 |
| P26452 | 40S ribosomal protein SA OS=Bos taurus OX=9913 GN=RPSA PE=2 SV=4 - [RSSA_BOVIN] | 0,000E0 | 0,000E0 | 0,000E0 | 3,791E8 | 1,371E9 | 4,922E8 | RPSA |
| Q95140 | 60S acidic ribosomal protein P0 OS=Bos taurus OX=9913 GN=RPLP0 PE=2 SV=3 - [RLA0_BOVIN] | 0,000E0 | 0,000E0 | 0,000E0 | 1,084E8 | 7,248E7 | 2,108E8 | RPLP0 |
| P61284 | 60S ribosomal protein L12 OS=Bos taurus OX=9913 GN=RPL12 PE=2 SV=1 - [RL12_BOVIN] | 0,000E0 | 0,000E0 | 0,000E0 | 2,219E8 | 3,845E7 | 3,384E7 | RPL12 |
| Q58DW5 | 60S ribosomal protein L5 OS=Bos taurus OX=9913 GN=RPL5 PE=2 SV=3 - [RL5_BOVIN] | 0,000E0 | 0,000E0 | 0,000E0 | 1,071E8 | 9,474E8 | 7,394E8 | RPL5 |
| Q4VQD6 | Actin (Fragment) OS=Cryptosporidium sp. ex Bos taurus OX=310049 PE=3 SV=1 - [Q4VQD6_9CRYT] | 1,203E8 | 0,000E0 | 0,000E0 | 2,143E9 | 3,818E9 | 3,586E9 | ACTB |
| Q148J6 | Actin-related protein 2/3 complex subunit 4 OS=Bos taurus OX=9913 GN=ARPC4 PE=1 SV=3 - [ARPC4_BOVIN] | 0,000E0 | 0,000E0 | 0,000E0 | 4,048E7 | 5,205E8 | 5,024E8 | ARPC4 |
| Q8SQH5 | ADP/ATP translocase 2 OS=Bos taurus OX=9913 GN=SLC25A5 PE=2 SV=3 - [ADT2_BOVIN] | 1,971E7 | 0,000E0 | 0,000E0 | 7,164E7 | 1,573E7 | 7,765E7 | SLC25A5 |
| P32007 | ADP/ATP translocase 3 OS=Bos taurus OX=9913 GN=SLC25A6 PE=1 SV=3 - [ADT3_BOVIN] | 1,541E7 | 0,000E0 | 0,000E0 | 3,837E7 | 1,573E7 | 7,765E7 | SLC25A6 |
| P04272 | Annexin A2 OS=Bos taurus OX=9913 GN=ANXA2 PE=1 SV=2 - [ANXA2_BOVIN] | 7,341E6 | 0,000E0 | 0,000E0 | 6,681E7 | 1,021E8 | 3,280E7 | ANXA2 |
| P63009 | AP-2 complex subunit beta OS=Bos taurus OX=9913 GN=AP2B1 PE=1 SV=2 - [AP2B1_BOVIN] | 0,000E0 | 0,000E0 | 0,000E0 | 1,060E7 | 1,097E8 | 7,708E7 | AP2B1 |
| F1MBT5 | BCR, RhoGEF and GTPase activating protein OS=Bos taurus OX=9913 GN=BCR PE=4 SV=2 - [F1MBT5_BOVIN] | 0,000E0 | 0,000E0 | 0,000E0 | 2,689E7 | 1,963E7 | 3,703E7 | BCR |
| F1MPH9 | Caprin-1 OS=Bos taurus OX=9913 GN=CAPRIN1 PE=4 SV=2 - [F1MPH9_BOVIN] | 0,000E0 | 0,000E0 | 0,000E0 | 1,178E8 | 2,185E8 | 1,780E8 | CAPRIN1 |
| P79132 | Caveolin-1 OS=Bos taurus OX=9913 GN=CAV1 PE=2 SV=1 - [CAV1_BOVIN] | 0,000E0 | 0,000E0 | 0,000E0 | 5,979E7 | 3,293E8 | 2,998E8 | CAV1 |
| A0A0F6QNP | Complement component 3 OS=Bos taurus OX=9913 GN=C3 PE=2 SV=1 - [A0A0F6QNP7_BOVIN] | 9,582E6 | 0,000E0 | 0,000E0 | 7,754E7 | 5,523E6 | 2,025E8 | C3 |
| P11179 | Dihydrolipoyllysine-residue succinyltransferase component of 2-oxoglutarate dehydrogenase complex, mitochondrial OS=Bos taurus OX=9913 GN=DLST PE=1 SV=2 - [ODO2_BOVIN] | 0,000E0 | 2,749E6 | 0,000E0 | 3,927E7 | 4,756E7 | 4,977E7 | DLST |
| P68103 | Elongation factor 1-alpha 1 OS=Bos taurus OX=9913 GN=EEF1A1 PE=1 SV=1 - [EF1A1_BOVIN] | 4,290E7 | 0,000E0 | 0,000E0 | 2,024E8 | 5,449E6 | 8,357E7 | EEF1A1 |
| E1B7R4 | Eukaryotic translation initiation factor 3 subunit A OS=Bos taurus OX=9913 GN=EIF3A PE=3 SV=2 - [E1B7R4_BOVIN] | 0,000E0 | 0,000E0 | 0,000E0 | 2,726E7 | 3,690E7 | 1,037E8 | EIF3A |
| Q3SYW6 | Eukaryotic translation initiation factor 3 subunit C OS=Bos taurus OX=9913 GN=EIF3C PE=2 SV=1 - [EIF3C_BOVIN] | 0,000E0 | 0,000E0 | 0,000E0 | 2,069E7 | 3,997E7 | 5,894E7 | EIF3C |
| A8E651 | EWS RNA binding protein 1 OS=Bos taurus OX=9913 GN=EWSR1 PE=2 SV=1 - [A8E651_BOVIN] | 1,273E8 | 1,041E7 | 4,251E7 | 1,902E8 | 9,836E5 | 8,826E7 | EWSR1 |
| Q3ZC44 | Heterogeneous nuclear ribonucleoprotein A/B OS=Bos taurus OX=9913 GN=HNRNPAB PE=1 SV=1 - [Q3ZC44_BOVIN] | 3,097E7 | 0,000E0 | 0,000E0 | 2,817E8 | 2,237E8 | 1,885E8 | HNRNPAB |
| Q5E949 | HMT1 hnRNP methyltransferase-like 2 isoform 3 OS=Bos taurus OX=9913 GN=PRMT1 PE=2 SV=1 - [Q5E949_BOVIN] | 0,000E0 | 0,000E0 | 0,000E0 | 8,383E7 | 4,567E7 | 4,871E7 | PRMT1 |
| Q2HJH4 | HNRPDL protein (Fragment) OS=Bos taurus OX=9913 GN=HNRPDL PE=2 SV=1 - [Q2HJH4_BOVIN] | 4,361E7 | 0,000E0 | 0,000E0 | 4,148E8 | 1,181E8 | 1,118E8 | HNRPDL |
| F1MMN6 | Integrin subunit alpha 1 OS=Bos taurus OX=9913 GN=ITGA1 PE=3 SV=2 - [F1MMN6_BOVIN] | 2,087E7 | 0,000E0 | 0,000E0 | 3,219E7 | 1,607E6 | 8,844E5 | ITGA1 |
| G3N0V2 | Keratin 1 OS=Bos taurus OX=9913 GN=KRT1 PE=1 SV=1 - [G3N0V2_BOVIN] | 3,489E8 | 3,147E7 | 1,634E7 | 2,256E9 | 4,360E7 | 6,381E7 | KRT1 |
| P08728 | Keratin, type I cytoskeletal 19 OS=Bos taurus OX=9913 GN=KRT19 PE=2 SV=1 - [K1C19_BOVIN] | 8,877E7 | 0,000E0 | 2,196E6 | 4,416E8 | 3,785E7 | 1,190E7 | KRT19 |
| G3MZU4 | Myosin XIX OS=Bos taurus OX=9913 GN=MYO19 PE=3 SV=1 - [G3MZU4_BOVIN] | 0,000E0 | 0,000E0 | 0,000E0 | 4,272E6 | 2,818E7 | 5,218E7 | MYO19 |
| E1BK63 | Ribosomal protein L15 OS=Bos taurus OX=9913 PE=3 SV=1 - [E1BK63_BOVIN] | 0,000E0 | 0,000E0 | 0,000E0 | 5,926E7 | 2,486E8 | 1,016E8 | RpL15 |
| F1MW57 | Serine/threonine-protein phosphatase OS=Bos taurus OX=9913 GN=PPP1CB PE=3 SV=2 - [F1MW57_BOVIN] | 0,000E0 | 0,000E0 | 0,000E0 | 2,519E7 | 6,897E7 | 1,588E8 | PPP1CB |
| Q2HJ81 | Tubulin beta-6 chain OS=Bos taurus OX=9913 GN=TUBB6 PE=2 SV=1 - [TBB6_BOVIN] | 4,519E7 | 0,000E0 | 0,000E0 | 3,323E7 | 1,818E8 | 4,385E7 | TUBB6 |
| E1BB71 | Uncharacterized protein OS=Bos taurus OX=9913 PE=3 SV=1 - [E1BB71_BOVIN] | 0,000E0 | 0,000E0 | 0,000E0 | 1,724E7 | 2,711E8 | 6,555E8 | E18871 |
| O18789 | 40S ribosomal protein S2 OS=Bos taurus OX=9913 GN=RPS2 PE=2 SV=2 - [RS2_BOVIN] | 0,000E0 | 0,000E0 | 0,000E0 | 5,188E7 | 1,240E7 | 3,647E7 | RPS2 |
| G3N0K3 | 40S ribosomal protein S4 OS=Bos taurus OX=9913 PE=3 SV=1 - [G3N0K3_BOVIN] | 0,000E0 | 0,000E0 | 0,000E0 | 8,644E7 | 1,553E7 | 2,180E8 | RPS4 |
| Q5E988 | 40S ribosomal protein S5 OS=Bos taurus OX=9913 GN=RPS5 PE=2 SV=3 - [RS5_BOVIN] | 0,000E0 | 0,000E0 | 0,000E0 | 1,026E8 | 1,667E8 | 2,072E8 | RPS5 |
| G8JKY0 | 40S ribosomal protein S8 OS=Bos taurus OX=9913 GN=RPS8 PE=3 SV=1 - [G8JKY0_BOVIN] | 0,000E0 | 0,000E0 | 0,000E0 | 8,684E7 | 5,272E8 | 8,540E8 | RPS8 |
| P42899 | 60S acidic ribosomal protein P2 OS=Bos taurus OX=9913 GN=RPLP2 PE=3 SV=1 - [RLA2_BOVIN] | 2,973E5 | 2,451E6 | 1,757E7 | 3,734E8 | 4,048E8 | 2,610E8 | RPLP2 |
| G3MXC8 | Actin-related protein 2/3 complex subunit 5 OS=Bos taurus OX=9913 GN=ARPC5 PE=3 SV=1 - [G3MXC8_BOVIN] | 0,000E0 | 0,000E0 | 0,000E0 | 3,334E7 | 2,791E8 | 5,231E8 | ARPC5 |
| E1BHK2 | Adducin 1 OS=Bos taurus OX=9913 GN=ADD1 PE=4 SV=1 - [E1BHK2_BOVIN] | 0,000E0 | 0,000E0 | 0,000E0 | 1,078E7 | 8,765E7 | 1,001E8 | ADD1 |
| F1MSZ6 | Antithrombin-III OS=Bos taurus OX=9913 GN=SERPINC1 PE=3 SV=1 - [F1MSZ6_BOVIN] | 0,000E0 | 0,000E0 | 0,000E0 | 1,089E8 | 3,783E7 | 2,297E7 | SERPINC1 |
| Q17QZ6 | Caveolae associated protein 2 (Fragment) OS=Bos taurus OX=9913 GN=CAVIN2 PE=1 SV=1 - [Q17QZ6_BOVIN] | 0,000E0 | 0,000E0 | 0,000E0 | 6,031E6 | 4,088E8 | 3,310E8 | CAVIN2 |
| Q148F1 | Cofilin-2 OS=Bos taurus OX=9913 GN=CFL2 PE=2 SV=1 - [COF2_BOVIN] | 2,300E7 | 0,000E0 | 0,000E0 | 4,595E8 | 1,853E9 | 1,916E8 | CFL2 |
| A5PKE2 | DnaJ homolog subfamily C member 10 OS=Bos taurus OX=9913 GN=DNAJC10 PE=2 SV=1 - [A5PKE2_BOVIN] | 0,000E0 | 5,678E6 | 4,686E6 | 3,724E7 | 2,923E7 | 1,292E7 | DNAJC10 |

| | | | | | | | | |
|---|---|---|---|---|---|---|---|---|
| Q3ZCE2 | ELAV-like protein OS=Bos taurus OX=9913 GN=ELAVL1 PE=2 SV=1 - [Q3ZCE2_BOVIN] | 0,000E0 | 0,000E0 | 0,000E0 | 2,994E7 | 8,460E6 | 2,082E6 | ELAVL1 |
| A7MB16 | Eukaryotic translation initiation factor 3 subunit B OS=Bos taurus OX=9913 GN=EIF3B PE=2 SV=1 - [EIF3B_BOVIN] | 0,000E0 | 0,000E0 | 0,000E0 | 4,213E7 | 6,440E7 | 9,518E7 | EIF3B |
| F1N0S9 | Family with sequence similarity 98 member A OS=Bos taurus OX=9913 GN=FAM98A PE=4 SV=2 - [F1N0S9_BOVIN] | 0,000E0 | 0,000E0 | 0,000E0 | 6,168E7 | 1,409E7 | 1,757E7 | FAM98A |
| Q08DC3 | KIAA1949 OS=Bos taurus OX=9913 GN=PPP1R18 PE=1 SV=1 - [Q08DC3_BOVIN] | 0,000E0 | 0,000E0 | 0,000E0 | 7,644E6 | 1,611E8 | 1,026E7 | PPP1R18 |
| Q3SZI2 | Lamin A/C OS=Bos taurus OX=9913 GN=LMNA PE=2 SV=1 - [Q3SZI2_BOVIN] | 0,000E0 | 0,000E0 | 0,000E0 | 1,192E7 | 3,000E7 | 9,390E6 | LMNA |
| A4IFN6 | Sorting nexin OS=Bos taurus OX=9913 GN=SNX18 PE=1 SV=1 - [A4IFN6_BOVIN] | 0,000E0 | 0,000E0 | 0,000E0 | 5,663E7 | 2,641E7 | 2,466E7 | SNX18 |
| Q28194 | Thrombospondin-1 (Fragment) OS=Bos taurus OX=9913 PE=2 SV=1 - [Q28194_BOVIN] | 9,682E5 | 0,000E0 | 0,000E0 | 1,277E7 | 1,020E8 | 7,533E7 | THBS1 |
| Q7M2Y2 | Tumor-specific transplantation antigen P198 homolog p23 (Fragment) OS=Bos taurus OX=9913 PE=3 SV=1 - [Q7M2Y2_BOVIN] | 0,000E0 | 0,000E0 | 0,000E0 | 5,632E7 | 2,310E8 | 2,325E8 | TSTAP198 |
| A0A140T832 | 40S ribosomal protein S17 OS=Bos taurus OX=9913 GN=RPS17 PE=3 SV=1 - [A0A140T832_BOVIN] | 0,000E0 | 0,000E0 | 0,000E0 | 1,104E8 | 2,705E8 | 3,804E8 | RPS17 |
| Q3T0U2 | 60S ribosomal protein L14 OS=Bos taurus OX=9913 GN=RPL14 PE=2 SV=3 - [RL14_BOVIN] | 0,000E0 | 4,570E6 | 0,000E0 | 1,233E7 | 8,023E6 | 1,448E8 | RPL14 |
| P61356 | 60S ribosomal protein L27 OS=Bos taurus OX=9913 GN=RPL27 PE=2 SV=2 - [RL27_BOVIN] | 0,000E0 | 0,000E0 | 0,000E0 | 9,133E7 | 7,446E8 | 7,456E8 | RPL27 |
| Q3T0D5 | 60S ribosomal protein L30 OS=Bos taurus OX=9913 GN=RPL30 PE=3 SV=3 - [RL30_BOVIN] | 0,000E0 | 0,000E0 | 0,000E0 | 7,082E7 | 1,011E8 | 2,929E7 | RPL30 |
| Q3T0S6 | 60S ribosomal protein L8 OS=Bos taurus OX=9913 GN=RPL8 PE=2 SV=3 - [RL8_BOVIN] | 0,000E0 | 0,000E0 | 0,000E0 | 1,140E8 | 4,068E8 | 5,003E8 | RPL8 |
| Q3SYR7 | 60S ribosomal protein L9 OS=Bos taurus OX=9913 GN=RPL9 PE=2 SV=1 - [RL9_BOVIN] | 0,000E0 | 0,000E0 | 0,000E0 | 1,010E8 | 3,361E8 | 6,339E8 | RPL9 |
| A5PKK3 | ABLIM3 protein OS=Bos taurus OX=9913 GN=ABLIM3 PE=2 SV=1 - [A5PKK3_BOVIN] | 0,000E0 | 0,000E0 | 0,000E0 | 5,829E6 | 4,105E7 | 1,068E7 | ABLIM3 |
| Q1JP79 | Actin-related protein 2/3 complex subunit 1A OS=Bos taurus OX=9913 GN=ARPC1A PE=2 SV=1 - [ARC1A_BOVIN] | 0,000E0 | 0,000E0 | 0,000E0 | 4,172E7 | 3,177E8 | 1,981E8 | ARPC1A |
| Q7SIH1 | Alpha-2-macroglobulin OS=Bos taurus OX=9913 GN=A2M PE=1 SV=2 - [A2MG_BOVIN] | 1,137E7 | 3,013E6 | 0,000E0 | 1,317E8 | 2,180E8 | 2,171E8 | A2M |
| E1BEQ5 | Angiomotin-like protein 1 OS=Bos taurus OX=9913 GN=AMOTL1 PE=3 SV=1 - [AMOL1_BOVIN] | 0,000E0 | 0,000E0 | 0,000E0 | 1,295E7 | 1,455E7 | 6,657E6 | AMOTL1 |
| E1BJ59 | Ataxin 2 like OS=Bos taurus OX=9913 GN=ATXN2L PE=4 SV=1 - [E1BJ59_BOVIN] | 0,000E0 | 0,000E0 | 0,000E0 | 1,875E7 | 1,501E7 | 9,101E6 | ATXN2L |
| Q3T0S7 | Chromosome 14 open reading frame 166 ortholog OS=Bos taurus OX=9913 GN=RTRAF PE=2 SV=1 - [Q3T0S7_BOVIN] | 0,000E0 | 0,000E0 | 0,000E0 | 1,074E7 | 6,778E7 | 6,546E7 | RTRAF |
| Q5E9F7 | Cofilin-1 OS=Bos taurus OX=9913 GN=CFL1 PE=2 SV=3 - [COF1_BOVIN] | 0,000E0 | 0,000E0 | 0,000E0 | 4,595E8 | 1,277E9 | 1,616E8 | CFL1 |
| Q3ZC35 | Cysteine rich angiogenic inducer 61 OS=Bos taurus OX=9913 GN=CYR61 PE=2 SV=1 - [Q3ZC35_BOVIN] | 0,000E0 | 0,000E0 | 0,000E0 | 6,670E7 | 1,494E8 | 4,259E7 | CYR61 |
| A0A140T862 | Eukaryotic translation initiation factor 2 subunit 2 OS=Bos taurus OX=9913 GN=EIF2S2 PE=4 SV=1 - [A0A140T862_BOVIN] | 0,000E0 | 0,000E0 | 0,000E0 | 1,468E7 | 2,911E7 | 3,245E7 | EIF2S2 |
| F1MQ56 | Focadhesin OS=Bos taurus OX=9913 GN=FOCAD PE=4 SV=2 - [F1MQ56_BOVIN] | 0,000E0 | 0,000E0 | 0,000E0 | 8,022E7 | 5,696E7 | 5,142E7 | FOCAD |
| F1N4M9 | Fragile X mental retardation syndrome-related protein 1 OS=Bos taurus OX=9913 GN=FXR1 PE=4 SV=1 - [F1N4M9_BOVIN] | 0,000E0 | 0,000E0 | 0,000E0 | 1,192E7 | 4,025E7 | 4,736E7 | FXR1 |
| A0A1B0Z542 | Heat shock 27 kDa protein 1 OS=Bos taurus OX=9913 GN=HspB1 PE=3 SV=1 - [A0A1B0Z542_BOVIN] | 0,000E0 | 0,000E0 | 0,000E0 | 8,268E7 | 8,538E7 | 1,155E8 | HspB1 |
| A5D9H5 | Heterogeneous nuclear ribonucleoprotein D OS=Bos taurus OX=9913 GN=HNRNPD PE=1 SV=1 - [A5D9H5_BOVIN] | 4,372E7 | 0,000E0 | 0,000E0 | 2,938E8 | 1,979E8 | 2,843E8 | HNRNPD |
| Q29RQ2 | KH RNA binding domain containing, signal transduction associated 1 OS=Bos taurus OX=9913 GN=KHDRBS1 PE=2 SV=1 - [Q29RQ2_BOVIN] | 3,071E7 | 0,000E0 | 0,000E0 | 1,917E7 | 4,182E6 | 2,765E7 | KHDRBS1 |
| A1L5B0 | Myosin regulatory light polypeptide 9 (Fragment) OS=Bos taurus OX=9913 GN=MYL9 PE=2 SV=1 - [A1L5B0_BOVIN] | 4,387E6 | 0,000E0 | 0,000E0 | 1,692E9 | 3,386E9 | 6,605E9 | MYL9 |
| F1MXK2 | NUFIP2, FMR1 interacting protein 2 OS=Bos taurus OX=9913 GN=NUFIP2 PE=4 SV=1 - [F1MXK2_BOVIN] | 0,000E0 | 0,000E0 | 0,000E0 | 1,574E7 | 4,698E7 | 4,306E7 | NUFIP2 |
| E1BIE7 | PGAM family member 5, mitochondrial serine/threonine protein phosphatase OS=Bos taurus OX=9913 GN=PGAM5 PE=4 SV=1 - [E1BIE7_BOVIN] | 0,000E0 | 0,000E0 | 0,000E0 | 5,324E7 | 1,114E8 | 1,166E7 | PGAM5 |
| P15246 | Protein-L-isoaspartate(D-aspartate) O-methyltransferase OS=Bos taurus OX=9913 GN=PCMT1 PE=1 SV=2 - [PIMT_BOVIN] | 0,000E0 | 0,000E0 | 0,000E0 | 1,224E7 | 4,604E7 | 2,820E7 | PCMT1 |
| Q0VCY7 | Serine/arginine-rich splicing factor 1 OS=Bos taurus OX=9913 GN=SRSF1 PE=2 SV=1 - [SRSF1_BOVIN] | 0,000E0 | 0,000E0 | 0,000E0 | 2,288E7 | 1,376E8 | 7,552E7 | SRSF1 |
| Q862N4 | Similar to ribosomal protein L26 (Fragment) OS=Bos taurus OX=9913 PE=2 SV=1 - [Q862N4_BOVIN] | 0,000E0 | 0,000E0 | 0,000E0 | 1,276E8 | 4,142E8 | 3,335E8 | RPL26 |
| F1MMU4 | Uncharacterized protein OS=Bos taurus OX=9913 GN=H1FX PE=1 SV=2 - [F1MMU4_BOVIN] | 0,000E0 | 0,000E0 | 0,000E0 | 4,781E7 | 2,250E8 | 1,146E8 | H1FX |
| F1MT25 | Uncharacterized protein OS=Bos taurus OX=9913 GN=HDLBP PE=1 SV=2 - [F1MT25_BOVIN] | 0,000E0 | 0,000E0 | 0,000E0 | 4,485E6 | 1,584E7 | 1,650E7 | HDLBP |
| G5E6K7 | Uncharacterized protein OS=Bos taurus OX=9913 PE=1 SV=1 - [G5E6K7_BOVIN] | 0,000E0 | 0,000E0 | 0,000E0 | 8,792E7 | 1,212E9 | 1,192E8 | G5E6K7 |
| E1BED8 | Uncharacterized protein OS=Bos taurus OX=9913 PE=4 SV=2 - [E1BED8_BOVIN] | 4,290E7 | 0,000E0 | 0,000E0 | 2,024E8 | 5,449E6 | 8,357E7 | E1BED8 |
| Q76I81 | 40S ribosomal protein S12 OS=Bos taurus OX=9913 GN=RPS12 PE=2 SV=1 - [RS12_BOVIN] | 0,000E0 | 0,000E0 | 0,000E0 | 1,684E8 | 6,996E7 | 2,046E7 | RPS12 |
| Q56JX8 | 40S ribosomal protein S13 OS=Bos taurus OX=9913 GN=RPS13 PE=2 SV=3 - [RS13_BOVIN] | 6,413E6 | 0,000E0 | 0,000E0 | 1,517E8 | 7,887E8 | 7,134E6 | RPS13 |
| Q3ZBH8 | 40S ribosomal protein S20 OS=Bos taurus OX=9913 GN=RPS20 PE=3 SV=1 - [RS20_BOVIN] | 0,000E0 | 0,000E0 | 0,000E0 | 2,328E8 | 3,548E8 | 4,791E8 | RPS20 |
| F1MKZ5 | 40S ribosomal protein S6 OS=Bos taurus OX=9913 PE=3 SV=1 - [F1MKZ5_BOVIN] | 0,000E0 | 0,000E0 | 0,000E0 | 8,869E7 | 5,191E8 | 6,972E8 | RPS6 |
| Q56JZ1 | 60S ribosomal protein L13 OS=Bos taurus OX=9913 GN=RPL13 PE=2 SV=3 - [RL13_BOVIN] | 0,000E0 | 0,000E0 | 0,000E0 | 5,088E7 | 3,066E8 | 1,328E8 | RPL13 |
| Q3T0L7 | 60S ribosomal protein L28 OS=Bos taurus OX=9913 GN=RPL28 PE=2 SV=3 - [RL28_BOVIN] | 0,000E0 | 0,000E0 | 0,000E0 | 3,291E7 | 2,485E8 | 2,020E8 | RPL28 |
| E1BMD5 | Apoptosis and caspase activation inhibitor OS=Bos taurus OX=9913 GN=AVEN PE=4 SV=1 - [E1BMD5_BOVIN] | 0,000E0 | 0,000E0 | 0,000E0 | 5,971E7 | 4,600E7 | 3,175E7 | AVEN |
| G3X6H0 | Centrosomal protein 170B OS=Bos taurus OX=9913 GN=CEP170B PE=4 SV=1 - [G3X6H0_BOVIN] | 0,000E0 | 0,000E0 | 0,000E0 | 1,348E7 | 7,762E6 | 6,083E6 | CEP170B |
| F1MBG0 | Coactivator associated arginine methyltransferase 1 OS=Bos taurus OX=9913 GN=CARM1 PE=3 SV=2 - [F1MBG0_BOVIN] | 0,000E0 | 0,000E0 | 0,000E0 | 8,712E7 | 1,873E7 | 1,653E7 | CARM1 |
| Q3T0B6 | Complement component 1 Q subcomponent-binding protein, mitochondrial OS=Bos taurus OX=9913 GN=C1QBP PE=2 SV=1 - [C1QBP_BOVIN] | 0,000E0 | 0,000E0 | 0,000E0 | 5,750E7 | 1,105E8 | 1,085E8 | C1QBP |
| A5D989 | Elongation factor 1-delta OS=Bos taurus OX=9913 GN=EEF1D PE=2 SV=2 - [EF1D_BOVIN] | 0,000E0 | 0,000E0 | 0,000E0 | 2,575E7 | 1,167E8 | 8,650E7 | EEF1D |
| Q1JPA2 | Eukaryotic translation elongation factor 1 gamma (Fragment) OS=Bos taurus OX=9913 GN=EEF1G PE=2 SV=1 - [Q1JPA2_BOVIN] | 0,000E0 | 0,000E0 | 0,000E0 | 1,101E7 | 3,934E7 | 3,985E7 | EEF1G |
| Q3ZCK1 | Eukaryotic translation initiation factor 3 subunit L OS=Bos taurus OX=9913 GN=EIF3L PE=2 SV=1 - [EIF3L_BOVIN] | 0,000E0 | 0,000E0 | 0,000E0 | 3,067E7 | 9,184E7 | 3,631E7 | EIF3L |
| F1MX04 | Eukaryotic translation initiation factor 4 gamma 1 OS=Bos taurus OX=9913 GN=EIF4G1 PE=1 SV=2 - [F1MX04_BOVIN] | 0,000E0 | 0,000E0 | 0,000E0 | 1,791E6 | 9,165E6 | 1,696E7 | EIF4G1 |
| F1MMA0 | Interleukin enhancer binding factor 3 OS=Bos taurus OX=9913 GN=ILF3 PE=4 SV=2 - [F1MMA0_BOVIN] | 0,000E0 | 0,000E0 | 0,000E0 | 9,194E6 | 2,612E7 | 3,575E7 | ILF3 |
| P13909 | Plasminogen activator inhibitor 1 OS=Bos taurus OX=9913 GN=SERPINE1 PE=1 SV=1 - [PAI1_BOVIN] | 0,000E0 | 0,000E0 | 0,000E0 | 1,088E7 | 1,376E7 | 1,483E6 | SERPINE1 |
| A7E3Q8 | Plastin-3 OS=Bos taurus OX=9913 GN=PLS3 PE=2 SV=1 - [PLST_BOVIN] | 0,000E0 | 0,000E0 | 0,000E0 | 1,804E7 | 2,058E6 | 5,515E7 | PLS3 |
| Q8WN55 | Polypyrimidine tract-binding protein 1 OS=Bos taurus OX=9913 GN=PTBP1 PE=2 SV=1 - [PTBP1_BOVIN] | 0,000E0 | 0,000E0 | 0,000E0 | 3,831E7 | 8,054E7 | 3,395E7 | PTBP1 |
| Q58D08 | Pyrroline-5-carboxylate reductase 3 OS=Bos taurus OX=9913 GN=PYCR3 PE=2 SV=1 - [P5CR3_BOVIN] | 0,000E0 | 0,000E0 | 0,000E0 | 1,569E7 | 1,484E7 | 1,339E7 | PYCR3 |
| Q862B7 | Similar to ribosomal protein S14 (Fragment) OS=Bos taurus OX=9913 PE=2 SV=1 - [Q862B7_BOVIN] | 4,964E6 | 0,000E0 | 1,138E7 | 1,135E8 | 8,427E7 | 5,332E7 | RPL14 |
| A6QNY4 | STAU1 protein (Fragment) OS=Bos taurus OX=9913 GN=STAU1 PE=2 SV=1 - [A6QNY4_BOVIN] | 0,000E0 | 0,000E0 | 0,000E0 | 1,558E7 | 1,091E7 | 1,820E7 | STAU1 |
| Q1RMR2 | U1 small nuclear ribonucleoprotein 70 kDa OS=Bos taurus OX=9913 GN=SNRNP70 PE=2 SV=1 - [RU17_BOVIN] | 0,000E0 | 0,000E0 | 0,000E0 | 5,682E7 | 1,176E7 | 6,602E7 | SNRNP70 |
| G3MWV5 | Uncharacterized protein OS=Bos taurus OX=9913 GN=HIST1H1E PE=3 SV=1 - [G3MWV5_BOVIN] | 0,000E0 | 0,000E0 | 0,000E0 | 6,812E6 | 9,031E7 | 5,153E7 | HIST1H1E |
| P62261 | 14-3-3 protein epsilon OS=Bos taurus OX=9913 GN=YWHAE PE=2 SV=1 - [1433E_BOVIN] | 0,000E0 | 0,000E0 | 0,000E0 | 4,173E7 | 4,978E7 | 3,479E7 | YWHAE |
| Q32PB8 | 40S ribosomal protein S21 OS=Bos taurus OX=9913 GN=RPS21 PE=3 SV=1 - [RS21_BOVIN] | 0,000E0 | 0,000E0 | 0,000E0 | 2,467E8 | 1,086E8 | 4,541E8 | RPS21 |
| Q2KHT7 | 40S ribosomal protein S27 OS=Bos taurus OX=9913 GN=RPS27 PE=3 SV=3 - [RS27_BOVIN] | 0,000E0 | 0,000E0 | 0,000E0 | 2,349E8 | 6,740E8 | 6,140E8 | RPS27 |
| Q3T087 | 60S ribosomal protein L11 OS=Bos taurus OX=9913 GN=RPL11 PE=2 SV=3 - [RL11_BOVIN] | 1,017E7 | 0,000E0 | 0,000E0 | 4,577E8 | 1,108E9 | 9,654E8 | RPL11 |
| Q3T171 | 60S ribosomal protein L36 OS=Bos taurus OX=9913 GN=RPL36 PE=3 SV=3 - [RL36_BOVIN] | 0,000E0 | 0,000E0 | 0,000E0 | 7,080E7 | 2,732E8 | 1,168E8 | RPL36 |
| F1MXX2 | Argonaute 1, RISC catalytic component OS=Bos taurus OX=9913 GN=AGO1 PE=3 SV=1 - [F1MXX2_BOVIN] | 0,000E0 | 0,000E0 | 0,000E0 | 3,785E7 | 1,025E8 | 4,834E7 | AGO1 |
| F1MPE1 | CD109 molecule OS=Bos taurus OX=9913 GN=CD109 PE=4 SV=2 - [F1MPE1_BOVIN] | 3,116E6 | 0,000E0 | 0,000E0 | 6,490E6 | 2,454E7 | 2,525E7 | CD109 |
| G3X6T9 | Flotillin-2 OS=Bos taurus OX=9913 GN=FLOT2 PE=4 SV=1 - [G3X6T9_BOVIN] | 0,000E0 | 0,000E0 | 0,000E0 | 3,018E6 | 2,113E7 | 3,457E7 | FLOT2 |
| A7MBH9 | G protein subunit alpha i2 OS=Bos taurus OX=9913 GN=GNAI2 PE=2 SV=1 - [A7MBH9_BOVIN] | 0,000E0 | 0,000E0 | 0,000E0 | 8,082E6 | 6,638E7 | 6,357E7 | GNAI2 |
| P18246 | Gap junction alpha-1 protein OS=Bos taurus OX=9913 GN=GJA1 PE=2 SV=2 - [CXA1_BOVIN] | 0,000E0 | 0,000E0 | 0,000E0 | 5,784E7 | 9,526E7 | 3,076E8 | GJA1 |
| Q27990 | Nonmuscle myosin heavy chain B (Fragment) OS=Bos taurus OX=9913 PE=2 SV=1 - [Q27990_BOVIN] | 0,000E0 | 0,000E0 | 0,000E0 | 1,415E7 | 1,667E8 | 2,043E8 | MYHCB |
| Q3ZBH5 | Proliferation-associated 2G4, 38kDa OS=Bos taurus OX=9913 GN=PA2G4 PE=1 SV=1 - [Q3ZBH5_BOVIN] | 0,000E0 | 0,000E0 | 0,000E0 | 3,003E7 | 8,378E7 | 1,121E8 | PA2G4 |
| F1MSA5 | Protein LSM14 homolog A OS=Bos taurus OX=9913 GN=LSM14A PE=4 SV=1 - [F1MSA5_BOVIN] | 0,000E0 | 0,000E0 | 0,000E0 | 1,185E7 | 1,164E7 | 1,508E7 | LSM14A |
| Q56JV6 | Twinfilin-1 OS=Bos taurus OX=9913 GN=TWF1 PE=2 SV=1 - [TWF1_BOVIN] | 0,000E0 | 0,000E0 | 0,000E0 | 8,981E6 | 5,799E7 | 6,810E7 | TWF1 |
| A5PJS6 | Ubiquitin carboxyl-terminal hydrolase 10 OS=Bos taurus OX=9913 GN=USP10 PE=2 SV=1 - [UBP10_BOVIN] | 0,000E0 | 0,000E0 | 0,000E0 | 4,721E6 | 2,971E7 | 1,435E7 | USP10 |
| G3X7D5 | Uncharacterized protein OS=Bos taurus OX=9913 GN=LOC787803 PE=3 SV=1 - [G3X7D5_BOVIN] | 0,000E0 | 0,000E0 | 0,000E0 | 8,109E7 | 1,198E8 | 3,586E7 | LOC787803 |
| G8JKX9 | Uncharacterized protein OS=Bos taurus OX=9913 PE=4 SV=1 - [G8JKX9_BOVIN] | 0,000E0 | 0,000E0 | 0,000E0 | 1,582E8 | 7,630E8 | 3,930E8 | G8JKX9 |
| E1BH80 | YTH N6-methyladenosine RNA binding protein 3 OS=Bos taurus OX=9913 GN=YTHDF3 PE=4 SV=2 - [E1BH80_BOVIN] | 0,000E0 | 0,000E0 | 7,660E6 | 2,072E7 | 3,117E6 | 5,268E6 | YTHDF3 |
| P63103 | 14-3-3 protein zeta/delta OS=Bos taurus OX=9913 GN=YWHAZ PE=1 SV=1 - [1433Z_BOVIN] | 0,000E0 | 0,000E0 | 0,000E0 | 6,211E7 | 5,559E7 | 2,238E7 | YWHAZ |
| G3N3U9 | 28S ribosomal protein S22, mitochondrial OS=Bos taurus OX=9913 GN=MRPS22 PE=4 SV=1 - [G3N3U9_BOVIN] | 0,000E0 | 0,000E0 | 0,000E0 | 1,821E6 | 1,326E7 | 2,570E7 | MRPS22 |
| Q3T0F4 | 40S ribosomal protein S10 OS=Bos taurus OX=9913 GN=RPS10 PE=2 SV=1 - [RS10_BOVIN] | 0,000E0 | 0,000E0 | 0,000E0 | 1,056E8 | 6,029E8 | 3,668E7 | RPS10 |
| Q3T0B7 | 40S ribosomal protein S27-like OS=Bos taurus OX=9913 GN=RPS27L PE=3 SV=3 - [RS27L_BOVIN] | 0,000E0 | 0,000E0 | 0,000E0 | 1,669E8 | 1,329E9 | 1,157E9 | RPS27L |
| Q3T035 | Actin-related protein 2/3 complex subunit 3 OS=Bos taurus OX=9913 GN=ARPC3 PE=1 SV=3 - [ARPC3_BOVIN] | 0,000E0 | 0,000E0 | 0,000E0 | 1,183E7 | 8,006E7 | 6,936E7 | ARPC3 |
| Q66WT7 | Caveolin-2 OS=Bos taurus OX=9913 GN=CAV2 PE=1 SV=1 - [CAV2_BOVIN] | 0,000E0 | 0,000E0 | 0,000E0 | 4,629E7 | 1,038E9 | 9,367E8 | CAV2 |
| Q5E9D5 | Destrin OS=Bos taurus OX=9913 GN=DSTN PE=2 SV=3 - [DEST_BOVIN] | 0,000E0 | 0,000E0 | 0,000E0 | 2,184E7 | 1,024E8 | 8,720E7 | DSTN |
| Q3T0V7 | Endothelial differentiation-related factor 1 OS=Bos taurus OX=9913 GN=EDF1 PE=2 SV=1 - [EDF1_BOVIN] | 0,000E0 | 0,000E0 | 0,000E0 | 1,233E8 | 7,445E6 | 5,071E8 | EDF1 |
| G8JKV2 | Eukaryotic translation initiation factor 3 subunit J OS=Bos taurus OX=9913 GN=EIF3J PE=3 SV=1 - [G8JKV2_BOVIN] | 0,000E0 | 0,000E0 | 0,000E0 | 9,036E6 | 4,120E7 | 5,003E7 | EIF3J |
| Q9N0T5 | Eukaryotic translation initiation factor 4E OS=Bos taurus OX=9913 GN=EIF4E PE=2 SV=2 - [IF4E_BOVIN] | 0,000E0 | 0,000E0 | 0,000E0 | 1,556E7 | 3,587E7 | 1,385E7 | EIF4E |
| F1N0F7 | Eukaryotic translation initiation factor 5 OS=Bos taurus OX=9913 GN=EIF5 PE=4 SV=1 - [F1N0F7_BOVIN] | 0,000E0 | 0,000E0 | 0,000E0 | 6,253E6 | 2,284E7 | 2,986E7 | EIF5 |
| G3X807 | Histone H4 OS=Bos taurus OX=9913 PE=3 SV=1 - [G3X807_BOVIN] | 8,849E6 | 0,000E0 | 0,000E0 | 7,682E7 | 2,274E8 | 1,096E8 | HIST1H4 |
| Q0V8D5 | Interleukin enhancer binding factor 3 isoform c (Fragment) OS=Bos taurus OX=9913 GN=ILF3 PE=2 SV=1 - [Q0V8D5_BOVIN] | 0,000E0 | 0,000E0 | 0,000E0 | 2,430E7 | 1,210E7 | 5,093E7 | ILF3 |
| A6QP32 | KRT85 protein OS=Bos taurus OX=9913 GN=KRT89 PE=2 SV=1 - [A6QP32_BOVIN] | 1,122E7 | 0,000E0 | 0,000E0 | 9,868E7 | 3,859E8 | 2,622E8 | KRT89 |
| Q5E9A1 | Nascent polypeptide-associated complex subunit alpha OS=Bos taurus OX=9913 GN=NACA PE=1 SV=1 - [NACA_BOVIN] | 0,000E0 | 0,000E0 | 0,000E0 | 2,870E7 | 3,322E7 | 3,219E7 | NACA |
| Q5E947 | Peroxiredoxin-1 OS=Bos taurus OX=9913 GN=PRDX1 PE=2 SV=1 - [PRDX1_BOVIN] | 0,000E0 | 0,000E0 | 0,000E0 | 5,179E7 | 3,470E7 | 3,403E7 | PRDX1 |
| E1B9K1 | Polyubiquitin-C OS=Bos taurus OX=9913 GN=UBC PE=4 SV=2 - [E1B9K1_BOVIN] | 2,804E7 | 0,000E0 | 0,000E0 | 1,080E9 | 4,948E9 | 4,621E9 | UBC |
| F1MQQ8 | PPFIA binding protein 1 OS=Bos taurus OX=9913 GN=PPFIBP1 PE=4 SV=1 - [F1MQQ8_BOVIN] | 0,000E0 | 0,000E0 | 0,000E0 | 3,977E6 | 2,054E7 | 3,003E7 | PPFIBP1 |
| Q58DM9 | Ribosomal protein L27a OS=Bos taurus OX=9913 GN=RPL27A PE=2 SV=1 - [Q58DM9_BOVIN] | 0,000E0 | 0,000E0 | 0,000E0 | 1,249E8 | 4,540E8 | 3,222E7 | RPL27A |
| Q3ZBP3 | RNA-binding motif, single-stranded-interacting protein 1 OS=Bos taurus OX=9913 GN=RBMS1 PE=2 SV=1 - [RBMS1_BOVIN] | 0,000E0 | 0,000E0 | 0,000E0 | 6,466E7 | 6,052E7 | 3,164E7 | RBMS1 |
| F1MSD2 | RuvB-like helicase OS=Bos taurus OX=9913 GN=RUVBL2 PE=3 SV=1 - [F1MSD2_BOVIN] | 0,000E0 | 0,000E0 | 0,000E0 | 6,548E6 | 8,940E6 | 8,113E6 | RUVBL2 |

| | | | | | | | | |
|---|---|---|---|---|---|---|---|---|
| Q3SZR8 | Serine/arginine-rich splicing factor 3 OS=Bos taurus OX=9913 GN=SRSF3 PE=2 SV=1 - [SRSF3_BOVIN] | 0,000E0 | 0,000E0 | 0,000E0 | 1,704E7 | 4,621E7 | 4,985E6 | SRSF3 |
| Q862R3 | Similar to ribosomal protein L18a (Fragment) OS=Bos taurus OX=9913 PE=2 SV=1 - [Q862R3_BOVIN] | 0,000E0 | 0,000E0 | 0,000E0 | 3,525E6 | 1,486E8 | 5,021E7 | RPL18 |
| F1MZ00 | Small nuclear ribonucleoprotein Sm D3 OS=Bos taurus OX=9913 GN=SNRPD3 PE=3 SV=2 - [F1MZ00_BOVIN] | 0,000E0 | 0,000E0 | 0,000E0 | 5,805E6 | 9,885E7 | 5,016E7 | SNRPD3 |
| E1BQ37 | Splicing factor proline and glutamine rich OS=Bos taurus OX=9913 GN=SFPQ PE=1 SV=1 - [E1BQ37_BOVIN] | 0,000E0 | 0,000E0 | 0,000E0 | 1,487E6 | 1,750E6 | 2,444E6 | SFPQ |
| A6H7J7 | Uncharacterized protein OS=Bos taurus OX=9913 PE=2 SV=1 - [A6H7J7_BOVIN] | 1,894E8 | 0,000E0 | 0,000E0 | 2,272E8 | 8,787E7 | 1,517E8 | A6H7J7 |
| Q56K10 | 40S ribosomal protein S15 OS=Bos taurus OX=9913 GN=RPS15 PE=2 SV=3 - [RS15_BOVIN] | 0,000E0 | 0,000E0 | 0,000E0 | 1,377E8 | 5,099E8 | 3,286E8 | RPS15 |
| Q3SZQ6 | 60S ribosomal protein L32 OS=Bos taurus OX=9913 GN=RPL32 PE=2 SV=3 - [RL32_BOVIN] | 0,000E0 | 0,000E0 | 0,000E0 | 5,706E7 | 9,771E7 | 6,501E7 | RPL32 |
| G3N2R1 | 60S ribosomal protein L6 OS=Bos taurus OX=9913 PE=3 SV=1 - [G3N2R1_BOVIN] | 0,000E0 | 0,000E0 | 0,000E0 | 1,007E8 | 4,719E7 | 4,830E6 | RPL6 |
| F1MLB8 | ATP synthase subunit alpha OS=Bos taurus OX=9913 GN=ATP5F1A PE=1 SV=1 - [F1MLB8_BOVIN] | 0,000E0 | 0,000E0 | 0,000E0 | 6,084E6 | 1,694E7 | 6,376E6 | ATP5F1A |
| Q32L92 | Calponin-3 OS=Bos taurus OX=9913 GN=CNN3 PE=2 SV=1 - [CNN3_BOVIN] | 0,000E0 | 0,000E0 | 0,000E0 | 6,621E6 | 1,696E6 | 2,887E6 | CNN3 |
| Q3ZBH3 | CD151 antigen OS=Bos taurus OX=9913 GN=CD151 PE=2 SV=1 - [CD151_BOVIN] | 1,829E7 | 0,000E0 | 0,000E0 | 2,437E8 | 1,726E8 | 1,498E8 | CD151 |
| Q2KII4 | Elongin-C OS=Bos taurus OX=9913 GN=ELOC PE=3 SV=1 - [ELOC_BOVIN] | 1,731E6 | 0,000E0 | 0,000E0 | 2,217E7 | 4,299E7 | 5,625E7 | ELOC |
| E1BLZ8 | Eukaryotic translation initiation factor 3 subunit F OS=Bos taurus OX=9913 GN=EIF3F PE=1 SV=2 - [E1BLZ8_BOVIN] | 0,000E0 | 0,000E0 | 0,000E0 | 3,096E7 | 1,714E6 | 4,833E7 | EIF3F |
| A5PK49 | IGL@ protein OS=Bos taurus OX=9913 GN=IGL@ PE=2 SV=1 - [A5PK49_BOVIN] | 5,852E8 | 1,489E7 | 6,150E6 | 1,867E9 | 1,979E9 | 1,824E9 | RBM8A |
| Q3ZCE8-2 | Isoform 2 of RNA-binding protein 8A OS=Bos taurus OX=9913 GN=RBM8A - [RBM8A_BOVIN] | 0,000E0 | 0,000E0 | 0,000E0 | 8,167E6 | 4,351E6 | 8,486E6 | RBM8A |
| P00974 | Pancreatic trypsin inhibitor OS=Bos taurus OX=9913 PE=1 SV=2 - [BPT1_BOVIN] | 0,000E0 | 0,000E0 | 3,765E7 | 1,766E9 | 7,076E8 | 2,062E8 | BPT1 |
| A5D9D7 | Procollagen-lysine, 2-oxoglutarate 5-dioxygenase 3 (Fragment) OS=Bos taurus OX=9913 GN=PLOD3 PE=2 SV=1 - [A5D9D7_BOVIN] | 0,000E0 | 0,000E0 | 0,000E0 | 4,242E6 | 5,760E6 | 9,798E6 | PLOD3 |
| Q0VCF9 | Protein LSM12 homolog OS=Bos taurus OX=9913 GN=LSM12 PE=2 SV=2 - [LSM12_BOVIN] | 0,000E0 | 0,000E0 | 0,000E0 | 2,548E7 | 4,155E7 | 5,952E7 | LSM12 |
| F1N301 | Uncharacterized protein OS=Bos taurus OX=9913 GN=RPL22 PE=1 SV=2 - [F1N301_BOVIN] | 0,000E0 | 0,000E0 | 0,000E0 | 1,154E8 | 2,741E8 | 6,289E8 | RPL22 |
| E1BHM9 | Uncharacterized protein OS=Bos taurus OX=9913 PE=3 SV=1 - [E1BHM9_BOVIN] | 0,000E0 | 0,000E0 | 0,000E0 | 5,423E6 | 1,843E8 | 7,843E7 | E18HM9 |
| Q2KI49 | 39S ribosomal protein L50, mitochondrial OS=Bos taurus OX=9913 GN=MRPL50 PE=1 SV=1 - [RM50_BOVIN] | 0,000E0 | 0,000E0 | 0,000E0 | 1,004E7 | 6,539E6 | 5,574E6 | MRPL50 |
| Q58DW3 | 60S ribosomal protein L29 OS=Bos taurus OX=9913 GN=RPL29 PE=2 SV=3 - [RL29_BOVIN] | 0,000E0 | 0,000E0 | 0,000E0 | 6,262E7 | 1,076E8 | 8,776E7 | RPL29 |
| P04973-2 | Isoform Non-brain of Clathrin light chain A OS=Bos taurus OX=9913 GN=CLTA - [CLCA_BOVIN] | 0,000E0 | 0,000E0 | 0,000E0 | 1,010E7 | 1,430E8 | 1,471E8 | CLTA |
| F1MKI2 | RAB12, member RAS oncogene family OS=Bos taurus OX=9913 GN=RAB12 PE=4 SV=2 - [F1MKI2_BOVIN] | 0,000E0 | 0,000E0 | 0,000E0 | 3,911E7 | 2,651E7 | 1,653E7 | RAB12 |
| Q0VCQ9 | Reticulocalbin 2, EF-hand calcium binding domain OS=Bos taurus OX=9913 GN=RCN2 PE=2 SV=1 - [Q0VCQ9_BOVIN] | 0,000E0 | 0,000E0 | 0,000E0 | 3,154E6 | 1,860E7 | 2,527E6 | RCN2 |
| Q32PA0 | U1 small nuclear ribonucleoprotein C OS=Bos taurus OX=9913 GN=SNRPC PE=2 SV=2 - [RU1C_BOVIN] | 0,000E0 | 0,000E0 | 0,000E0 | 8,697E7 | 1,857E7 | 4,872E6 | SNRPC |
| F1MI18 | Uncharacterized protein OS=Bos taurus OX=9913 PE=4 SV=2 - [F1MI18_BOVIN] | 0,000E0 | 0,000E0 | 0,000E0 | 1,437E8 | 2,459E8 | 3,093E8 | F1MI18 |

**Supplementary Table 3. Differentially expressed genes analyzed (AmotL2 KO vs WT aortae) with enriched GO**

| Gene_ID | readcount_amotl2 K0 aorta | readcount_wt_aorta | log2FoldChange | pvalue | padj | Gene_Name |
|---|---|---|---|---|---|---|
| ENSMUSG00000000290 | 158.927701543679 | 59.7030557544915 | 0.64533 | 0.00020619 | 0.047359 | Itgb2 |
| ENSMUSG00000002111 | 216.961569643465 | 100.502648381335 | 0.6889 | 4.0862e-05 | 0.020194 | Spi1 |
| ENSMUSG00000003032 | 1053.73474574084 | 664.16622403341 | 0.53107 | 0.00014112 | 0.040633 | Klf4 |
| ENSMUSG00000005338 | 173.756385209253 | 82.7869817301975 | 0.67988 | 4.3562e-05 | 0.020478 | Cadm3 |
| ENSMUSG00000010064 | 32.5248378550642 | 69.2139660522236 | -0.651 | 0.0001397 | 0.040633 | Slc38a3 |
| ENSMUSG00000010797 | 111.874554274348 | 15.1184535312192 | 0.70298 | 1.2295e-05 | 0.0091597 | Wnt2 |
| ENSMUSG00000012705 | 239.844667429362 | 52.6885310870578 | 0.79769 | 3.1281e-06 | 0.0065624 | Retn |
| ENSMUSG00000015340 | 143.040000460753 | 57.6902339080253 | 0.65278 | 0.00017391 | 0.045604 | Cybb |
| ENSMUSG00000015396 | 78.5125901989447 | 29.0719859870467 | 0.70756 | 4.74e-05 | 0.020478 | Cd83 |
| ENSMUSG00000018339 | 2485.71158356384 | 1233.54481642737 | 0.74646 | 9.1359e-07 | 0.003354 | Gpx3 |
| ENSMUSG00000018927 | 707.295207631469 | 170.544551968579 | 0.93728 | 7.1612e-08 | 0.00035054 | Ccl6 |
| ENSMUSG00000019122 | 476.329244367314 | 119.344035032349 | 1.0472 | 1.4243e-09 | 1.0458e-05 | Ccl9 |
| ENSMUSG00000020099 | 345.887697572691 | 496.115281245493 | -0.44752 | 0.00019136 | 0.046835 | Unc5b |
| ENSMUSG00000020437 | 77.1633165981257 | 22.0118072197688 | 0.77686 | 7.6345e-06 | 0.0091597 | Myo1g |
| ENSMUSG00000020607 | 538.408993623667 | 750.953249029995 | -0.4257 | 0.00011525 | 0.037515 | Fam84a |
| ENSMUSG00000021388 | 1406.02622944515 | 673.183669052565 | 0.71181 | 1.2079e-05 | 0.0091597 | Aspn |
| ENSMUSG00000021457 | 213.973625481079 | 106.330862475272 | 0.66452 | 4.9727e-05 | 0.020478 | Syk |
| ENSMUSG00000022102 | 145.695459403768 | 51.5169124495549 | 0.78403 | 6.0794e-06 | 0.0091597 | Dok2 |
| ENSMUSG00000022488 | 205.41780877786 | 98.2592233833908 | 0.64108 | 0.00015277 | 0.042328 | Nckap1l |
| ENSMUSG00000022901 | 66.4110773964363 | 29.7301164689561 | 0.68252 | 5.8594e-05 | 0.022063 | Cd86 |
| ENSMUSG00000023046 | 2884.24901370756 | 1480.29003107507 | 0.61981 | 0.00018518 | 0.046835 | Igfbp6 |
| ENSMUSG00000024679 | 76.0534731344345 | 33.0719080199057 | 0.66918 | 0.00010042 | 0.034294 | Ms4a6d |
| ENSMUSG00000024965 | 165.823698438286 | 90.238439882096 | 0.65277 | 1.4921e-05 | 0.010434 | Fermt3 |
| ENSMUSG00000026109 | 428.493277023279 | 244.168057152686 | 0.61223 | 3.7101e-05 | 0.020179 | Tmeff2 |
| ENSMUSG00000026177 | 252.967326692869 | 135.598140854474 | 0.6467 | 3.0292e-05 | 0.017794 | Slc11a1 |
| ENSMUSG00000026395 | 120.375597657209 | 44.8203082114949 | 0.71883 | 3.5148e-05 | 0.019852 | Ptprc |
| ENSMUSG00000026399 | 297.856093868171 | 113.297994385314 | 0.98022 | 4.6362e-10 | 6.8082e-06 | Cd55 |
| ENSMUSG00000028581 | 465.732257990103 | 238.530901942934 | 0.65498 | 4.9404e-05 | 0.020478 | Laptm5 |
| ENSMUSG00000029771 | 112.314140769318 | 54.4093371094395 | 0.71349 | 9.359e-06 | 0.0091597 | Irf5 |
| ENSMUSG00000030787 | 542.979717509508 | 221.284214010384 | 0.74918 | 1.1737e-05 | 0.0091597 | Lyve1 |
| ENSMUSG00000031827 | 420.977724023766 | 199.139679950657 | 0.70633 | 1.7501e-05 | 0.011682 | Cotl1 |
| ENSMUSG00000031906 | 164.547241376456 | 84.2907934668227 | 0.62867 | 0.00013349 | 0.040042 | Smpd3 |
| ENSMUSG00000032332 | 349.146794512607 | 192.498035360874 | 0.6278 | 4.1255e-05 | 0.020194 | Col12a1 |
| ENSMUSG00000035095 | 44.906359424677 | 4.66152356471479 | 0.63442 | 3.904e-05 | 0.020194 | Fam167a |
| ENSMUSG00000035208 | 84.8499252503698 | 29.273182687006 | 0.78264 | 6.5017e-06 | 0.0091597 | Slfn8 |
| ENSMUSG00000036887 | 1888.43045766811 | 1033.03008726011 | 0.59504 | 0.00021974 | 0.049644 | C1qa |
| ENSMUSG00000036905 | 1436.86135647427 | 723.356029904941 | 0.62421 | 0.00018833 | 0.046835 | C1qb |
| ENSMUSG00000037379 | 88.0293142776631 | 9.28999489950067 | 0.7633 | 2.2044e-06 | 0.0064744 | Spon2 |
| ENSMUSG00000038147 | 68.1694590534354 | 23.4935738710137 | 0.65282 | 0.00016487 | 0.04402 | Cd84 |
| ENSMUSG00000038642 | 464.667808327248 | 211.512395936203 | 0.686 | 5.0664e-05 | 0.020478 | Ctss |
| ENSMUSG00000039109 | 1503.97106070614 | 747.972796177522 | 0.6244 | 0.00020402 | 0.047359 | F13a1 |
| ENSMUSG00000039476 | 249.429624873089 | 141.757473194012 | 0.63225 | 1.2475e-05 | 0.0091597 | Prrx2 |
| ENSMUSG00000040552 | 132.466267916088 | 65.2414056829882 | 0.62538 | 0.0002064 | 0.047359 | C3ar1 |
| ENSMUSG00000040612 | 229.441118259121 | 88.319984284168 | 0.69464 | 6.4261e-05 | 0.023592 | Ildr2 |
| ENSMUSG00000040950 | 461.518760323747 | 205.384453243512 | 0.6718 | 8.7349e-05 | 0.030541 | Mgl2 |
| ENSMUSG00000044337 | 1335.306800426 | 691.095361709203 | 0.68272 | 1.154e-05 | 0.0091597 | Ackr3 |
| ENSMUSG00000045573 | 284.435378949316 | 164.418066334223 | 0.57468 | 0.00020262 | 0.047359 | Penk |
| ENSMUSG00000046743 | 1173.27680614689 | 1606.07262123916 | -0.41348 | 2.4734e-05 | 0.015134 | Fat4 |
| ENSMUSG00000049130 | 193.925359200559 | 93.3120301481718 | 0.64854 | 0.00011751 | 0.037515 | C5ar1 |
| ENSMUSG00000052384 | 161.609513064848 | 75.5828244406309 | 0.67969 | 5.1576e-05 | 0.020478 | Nrros |
| ENSMUSG00000052688 | 311.450918170674 | 175.345615022865 | 0.59411 | 0.00013361 | 0.040042 | Rab7b |
| ENSMUSG00000054203 | 58.3391527588879 | 21.2200107598094 | 0.65648 | 0.00015872 | 0.043162 | Ifi205 |
| ENSMUSG00000056069 | 52.0027327519442 | 21.5993828748247 | 0.69577 | 5.2991e-05 | 0.020478 | Fam105a |
| ENSMUSG00000056938 | 1667.9772179887 | 2239.70157240921 | -0.3859 | 0.00012605 | 0.039383 | Acbd4 |
| ENSMUSG00000059412 | 121.602644876039 | 36.8721972818891 | 0.69779 | 5.2982e-05 | 0.020478 | Fxyd2 |
| ENSMUSG00000059498 | 404.525738192865 | 205.145200130876 | 0.62295 | 0.00018416 | 0.046835 | Fcgr3 |
| ENSMUSG00000060586 | 1199.0007722268 | 444.640925833898 | 0.65852 | 0.00015262 | 0.042328 | H2-Eb1 |
| ENSMUSG00000062593 | 78.9260524892479 | 28.3739988423478 | 0.75867 | 1.2294e-05 | 0.0091597 | Lilrb4a |
| ENSMUSG00000064372 | 1173.17730974367 | 724.65728653727 | 0.56873 | 2.1634e-05 | 0.013813 | -//- |
| ENSMUSG00000069763 | 308.731091223443 | 94.2885701350182 | 0.81497 | 2.82e-06 | 0.0065624 | Tmem100 |
| ENSMUSG00000072596 | 37.6559320129037 | 5.85817747873321 | 0.73401 | 8.3062e-06 | 0.0091597 | Ear2 |
| ENSMUSG00000072620 | 133.548729536474 | 52.3537752797979 | 0.77541 | 5.7835e-06 | 0.0091597 | Slfn2 |
| ENSMUSG00000073421 | 1230.3363026074 | 440.149774472324 | 0.68531 | 8.1451e-05 | 0.029173 | H2-Ab1 |
| ENSMUSG00000079018 | 1618.26627631113 | 772.609348595751 | 0.65121 | 0.00011528 | 0.037515 | Ly6c1 |
| ENSMUSG00000097233 | 64.9621089152712 | 135.658897939805 | -0.72186 | 9.947e-06 | 0.0091597 | -//- |

| Term | Overlap | P-value | Adjusted P-value | Genes |
|---|---|---|---|---|
| neutrophil activation involved in immune response (GO:0002283) | 12/483 | 2.9981738632589026E-8 | 1.7959061440920828E-5 | PTPRC;SYK;SLC11A1;ITGB2;C5AR1;COTL1;C3AR1;CYBB;NCKAP1L;RETN;CTSS;CD55 |
| neutrophil degranulation (GO:0043312) | 11/479 | 2.655867971923774E-7 | 6.252698176770551E-5 | PTPRC;SLC11A1;ITGB2;C5AR1;COTL1;C3AR1;CYBB;NCKAP1L;RETN;CTSS;CD55 |
| neutrophil mediated immunity (GO:0002446) | 11/487 | 3.131568369000276E-7 | 6.252698176770551E-5 | PTPRC;SLC11A1;ITGB2;C5AR1;COTL1;C3AR1;CYBB;NCKAP1L;RETN;CTSS;CD55 |
| inflammatory response (GO:0006954) | 7/252 | 1.4068317157358681E-5 | 0.0021067304943144624 | NRROS;SYK;SLC11A1;ITGB2;C5AR1;C3AR1;CYBB |
| regulation of protein activation cascade (GO:2000257) | 5/108 | 2.291690762006761E-5 | 0.002227345730811466 | C1QB;C1QA;C5AR1;C3AR1;CD55 |
| regulation of complement activation (GO:0030449) | 5/109 | 2.3960804936406545E-5 | 0.002227345730811466 | C1QB;C1QA;C5AR1;C3AR1;CD55 |
| regulation of humoral immune response (GO:0002920) | 5/113 | 2.851149974163726E-5 | 0.002227345730811466 | C1QB;C1QA;C5AR1;C3AR1;CD55 |
| regulation of immune effector process (GO:0002697) | 5/114 | 2.9747522281288362E-5 | 0.002227345730811466 | C1QB;C1QA;C5AR1;C3AR1;CD55 |
| regulation of acute inflammatory response (GO:0002673) | 5/121 | 3.9607990367493474E-5 | 0.0024421469430699785 | C1QB;C1QA;C5AR1;C3AR1;CD55 |
| positive regulation of T cell proliferation (GO:0042102) | 4/61 | 4.0770399717361916E-5 | 0.0024421469430699785 | CD86;PTPRC;SYK;NCKAP1L |

**Supplementary Table 4. Top 500 differentially expressed genes analyzed by RNA-seq (AmotL2 KD vs WT in MS1) with enriched GO**

| Gene id | readcount MS1 AmotL2 KD | readcount MS1 wt | log2FoldChange | pval | padj | Gene Name |
|---|---|---|---|---|---|---|
| ENSMUSG00000042379 | 5259.3841596065 | 27408.3893708859 | -2.3817 | 2.0038e-291 | 4.6358e-287 | Esm1 |
| ENSMUSG00000025348 | 3608.35727844571 | 567.00397865601 | 2.6699 | 1.2177e-282 | 1.4086e-278 | Itga7 |
| ENSMUSG00000021367 | 359.956797901959 | 2314.55896240022 | -2.6848 | 1.9038e-233 | 1.4682e-229 | Edn1 |
| ENSMUSG00000017466 | 9073.66031585982 | 2219.32995276063 | 2.0316 | 7.283e-179 | 4.2123e-175 | Timp2 |
| ENSMUSG00000015568 | 311.773565666378 | 8.25511611154253 | 5.2391 | 6.7163e-160 | 3.1076e-156 | Lpl |
| ENSMUSG00000029338 | 3860.2508683871 | 1061.18211999841 | 1.863 | 2.2322e-155 | 8.6068e-152 | Antxr2 |
| ENSMUSG00000024066 | 1182.51064646542 | 241.782322059228 | 2.2901 | 8.9644e-151 | 2.9627e-147 | Xdh |
| ENSMUSG00000031274 | 2865.5548895776 | 536.108599229443 | 2.4182 | 1.7578e-147 | 5.0835e-144 | Col4a5 |
| ENSMUSG00000040322 | 2112.87119805396 | 538.677035172656 | 1.9717 | 1.714e-146 | 4.4059e-143 | Slc25a24 |
| ENSMUSG00000042842 | 1205.81383634838 | 255.208967159824 | 2.2403 | 4.4294e-146 | 1.0247e-142 | Serpinb6b |
| ENSMUSG00000021831 | 10723.4565497625 | 3173.00936386782 | 1.7568 | 6.603e-139 | 1.3887e-135 | Ero1l |
| ENSMUSG00000062488 | 3143.47835728854 | 934.339071682387 | 1.7503 | 3.6051e-132 | 6.9504e-129 | Ifit3b |
| ENSMUSG00000018217 | 3813.76664107883 | 1074.97828359022 | 1.8269 | 5.4619e-128 | 9.7201e-125 | Pmp22 |
| ENSMUSG00000021998 | 212.718298963859 | 957.985465077604 | -2.1711 | 1.2828e-123 | 2.1198e-120 | Lcp1 |
| ENSMUSG00000024659 | 13192.3664831882 | 4629.36652420929 | 1.5108 | 6.6315e-122 | 1.0228e-118 | Anxa1 |
| ENSMUSG00000023092 | 365.538818187736 | 35.5679281962848 | 3.3614 | 4.3189e-121 | 6.2448e-118 | Fhl1 |
| ENSMUSG00000027102 | 2440.12260417088 | 7013.75163050779 | -1.5232 | 3.0457e-118 | 4.1449e-115 | Hoxd8 |
| ENSMUSG00000026532 | 2770.50776862003 | 353.631158919681 | 2.9698 | 3.6616e-114 | 4.7062e-111 | Spta1 |
| ENSMUSG00000003849 | 3844.14774684543 | 1328.27562843434 | 1.5331 | 2.5692e-109 | 3.1284e-106 | Nqo1 |
| ENSMUSG00000044734 | 410.136196294577 | 57.0657694690756 | 2.8454 | 2.9439e-108 | 3.4054e-105 | Serpinb1a |
| ENSMUSG00000023904 | 995.306457340794 | 260.629501439919 | 1.9331 | 7.8098e-106 | 8.6038e-103 | Hcfc1r1 |
| ENSMUSG00000005413 | 13315.2542428233 | 3553.37564931676 | 1.9058 | 4.7944e-99 | 5.0417e-96 | Hmox1 |
| ENSMUSG00000034674 | 414.652970316461 | 1354.06104761672 | -1.7073 | 7.1128e-99 | 7.1545e-96 | Tdg |
| ENSMUSG00000001300 | 1021.51052703266 | 291.40020782279 | 1.8096 | 1.9001e-96 | 1.8316e-93 | Efnb2 |
| ENSMUSG00000032060 | 1002.25425465035 | 41.7452796675971 | 4.5855 | 3.53e-94 | 3.2666e-91 | Cryab |
| ENSMUSG00000005803 | 545.07068877765 | 77.1726081246563 | 2.8203 | 1.3883e-88 | 1.2353e-85 | Sqrdl |
| ENSMUSG00000074183 | 527.585090330456 | 6.85557268191534 | 6.266 | 6.0923e-88 | 5.2202e-85 | Gsta1 |
| ENSMUSG00000039457 | 335.970056804277 | 52.8268082515471 | 2.669 | 4.1542e-86 | 3.4324e-83 | Ppl |
| ENSMUSG00000031555 | 13473.2202559297 | 5778.08831369475 | 1.2214 | 1.5866e-82 | 1.2657e-79 | Adam9 |
| ENSMUSG00000031465 | 2492.13399120134 | 15865.7101148076 | -2.6705 | 1.29e-81 | 9.9484e-79 | Angpt2 |
| ENSMUSG00000031119 | 714.630055675109 | 154.357971457575 | 2.2109 | 1.3709e-80 | 1.0231e-77 | Gpc4 |
| ENSMUSG00000050148 | 5815.32733383099 | 13377.7202201727 | -1.2019 | 5.762e-80 | 4.1657e-77 | Ubqln2 |
| ENSMUSG00000033446 | 1271.89334418744 | 3551.57345059071 | -1.4815 | 8.6307e-80 | 6.0506e-77 | Lpar6 |
| ENSMUSG00000032366 | 4938.62011206794 | 11349.9187580179 | -1.2005 | 6.4917e-79 | 4.4172e-76 | Tpm1 |
| ENSMUSG00000026482 | 1096.57811488489 | 3754.44490836453 | -1.7756 | 1.5372e-78 | 1.0161e-75 | Rgl1 |
| ENSMUSG00000030774 | 942.174796438215 | 304.931965419674 | 1.6275 | 1.0223e-77 | 6.5697e-75 | Pak1 |
| ENSMUSG00000050335 | 270.205357865883 | 40.064017435903 | 2.7537 | 1.1595e-75 | 7.25e-73 | Lgals3 |
| ENSMUSG00000031647 | 2395.87880731676 | 7374.24125755686 | -1.6219 | 2.5629e-75 | 1.5603e-72 | Mfap3l |
| ENSMUSG00000025856 | 5369.53887999815 | 2310.35740169772 | 1.2167 | 2.9335e-75 | 1.7401e-72 | Pdgfa |
| ENSMUSG00000000489 | 2741.91937244839 | 10027.9260169223 | -1.8708 | 1.2077e-74 | 6.9853e-72 | Pdgfb |
| ENSMUSG00000024664 | 4672.04932449419 | 1943.09586603002 | 1.2657 | 4.374e-74 | 2.4681e-71 | Fads3 |
| ENSMUSG00000060600 | 1926.85959411789 | 748.287123506264 | 1.3646 | 1.5356e-73 | 8.4586e-71 | Eno3 |
| ENSMUSG00000039697 | 627.349078711775 | 1892.94820894016 | -1.5933 | 1.0579e-70 | 5.692e-68 | Ncoa7 |
| ENSMUSG00000087141 | 4460.52002722628 | 1765.3563121303 | 1.3373 | 4.6094e-70 | 2.4236e-67 | Plcxd2 |
| ENSMUSG00000021959 | 3893.02411727302 | 1700.91798428832 | 1.1946 | 1.0682e-69 | 5.492e-67 | Lats2 |
| ENSMUSG00000097636 | 1893.61350669377 | 4296.49586512696 | -1.182 | 2.8537e-69 | 1.4352e-66 | -//- |
| ENSMUSG00000027381 | 1366.48498547417 | 4370.81202602589 | -1.6774 | 4.0463e-69 | 1.9917e-66 | Bcl2l11 |
| ENSMUSG00000063011 | 927.48176990284 | 18.4060600343647 | 5.6551 | 5.0366e-69 | 2.4276e-66 | Msln |
| ENSMUSG00000000184 | 3162.47579601131 | 1362.74417145277 | 1.2145 | 6.8294e-69 | 3.2244e-66 | Ccnd2 |
| ENSMUSG00000107061 | 536.057031139816 | 1389.13621704779 | -1.3737 | 8.8581e-69 | 4.0986e-66 | -//- |
| ENSMUSG00000026872 | 1110.87985640187 | 342.990174749441 | 1.6955 | 4.3738e-67 | 1.9841e-64 | Zeb2 |
| ENSMUSG00000056091 | 730.481465094279 | 1783.34270151788 | -1.2877 | 1.6959e-66 | 7.545e-64 | St3gal5 |
| ENSMUSG00000024846 | 948.877922586051 | 340.767829688481 | 1.4774 | 5.3075e-66 | 2.3168e-63 | Cst6 |
| ENSMUSG00000052397 | 3359.23418004814 | 1464.13246556516 | 1.1981 | 1.2177e-65 | 5.2171e-63 | Ezr |
| ENSMUSG00000026837 | 3273.37313139934 | 1285.55715954256 | 1.3484 | 4.2019e-64 | 1.7675e-61 | Col5a1 |
| ENSMUSG00000039377 | 2520.11518369099 | 8070.01490014126 | -1.6791 | 9.7118e-64 | 4.0122e-61 | Hlx |
| ENSMUSG00000020173 | 667.749519294603 | 210.559925301666 | 1.6651 | 2.3265e-63 | 9.4426e-61 | Cobl |
| ENSMUSG00000016382 | 5005.54911173329 | 2319.90753637809 | 1.1095 | 5.7469e-63 | 2.2923e-60 | Pls3 |
| ENSMUSG00000009654 | 71.0024776362824 | 322.901002746124 | -2.1852 | 6.8089e-63 | 2.6699e-60 | Oit3 |
| ENSMUSG00000072872 | 1919.04965728453 | 4160.89901203222 | -1.1165 | 1.83e-62 | 7.0563e-60 | Rybp |
| ENSMUSG00000081534 | 6121.81692541213 | 2883.81402235899 | 1.086 | 2.2474e-62 | 8.5237e-60 | Slc48a1 |
| ENSMUSG00000038776 | 8203.10019870749 | 3933.42649080918 | 1.0604 | 1.9396e-61 | 7.2374e-59 | Ephx1 |
| ENSMUSG00000027276 | 1910.73660616497 | 726.463663075908 | 1.3952 | 5.1389e-61 | 1.8871e-58 | Jag1 |
| ENSMUSG00000001870 | 393.367246436996 | 83.7028245480704 | 2.2325 | 1.1164e-60 | 4.0357e-58 | Ltbp1 |
| ENSMUSG00000049112 | 16.5202283923974 | 161.016344056911 | -3.2849 | 4.0679e-60 | 1.4479e-57 | Oxtr |
| ENSMUSG00000026576 | 898.984548050063 | 136.438323783869 | 2.72 | 6.6328e-60 | 2.325e-57 | Atp1b1 |
| ENSMUSG00000032035 | 3107.43143675976 | 6481.99398734836 | -1.0607 | 8.2571e-60 | 2.8512e-57 | Ets1 |
| ENSMUSG00000055555 | 58.0338209682038 | 534.437113294149 | -3.2031 | 2.4852e-59 | 8.4552e-57 | 4930502E18Rik |
| ENSMUSG00000025037 | 4573.65807432182 | 1954.36785918408 | 1.2266 | 7.3475e-59 | 2.4635e-56 | Maoa |
| ENSMUSG00000056220 | 1770.80559236253 | 687.979206964062 | 1.364 | 5.3651e-58 | 1.7732e-55 | Pla2g4a |
| ENSMUSG00000036875 | 352.48867113423 | 914.746235844811 | -1.3758 | 1.4374e-57 | 4.6837e-55 | Dna2 |
| ENSMUSG00000034459 | 4101.38019931467 | 1274.0336892866 | 1.6867 | 3.9334e-57 | 1.2639e-54 | Ifit1 |
| ENSMUSG00000045362 | 5135.6902971291 | 2316.13448917244 | 1.1488 | 6.6688e-57 | 2.1134e-54 | Tnfrsf26 |
| ENSMUSG00000049580 | 1099.62881588625 | 447.932220122492 | 1.2957 | 1.1592e-56 | 3.6241e-54 | Tsku |
| ENSMUSG00000001025 | 18471.4126562055 | 7204.24958054712 | 1.3584 | 4.8654e-56 | 1.5008e-53 | S100a6 |
| ENSMUSG00000057346 | 4062.64371284751 | 1945.6478579482 | 1.0622 | 7.5988e-56 | 2.3131e-53 | Apol9a |
| ENSMUSG00000026674 | 881.877936132414 | 173.194293274041 | 2.3482 | 8.9876e-56 | 2.7004e-53 | Ddr2 |
| ENSMUSG00000004317 | 1512.61060974071 | 662.28209344299 | 1.1915 | 3.2264e-55 | 9.5697e-53 | Clcn5 |
| ENSMUSG00000064215 | 2886.18000891536 | 5866.93470609421 | -1.0234 | 3.9638e-55 | 1.1608e-52 | Ifi27 |
| ENSMUSG00000040447 | 1936.22158826575 | 4016.40658365156 | -1.0527 | 4.968e-55 | 1.4367e-52 | Spns2 |
| ENSMUSG00000056602 | 130.231495801032 | 431.286460432733 | -1.7276 | 9.5877e-55 | 2.7384e-52 | Fry |
| ENSMUSG00000105703 | 702.383022739657 | 1587.84019389653 | -1.1767 | 1.3372e-54 | 3.7727e-52 | -//- |
| ENSMUSG00000029082 | 3632.65335140063 | 1750.10867742636 | 1.0536 | 1.8744e-54 | 5.2247e-52 | Bst1 |
| ENSMUSG00000025185 | 3322.84194821492 | 1598.60231548818 | 1.0556 | 6.8967e-54 | 1.8995e-51 | Loxl4 |
| ENSMUSG00000024986 | 2396.12629952897 | 4845.5524430244 | -1.016 | 1.5404e-53 | 4.1926e-51 | Hhex |
| ENSMUSG00000002020 | 40.8552376139682 | 218.507175494367 | -2.4191 | 1.8983e-53 | 5.1066e-51 | Ltbp2 |
| ENSMUSG00000037010 | 21074.5344693616 | 40803.2603124499 | -0.95318 | 2.3757e-53 | 6.3174e-51 | Apln |
| ENSMUSG00000050708 | 87458.8074818202 | 42082.1700522303 | 1.0554 | 2.575e-53 | 6.7697e-51 | Ftl1 |
| ENSMUSG00000015312 | 679.737843615539 | 253.052722884478 | 1.4255 | 9.5686e-53 | 2.4873e-50 | Gadd45b |
| ENSMUSG00000001131 | 1332.27791186942 | 583.53092339057 | 1.191 | 1.8884e-52 | 4.8542e-50 | Timp1 |
| ENSMUSG00000030257 | 616.039496875924 | 1399.02023622 | -1.1833 | 3.3093e-52 | 8.4132e-50 | Srgap3 |
| ENSMUSG00000042985 | 87.2332952259815 | 0.719957334472493 | 6.9208 | 7.2666e-52 | 1.8273e-49 | Upk3b |
| ENSMUSG00000032231 | 15792.1264122333 | 6986.49776870799 | 1.1766 | 8.4885e-52 | 2.1116e-49 | Anxa2 |
| ENSMUSG00000025352 | 364.588228783538 | 1260.22117548597 | -1.7893 | 1.0742e-51 | 2.6437e-49 | Gdf11 |
| ENSMUSG00000028602 | 150.341178244863 | 463.106165157585 | -1.6231 | 1.1742e-51 | 2.8596e-49 | Tnfrsf8 |
| ENSMUSG00000035916 | 138.35151966156 | 15.080073776915 | 3.1976 | 1.355e-51 | 3.2654e-49 | Ptprq |
| ENSMUSG00000028713 | 145.2314974937 | 450.863274575398 | -1.6343 | 3.9794e-51 | 9.491e-49 | Cyp4b1 |
| ENSMUSG00000024803 | 5134.07038343519 | 1446.10341864672 | 1.8279 | 7.814e-51 | 1.8447e-48 | Ankrd1 |
| ENSMUSG00000049804 | 1074.03656170519 | 2251.57508905257 | -1.0679 | 1.1622e-50 | 2.7158e-48 | Armcx4 |
| ENSMUSG00000017309 | 31.337380794129 | 185.101386317401 | -2.5624 | 1.3004e-50 | 3.0084e-48 | Cd300lg |
| ENSMUSG00000029561 | 12838.463120444 | 6269.6647013576 | 1.034 | 1.7264e-50 | 3.9545e-48 | Oasl2 |
| ENSMUSG00000002222 | 669.37791942232 | 1545.6851802962 | -1.2074 | 3.7964e-50 | 8.6108e-48 | Rmnd5a |
| ENSMUSG00000022637 | 1152.7501231725 | 2395.19284572604 | -1.0551 | 5.1775e-50 | 1.1629e-47 | Cblb |
| ENSMUSG00000022055 | 2091.11817577702 | 576.37663328274 | 1.8592 | 2.6004e-49 | 5.7846e-47 | Nefl |
| ENSMUSG00000060594 | 619.214269640112 | 233.419180443159 | 1.4075 | 3.3494e-49 | 7.3799e-47 | Layn |
| ENSMUSG00000020422 | 4934.73136265334 | 2516.26854445365 | 0.97169 | 3.894e-49 | 8.4988e-47 | Tns3 |
| ENSMUSG00000013415 | 780.017747626936 | 178.483458460174 | 2.1277 | 6.82e-49 | 1.4746e-46 | Igf2bp1 |
| ENSMUSG00000031963 | 377.999463432245 | 87.719338165558 | 2.1074 | 2.6834e-48 | 5.7481e-46 | Bmper |
| ENSMUSG00000027447 | 7197.90293539115 | 3756.1721479715 | 0.93831 | 3.1403e-48 | 6.6303e-46 | Cst3 |
| ENSMUSG00000055737 | 1707.40176859791 | 3398.60250620413 | -0.99314 | 3.1525e-48 | 6.6303e-46 | Ghr |
| ENSMUSG00000038482 | 1672.34030982367 | 3329.03814193206 | -0.99324 | 3.3145e-48 | 6.9082e-46 | Tfdp1 |
| ENSMUSG00000074896 | 7510.46618298711 | 2131.24701681792 | 1.8172 | 4.1975e-48 | 8.6705e-46 | Ifit3 |
| ENSMUSG00000027078 | 3038.12305562134 | 1125.80836779724 | 1.4322 | 5.2543e-48 | 1.0757e-45 | Ube2l6 |
| ENSMUSG00000063531 | 473.861852774841 | 142.550459025996 | 1.733 | 1.0892e-46 | 2.2104e-44 | Sema3e |
| ENSMUSG00000027962 | 5329.98168423507 | 1759.79285988836 | 1.5987 | 5.059e-46 | 1.0177e-43 | Vcam1 |
| ENSMUSG00000022464 | 1567.66330874174 | 671.903586506724 | 1.2223 | 1.0182e-45 | 2.0308e-43 | Slc38a4 |
| ENSMUSG00000027004 | 168.815601203469 | 471.705843842815 | -1.4824 | 4.7153e-45 | 9.3238e-43 | Frzb |
| ENSMUSG00000028961 | 5196.66384801167 | 2745.98154964623 | 0.92026 | 4.7559e-45 | 9.3245e-43 | Pgd |
| ENSMUSG00000058135 | 2303.59902768913 | 880.10453877063 | 1.3881 | 5.2139e-45 | 1.0136e-42 | Gstm1 |
| ENSMUSG00000031093 | 285.028440202731 | 45.1163567753186 | 2.6594 | 9.6402e-45 | 1.8585e-42 | Dock11 |
| ENSMUSG00000041827 | 7421.6118148342 | 3813.00101339628 | 0.96081 | 3.8278e-44 | 7.3186e-42 | Oasl1 |
| ENSMUSG00000062585 | 6.4219515733932 | 93.6240543157966 | -3.8658 | 8.1352e-44 | 1.5427e-41 | Cnr2 |
| ENSMUSG00000036087 | 1532.93187542022 | 2961.59975145057 | -0.95008 | 2.3094e-43 | 4.3437e-41 | Slain2 |
| ENSMUSG00000029635 | 2120.80569687178 | 1070.43881308881 | 0.98641 | 2.4906e-43 | 4.6468e-41 | Cdk8 |
| ENSMUSG00000038650 | 3772.32967473529 | 1989.75747583492 | 0.92286 | 4.1234e-43 | 7.6316e-41 | Rnh1 |
| ENSMUSG00000023088 | 5080.33925733489 | 2090.75809933885 | 1.2809 | 4.2021e-43 | 7.7155e-41 | Abcc1 |
| ENSMUSG00000033855 | 3924.34218722248 | 2080.56030997554 | 0.91548 | 6.7026e-43 | 1.221e-40 | Ston1 |
| ENSMUSG00000027068 | 423.669995025903 | 91.9720334634945 | 2.2037 | 7.4167e-43 | 1.3405e-40 | Dhrs9 |
| ENSMUSG00000028124 | 6465.83765366166 | 3494.84577324088 | 0.88761 | 1.1349e-42 | 2.0354e-40 | Gclm |
| ENSMUSG00000020641 | 11924.6477947054 | 4964.63762734851 | 1.2642 | 1.9602e-42 | 3.4885e-40 | Rsad2 |
| ENSMUSG00000032661 | 5144.07772168216 | 1453.53602592142 | 1.8233 | 4.9304e-42 | 8.7072e-40 | Oas3 |
| ENSMUSG00000052428 | 1468.94474691362 | 718.180605692296 | 1.0324 | 4.9738e-42 | 8.7173e-40 | Tmco1 |
| ENSMUSG00000019564 | 79.4498282175938 | 274.354810992519 | -1.7879 | 7.391e-42 | 1.2856e-39 | Arid3a |
| ENSMUSG00000029718 | 409.679791659304 | 67.6789181611141 | 2.5977 | 7.5894e-42 | 1.3103e-39 | Pcolce |
| ENSMUSG00000086316 | 1054.18751356642 | 2068.09391368472 | -0.97217 | 9.5125e-42 | 1.6302e-39 | 2210013O21Rik |
| ENSMUSG00000028634 | 1478.34562656128 | 490.573488919649 | 1.5914 | 1.0095e-41 | 1.7173e-39 | Hivep3 |
| ENSMUSG00000021187 | 117.939769284584 | 422.125852575832 | -1.8396 | 1.649e-41 | 2.7846e-39 | Tc2n |
| ENSMUSG00000035407 | 53.4404183468191 | 223.069179201148 | -2.0615 | 2.7551e-41 | 4.6187e-39 | Kank4 |
| ENSMUSG00000024219 | 777.04415898949 | 1565.167622898 | -1.0102 | 4.2425e-41 | 7.0612e-39 | Anks1 |
| ENSMUSG00000005763 | 97.005030460204 | 307.498784253201 | -1.6644 | 5.0579e-41 | 8.3581e-39 | Cd247 |
| ENSMUSG00000036136 | 363.931790360006 | 812.624568798512 | -1.1589 | 7.0946e-41 | 1.1641e-38 | Fam110c |
| ENSMUSG00000029484 | 9971.50954865036 | 4490.33695692815 | 1.151 | 7.6584e-41 | 1.2477e-38 | Anxa3 |
| ENSMUSG00000015837 | 23636.5534834131 | 13315.0435489779 | 0.82796 | 1.2766e-40 | 2.0654e-38 | Sqstm1 |
| ENSMUSG00000005686 | 3330.9393884886 | 1788.53085299737 | 0.89715 | 2.9045e-40 | 4.6663e-38 | Ampd3 |
| ENSMUSG00000075595 | 702.145931755174 | 1418.94910195444 | -1.015 | 3.8931e-40 | 6.2114e-38 | Zfp652 |
| ENSMUSG00000020399 | 314.091827049647 | 75.1486208994504 | 2.0634 | 1.1605e-39 | 1.8389e-37 | Havcr2 |
| ENSMUSG00000030101 | 1428.12924626203 | 714.235087663881 | 0.99966 | 1.9873e-39 | 3.1277e-37 | Sumf1 |
| ENSMUSG00000003032 | 802.5818843913 | 301.77555612853 | 1.4112 | 2.228e-39 | 3.4828e-37 | Klf4 |
| ENSMUSG00000028792 | 945.384748972414 | 1829.39428482974 | -0.95239 | 3.9221e-39 | 6.0898e-37 | Ak2 |
| ENSMUSG00000004880 | 1984.89088424092 | 3647.87404402786 | -0.878 | 4.2726e-39 | 6.5897e-37 | Lbr |
| ENSMUSG00000000420 | 3548.95718374664 | 6362.47341636467 | -0.84219 | 5.4334e-39 | 8.3247e-37 | Galnt1 |
| ENSMUSG00000046546 | 3872.98548620916 | 10626.3104194543 | -1.4561 | 5.8921e-39 | 8.968e-37 | Fam43a |
| ENSMUSG00000032870 | 2362.00367570148 | 4288.74678057068 | -0.86054 | 1.6478e-38 | 2.4916e-36 | Smap2 |
| ENSMUSG00000041957 | 490.353117916963 | 1022.05362191733 | -1.0596 | 1.8168e-38 | 2.7293e-36 | Pkp2 |
| ENSMUSG00000027907 | 16512.2569553408 | 9409.21974613033 | 0.81139 | 2.0277e-38 | 3.0265e-36 | S100a11 |
| ENSMUSG00000041012 | 40.4472492412427 | 177.294045513892 | -2.132 | 2.4406e-38 | 3.6195e-36 | Cmtm8 |
| ENSMUSG00000024661 | 72337.3423837061 | 33383.7450334939 | 1.1156 | 2.5172e-38 | 3.7093e-36 | Fth1 |
| ENSMUSG00000032802 | 5319.88563195429 | 2965.0895080333 | 0.84332 | 3.9465e-38 | 5.7787e-36 | Srxn1 |
| ENSMUSG00000030787 | 765.238460645003 | 1739.08966215193 | -1.1844 | 7.7283e-38 | 1.1245e-35 | Lyve1 |
| ENSMUSG00000027011 | 1434.17646365733 | 731.082332669672 | 0.97212 | 9.1654e-38 | 1.3253e-35 | Ube2e3 |
| ENSMUSG00000004319 | 1413.10409356065 | 2894.16183098438 | -1.0343 | 2.1346e-37 | 3.0674e-35 | Clcn3 |
| ENSMUSG00000066894 | 1116.43953101039 | 2100.24328725232 | -0.91165 | 3.2943e-37 | 4.7046e-35 | Vsig10 |
| ENSMUSG00000024063 | 2634.70149864351 | 1047.61396655123 | 1.3305 | 4.9687e-37 | 7.0522e-35 | Lbh |
| ENSMUSG00000014599 | 13463.6368900785 | 4224.99796400083 | 1.672 | 6.0852e-37 | 8.5842e-35 | Csf1 |
| ENSMUSG00000020262 | 1190.54464474032 | 2220.52685442687 | -0.89928 | 8.0422e-37 | 1.1276e-34 | Adarb1 |
| ENSMUSG00000030790 | 1156.13345297191 | 2605.38597726705 | -1.1722 | 1.4637e-36 | 2.0399e-34 | Adm |
| ENSMUSG00000021745 | 1294.04429893475 | 629.322461152252 | 1.04 | 1.4976e-36 | 2.0746e-34 | Ptprg |
| ENSMUSG00000035107 | 2036.60771697894 | 994.268205124853 | 1.0345 | 3.2076e-36 | 4.4171e-34 | Dcbld2 |
| ENSMUSG00000005087 | 4118.19708633062 | 2308.2346246202 | 0.83522 | 3.9874e-36 | 5.4584e-34 | Cd44 |
| ENSMUSG00000015365 | 227.266455160667 | 35.2918466569641 | 2.687 | 4.3103e-36 | 5.8659e-34 | Mov10l1 |
| ENSMUSG00000071637 | 239.448261334665 | 738.44835141701 | -1.6248 | 4.6207e-36 | 6.2515e-34 | -//- |
| ENSMUSG00000029366 | 1083.72098283776 | 2012.92127410773 | -0.8933 | 5.5325e-36 | 7.4416e-34 | Dck |
| ENSMUSG00000028063 | 6935.46526801434 | 3970.79645936072 | 0.80456 | 6.2388e-36 | 8.3431e-34 | Lmna |
| ENSMUSG00000037138 | 106.296134135811 | 309.731305411042 | -1.5429 | 6.4056e-36 | 8.5168e-34 | Aff3 |
| ENSMUSG00000037071 | 1184.82211729394 | 599.916056024769 | 0.98184 | 7.8427e-36 | 1.0368e-33 | Scd1 |
| ENSMUSG00000060519 | 2221.61568867582 | 1202.89893297594 | 0.88509 | 8.824e-36 | 1.1599e-33 | Tor3a |
| ENSMUSG00000002489 | 417.791831575725 | 992.312017658688 | -1.248 | 1.5425e-35 | 2.0162e-33 | Tiam1 |
| ENSMUSG00000024892 | 766.593529893772 | 364.477777478787 | 1.0726 | 3.5225e-35 | 4.5782e-33 | Pcx |
| ENSMUSG00000031827 | 805.069039899454 | 1544.67126300178 | -0.94012 | 4.028e-35 | 5.2061e-33 | Cotl1 |
| ENSMUSG00000022106 | 571.976145942783 | 1131.59905000457 | -0.98434 | 4.1652e-35 | 5.3535e-33 | Rcbtb2 |
| ENSMUSG00000019791 | 249.653515997908 | 608.622766978427 | -1.2856 | 4.8622e-35 | 6.2148e-33 | Hint3 |
| ENSMUSG00000031273 | 256.496127286772 | 82.3724248887433 | 1.6387 | 4.907e-35 | 6.2376e-33 | Col4a6 |
| ENSMUSG00000045095 | 1168.68538814426 | 2375.73204355072 | -1.0235 | 5.4983e-35 | 6.951e-33 | Magi1 |
| ENSMUSG00000019970 | 670.790653760037 | 1295.52476808526 | -0.9496 | 8.1818e-35 | 1.0287e-32 | Sgk1 |
| ENSMUSG00000013662 | 3399.17783629291 | 1912.92713870317 | 0.8294 | 1.1069e-34 | 1.3842e-32 | Atad1 |
| ENSMUSG00000048120 | 3126.07228768698 | 1677.8643189622 | 0.89773 | 1.4695e-34 | 1.8278e-32 | Entpd1 |
| ENSMUSG00000054871 | 889.588932823954 | 2288.65175021388 | -1.3633 | 1.5698e-34 | 1.9422e-32 | Tmem158 |
| ENSMUSG00000038267 | 786.39029147021 | 1489.77967163631 | -0.92178 | 2.0469e-34 | 2.5189e-32 | Slc22a23 |
| ENSMUSG00000041301 | 0.726430787056432 | 108.669261607656 | -7.2249 | 2.1522e-34 | 2.6344e-32 | Cftr |
| ENSMUSG00000005672 | 2421.6747808258 | 9177.78561244667 | -1.9221 | 2.3825e-34 | 2.9011e-32 | Kit |
| ENSMUSG00000025241 | 953.190356896481 | 474.692183142959 | 1.0058 | 3.2175e-34 | 3.8772e-32 | Fyco1 |
| ENSMUSG00000039155 | 107.839137083447 | 405.043948447361 | -1.9092 | 3.2177e-34 | 3.8772e-32 | Cdh26 |
| ENSMUSG00000022707 | 745.206984359393 | 355.295962816459 | 1.0686 | 3.3626e-34 | 4.0307e-32 | Gbe1 |
| ENSMUSG00000056025 | 6166.66478104541 | 1929.46815982013 | 1.6763 | 3.4332e-34 | 4.0941e-32 | Clca3a1 |
| ENSMUSG00000102204 | 590.416504202135 | 96.3839614538498 | 2.6149 | 4.1465e-34 | 4.9195e-32 | -//- |
| ENSMUSG00000043873 | 146.371791969624 | 373.801650699807 | -1.3526 | 4.5979e-34 | 5.4271e-32 | Chil5 |
| ENSMUSG00000023048 | 3273.49241675248 | 1852.62799819013 | 0.82126 | 7.0585e-34 | 8.206e-32 | Prr13 |
| ENSMUSG00000029191 | 2957.92415780519 | 1665.20437532922 | 0.82889 | 7.0074e-34 | 8.206e-32 | Rfc1 |
| ENSMUSG00000104045 | 351.398725157448 | 94.4321337914547 | 1.8958 | 7.0488e-34 | 8.206e-32 | -//- |
| ENSMUSG00000015090 | 168.871913860406 | 16.3440044685866 | 3.3691 | 8.2436e-34 | 9.5357e-32 | Ptgds |
| ENSMUSG00000047045 | 1017.56397662623 | 1869.39164639736 | -0.87745 | 9.6523e-34 | 1.111e-31 | Tmem164 |
| ENSMUSG00000043993 | 155.068785329773 | 389.976036576239 | -1.3305 | 1.1071e-33 | 1.2679e-31 | -//- |
| ENSMUSG00000026308 | 84.597462244844 | 0.234883275400345 | 8.4925 | 1.269e-33 | 1.4462e-31 | Klhl30 |
| ENSMUSG00000018169 | 3708.30076114188 | 6361.87601434999 | -0.77869 | 1.4092e-33 | 1.5981e-31 | Mfng |
| ENSMUSG00000016087 | 2584.27123222288 | 4490.30260556789 | -0.79706 | 1.7154e-33 | 1.9359e-31 | Fli1 |
| ENSMUSG00000040133 | 640.859728192097 | 296.124464823128 | 1.1138 | 1.7267e-33 | 1.9392e-31 | Gpr176 |
| ENSMUSG00000024423 | 3108.83798667915 | 1738.69795657576 | 0.83837 | 1.7379e-33 | 1.9424e-31 | Impact |
| ENSMUSG00000025089 | 294.050644017971 | 104.87606196407 | 1.4874 | 2.4792e-33 | 2.7575e-31 | Gfra1 |
| ENSMUSG00000019966 | 2089.56980348197 | 1153.46890454 | 0.85723 | 2.68e-33 | 2.9666e-31 | Kitl |
| ENSMUSG00000022749 | 1192.5226532838 | 2493.0898978975 | -1.0639 | 3.0828e-33 | 3.3962e-31 | Tbc1d23 |
| ENSMUSG00000030107 | 2586.86062240895 | 1110.65042850839 | 1.2198 | 3.904e-33 | 4.2805e-31 | Usp18 |
| ENSMUSG00000030591 | 3891.89271625579 | 2238.44813926159 | 0.79797 | 4.3928e-33 | 4.7937e-31 | Psmd8 |
| ENSMUSG00000003948 | 953.93350766075 | 2130.75505103504 | -1.1594 | 6.1789e-33 | 6.707e-31 | Mmd |
| ENSMUSG00000038347 | 355.743119200983 | 139.239919967278 | 1.3533 | 6.204e-33 | 6.707e-31 | Tcte2 |
| ENSMUSG00000069874 | 1883.5849746391 | 878.798974993309 | 1.0999 | 9.2474e-33 | 9.9506e-31 | Irgm2 |
| ENSMUSG00000042659 | 455.1182574969 | 193.315907798507 | 1.2353 | 1.0423e-32 | 1.1164e-30 | Arrdc4 |
| ENSMUSG00000028780 | 156.979348750097 | 10.0185563853443 | 3.9698 | 1.0518e-32 | 1.1213e-30 | Sema3c |
| ENSMUSG00000094441 | 663.787291056091 | 312.952026785172 | 1.0848 | 1.1424e-32 | 1.2124e-30 | Zfp955a |
| ENSMUSG00000017929 | 8458.50326807486 | 4181.72609419477 | 1.0163 | 1.5296e-32 | 1.6158e-30 | B4galt5 |
| ENSMUSG00000030256 | 1031.6202324391 | 531.714174981287 | 0.95619 | 1.6066e-32 | 1.6895e-30 | Bhlhe41 |
| ENSMUSG00000026475 | 971.377372676708 | 2070.78394975207 | -1.0921 | 1.7916e-32 | 1.8755e-30 | Rgs16 |
| ENSMUSG00000018474 | 1790.83180966618 | 3914.64451909877 | -1.1283 | 1.805e-32 | 1.881e-30 | Chd3 |
| ENSMUSG00000060639 | 347.913435948814 | 136.246253631024 | 1.3525 | 1.9127e-32 | 1.9844e-30 | Hist1h4i |
| ENSMUSG00000038059 | 496.401472099773 | 219.064181075192 | 1.1802 | 2.542e-32 | 2.6254e-30 | Smim3 |
| ENSMUSG00000026509 | 8124.98599852392 | 4805.68041173416 | 0.75762 | 2.613e-32 | 2.6867e-30 | Capn2 |
| ENSMUSG00000042284 | 1115.42826631791 | 2050.41089742616 | -0.87832 | 2.7729e-32 | 2.8386e-30 | Itga1 |
| ENSMUSG00000030353 | 822.857103425006 | 393.487464569203 | 1.0643 | 2.8797e-32 | 2.9349e-30 | Tead4 |
| ENSMUSG00000087028 | 839.961690199659 | 1544.83017979419 | -0.87905 | 4.4989e-32 | 4.565e-30 | -//- |
| ENSMUSG00000032388 | 1478.75252179838 | 802.792607599388 | 0.88128 | 5.2509e-32 | 5.3047e-30 | Spg21 |
| ENSMUSG00000055407 | 980.82166623455 | 355.783229040131 | 1.463 | 5.6097e-32 | 5.6426e-30 | Map6 |
| ENSMUSG00000024440 | 901.083278016599 | 1904.23551170019 | -1.0795 | 5.6799e-32 | 5.664e-30 | Pcdh12 |
| ENSMUSG00000030096 | 1287.75347032292 | 675.767630978608 | 0.93026 | 5.672e-32 | 5.664e-30 | Slc6a6 |
| ENSMUSG00000020156 | 2711.06266743391 | 4624.17906184469 | -0.77034 | 7.7232e-32 | 7.6685e-30 | Mum1 |
| ENSMUSG00000032601 | 1074.86607519504 | 1928.97153025069 | -0.84367 | 9.2662e-32 | 9.1612e-30 | Prkar2a |
| ENSMUSG00000076441 | 4518.13676771392 | 2645.70170790436 | 0.77208 | 1.18e-31 | 1.1617e-29 | Ass1 |
| ENSMUSG00000024420 | 1157.05423373457 | 2120.6683427005 | -0.87406 | 1.4054e-31 | 1.3719e-29 | Zfp521 |
| ENSMUSG00000038963 | 162.315343801665 | 797.222512820765 | -2.2962 | 1.4018e-31 | 1.3719e-29 | Slco4a1 |
| ENSMUSG00000031628 | 1163.2887729773 | 2066.7003536415 | -0.82912 | 1.5759e-31 | 1.5319e-29 | Casp3 |
| ENSMUSG00000029334 | 81.3490260551527 | 9.30045542282583 | 3.1288 | 1.778e-31 | 1.7211e-29 | Prkg2 |
| ENSMUSG00000021403 | 2497.42647755553 | 517.727119079014 | 2.2702 | 1.8125e-31 | 1.7472e-29 | Serpinb9b |
| ENSMUSG00000064373 | 2108.76716854542 | 7408.87401983129 | -1.8129 | 3.0223e-31 | 2.9013e-29 | Sepp1 |
| ENSMUSG00000006373 | 4537.09132662321 | 2670.07735388035 | 0.76489 | 5.3556e-31 | 5.1199e-29 | Pgrmc1 |
| ENSMUSG00000028883 | 370.319682209112 | 152.255832952478 | 1.2823 | 5.7216e-31 | 5.4473e-29 | Sema3a |
| ENSMUSG00000021662 | 794.861558277742 | 1452.80786644518 | -0.87007 | 9.1639e-31 | 8.6888e-29 | Arhgef28 |
| ENSMUSG00000004655 | 300.882148602449 | 114.350107132266 | 1.3957 | 9.2292e-31 | 8.715e-29 | Aqp1 |
| ENSMUSG00000035692 | 2475.08211923528 | 879.441017497137 | 1.4928 | 1.0363e-30 | 9.7455e-29 | Isg15 |
| ENSMUSG00000021466 | 391.203846747413 | 893.117652205793 | -1.1909 | 1.344e-30 | 1.2588e-28 | Ptch1 |
| ENSMUSG00000031442 | 205.926385025409 | 463.109385708123 | -1.1692 | 1.6706e-30 | 1.5584e-28 | Mcf2l |
| ENSMUSG00000018849 | 985.419474998121 | 318.410910295452 | 1.6298 | 2.2392e-30 | 2.0805e-28 | Wwc1 |
| ENSMUSG00000032782 | 552.954969963923 | 1141.17481836564 | -1.0453 | 2.2524e-30 | 2.0844e-28 | Cntrob |
| ENSMUSG00000028212 | 332.214252409412 | 675.746256708439 | -1.0244 | 2.4867e-30 | 2.292e-28 | Ccne2 |
| ENSMUSG00000018983 | 96.2164712924599 | 267.315859372451 | -1.4742 | 2.6375e-30 | 2.4214e-28 | E2f2 |
| ENSMUSG00000026701 | 16480.0788199782 | 10039.4634507146 | 0.71504 | 2.7712e-30 | 2.5341e-28 | Prdx6 |
| ENSMUSG00000028832 | 1788.63976201002 | 3066.02381078254 | -0.77751 | 2.9692e-30 | 2.7044e-28 | Stmn1 |
| ENSMUSG00000023034 | 352.766254777515 | 748.950292794031 | -1.0862 | 3.7791e-30 | 3.4286e-28 | Nr4a1 |
| ENSMUSG00000024646 | 2083.87553034921 | 1190.23871140101 | 0.80802 | 5.0482e-30 | 4.5621e-28 | Cyb5a |
| ENSMUSG00000071656 | 549.551089469518 | 148.697334465257 | 1.8859 | 6.1478e-30 | 5.5342e-28 | Lrrn4cl |
| ENSMUSG00000020811 | 113.155387994605 | 295.576208923199 | -1.3852 | 6.4056e-30 | 5.7439e-28 | Wscd1 |
| ENSMUSG00000051855 | 3241.64983356363 | 9962.28948974473 | -1.6197 | 7.7492e-30 | 6.9219e-28 | Mest |
| ENSMUSG00000040339 | 967.115300188709 | 2657.21070214416 | -1.4582 | 8.7927e-30 | 7.8238e-28 | Fam102b |
| ENSMUSG00000026104 | 1075.74390118242 | 555.030582123485 | 0.9547 | 1.2401e-29 | 1.0992e-27 | Stat1 |
| ENSMUSG00000057329 | 452.643373157264 | 871.420715960535 | -0.94499 | 1.5599e-29 | 1.3774e-27 | Bcl2 |
| ENSMUSG00000086763 | 754.768364927907 | 1361.12466803635 | -0.85069 | 1.9229e-29 | 1.6915e-27 | -//- |
| ENSMUSG00000005054 | 1614.84543649881 | 907.829537070573 | 0.8309 | 2.1934e-29 | 1.9222e-27 | Cstb |
| ENSMUSG00000029470 | 906.20338830803 | 476.894647798443 | 0.92616 | 3.0232e-29 | 2.6393e-27 | P2rx4 |
| ENSMUSG00000057561 | 3233.76384385112 | 1909.72099598344 | 0.75985 | 3.6084e-29 | 3.1384e-27 | Eif1a |
| ENSMUSG00000041959 | 5899.25723002484 | 3388.66704290104 | 0.79982 | 4.3721e-29 | 3.7884e-27 | S100a10 |
| ENSMUSG00000002068 | 375.246404652863 | 736.489078291493 | -0.97283 | 5.7172e-29 | 4.9354e-27 | Ccne1 |
| ENSMUSG00000060147 | 3172.65795135687 | 1647.07966258791 | 0.94578 | 7.2898e-29 | 6.2695e-27 | Serpinb6a |
| ENSMUSG00000030494 | 422.985619973323 | 84.4949437219446 | 2.3237 | 7.4964e-29 | 6.4233e-27 | Rhpn2 |
| ENSMUSG00000045827 | 445.827358715052 | 200.597904801318 | 1.1522 | 7.7814e-29 | 6.6429e-27 | Serpinb9 |
| ENSMUSG00000062168 | 101.011378438745 | 12.3487711593169 | 3.0321 | 8.0978e-29 | 6.8876e-27 | Ppef1 |
| ENSMUSG00000020393 | 1132.23640890977 | 1989.13434605074 | -0.81297 | 8.2031e-29 | 6.9516e-27 | Kremen1 |
| ENSMUSG00000019768 | 195.040859523615 | 61.8155272686842 | 1.6577 | 9.9184e-29 | 8.3746e-27 | Esr1 |
| ENSMUSG00000050565 | 6571.81913272336 | 3998.34904912433 | 0.71689 | 1.0542e-28 | 8.8689e-27 | Tor1aip2 |
| ENSMUSG00000029086 | 6005.4239022046 | 11711.6722394555 | -0.96361 | 1.3305e-28 | 1.1153e-26 | Prom1 |
| ENSMUSG00000019528 | 1524.56802032734 | 860.752029693024 | 0.82473 | 1.8505e-28 | 1.5455e-26 | Gyg |
| ENSMUSG00000054932 | 109.539933365927 | 21.885433019488 | 2.3234 | 2.0073e-28 | 1.6704e-26 | Afp |
| ENSMUSG00000038151 | 68.6001897819887 | 369.693955149425 | -2.43 | 2.1615e-28 | 1.7924e-26 | Prdm1 |
| ENSMUSG00000057604 | 898.333541754494 | 1935.56934538445 | -1.1074 | 2.3358e-28 | 1.93e-26 | Lmcd1 |
| ENSMUSG00000040274 | 983.213516797233 | 502.431409931343 | 0.96858 | 2.6808e-28 | 2.2072e-26 | Cdk6 |
| ENSMUSG00000038648 | 2029.8401738336 | 3709.04584593503 | -0.86968 | 4.9837e-28 | 4.0886e-26 | Creb3l2 |
| ENSMUSG00000030785 | 66.5740333443078 | 6.63742853428644 | 3.3263 | 5.3746e-28 | 4.3937e-26 | Cox6a2 |
| ENSMUSG00000027210 | 3448.9664680238 | 5654.37349286041 | -0.7132 | 7.0389e-28 | 5.734e-26 | Meis2 |
| ENSMUSG00000023960 | 816.833824798435 | 411.419972962558 | 0.98943 | 9.5729e-28 | 7.7708e-26 | Enpp5 |
| ENSMUSG00000054951 | 225.678983870173 | 45.3265375751824 | 2.3158 | 1.0473e-27 | 8.4717e-26 | 9130008F23Rik |
| ENSMUSG00000021589 | 1293.81937541192 | 724.06043340075 | 0.83745 | 1.0598e-27 | 8.5427e-26 | Rhobtb3 |
| ENSMUSG00000095550 | 56.6856633567448 | 180.458452948877 | -1.6706 | 1.0783e-27 | 8.662e-26 | Gm21671 |
| ENSMUSG00000048482 | 201.314248141963 | 62.0391631488376 | 1.6982 | 1.3141e-27 | 1.0519e-25 | Bdnf |
| ENSMUSG00000046873 | 595.901881953344 | 1084.35106853781 | -0.86369 | 1.3998e-27 | 1.1167e-25 | Mbtps2 |
| ENSMUSG00000046718 | 26162.2285111977 | 16364.6312839634 | 0.6769 | 1.5464e-27 | 1.2294e-25 | Bst2 |
| ENSMUSG00000034118 | 703.517125431201 | 1253.00704992198 | -0.83274 | 1.604e-27 | 1.2709e-25 | Tpst1 |
| ENSMUSG00000027339 | 301.827880043238 | 885.472213313031 | -1.5527 | 1.7161e-27 | 1.355e-25 | Rassf2 |
| ENSMUSG00000072235 | 3438.09917503062 | 2072.77711426433 | 0.73005 | 1.7795e-27 | 1.4003e-25 | Tuba1a |
| ENSMUSG00000035790 | 132.29861565601 | 318.812904823608 | -1.2689 | 2.4201e-27 | 1.898e-25 | Cep19 |
| ENSMUSG00000078532 | 403.921904362301 | 181.664805933744 | 1.1528 | 2.5098e-27 | 1.9616e-25 | Nkain1 |
| ENSMUSG00000063524 | 22193.119833895 | 13945.0490791514 | 0.67036 | 2.7229e-27 | 2.121e-25 | Eno1 |
| ENSMUSG00000041126 | 1068.24403268492 | 1836.74222089487 | -0.78191 | 2.7417e-27 | 2.1285e-25 | H2afv |
| ENSMUSG00000033499 | 907.243287275074 | 1576.38144977714 | -0.79706 | 3.1273e-27 | 2.4198e-25 | Larp4b |
| ENSMUSG00000023143 | 871.195930567868 | 469.208273064763 | 0.89277 | 5.1525e-27 | 3.9734e-25 | Nagpa |
| ENSMUSG00000037428 | 111.141846162995 | 278.864633606577 | -1.3272 | 6.236e-27 | 4.793e-25 | Vgf |
| ENSMUSG00000029377 | 456.165522336796 | 99.8539492554421 | 2.1917 | 6.4188e-27 | 4.9172e-25 | Ereg |
| ENSMUSG00000047604 | 130.291080506572 | 412.811942391588 | -1.6637 | 6.6062e-27 | 5.044e-25 | Frat2 |
| ENSMUSG00000068246 | 3343.23174813868 | 1846.70039711053 | 0.85629 | 8.0125e-27 | 6.0977e-25 | Apol9b |
| ENSMUSG00000032006 | 711.087046125749 | 1285.20404731645 | -0.8539 | 9.0634e-27 | 6.8748e-25 | Pdgfd |
| ENSMUSG00000055723 | 4687.14115996062 | 2390.58654396194 | 0.97134 | 1.4751e-26 | 1.1153e-24 | Rras2 |
| ENSMUSG00000026715 | 65.4989139236176 | 7.09003120486167 | 3.2076 | 1.5614e-26 | 1.1766e-24 | Serpinc1 |
| ENSMUSG00000031586 | 469.581128604649 | 868.14300220516 | -0.88656 | 1.6584e-26 | 1.2457e-24 | Rbpms |
| ENSMUSG00000031453 | 2284.81820027111 | 3741.44063582436 | -0.71151 | 2.0671e-26 | 1.5477e-24 | Rasa3 |
| ENSMUSG00000024087 | 1238.74164684943 | 337.714924414241 | 1.875 | 2.1053e-26 | 1.5711e-24 | Cyp1b1 |
| ENSMUSG00000033470 | 5.1297972432995 | 58.8922690176718 | -3.5211 | 2.1595e-26 | 1.6064e-24 | Cysltr2 |
| ENSMUSG00000055421 | 203.695100278717 | 71.1599628671208 | 1.5173 | 2.506e-26 | 1.8582e-24 | Pcdh9 |
| ENSMUSG00000079852 | 1521.90433241175 | 512.381939988892 | 1.5706 | 2.7864e-26 | 2.0595e-24 | Klra4 |
| ENSMUSG00000025498 | 8587.44457879762 | 3506.04543948525 | 1.2924 | 2.8301e-26 | 2.0852e-24 | Irf7 |
| ENSMUSG00000073490 | 1151.40857417478 | 646.660497561715 | 0.83232 | 3.0278e-26 | 2.2238e-24 | AI607873 |
| ENSMUSG00000036585 | 12.6941595583091 | 158.990065923563 | -3.6467 | 3.1509e-26 | 2.3068e-24 | Fgf1 |
| ENSMUSG00000026582 | 1000.22174223783 | 528.622405644498 | 0.92001 | 3.5151e-26 | 2.5654e-24 | Sele |
| ENSMUSG00000068747 | 245.2498038812 | 531.97094331849 | -1.1171 | 4.0338e-26 | 2.9347e-24 | Sort1 |

| | | | | | | |
|---|---|---|---|---|---|---|
| ENSMUSG00000030313 | 628.264288177177 | 1118.01573037687 | -0.8315 | 5.0124e-26 | 3.6352e-24 | Dennd5b |
| ENSMUSG00000040033 | 3674.38591647911 | 2251.42809122146 | 0.70666 | 5.1185e-26 | 3.7005e-24 | Stat2 |
| ENSMUSG00000029070 | 1021.18551800977 | 568.941874890355 | 0.84389 | 5.1713e-26 | 3.727e-24 | Mxra8 |
| ENSMUSG00000036606 | 9887.71696088441 | 5983.4280579045 | 0.72467 | 5.5224e-26 | 3.9677e-24 | Plxnb2 |
| ENSMUSG00000031133 | 42.7528097062698 | 167.033957779383 | -1.9661 | 7.8832e-26 | 5.6464e-24 | Arhgef6 |
| ENSMUSG00000027993 | 560.71312804223 | 282.322387958801 | 0.98992 | 8.2097e-26 | 5.8621e-24 | Trim2 |
| ENSMUSG00000022598 | 35.6162724560469 | 0.234413562335488 | 7.2473 | 8.4934e-26 | 6.046e-24 | Psca |
| ENSMUSG00000020808 | 2023.56584680507 | 3301.51284261257 | -0.70623 | 9.8663e-26 | 7.0018e-24 | Fam64a |
| ENSMUSG00000040010 | 3765.36545861123 | 1891.27588638848 | 0.99343 | 1.1299e-25 | 7.9941e-24 | Slc7a5 |
| ENSMUSG00000031355 | 790.601852422149 | 283.77818349801 | 1.4782 | 1.1933e-25 | 8.4167e-24 | Arhgap6 |
| ENSMUSG00000030126 | 946.238058315422 | 523.382530150708 | 0.85434 | 1.2131e-25 | 8.5303e-24 | Tmcc1 |
| ENSMUSG00000024501 | 6985.1682542741 | 4078.88436121251 | 0.77612 | 1.2736e-25 | 8.9285e-24 | Dpysl3 |
| ENSMUSG00000026833 | 262.395378866637 | 893.740922597663 | -1.7681 | 1.513e-25 | 1.0575e-23 | Olfm1 |
| ENSMUSG00000060791 | 300.743311549572 | 736.459694556457 | -1.2921 | 1.5328e-25 | 1.0681e-23 | Gmfg |
| ENSMUSG00000031075 | 1035.72688475455 | 453.958159701488 | 1.19 | 1.5726e-25 | 1.0926e-23 | Ano1 |
| ENSMUSG00000021285 | 1368.43381797859 | 2271.95685969962 | -0.73141 | 1.8955e-25 | 1.3129e-23 | Ppp1r13b |
| ENSMUSG00000086145 | 7.12012398317564 | 63.6072782483476 | -3.1592 | 2.1484e-25 | 1.4837e-23 | -//- |
| ENSMUSG00000072437 | 193.955481952095 | 411.917618921036 | -1.0866 | 4.239e-25 | 2.9187e-23 | Nanos1 |
| ENSMUSG00000044167 | 1052.39742727042 | 1767.36365107554 | -0.74792 | 4.409e-25 | 3.0268e-23 | Foxo1 |
| ENSMUSG00000030403 | 2742.33106086425 | 4387.52236126722 | -0.678 | 4.868e-25 | 3.332e-23 | Vasp |
| ENSMUSG00000054203 | 275.027420576173 | 54.121377146035 | 2.3453 | 5.6332e-25 | 3.8444e-23 | Ifi205 |
| ENSMUSG00000045679 | 1593.50714170261 | 941.618864515908 | 0.75899 | 6.3115e-25 | 4.2946e-23 | Pqlc3 |
| ENSMUSG00000049791 | 619.275534104047 | 1089.90393060733 | -0.81555 | 6.3767e-25 | 4.3262e-23 | Fzd4 |
| ENSMUSG00000025510 | 6604.10930046691 | 4179.03175536863 | 0.6602 | 6.8605e-25 | 4.6409e-23 | Cd151 |
| ENSMUSG00000037362 | 1166.42811107011 | 580.506009225862 | 1.0067 | 6.9875e-25 | 4.713e-23 | Nov |
| ENSMUSG00000047205 | 921.416398374102 | 515.115246009567 | 0.83896 | 7.3822e-25 | 4.9648e-23 | Dusp18 |
| ENSMUSG00000000278 | 6940.40684926789 | 4404.14183736527 | 0.65616 | 8.2152e-25 | 5.509e-23 | Scpep1 |
| ENSMUSG00000032487 | 6772.34763391811 | 2099.43189753895 | 1.6897 | 9.1291e-25 | 6.1041e-23 | Ptgs2 |
| ENSMUSG00000021750 | 471.393149887933 | 189.391546772948 | 1.3156 | 9.3935e-25 | 6.2628e-23 | Fam107a |
| ENSMUSG00000018395 | 1184.55345674297 | 1983.44259999541 | -0.74366 | 1.084e-24 | 7.2065e-23 | Kif3a |
| ENSMUSG00000020911 | 102.510457329747 | 1.43201481812412 | 6.1616 | 1.1029e-24 | 7.3114e-23 | Krt19 |
| ENSMUSG00000026896 | 1788.04951531457 | 911.981944666336 | 0.97131 | 1.1181e-24 | 7.3906e-23 | Ifih1 |
| ENSMUSG00000026483 | 105.792497686634 | 24.2589907293334 | 2.1246 | 1.1288e-24 | 7.4404e-23 | Fam129a |
| ENSMUSG00000030002 | 2820.54145795621 | 1736.05216238241 | 0.70016 | 1.1822e-24 | 7.7698e-23 | Dusp11 |
| ENSMUSG00000069662 | 10581.3877159823 | 6754.05577948067 | 0.6477 | 1.2821e-24 | 8.4028e-23 | Marcks |
| ENSMUSG00000026750 | 3041.09218425685 | 1877.66922273173 | 0.69565 | 1.4917e-24 | 9.7486e-23 | Psmb7 |
| ENSMUSG00000025314 | 314.371618820435 | 83.4308547868021 | 1.9138 | 1.5062e-24 | 9.8157e-23 | Ptprj |
| ENSMUSG00000073016 | 322.096958969174 | 616.671282233695 | -0.93701 | 1.5853e-24 | 1.0302e-22 | Uprt |
| ENSMUSG00000052373 | 108.967687798279 | 264.976530749236 | -1.282 | 1.7325e-24 | 1.1227e-22 | Mpp3 |
| ENSMUSG00000020986 | 1242.61988552357 | 2049.97677150463 | -0.72222 | 1.7978e-24 | 1.1618e-22 | Sec23a |
| ENSMUSG00000041729 | 118.369427872623 | 282.449767840772 | -1.2547 | 1.9027e-24 | 1.2262e-22 | Coro2b |
| ENSMUSG00000022964 | 456.805971740446 | 827.248353710577 | -0.85674 | 1.9823e-24 | 1.2739e-22 | Tmem50b |
| ENSMUSG00000029461 | 714.168594337052 | 1231.13941969613 | -0.78566 | 2.0478e-24 | 1.3124e-22 | Fam168a |
| ENSMUSG00000039197 | 2203.24050762166 | 1065.48994778665 | 1.0481 | 2.3621e-24 | 1.5096e-22 | Adk |
| ENSMUSG00000044042 | 1695.1245782019 | 1017.06322361527 | 0.73698 | 2.7706e-24 | 1.7658e-22 | Fmn1 |
| ENSMUSG00000027763 | 6804.73799228603 | 4336.86220361815 | 0.64989 | 2.8746e-24 | 1.8271e-22 | Mbnl1 |
| ENSMUSG00000021482 | 197.074680216746 | 441.85641935162 | -1.1648 | 3.8966e-24 | 2.4686e-22 | Aaed1 |
| ENSMUSG00000068245 | 2782.42875401174 | 1596.44478076072 | 0.80148 | 3.9054e-24 | 2.4686e-22 | Phf11d |
| ENSMUSG00000022906 | 1749.24579858979 | 1049.29357355019 | 0.73731 | 3.954e-24 | 2.4925e-22 | Parp9 |
| ENSMUSG00000068876 | 193.495365898991 | 21.8099248254821 | 3.1492 | 4.3182e-24 | 2.7147e-22 | Cgn |
| ENSMUSG00000074825 | 936.068492802107 | 1571.61706637777 | -0.74756 | 4.6297e-24 | 2.9027e-22 | Itpripl1 |
| ENSMUSG00000027660 | 557.366122802763 | 979.707529637621 | -0.81373 | 5.3729e-24 | 3.3595e-22 | Skil |
| ENSMUSG00000019997 | 1642.95260733391 | 4779.2874650396 | -1.5405 | 5.4108e-24 | 3.3659e-22 | Ctgf |
| ENSMUSG00000031176 | 3553.56965834999 | 2224.76586780264 | 0.67562 | 5.4122e-24 | 3.3659e-22 | Dynlt3 |
| ENSMUSG00000062960 | 13024.6361905367 | 20780.3958641941 | -0.67398 | 6.838e-24 | 4.2412e-22 | Kdr |
| ENSMUSG00000020116 | 1513.62481569495 | 899.933545377028 | 0.75012 | 7.6277e-24 | 4.7184e-22 | Pno1 |
| ENSMUSG00000045838 | 3378.91012603417 | 2088.60494733663 | 0.69402 | 8.5019e-24 | 5.2451e-22 | A430105I19Rik |
| ENSMUSG00000022090 | 184.546385647804 | 65.1516534847241 | 1.5021 | 9.3558e-24 | 5.7566e-22 | Pdlim2 |
| ENSMUSG00000024885 | 421.175332023912 | 808.329346490394 | -0.94052 | 1.1275e-23 | 6.9189e-22 | Aldh3b1 |
| ENSMUSG00000026305 | 2909.75430101618 | 1810.00493619586 | 0.6849 | 1.2848e-23 | 7.8635e-22 | Lrrfip1 |
| ENSMUSG00000021701 | 3007.81607889622 | 1603.56184165097 | 0.90744 | 1.2931e-23 | 7.8937e-22 | Plk2 |
| ENSMUSG00000026478 | 22020.3545374023 | 13417.1680833891 | 0.71476 | 1.307e-23 | 7.9572e-22 | Lamc1 |
| ENSMUSG00000012443 | 4045.20728767244 | 6322.5529085432 | -0.64429 | 1.3568e-23 | 8.2389e-22 | Kif11 |
| ENSMUSG00000097252 | 116.549075673168 | 30.5629622434848 | 1.9311 | 1.7636e-23 | 1.0681e-21 | -//- |
| ENSMUSG00000073119 | 58.0253304802357 | 239.489378650603 | -2.0452 | 1.8262e-23 | 1.1031e-21 | Speer4a |
| ENSMUSG00000025507 | 555.889009653955 | 972.741620532839 | -0.80726 | 1.8838e-23 | 1.135e-21 | Pidd1 |
| ENSMUSG00000106627 | 415.500784552705 | 1585.99816074853 | -1.9325 | 2.2101e-23 | 1.3281e-21 | RP23-56M18.6 |
| ENSMUSG00000026074 | 4884.17488639705 | 7633.75495881716 | -0.64428 | 2.2803e-23 | 1.3667e-21 | Map4k4 |
| ENSMUSG00000002997 | 1569.48481612091 | 724.149792312498 | 1.1159 | 2.3254e-23 | 1.383e-21 | Prkar2b |
| ENSMUSG00000018567 | 5254.46458638498 | 3360.65151700483 | 0.6448 | 2.3254e-23 | 1.383e-21 | Gabarap |
| ENSMUSG00000051517 | 437.876982893228 | 789.646333762819 | -0.85068 | 2.3222e-23 | 1.383e-21 | Arhgef39 |
| ENSMUSG00000024640 | 10519.8492631794 | 5812.32077911749 | 0.85593 | 2.3956e-23 | 1.4211e-21 | Psat1 |
| ENSMUSG00000039886 | 2166.55627650145 | 1332.1975287485 | 0.7016 | 2.7874e-23 | 1.6493e-21 | Tmem120a |
| ENSMUSG00000021477 | 8706.83825147979 | 13397.0768153949 | -0.6217 | 2.9615e-23 | 1.7478e-21 | Ctsl |
| ENSMUSG00000028382 | 2115.35684730712 | 3668.88280924106 | -0.79444 | 3.1956e-23 | 1.8812e-21 | Ptbp3 |
| ENSMUSG00000060477 | 212.757393369159 | 431.146098421542 | -1.019 | 3.5458e-23 | 2.082e-21 | Irak2 |
| ENSMUSG00000038295 | 776.174957470685 | 1674.97344646685 | -1.1097 | 3.611e-23 | 2.1149e-21 | Atg9b |
| ENSMUSG00000059534 | 2732.55751683857 | 1709.7191036572 | 0.67649 | 3.8484e-23 | 2.2483e-21 | Uqcr10 |
| ENSMUSG00000030539 | 622.339548420142 | 331.469267481867 | 0.90883 | 4.1281e-23 | 2.4057e-21 | Sema4b |
| ENSMUSG00000038203 | 105.999068201379 | 254.768033170537 | -1.2651 | 4.4134e-23 | 2.5654e-21 | Hoxa13 |
| ENSMUSG00000022150 | 8856.86850515508 | 5742.1496496137 | 0.62521 | 4.6989e-23 | 2.7246e-21 | Dab2 |
| ENSMUSG00000030057 | 17609.7528643641 | 11487.1546527425 | 0.61635 | 4.782e-23 | 2.7658e-21 | Cnbp |
| ENSMUSG00000026626 | 3216.97666769945 | 5018.70969509056 | -0.64161 | 6.0439e-23 | 3.4869e-21 | Ppp2r5a |
| ENSMUSG00000042807 | 220.460598180378 | 443.173413090685 | -1.0074 | 6.193e-23 | 3.5574e-21 | Hecw2 |
| ENSMUSG00000097184 | 32.0664617718409 | 120.057549538446 | -1.9046 | 6.1967e-23 | 3.5574e-21 | -//- |
| ENSMUSG00000034993 | 13860.8794866763 | 8743.93854065514 | 0.66466 | 7.166e-23 | 4.1036e-21 | Vat1 |
| ENSMUSG00000022265 | 567.531599187192 | 984.552954616404 | -0.79477 | 7.8788e-23 | 4.5007e-21 | Ank |
| ENSMUSG00000032349 | 2148.2103271088 | 1308.85499604025 | 0.71483 | 9.256e-23 | 5.2743e-21 | Elovl5 |
| ENSMUSG00000044573 | 2119.81573177731 | 1309.39058056572 | 0.69504 | 9.3826e-23 | 5.3333e-21 | Acp1 |
| ENSMUSG00000070436 | 24707.1045377945 | 16222.1609121288 | 0.60696 | 9.6966e-23 | 5.4983e-21 | Serpinh1 |
| ENSMUSG00000023963 | 161.007910739506 | 342.769168227468 | -1.0901 | 1.0099e-22 | 5.7123e-21 | Cyp39a1 |
| ENSMUSG00000048807 | 1327.91840662957 | 767.143844753691 | 0.7916 | 1.0165e-22 | 5.736e-21 | Slc35e4 |
| ENSMUSG00000032300 | 1567.07387980536 | 2510.07367293772 | -0.67966 | 1.1148e-22 | 6.2751e-21 | 1700017B05Rik |
| ENSMUSG00000030605 | 6109.39718338514 | 3945.87477159807 | 0.63068 | 1.1249e-22 | 6.3169e-21 | Mfge8 |
| ENSMUSG00000035093 | 3533.47888713318 | 5487.74152024282 | -0.63512 | 1.1899e-22 | 6.6653e-21 | Secisbp2l |
| ENSMUSG00000041633 | 4260.65809542335 | 9741.39203255407 | -1.1931 | 1.2202e-22 | 6.8189e-21 | Kctd12b |
| ENSMUSG00000000093 | 314.671850919672 | 1281.71916392147 | -2.0262 | 1.5342e-22 | 8.5529e-21 | Tbx2 |
| ENSMUSG00000043008 | 704.169406834406 | 1353.55891681507 | -0.94276 | 1.6245e-22 | 9.0342e-21 | Klhl6 |
| ENSMUSG00000026814 | 5392.03182903034 | 8511.43058023024 | -0.65857 | 1.6412e-22 | 9.1055e-21 | Eng |
| ENSMUSG00000079037 | 5177.06867908764 | 3331.75492259577 | 0.63585 | 1.9775e-22 | 1.0945e-20 | Prnp |
| ENSMUSG00000031198 | 2019.42903309759 | 1248.7857190041 | 0.69342 | 2.0416e-22 | 1.1273e-20 | Fundc2 |
| ENSMUSG00000001847 | 10520.6465571949 | 6885.8162743094 | 0.61152 | 2.1109e-22 | 1.1627e-20 | Rac1 |
| ENSMUSG00000067297 | 70.7717348726539 | 11.6376306214896 | 2.6044 | 2.484e-22 | 1.365e-20 | Ifit1bl2 |
| ENSMUSG00000030717 | 1107.36414714166 | 573.530104572611 | 0.94919 | 2.5511e-22 | 1.3986e-20 | Nupr1 |
| ENSMUSG00000028556 | 1288.81343808514 | 586.657681342041 | 1.1355 | 2.937e-22 | 1.6063e-20 | Dock7 |
| ENSMUSG00000032531 | 2277.88993268375 | 4642.67692266013 | -1.0273 | 3.7171e-22 | 2.0282e-20 | Amotl2 |
| ENSMUSG00000022895 | 4003.92654933865 | 7063.40865499672 | -0.81895 | 3.8094e-22 | 2.0737e-20 | Ets2 |
| ENSMUSG00000047344 | 176.187436647831 | 31.6460297082001 | 2.477 | 4.0382e-22 | 2.193e-20 | Lancl3 |
| ENSMUSG00000027955 | 3034.99919862021 | 1609.26000726451 | 0.9153 | 4.0549e-22 | 2.197e-20 | Fam198b |
| ENSMUSG00000031434 | 1351.37224695019 | 2975.75528645347 | -1.1388 | 4.1403e-22 | 2.238e-20 | Morc4 |
| ENSMUSG00000020181 | 337.464621687517 | 134.110974780933 | 1.3313 | 4.381e-22 | 2.3626e-20 | Nav3 |
| ENSMUSG00000022421 | 1293.52187962145 | 2321.35868309347 | -0.84366 | 4.4275e-22 | 2.3821e-20 | Nptxr |
| ENSMUSG00000041736 | 5412.85620551501 | 3507.92335323903 | 0.62577 | 4.8358e-22 | 2.5957e-20 | Tspo |
| ENSMUSG00000052056 | 791.163901932325 | 1316.00956280968 | -0.73412 | 4.8933e-22 | 2.6205e-20 | Zfp217 |
| ENSMUSG00000037060 | 4604.88223080758 | 2973.7731030136 | 0.63087 | 4.9323e-22 | 2.6353e-20 | Prkcdbp |
| ENSMUSG00000056204 | 2445.44250750601 | 1542.14076601266 | 0.66516 | 4.9561e-22 | 2.6419e-20 | Pgpep1 |
| ENSMUSG00000037606 | 1631.41586954536 | 1004.38702473646 | 0.69981 | 5.3041e-22 | 2.8209e-20 | Osbpl5 |
| ENSMUSG00000002227 | 2396.71034644757 | 1510.53517430576 | 0.666 | 5.9271e-22 | 3.145e-20 | Mov10 |
| ENSMUSG00000041890 | 3299.13258377494 | 5102.06215835035 | -0.62899 | 6.0974e-22 | 3.228e-20 | Git2 |
| ENSMUSG00000061119 | 907.106350678947 | 523.886642133638 | 0.79202 | 7.0903e-22 | 3.7451e-20 | Prcp |
| ENSMUSG00000039043 | 940.035702587552 | 548.344902924889 | 0.77763 | 7.3506e-22 | 3.8737e-20 | Arpin |
| ENSMUSG00000070047 | 2864.62542232925 | 1190.76767573845 | 1.2665 | 7.9779e-22 | 4.1947e-20 | Fat1 |
| ENSMUSG00000041324 | 280.455211413348 | 528.167012239912 | -0.91322 | 8.4293e-22 | 4.422e-20 | Inhba |
| ENSMUSG00000071454 | 495.860613084491 | 862.840409705228 | -0.79916 | 9.4393e-22 | 4.9407e-20 | Dtnb |
| ENSMUSG00000038855 | 1279.271739588 | 2249.24923772515 | -0.81412 | 9.7143e-22 | 5.0732e-20 | Itpkb |
| ENSMUSG00000021996 | 20310.5102305859 | 13435.924691463 | 0.59613 | 1.0035e-21 | 5.2291e-20 | Esd |
| ENSMUSG00000073116 | 26.9353248228406 | 119.875257674549 | -2.154 | 1.0173e-21 | 5.2888e-20 | Gm10471 |
| ENSMUSG00000026655 | 5658.02100924151 | 8607.7729861045 | -0.60534 | 1.073e-21 | 5.5659e-20 | Fam107b |
| ENSMUSG00000024168 | 198.085460186879 | 647.961524031342 | -1.7098 | 1.1644e-21 | 6.0264e-20 | Tmem204 |
| ENSMUSG00000017314 | 367.04144221802 | 700.985928513812 | -0.93344 | 1.178e-21 | 6.0831e-20 | Mpp2 |
| ENSMUSG00000047735 | 4739.6656514408 | 2900.21439526971 | 0.70863 | 1.2113e-21 | 6.2412e-20 | Samd9l |
| ENSMUSG00000021180 | 191.885277432859 | 385.903427733748 | -1.008 | 1.5923e-21 | 8.1862e-20 | Rps6ka5 |
| ENSMUSG00000062609 | 89.4584888475893 | 218.554895322018 | -1.2887 | 1.607e-21 | 8.2436e-20 | Kcnj15 |
| ENSMUSG00000005611 | 435.063216307696 | 763.323998465592 | -0.81107 | 1.6124e-21 | 8.2526e-20 | Mrvi1 |
| ENSMUSG00000001768 | 586.754176394769 | 1304.23704131721 | -1.1524 | 1.6976e-21 | 8.6699e-20 | Rin2 |
| ENSMUSG00000045930 | 320.323349902834 | 652.783227786377 | -1.0271 | 1.8246e-21 | 9.2976e-20 | Clec14a |
| ENSMUSG00000016128 | 1131.29530087758 | 676.330811707667 | 0.74217 | 2.0584e-21 | 1.0466e-19 | Stard13 |
| ENSMUSG00000018648 | 324.157386982742 | 152.152497168214 | 1.0912 | 2.157e-21 | 1.0944e-19 | Dusp14 |
| ENSMUSG00000056144 | 695.449620129815 | 388.628102931819 | 0.83956 | 2.1948e-21 | 1.1111e-19 | Trim34a |
| ENSMUSG00000068220 | 29242.3651283734 | 19447.3173352356 | 0.58849 | 2.2867e-21 | 1.1551e-19 | Lgals1 |
| ENSMUSG00000069833 | 20817.8622354072 | 10622.4993351875 | 0.9707 | 2.4964e-21 | 1.2582e-19 | Ahnak |
| ENSMUSG00000022013 | 795.22783308748 | 453.67195808842 | 0.80972 | 2.6827e-21 | 1.3492e-19 | Dnajc15 |
| ENSMUSG00000033365 | 4297.84565245235 | 2684.0956890312 | 0.67918 | 2.7416e-21 | 1.3759e-19 | Ipo13 |
| ENSMUSG00000030207 | 513.040909924013 | 880.335361607839 | -0.77898 | 2.8955e-21 | 1.45e-19 | 8430419L09Rik |
| ENSMUSG00000018042 | 7777.86307512858 | 5128.44144786573 | 0.60085 | 3.1728e-21 | 1.5854e-19 | Cyb5r3 |
| ENSMUSG00000029798 | 1353.20359211367 | 708.467904279183 | 0.9336 | 3.422e-21 | 1.7062e-19 | Herc6 |
| ENSMUSG00000037005 | 12.1928221959891 | 69.4422460333888 | -2.5098 | 4.5241e-21 | 2.2509e-19 | Xpnpep2 |
| ENSMUSG00000019888 | 66.2790609712638 | 11.2115457817661 | 2.5636 | 5.0579e-21 | 2.5057e-19 | Mgat4c |
| ENSMUSG00000024269 | 451.190871150438 | 212.563285207244 | 1.0858 | 5.0525e-21 | 2.5057e-19 | Tpgs2 |
| ENSMUSG00000024899 | 209.857981525184 | 42.7525328079427 | 2.2953 | 6.0333e-21 | 2.9825e-19 | Papss2 |
| ENSMUSG00000016496 | 660.94998346064 | 272.568991090196 | 1.2779 | 6.2389e-21 | 3.0775e-19 | Cd274 |
| ENSMUSG00000050919 | 426.577922015412 | 1078.63870341918 | -1.3383 | 6.5056e-21 | 3.2023e-19 | Zfp366 |
| ENSMUSG00000022419 | 445.135345160001 | 176.582230408215 | 1.3339 | 6.7319e-21 | 3.3067e-19 | Deptor |
| ENSMUSG00000022146 | 270.194976486279 | 121.622802810892 | 1.1516 | 7.754e-21 | 3.8006e-19 | Osmr |
| ENSMUSG00000024781 | 3215.41103413306 | 2043.2426513778 | 0.65414 | 8.9404e-21 | 4.3729e-19 | Lipa |
| ENSMUSG00000039318 | 1869.54011173617 | 1179.91843555198 | 0.664 | 9.8945e-21 | 4.8293e-19 | Rab3gap2 |
| ENSMUSG00000064120 | 1488.94700154049 | 923.329412617852 | 0.68938 | 1.0411e-20 | 5.0705e-19 | Mocs1 |
| ENSMUSG00000053477 | 3117.80942904389 | 4758.53055812219 | -0.60998 | 1.1726e-20 | 5.6992e-19 | Tcf4 |
| ENSMUSG00000034906 | 1928.51722948384 | 2985.51868326653 | -0.63049 | 1.3527e-20 | 6.561e-19 | Ncaph |
| ENSMUSG00000032966 | 32044.7202043725 | 47623.1525271984 | -0.57158 | 1.3931e-20 | 6.7424e-19 | Fkbp1a |
| ENSMUSG00000015289 | 541.231794559 | 294.691371594382 | 0.87704 | 1.409e-20 | 6.8055e-19 | Lage3 |
| ENSMUSG00000021485 | 306.67505222697 | 564.494839123739 | -0.88025 | 1.474e-20 | 7.1043e-19 | Mxd3 |
| ENSMUSG00000004791 | 1359.73062560548 | 3306.78093818372 | -1.2821 | 1.5043e-20 | 7.2351e-19 | Pgf |
| ENSMUSG00000107768 | 1083.5233347551 | 466.145197533405 | 1.2169 | 1.7486e-20 | 8.3927e-19 | -//- |
| ENSMUSG00000025473 | 409.931715574187 | 210.58058966063 | 0.96101 | 1.7966e-20 | 8.6054e-19 | Adam8 |
| ENSMUSG00000021281 | 4853.10652864543 | 3192.6770952074 | 0.60414 | 1.8051e-20 | 8.6283e-19 | Tnfaip2 |
| ENSMUSG00000040265 | 125.540407205372 | 338.582097234237 | -1.4314 | 1.9504e-20 | 9.3037e-19 | Dnm3 |
| ENSMUSG00000027254 | 337.15167536641 | 145.175696796195 | 1.2156 | 2.1482e-20 | 1.0226e-18 | Map1a |
| ENSMUSG00000021226 | 363.659739066321 | 181.919924871681 | 0.99929 | 2.1796e-20 | 1.0354e-18 | Acot2 |
| ENSMUSG00000025408 | 1190.51043875672 | 570.911973234883 | 1.0602 | 2.3021e-20 | 1.0914e-18 | Ddit3 |
| ENSMUSG00000042216 | 238.937795986272 | 518.455909876214 | -1.1176 | 2.3397e-20 | 1.1069e-18 | Sgsm1 |
| ENSMUSG00000029722 | 1584.77062126583 | 2471.34610784313 | -0.64102 | 2.6932e-20 | 1.2716e-18 | Agfg2 |
| ENSMUSG00000016239 | 298.098988273423 | 101.044051361935 | 1.5608 | 2.8596e-20 | 1.3474e-18 | Lonrf3 |
| ENSMUSG00000001018 | 894.647957791136 | 530.588919254557 | 0.75373 | 2.9973e-20 | 1.4094e-18 | Snapin |
| ENSMUSG00000033904 | 1168.78327143161 | 1847.9456240623 | -0.66091 | 3.1046e-20 | 1.4569e-18 | Ccp110 |
| ENSMUSG00000024014 | 1031.78699597968 | 2181.21836412184 | -1.08 | 3.4703e-20 | 1.6252e-18 | Pim1 |
| ENSMUSG00000078349 | 419.671157922318 | 217.654320376567 | 0.94722 | 3.7324e-20 | 1.7444e-18 | AW011738 |
| ENSMUSG00000040483 | 2736.62760397043 | 1764.95958783784 | 0.63276 | 3.7943e-20 | 1.7698e-18 | Xaf1 |
| ENSMUSG00000001995 | 1003.33439786165 | 1988.59717723469 | -0.98695 | 4.0521e-20 | 1.8862e-18 | Sipa1l2 |
| ENSMUSG00000031924 | 4005.62173462314 | 7112.53622033551 | -0.82834 | 4.1262e-20 | 1.9138e-18 | Cyb5b |
| ENSMUSG00000059027 | 141.713273530647 | 298.001875151057 | -1.0723 | 4.1278e-20 | 1.9138e-18 | -//- |
| ENSMUSG00000037594 | 311.669667043065 | 149.963619078161 | 1.0554 | 4.6987e-20 | 2.1741e-18 | BC022687 |

| Term | Overlap | P-value | Adjusted P-value | Genes |
|---|---|---|---|---|
| regulation of cell migration (GO:0030334) | 34/316 | 4.168055490873482E-13 | 1.0586860946818643E-9 | CD151;CSF1;SEMA3C;SEMA3A;PDGFB;PDGFA;PTPRJ;SEMA3E;ADARB1; FGF1;PAK1;FAM110C;PDGFD;DPYSL3;LMNA;KDR;CYP1B1;RAC1;SRGAP3; ARPIN;EDN1;JAG1;ANXA1;ARHGEF39;SEMA4B;TPM1;BST2; TIAM1;DAB2;NAV3;PLXNB2;AMOTL2;SGK1;ENG |
| extracellular matrix organization (GO:0030198) | 21/229 | 2.1678849334941394E-7 | 2.753213865537557E-4 | VCAM1;ITGA1;PDGFB;PDGFA;LOXL4;LAMC1;COL5A1;CTSL;KDR; SERPINH1;TIMP2;CAPN2;COL4A6;CYP1B1;COL4A5; ITGA7;ADAM8;TIMP1;LCP1;CD44;DDR2 |
| cellular response to type I interferon (GO:0071357) | 11/65 | 4.356552813789541E-7 | 2.7664110367563584E-4 | BST2;RSAD2;IFI27;STAT1;OAS3;STAT2;IRF7;ISG15;IFIT1;XAF1;IFIT3 |
| type I interferon signaling pathway (GO:0060337) | 11/65 | 4.356552813789541E-7 | 2.7664110367563584E-4 | BST2;RSAD2;IFI27;STAT1;OAS3;STAT2;IRF7;ISG15;IFIT1;XAF1;IFIT3 |
| response to cytokine (GO:0034097) | 15/138 | 1.418875327663055E-6 | 6.344663285074403E-4 | CD274;STAT1;ISG15;AFF3;OSMR;SELE;BST2;IRAK2;CASP3;DPYSL3;BCL2; TIMP2;ADAM9;TIMP1;XAF1 |
| negative regulation of cell migration (GO:0030336) | 14/121 | 1.498739358678993E-6 | 6.344663285074403E-4 | JAG1;TPM1;PTPRJ;ADARB1;PTPRG;BST2;NAV3;DPYSL3;CYP1B1;HMOX1; SRGAP3;TIMP1;ARPIN;ENG |
| positive regulation of cell motility (GO:2000147) | 17/179 | 1.9113769513245516E-6 | 6.93556779480623E-4 | EDN1;CD151;CSF1;SEMA3C;ARHGEF39;SEMA3A;SEMA4B;PDGFB;PDGFA; SEMAE;FGF1;TIAM1;DAB2;PAK1;FAM110C;PDGFD;KDR |
| positive regulation of fibroblast proliferation (GO:0048146) | 07/28 | 3.693471784353107E-6 | 0.0011726772915321114 | CDK6;PDGFB;S100A6;PDGFA;AQP1;EREG;DDR2 |
| positive regulation of cell proliferation (GO:0008284) | 27/424 | 5.994210083927416E-6 | 0.001570063310885789 | CSF1;PDGFB;PDGFA;LAMC1;FGF1;AQP1;EFNB2;ESM1;PAK1;CCND2; PDGFD;PIM1;KDR;HMOX1;TIMP1;EDN1;STAT1;CNBP;OSMR;PGF; TBX2;EREG;TIAM1;BST1;CDK6;S100A6;DDR2 |
| cytokine-mediated signaling pathway (GO:0019221) | 35/633 | 6.1813516176605865E-6 | 0.001570063310885789 | CSF1;MAOA;PDGFB;PTGS2;IFIT1;FOXO1;IFIT3;PSMD8;PSMB7;RPS6KA5; IRAK2;CASP3;PIM1;HMOX1;TNFRSF8;TIMP1;ANXA1; VCAM1;ANXA2;RSAD2;STAT1;STAT2;ISG15;OSMR; EREG;BST2;IFI27;OAS3;KIT;BCL2;IRF7;LCP1;XAF1;SQSTM1;CD44 |

**Primary Antibodies**

| Antibody | Full name | Company | Catalog number | Dilution |
|---|---|---|---|---|
| Isolectin B4 | Biotinylated griffonia simplicifolia lectin 1 | VECTOR | B-1205 | IF 1:300 |
| GFP | Chk pAb to GFP | Abcam | ab13970 | IF 1:200 |
| Phalloidin | Texas Red-X-Phalloidin | Life Technologies | T7471 | IF 1:300 for cell; 1:200 for retina |
| Phalloidin | Phalloidin-Atto 647N | Sigma | 65906-10nmol | IF 1:300 for cell; 1:200 for retina |
| TO-PRO-3 | TO-PRO- iodide (642/661) | Life Technologies | T3605 | IF 1:1000 |
| β-actin | Ms mAb to Actin | Abcam | ab3284 | WB-1:2000 |
| VE-cadherin | Purified anti human CD144 (BV9) | Bio-legend | 348502 | IF 1:300 |
| VE-cadherin | Purified rat anti mouse CD144 (11d4.1) | BD | 555289 | IF 1:300 |
| VE-cadherin | Rabbit polyclonal VE Cadherin antibody | Abcam | 33168 | WB 1:500 |
| Lamin A/C | Monocolonal Anti-Lamin A/C ab produced in mouse | Sigma | SAB4200236 | WB 1:500 |
| Lamin B | Rb pAb to LaminB1 | Abcam | ab16048 | WB 1:500 |
| SUN2 | Rb mAb to SUN2 | Abcam | ab124916 | WB 1:500 |
| GAPDH | Monocolonal Anti-GAPDH ab produced in mouse | Sigma | G8795 | WB 1:3000 |
| CD31 (Pecam-1) | Purified rat anti mouse CD31 | BD | 553370 | IF 1:300 |
| β-catenin | Purified mouse anti β-catenin | BD | 810154 | WB 1:500 |
| CD45 | Mouse CD45 Antibody | R&D systems | AF-114 | IF 1:300 |
| AmotL2 | Angiomotin like 2 | Innovagen, Lund, Sweden | Purified from rabbit serum | WB-1:500, IP-1:1:200 (v/v) IF/IHC 1:100 |

**Secondary Antibodies**

| | Secondary Antibody | Conjugation | Catalog number | Company | Dilution |
|---|---|---|---|---|---|
| For WB | ECL® Anti-rabbit IgG | HRP linked whole antibody from donkey | NA934V | GE Healthcare | 1:10000 |
| | ECL® Anti-mouse IgG | HRP linked whole antibody from donkey | NA931V | GE Healthcare | 1:10000 |
| | ECL® Anti-rat IgG | HRP linked whole antibody from donkey | NA935V | GE Healthcare | 1:10000 |
| For IF staining | goat anti-Mouse IgG (H+L) | Alexa Fluor® 405 conjugate | A31553 | LifeTechnologies | 1:500 |
| | goat anti-Rat IgG (H+L) | Alexa Fluor® 488 conjugate | A11006 | LifeTechnologies | 1:500 |
| | donkey anti-Sheep IgG (H+L) | Alexa Fluor® 488 conjugate | A11015 | LifeTechnologies | 1:500 |
| | goat anti-Chicken IgG (H+L) | Alexa Fluor® 488 conjugate | A11039 | LifeTechnologies | 1:500 |
| | donkey anti-Goat IgG (H+L) | Alexa Fluor® 488 conjugate | A11055 | LifeTechnologies | 1:500 |
| | chicken anti-Rabbit IgG (H+L) | Alexa Fluor® 488 conjugate | A21441 | LifeTechnologies | 1:500 |
| | donkey anti-Mouse IgG (H+L) | Alexa Fluor® 555 conjugate | A31570 | LifeTechnologies | 1:500 |
| | donkey anti-Rabbit IgG (H+L) | Alexa Fluor® 555 conjugate | A31572 | LifeTechnologies | 1:500 |
| | goat anti-Mouse IgG (H+L) | Alexa Fluor® 594 conjugate | A11005 | LifeTechnologies | 1:500 |
| | goat anti-Rat IgG (H+L) | Alexa Fluor® 594 conjugate | A11007 | LifeTechnologies | 1:500 |
| | goat anti-Rabbit IgG (H+L) | Alexa Fluor® 594 conjugate | A11037 | LifeTechnologies | 1:500 |
| | goat anti-Rat IgG (H+L) | Alexa Fluor® 633 conjugate | A21094 | LifeTechnologies | 1:500 |
| | donkey anti-Mouse IgG (H+L) | Alexa Fluor® 647 conjugate | A31571 | LifeTechnologies | 1:500 |
| | donkey anti-Rabbit IgG (H+L) | Alexa Fluor® 647 conjugate | A31573 | LifeTechnologies | 1:500 |
| | goat anti-Mouse IgG (H+L) | Cy3® | A10521 | LifeTechnologies | 1:500 |
| | goat anti-Rabbit IgG (H+L) | Cy3 | A10520 | LifeTechnologies | 1:500 |
| | Streptavidin | Alexa Fluor® 405 conjugate | S32351 | LifeTechnologies | 1:500 |
| | Streptavidin | Alexa Fluor® 488 conjugate | S32354 | LifeTechnologies | 1:500 |

**Reagents**

| Reagent | Company | Catalog number |
|---|---|---|
| *In vivo* | | |
| Tamoxifen | Sigma | T5648 |
| *In vitro* | | |
| Blebbistatin | Sigma | B0560 |
| Cytoclaxin D | Sigma | C8273 |
| Paraformaldehyde solution 4% in PBS | ChemCruz | sc-281692 |
| Fluoroshield with DAPI | Sigma | F6057 |
| Western Lightning Plus-ECL | PerkinElmer | 203-170071 |
| Protein G Sepharose 4 fast flow | GE Healthcare | 17-0618-01 |
| IgG from rabbit serum | Sigma | I8140 |
| IgG from mouse serum | Sigma | I8765 |
| Polybrene (Hexadimethrine Bromide) | VectorBuilder | provided with lentivirus |
| Polybrene (Hexadimethrine Bromide) | Sigma | H9268-5G |

**Lenti-virus**

| Name | Company | Sequence if it's customized product |
|---|---|---|
| Non-Targeting (scrambled) shRNA Control | Sigma | - |
| human AmotL2 shRNA virus | Sigma | 5’-GCGAGAGAAGGAGGAGCAGATC-3’ |

**TaqMan Probes**

| | |
|---|---|
| Company | Thermo-Fisher |
| Species | Mouse |
| **Gene** | **Taqman reference** |
| TNF | Mm00443258_m1 |
| CD68 | Mm03047343_m1 |
| Cxcl10 | Mm00445235_m1 |
| Icam1 | Mm00516023_m1 |
| Vcam1 | Mm01320970_m1 |
| Cd8a | Mm01188922_m1 |
| Cd4 | Mm01185100_m1 |
| Ccl2 | Mm00441242_m1 |
| Ccl5 | Mm01302427_m1 |
| Cd19 | Mm00515420_m1 |
| IL6 | Mm00446190_m1 |
| HPRT | Mm03024075_m1 |
